# Phylogenomics Illuminates the Evolutionary History of Wild Silkmoths in Space and Time (Lepidoptera: Saturniidae)

**DOI:** 10.1101/2022.03.29.486224

**Authors:** Rodolphe Rougerie, Astrid Cruaud, Pierre Arnal, Liliana Ballesteros-Mejia, Fabien L. Condamine, Thibaud Decaëns, Marianne Elias, Delphine Gey, Paul D. N. Hebert, Ian J. Kitching, Sébastien Lavergne, Carlos Lopez-Vaamonde, Jérôme Murienne, Yves Cuenot, Sabine Nidelet, Jean-Yves Rasplus

## Abstract

Wild silkmoths (Saturniidae) are one of the most emblematic and most studied families of moths. Yet, the absence of a robust phylogenetic framework based on a comprehensive taxonomic sampling impedes our understanding of their evolutionary history. We analyzed 1,024 ultraconserved elements (UCEs) and their flanking regions to infer the relationships among 338 species of Saturniidae representing all described subfamilies, tribes, and genera. We investigated systematic biases in genomic data and performed dating and historical biogeographic analyses to reconstruct the evolutionary history of wild silkmoths in space and time. Using Gene Genealogy Interrogation, we showed that saturation of nucleotide sequence data blurred our understanding of early divergences and first biogeographic events. Our analyses support a Neotropical origin of saturniids, shortly after the Cretaceous-Paleogene extinction event (*ca* 64.0 [stem] - 52.0 [crown] Ma), and two independent colonization events of the Old World during the Eocene, presumably through the Bering Land Bridge. Early divergences strongly shaped the distribution of extant subfamilies as they showed very limited mobility across biogeographical regions, except for Saturniinae, a subfamily now present on all continents but Antarctica. Overall, our results provide a framework for in-depth investigations into the spatial and temporal dynamics of all saturniid lineages and for the integration of their evolutionary history into further global studies of biodiversity and conservation. Rather unexpectedly for a taxonomically well-known family such as Saturniidae, the proper alignment of taxonomic divisions and ranks with our phylogenetic results leads us to propose substantial rearrangements of the family classification, including the description of one new subfamily and two new tribes.

---

Thanks to generations of naturalists, moths in the family Saturniidae – also known as wild silkmoths – are now among the best documented insects, particularly because of their spectacular variation in size and form in both adult and immature stages. The family encompasses some of the largest (e.g., *Coscinocera hercules* and *Attacus* species that reach 30cm wingspan) and most emblematic moths (e.g., the moon moths, *Actias* species and the comet moth, *Argema mittrei*). Wild silkmoths also bear importance to humans for silk production (Kundu et al. 2012, Dong et al. 2015), as a food resource (Fogang Mba et al. 2019), and for health concern (Carrijo-Carvalho and Chudzinski-Tavassi 2007, Battisti et al. 2011, Ciminera et al. 2018). However, despite being one of the best-known families of Lepidoptera, a robust phylogenetic hypothesis of interrelationships among its genera based on extensive sampling is lacking. Saturniids are considered to be poor-dispersers, because of their large and short-lived non-feeding adults that represent an extreme case of capital-breeding organisms (Davis et al. 2016). These moths nonetheless managed to colonize and diversify on almost all continents, but the spatial and temporal dynamics of their evolutionary history remain largely unknown, for neither formal biogeographical analyses nor inferences from time trees have been drawn for the family.

Saturniidae comprise 3,454 species and 180 genera worldwide, classified into eight subfamilies and 11 tribes (Kitching et al. 2018). Five of these subfamilies (Arsenurinae, Ceratocampinae, Cercophaninae, Hemileucinae and Oxyteninae) are restricted to the New World and comprise 60% of the diversity of the family (>2,100 species). The subfamily Agliinae (5 species) occurs throughout the Palearctic region while most species of Salassinae (32 species) are in the Sino-Himalayan region with a few species in the East Palearctic and Oriental regions. Finally, the highly diversified Saturniinae (>1,000 species) are present in all biogeographical regions, except for polar and desertic regions. The family is considered monophyletic based on both morphological (Minet 1994) and molecular evidence (Regier et al. 2008, Breinholt et al. 2018, Hamilton et al. 2019). A few studies have addressed higher-level relationships within the Saturniidae although with sparse taxon and often reduced gene sampling (Regier et al. 2008: 8 species, 7 subfamilies, 5 loci, Barber et al. 2015: 80 species, 8 subfamilies, 6 loci, Hamilton et al. 2019: 16 species, 8 subfamilies, 650 loci). Nevertheless, a consensus has emerged that the Neotropical Oxyteninae is sister to all other Saturniidae while the Cercophaninae is the second earliest-diverging subfamily. The remaining lineages are divided into two large clades, a Neotropical clade that includes three subfamilies (Arsenurinae + Ceratocampinae + Hemileucinae), and an old-world clade composed of Salassinae as sister to Saturniinae. However, the position of the eighth subfamily – the palearctic Agliinae – remains contentious. In recent phylogenomic analyses based on anchored hybrid enrichment (Rubin et al. 2018, Hamilton et al. 2019), Agliinae is recovered as sister to the old-world clade, but with low support, rather than as sister to the new-world clade as found in studies based on Sanger sequencing of a few loci and with low support as well (Regier et al. 2008, Barber et al. 2015, Rubinoff and Doorenweerd 2020). Beside this uncertainty, previous studies have also raised questions regarding the placement of certain taxa, such as the genus *Rhodinia* which has been placed as a member of the tribes Saturniini (Regier et al. 2002) or Attacini (Rubin et al 2018; Chen et al. 2021), or poorly known genera such as the African *Eosia* and *Eochroa* – doubtfully placed, respectively, in Micragonini and Bunaeini tribes within Saturniinae (Oberprieler and Nässig 1994), or *Hirpida* and *Catharisa* of unclear affinities within Hemileucinae (Michener 1952, Lemaire 2002). The latter is the most speciose subfamily with more than 1,600 species in 55 genera, of which just eleven have been included in a formal phylogenetic analysis (Barber et al 2015). Furthermore, although both morphology and molecules have revealed the paraphyly of Urotini with respect to Afrotropical Bunaeini and Micragonini (Regier et al 2008, Rubin et al 2018), incomplete taxon sampling led taxonomists to refrain from revising a classification recognized as unnatural for African Saturniidae (Oberprieler, 1997).

While genome-scale data can often help resolving phylogenetic relationships, they have proven frustrating when addressing difficult nodes within the Saturniidae, presumably because of the lack of signal associated with an old, rapid radiation (Hamilton et al. 2021) and of the increasing probability of observing conflicting signals between markers (Kumar et al. 2012). In particular, although these properties are still rarely thoroughly explored, heterogeneity in base composition and evolutionary rates of taxa and markers (e.g., Boussau et al. 2014, Romiguier et al. 2016, Bossert et al. 2017, Borowiec et al. 2019, Cruaud et al. 2021, Rasplus et al. 2021) as well as saturation (Borowiec 2019, Duchêne et al. 2021) were identified as major causes of analytical bias in phylogenomic analyses. These factors, in concert with limited taxon sampling, may, for instance, explain the poorly resolved placement of Agliinae within the Saturniidae (Hamilton et al. 2019) and constrain our understanding of the early evolutionary history of this family.

In this study, we present a new time-calibrated phylogeny of Saturniidae reconstructed with 1,024 ultraconserved elements (UCEs) and their flanking regions obtained from 338 species representing all described subfamilies, tribes, and genera of Saturniidae. We analyzed potential systematic bias in genomic data and used the best-supported phylogenomic hypothesis to (1) propose a revised classification of the family that reflects these inferred relationships, and (2) infer a timeline for the origin and dispersal patterns of the family through the estimation of its biogeographical history.

## Materials and Methods

### Taxon Sampling

Our taxonomic sampling included 9.8% of all species of Saturniidae (338/3454) and representatives of all recognized subfamilies (8), tribes (11), and genera (180) (Kitching et al. 2018). Our samples also included representatives of all subgenera of *Psilopygida*, *Meroleuca*, *Gonimbrasia,* and *Antheraea*, and five of the six subgenera of *Hylesia* (Table S1). Sixteen outgroup species were used, including (1) ten species of Sphingidae, the sister family of Saturniidae (Hamilton et al. 2019), with representatives from all four of its subfamilies, and (2) six species from four other families of Bombycoidea (Brahmaeidae, Endromidae, Bombycidae and Eupterotidae).

### Library Preparations and Sequencing

Samples were obtained from tissue collections at CBGP and MNHN, and from historical specimen collections at MNHN when necessary. DNA was extracted from thorax muscles or legs using the Qiagen DNeasy Blood and Tissue kit and following manufacturer’s protocol with modifications detailed in Cruaud et al. (2019). Alternatively, we also used DNA extracts prepared for DNA barcoding studies at the Centre for Biodiversity Genomics (University of Guelph, Ontario, Canada) following the protocol in Ivanova et al. (2006).

Library preparations followed Cruaud et al. (2019). Pools of 16 specimens made at equimolar ratio were enriched in 1,381 UCEs using the 14,363 baits designed by Faircloth (2017) (ordered as a myBaits UCE kit; Arbor Biosciences). Batches of 96 samples were sequenced on different flow cells of an Illumina MiSeq platform with a 2×250 pb paired-end sequencing.

### Assembly of Data Sets into Loci, Data Set Cleaning

Filtering of raw data and assembly into loci followed Cruaud et al. (2019). Only UCEs that had a sequence from at least 50% of the samples were retained for analysis (n=1,024). Loci were aligned with MAFFT (*-linsi* option; Katoh and Standley 2013). Sites with more than 50% gaps and sequences with more than 25% gaps were removed from each UCE using SEQTOOLS (package *PASTA*; Mirarab et al. 2015)). For all loci, two successive rounds of TreeShrink (Mai and Mirarab 2018) were performed to detect and remove abnormally long branches in individual gene trees, using the *per-species* mode and *b* parameter set to 20. Loci were realigned with MAFFT between and after the two rounds of TreeShrink; gene trees were inferred with IQ-TREE 2.0.6 (Minh et al. 2020) with the best fit model selected by ModelFinder (Kalyaanamoorthy et al. 2017).

### Phylogenetic Analyses

UCEs were analyzed with concatenation (IQ-TREE 2.0.6) and tree reconciliation (ASTRAL-III, Zhang et al. 2018) approaches. For the concatenation approach, aligned UCEs were merged and the resulting data set was analyzed without (hereafter referred to as IQTREE-unpartitioned) and with partitions within the alignment. For the partitioned analysis, each UCE was first divided into one core and two flanking regions using the Sliding-Window Site Characteristics (SWSC) method (Tagliacollo et al. 2018). We then tested two partitioning schemes: (1) a partition that grouped all core regions and another that grouped all flanking regions (hereafter referred to as IQTREE-CoreVsFlanking); and (2) the best partitioning scheme inferred by PartitionFinder2 (Lanfear et al. 2017) that joins core and flanking regions under a similar evolutionary model (model selection = AICc; algorithm = rclusterf; branch lengths linked; hereafter referred to as IQTREE-PartitionFinder). Best-fit models for the unpartitioned data set and for all partitions were selected with the Bayesian Information Criterion (BIC) as implemented in ModelFinder (Kalyaanamoorthy et al. 2017). FreeRate models with up to ten categories of rates were included in tests for the unpartitioned data set and for partitions that grouped all core regions or all flanking regions, but only common substitution models were tested for the subsets identified by PartitionFinder. The candidate tree set for all tree searches was composed of 98 parsimony trees + 1 BIONJ tree and only the 20 best initial trees were retained for NNI search. Node supports were assessed with ultrafast bootstrap (UFBoot) (Minh et al. 2013) with a minimum correlation coefficient set to 0.99 and 1,000 replicates of SH-aLRT tests (Guindon et al. 2010). UFBoot values ≥ 95 and SH-aLRT ≥ 80 were considered as strong support for a node. In addition, we computed gene (gCF) and site (sCF) concordance factors (Minh et al. 2020) with IQ-TREE.

For ASTRAL analyses, nodes in gene trees with UFBoot support lower than 50, 70, or 90 were collapsed following the approach of Zhu (2014, perl script AfterPhylo.pl) before reconciliation (trees are hereafter referred to as ASTRAL50, ASTRAL70 and ASTRAL90, respectively). Node supports for the ASTRAL trees were evaluated with local posterior probabilities (local PP). RF distances (Robinson and Foulds 1981) among recovered trees were calculated with RAxML-NG_v0.9.0 (Kozlov et al. 2019) and Approximately Unbiased (AU) tests (Shimodaira 2002) were performed with IQ-TREE (number of bootstrap replicates set to 20,000).

### Analysis of Conflicting Topologies and Bias Exploration

Systematic biases were investigated with respect to influences from both loci and taxa. For analyses based on loci, Gene Genealogy Interrogation (GGI) (Arcila et al. 2017, Zhong and Betancur-R 2017, Betancur-R et al. 2019) was used to identify UCEs that supported each of the competing topologies. Each UCE was analyzed with IQ-TREE to infer trees using each of the competing topologies as multifurcating constraint trees (the structure of the backbone was fixed, but taxa within identified clades were free to move around). Per-site log-likelihood scores for all constrained trees were then calculated with RAxML-8.2.11 (Stamatakis 2014) and used to perform AU tests in CONSEL (Shimodaira and Hasegawa 2001). The program *makermt* in CONSEL was used to generate K = 10 sets of bootstrap replicates with each set consisting of 100,000 replicates of the row sums. Properties of the UCEs that supported each topology were compared to detect whether significant differences could be identified between groups of UCEs and whether any topology could result from analytical bias. Two properties that are known to bias phylogenetic analyses, GC content (Romiguier and Roux 2017) and saturation (Philippe et al. 2001, Duchêne et al. 2021), were analyzed. GC content was calculated with AMAS (Borowiec 2016) and saturation (R squared of the linear regression of uncorrected p-distances against inferred distances in individual gene trees) was calculated following Borowiec et al. (2015). Results were analyzed in R (R Core Team, 2018) using the packages *ggpubr* (Kassambara 2020) and *rich* (Rossi 2011).

For analyses based on taxa, GC content and long branch (LB) score heterogeneity were calculated for each taxon in each UCE and in the concatenated data set. GC content was calculated with AMAS and LB heterogeneity scores were calculated with TreSpEx (Struck 2014). In each tree, the per sample LB score measures how the patristic distance (PD) of a sample to all others deviates (as a percentage) from the average PD across all samples (Struck, 2014). The calculation of the heterogeneity in LB scores can therefore help to reveal possible Long Branch Attraction (LBA) artefacts (Bergsten 2005). Hierarchical clustering of taxa based on GC content and LB scores was performed with the R package *cluster* (Maechler et al. 2018).

### Divergence Time Estimates

We estimated divergence times for Saturniidae with MCMCtree (Yang and Rannala 2006), using uniform distributions as calibration priors. A sphingid fossil (†*Mioclanis shanwangiana*) provided a minimum age for the crown of Smerinthinii set to 15.0 Ma (Zhang et al. 1994, Yang et al. 2007), while a maximum of 44.9 Ma was set following the maximum age estimate for Sphingidae in Kawahara et al. (2019). For secondary calibration, the minimum and maximum bounds of four selected nodes were respectively set to the mean of minimum ages and the mean of maximum ages obtained by Kawahara et al. (2019) with different calibration schemes and relaxed-clock models: (1) root (crown Bombycoidea): 59.6-81.1 Ma; (2) crown Saturniidae + Sphingidae: 48.8-69.0 Ma; (3) crown Saturniidae: 35.1-52.5 Ma; and (4) crown Sphingidae: 29.6-44.9 Ma. Analyses were run with uncorrelated and correlated relaxed clock models. The IQ-TREE topology obtained with the 50% least saturated UCEs loci was used as input tree. Five data sets composed of 30,000 randomly selected sites (custom script) were used as input data to make computation tractable. Two chains were run for each data set, 20,000 generations were discarded as burnin, and chains were run for at least 2 million generations with sampling every 200 generations and until effective sample sizes of parameters reached at least 200 as reported by Tracer (Rambaut et al. 2018). Possible conflicts between priors and data were controlled by sampling directly from the prior of times and rates. Prior and posterior distributions were then compared in Tracer. Marginal densities of parameters obtained with different data sets were compared with Tracer. Posterior estimates obtained with the different chains were combined with LogCombiner 2.6.0 (Bouckaert et al. 2019).

### Historical Biogeography

Species distributions were assigned as presence/absence to the nine following biogeographical areas: Neotropical, West Nearctic, East Nearctic, Afrotropical, Australasian, Madagascar, Oriental, West Palearctic, and East Palearctic (Table S2). Ancestral area estimations were performed using the R package *BioGeoBEARS 1.1.1* (Matzke 2014). The chronogram built with MCMCtree was used as input but only one specimen per genus/subgenus was included in the analysis and outgroups were removed to avoid artefacts (taxa were pruned from the chronogram with the R package *ape* (Paradis and Schliep 2018)).

Dispersal-Extinction-Cladogenesis (DEC; Ree and Smith 2008), BAYAREALIKE (Landis et al. 2013) and DIVALIKE (Ronquist 1997) models were used with and without considering the jump parameter for founder events (*+J*; Matzke 2014). Model selection was performed by considering statistical (AICc; Klaus and Matzke 2020, Matzke 2021) and non-statistical (i.e., biological and geographical) information (Ree and Sanmartin 2018). The maximum number of areas that a species could occupy was set to 5. To consider the main geological events that occurred during the diversification of Saturniidae, we defined five time periods with different dispersal rate scalers: (1) from the mean crown age of Saturniidae (ca. 52 Ma) to 34 Ma; (2) 34 to 23 Ma (Oligocene); (3) 23 to 14 Ma (early to mid-Miocene); (4) 14 to 5 Ma (mid to late Miocene); (5) 5 Ma to present. Dispersal rate matrices between areas and adjacency matrices are provided in Table S2.

## Results

### Initial Data Set and Trees

The initial phylogenomic data set included sequence information for 354 taxa (338 species from 185 genera of Saturniidae + 16 outgroup species) and 1,024 UCEs representing 579,775 aligned bp (215,449 bp for core regions and 364,326 bp for flanking regions) of which 66.7% were parsimony informative (Table S3). PartitionFinder grouped the 3,072 core/flanking regions into 1,943 subsets. The six trees (IQTREE-unpartitioned, IQTREE-CoreVsFlanking, IQTREE-PartitionFinder, ASTRAL50, ASTRAL70, ASTRAL90) are shown in Figure S1. Relationships within Saturniidae inferred by IQTREE were well resolved with only 6-7% of the nodes receiving SH-aLRT/UFBoot scores below 100/100. IQ-TREE trees were considered as equally good explanations of the data set (AU test *P*-value > 0.05; Table S3), regardless of the partitioning scheme.

The differences between IQ-TREE topologies (RF pairwise distances 4-8; Table S3) were located in a few poorly supported regions of the topology (within Bunaeini and Arsenurinae). Except for 8-10% of the nodes, local PP were above 0.99 in ASTRAL trees. The differences between ASTRAL topologies (RF pairwise distances 6-14; Table S3) were restricted to parts that were poorly supported. Depending on the pairs of topologies compared, from 19 to 24 clades were observed in IQ-TREE trees but not in ASTRAL trees and vice versa. All conflicts (but one, see next section) were either unsupported in IQ-TREE trees, ASTRAL trees, or in both types of trees.

Saturniidae was recovered monophyletic with maximal node support in all analyses. All subfamilies were recovered as monophyletic with strong support except Hemileucinae which was polyphyletic with *Hirpida + Hirpsinjaevia* well separated from other species. The Oxyteninae and Cercophaninae formed a grade sister to all other saturniids. Except for Agliinae and *Hirpida + Hirpsinjaevia* (Fig. 1; see next section), relationships among the main lineages were strongly supported with Arsenurinae recovered as sister to Ceratocampinae + Hemileucinae (excluding *Hirpida + Hirpsinjaevia*) and Salassinae recovered as sister to Saturniinae. All but five tribes (Hemileucini, Urotini, Micragonini, Saturniini, Bunaeini) were recovered as monophyletic with strong support. Finally, 95 (92%) of the 103 genera represented by multiple species were recovered as monophyletic. The six genera *Automeris*, *Eacles, Ludia*, *Neodiphthera*, and *Pseudodirphia* were problematic as they were rendered paraphyletic, respectively, by *Eubergioides, Catharisa, Vegetia, Pararhodia,* and *Kentroleuca*. Two genera (*Gonimbrasia*, *Molippa*) appeared polyphyletic while *Syssphinx* was recovered as a grade that also included the North American *Dryocampa* and *Anisota*.

**Figure 1.**
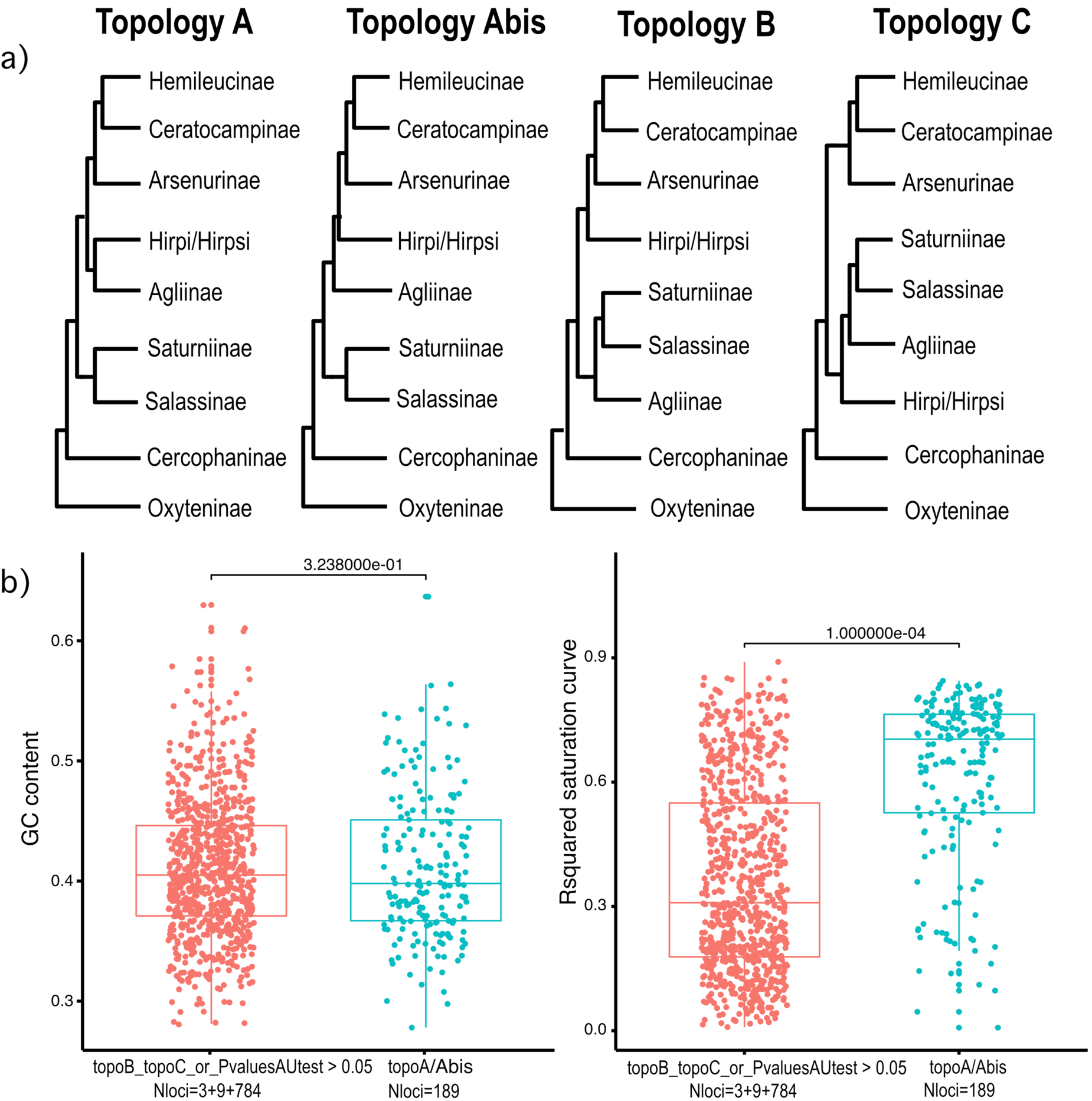
Conflicting topologies observed after analysis of the initial data set (a) and results of the gene genealogy interrogation (GGI) approach (b). In panel (a), topology *A* is supported by IQ-TREE analyses; *Abis* is supported by ASTRAL90; *B* is supported by ASTRAL70 and *C* is supported by ASTRAL50. All corresponding trees are available in Figure S1. In simplified topologies, Hemileucinae is used for Hemileucinae minus *Hirpida* and *Hirpsinjaevia* and Hirpi/Hirpsi stands for *Hirpida* + *Hirpsinjaevia.* For the GGI approach, Hemileucinae (minus *Hirpida* and *Hirpsinjaevia*) + Ceratocampinae + Arsenurinae; Saturniinae + Salassinae; *Hirpida* + *Hirpsinjaevia* and Agliinae were considered as different clades and trees were inferred using each of the competing topologies as multifurcating constraint trees. The structure of the backbone was fixed, but taxa within clades were free to move around. Topology *A* was considered equivalent to *Abis* as we focused on sister taxa relationships of Agliinae / *Hirpida* + *Hirpsinjaevia* with either Hemileucinae + Ceratocampinae + Arsenurinae or Saturniinae + Salassinae. In panel (b), average values of GC content and saturation (R-squared of the regression between pairwise distances calculated from aligned sequences and branch length) for loci that significantly supported topologies *A*/*Abis* (i.e., Pvalue of the AU test ≦ 0.05; N loci = 189; Table S3) or not (Nloci=796) were compared using a randomization test (N randomization = 9999; c2m function of the R package *rich*) and P values are reported on graph.

### Analysis of Conflicting Topologies

A single important conflict was observed among the trees with respect to the position of Agliinae and of species in the *Hirpida* + *Hirpsinjaevia* clade (Figs. 1 and S1): (1) Agliinae and *Hirpida* + *Hirpsinjaevia* either formed a clade sister to Arsenurinae + Ceratocampinae + other Hemileucinae (ACH) (IQ-TREE; topology A), or branched sequentially with Agliinae sister to a clade grouping *Hirpida* + *Hirpsinjaevia* and ACH (ASTRAL90; topology Abis); or (2) *Hirpida* + *Hirpsinjaevia* grouped with ACH while Agliinae grouped with Salassinae + Saturniinae (ASTRAL70; topology B); or (3) Agliinae and *Hirpida* + *Hirpsinjaevia* formed a grade basal to Salassinae + Saturniinae (ASTRAL50; topology C). The contrasting results obtained with ASTRAL and the gCFs of the Agliinae + *Hirpida* + *Hirpsinjaevia* clade in the IQ-TREE trees (8.6%; Fig. S1) highlight conflict among gene trees. GGI analyses show that of the 985 UCEs available for both Agliinae and *Hirpida* + *Hirpsinjaevia* species, 540 (54.8%) favored topologies A/Abis; 228 (23.2%) favored topology B; and 217 (22.0%) favored topology C (Table S3). In addition, topologies A/Abis were a significantly better explanation for 189 (19.2%) of the gene trees while topology B was a significantly better explanation for only 3 (0.3%) of the gene trees, and topology C was a significantly better explanation for 9 (0.9%) of the gene trees. Statistical analyses of UCE properties show that topologies A/Abis were supported by UCEs with lower saturation (Fig. 1(2)). Further analyses of data subsets comprising the 20% least saturated (*n*=204) or 50% most saturated (*n*=512) loci confirmed that the placement of Agliinae as sister to Salassinae + Saturniinae was likely an artefact caused by saturation of phylogenetic signal (Fig. S2). Both IQ-TREE and ASTRAL90 analyses of the least saturated UCEs recovered Agliinae close to ACH (topologies A/Abis; Fig. 1a), while Agliinae always jumped into the Saturniinae + Salassinae part of the tree when analysis focused on the most saturated loci (Fig. S2). In addition, taxon-based analysis of systematic bias did not reveal evidence that the sister taxa relationship between Agliinae and *Hirpida* + *Hirpsinjaevia* (topologies A/Abis; Fig. 1a) resulted from compositional (GC) bias or LBA (Fig. S3).

**Figure 2.**
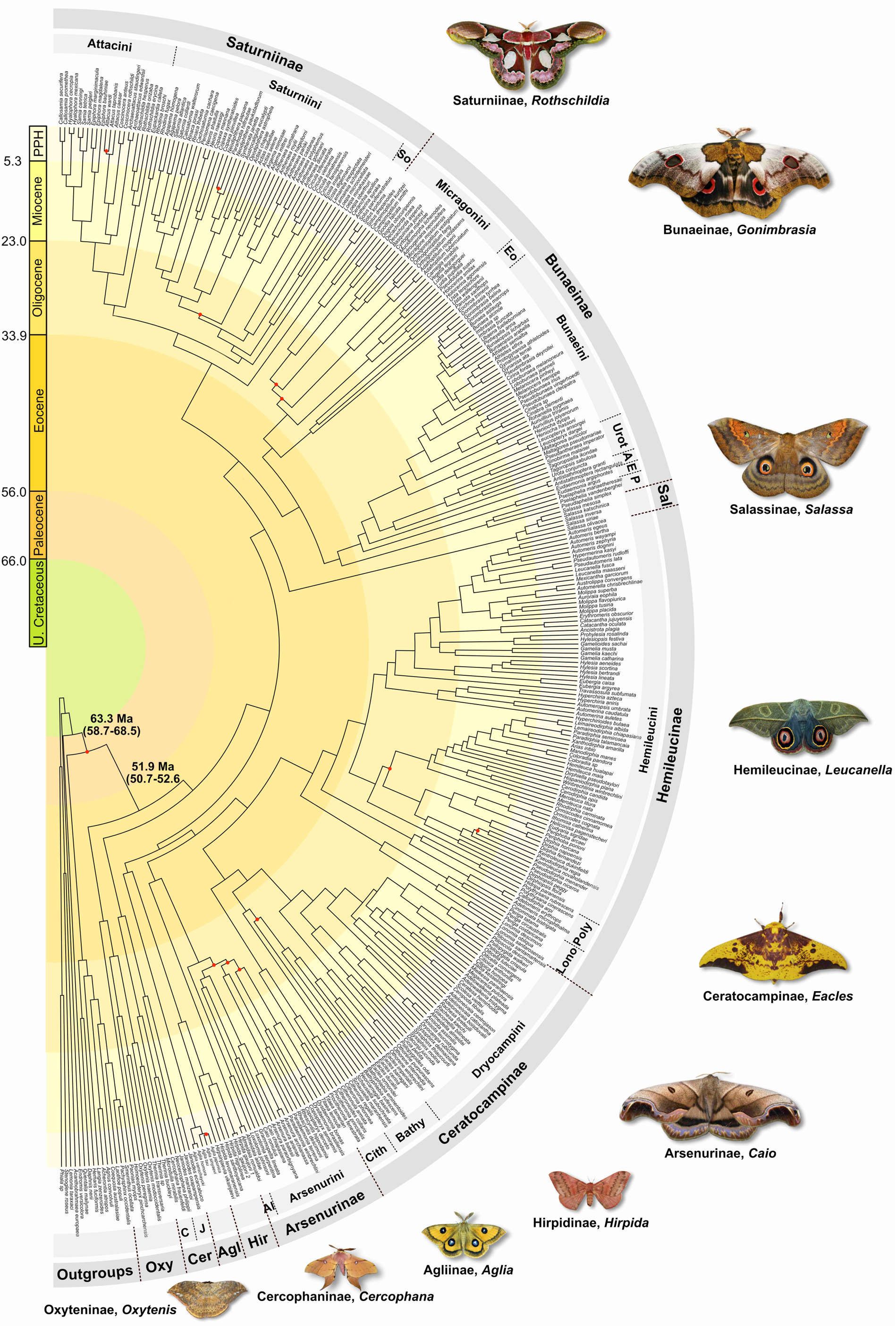
Saturniidae tree of life. The classification proposed in this study is mapped on the chronogram (MCMCtree, uncorrelated relaxed clock model) inferred from the topology selected after exploration of systematic bias (i.e., the IQ-TREE tree inferred from the 50% least saturated loci). Red dots at nodes identify poorly supported nodes (SHaLRT<80 or UFBoot<95). Abbreviations used: A = Antistathmopterini; Agl = Agliinae; Al = Almeidaiini; Bathy = Bathyphlebiini; C = Cercophanini; Cer = Cercophaninae; Cith = Citheroniini; E = Eudaemoniini; Eo = Eochroini; Hir = Hirpidinae; J = Janiodini; Lono = Lonomiini; Oxyteninae; P = Pseudapheliini; Poly = Polythysanini; Sal = Salassinae; So = Solini; Urot = Urotini.

Finally, regardless of the conflicting placement of the subfamily Agliinae, IQ-TREE and ASTRAL analyses based on the least saturated half of the loci produced the most similar topologies (RF distances; Table S3). Consequently, subsequent analyses focused on the phylogram derived from the IQ-TREE analyses of the 50% least saturated UCEs (*n*=512; Figs. 2 and S2I).

### Divergence Time Estimates and Historical Biogeography

Analyses based on both autocorrelated and uncorrelated relaxed clock models provided similar results (median of differences between mean or median ages of all nodes = 1.65 Myr; Table S4), therefore we only represent (Figs. 2 and 3) and cite estimates based on the uncorrelated relaxed clock model in main text for brevity. The divergence times and confidence intervals for correlated and uncorrelated clock models are provided in Table S4 for all nodes (see also Figure S4 for the chronograms), while those of clades discussed below are provided in Table 1.

**Figure 3.**
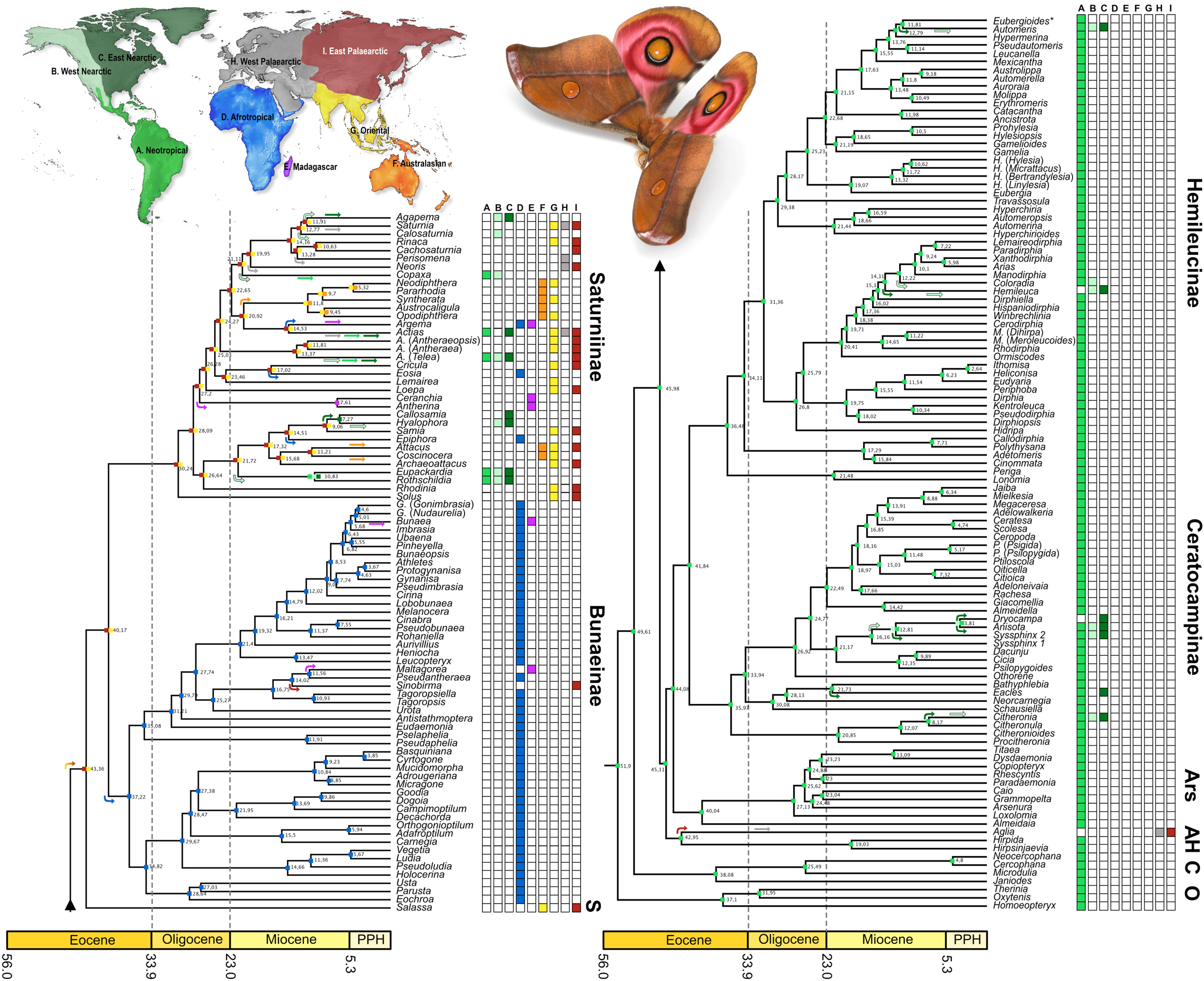
Global historical biogeography of Saturniidae. Ancestral ranges were inferred with the BAYAREALIKE+J model (best-fit and most plausible model according to empirical considerations; see text) as implemented in *BioGeoBEARS*. Alternative inferences of ancestral ranges are provided in Figure S5. The chronogram is derived from the species-level chronogram obtained with the uncorrelated relaxed clock model implemented in MCMCtree (see Fig. 2) that was pruned to keep only one specimen per genus/subgenus. The classification proposed in this study is used to annotate the tree. Colored boxes following each terminal name represent biogeographical regions A to I as illustrated in the upper-left map; colored arrows refer to inferred dispersal/colonization events. Abbreviations used: A = Agliinae; Ars = Arsenurinae; C=Cercophaninae; H = Hirpidinae; O = Oxyteninae; S = Salassinae. PPH=Pliocene, Pleistocene and Holocene. Illustrative photograph = *Antherina suraka* (Saturniinae); image courtesy of Armin Dett.

**Table 1.**
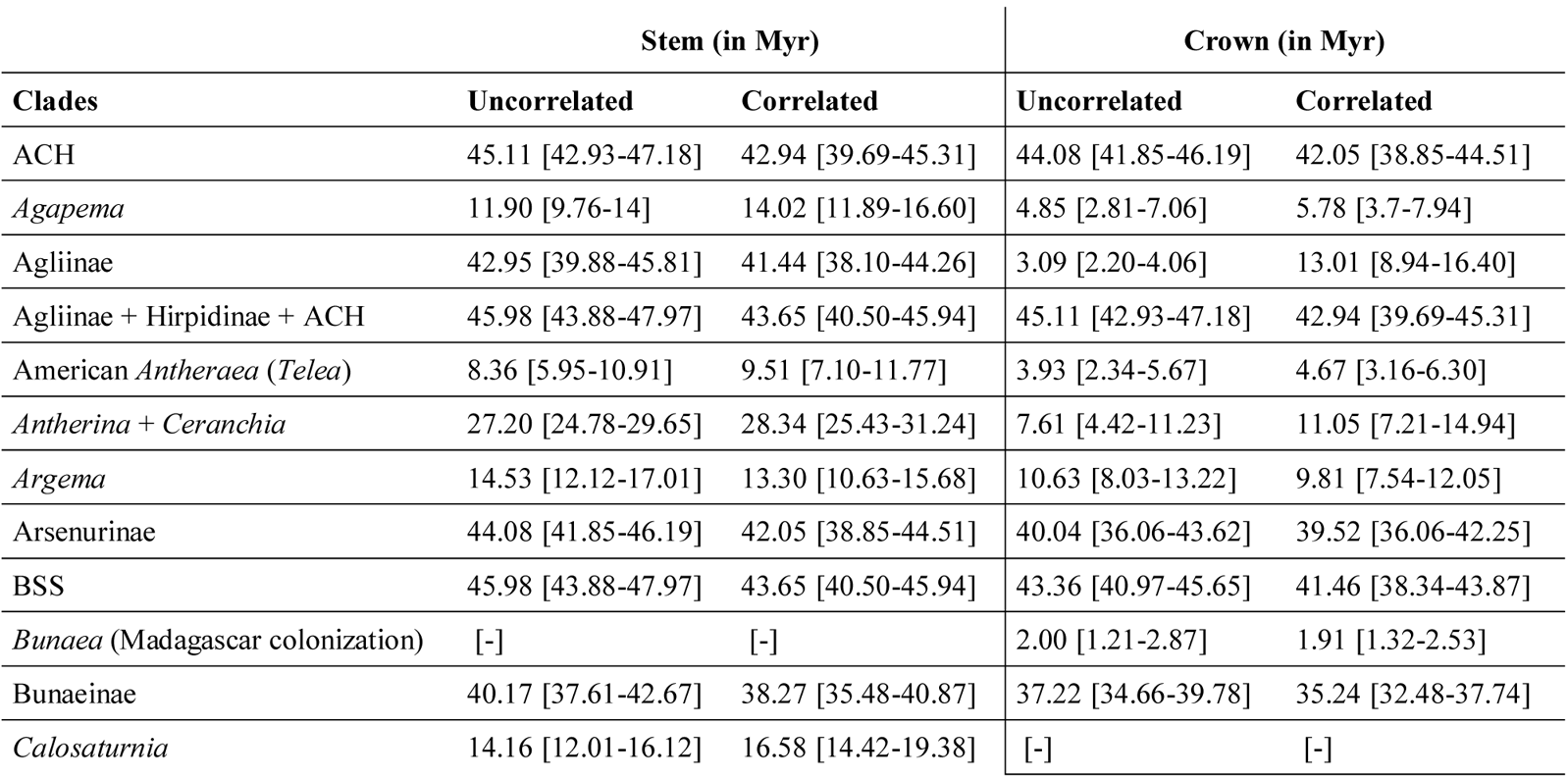
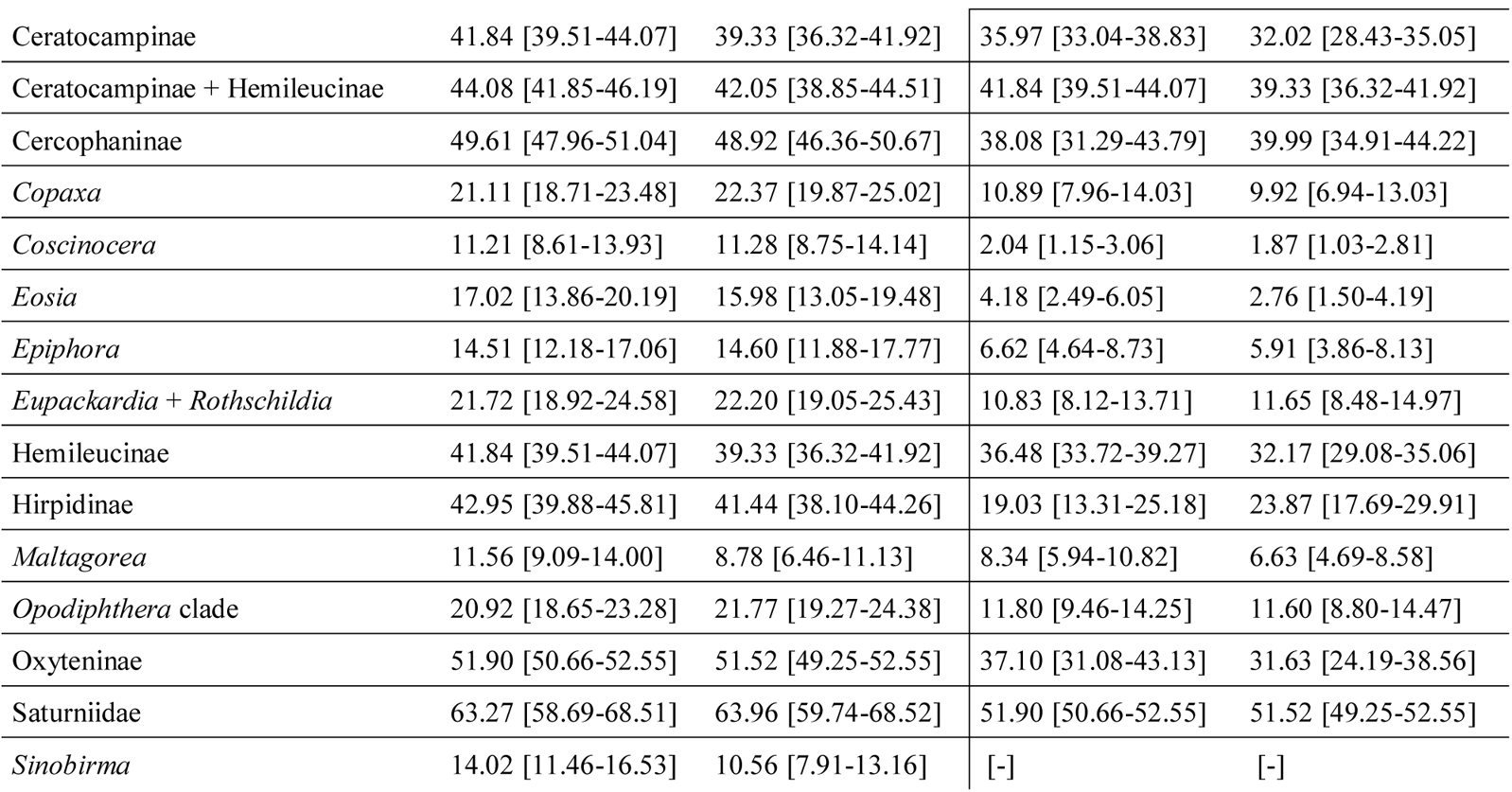
Summary of mean divergence time estimates and 95% highest posterior density (HPD) for groups or colonization events discussed. Mean and median ages with 95% HPD for all nodes are provided in Table S4. Units are in Myr. Our new classification is used for taxon names. ACH = Arsenurinae + Ceratocampinae + Hemileucinae; BSS = Bunaeinae + Salassinae + Saturniinae.

The stem age of Saturniidae was estimated in the early Cenozoic at about 63 Ma. The Oxyteninae and Cercophaninae originated about 52 Ma and 50 Ma, respectively. All other subfamilies originated during a brief interval (4 Myr) in the middle Eocene (46-42 Ma) and diversified during the Oligocene and Miocene. While only 32 (18%) of the 180 extant genera had appeared before the end of the Oligocene (including all 9 genera of Arsenurini), the remaining bulk of extant saturniid genera originated during the Miocene (Fig. 3).

With respect to historical biogeography, inclusion of the parameter *J* (jump dispersal events during cladogenesis) in models of geographic range evolution remains controversial, as is the validity of statistical comparisons of models with and without *J* (Ree and Sanmartín 2018; Matzke 2021). Empirical considerations have been advocated as an alternative way to select among biogeographical scenarios (Ree and Sanmartin 2018). Here, both AICc and the observation that most extant subfamilies occupy a single biogeographical area favor biogeographical models that include the *J* parameter (Table S2, Fig. S5). Without it, the DEC model had the best fit, but inferred a widespread ancestor for the family with multiple subsequent local extinctions. When the *J* parameter was included, BAYAREALIKE+J was inferred as the best-fit model (Table S2) and all ancestral range estimations unambiguously supported a Neotropical origin for saturniids.

Given the most likely ancestral areas estimation, saturniid biology, and palaeogeological knowledge, following their origin in the Neotropics, the Saturniidae likely colonized the Old World twice during the mid-Eocene, probably through Beringia. These two dispersal events involved an ancestor of Salassinae + Saturniinae between 46 and 43 Ma, and an ancestor of Agliinae at 43 Ma (red arrows in Figure 3). Shortly later, a lineage (represented by three tribes - Bunaeini, Micragonini and Urotini - in the current classification) colonized Africa between 40 and 37 Ma and subsequently diversified extensively (with 55 extant genera). This lineage remained restricted to this continent except for three lineages which dispersed to new areas: *Sinobirma* invaded the Oriental region (*ca*. 14 Ma), while *Maltagorea* (*ca*. 12 Ma) and *Bunaea* (*ca*. 2 Ma) colonized Madagascar. By contrast, the Attacini and Saturniini tribes in Saturniinae not only diversified in the Oriental and East Palearctic regions, but also dispersed multiple times from there to adjacent areas (Fig. 3). They colonized Africa and Madagascar on four occasions during the mid-Miocene Climatic Optimum (*Epiphora* 14 Ma, *Argema* 14 Ma, *Eosia* 17 Ma, and the clade *Antherina + Ceranchia* 27 Ma). They also dispersed through Sunda and Wallacea on multiple times, crossing Lydekker’s line on three occasions (*Attacus*, *Coscinocera*, *Austrocaligula* clade). Finally, they returned seven times to the New World with two relatively ancient dispersal events (early Miocene; *Copaxa* 21 Ma and the clade *Eupackardia* + *Rothschildia* 21 Ma) and five others during the second half of Miocene (*Actias* 14 Ma, *Antheraea* (*Telea*) 13 Ma, *Calosaturnia* 13 Ma, *Agapema* 12 Ma, and the clade *Hyalophora* + *Callosamia* 7 Ma). The diverse Neotropical fauna of saturniids colonized the Nearctic region only seven times (Fig. 3, right-hand section of tree, green arrows) and all these dispersal events postdated the final formation of the Isthmus of Panama (*ca*. 15 Ma).

## Discussion

### A Comprehensive Genus-level Phylogeny of Saturniidae

This study expands upon earlier investigations (Regier et al. 2008, Barber et al. 2015, Hamilton et al. 2019) to yield a comprehensive and robust phylogenetic framework for higher relationships within Saturniidae. The results confirm that Oxyteninae and Cercophaninae were the first two subfamilies to diverge and that Salassinae is sister to Saturniinae. The present results agree with those of Barber et al. (2015) and Hamilton et al. (2019) as they place Arsenurinae as sister to Ceratocampinae + Hemileucinae (but without *Hirpida* + *Hirpsinjaevia*; see below). However, in-depth analysis of conflict among gene trees shows that subfamily Agliinae, whose position remained unelucidated, belongs to a lineage that also includes the new-world subfamilies Arsenurinae, Ceratocampinae, and Hemileucinae. This suggests that its alternative placement as sister to Salassinae and Saturniinae as obtained by Hamilton et al. (2019) was also likely an artefact of tree reconstruction, possibly caused by the saturation of nucleotide sequences. Indeed, when these authors analyzed the AHE loci using protein sequence data which saturate less rapidly owing to a larger state space (20 possible amino acids; see their additional file 8), they recovered Agliinae as more closely related to new-world subfamilies. These relationships are corroborated by larval characters shared with Arsenurinae (Michener 1952; e.g., D2 setae on abdominal segment A9 of first instar larva form a median scolus or chalaza; first larval stages bear two or more pairs of large thoracic horns forked at their apices). Interestingly, our analyses are the first to examine the genera *Hirpida* and *Hirpsinjaevia* and they reveal that these two closely related Neotropical genera are sister to Agliinae. Unfortunately, the host plant(s) and immature stages of *Hirpida* and *Hirpsinjaevia* are unknown but, based on their phylogenetic placement, we predict that their first instar larvae bear large thoracic horns and may feed on Fagaceae and/or Betulaceae like the Agliinae.

In the arsenurine tribe Arsenurini, Hamilton et al. (2021) found from a thorough phylogenomic analysis of hundreds of AHE genes that phylogenetic signal remained elusive, likely reflecting a burst of diversification. Our analysis also found that relationships among its component genera are poorly supported (Figs. 2 and S2I) and are discordant from the hypothesis preferred by Hamilton et al. (2021). This casts doubt on current understanding of their evolutionary history, an uncertainty that can only be resolved by further investigation, perhaps by combining AHE and UCE loci and expanding taxon sampling. Within the Ceratocampinae and Hemileucinae, our results provide the first robust, comprehensive hypothesis of relationships among its genera. They fully agree with conclusions based on a 6-gene data set and limited taxon sampling (Barber et al 2015), but largely contradict groupings proposed for Ceratocampinae by Balcazar-Lara & Wolfe (1997) based on their cladistic analysis of both adult and immature morphology.

Within Saturniinae, our results unequivocally support the position of *Rhodinia* as sister to members of the Attacini, in agreement with AHE-based conclusions (Rubin et al. 2018) and recent findings from the analysis of mitogenomes (Chen et al. 2021). In addition, as our sampling examined all genera in the tribe Urotini – considered to represent an artificial grouping of several unrelated lineages (Oberprieler 1997; Rougerie 2005; Barber et al 2015; Rubin et al 2018) – the outcome of our analyses offers the first robust framework of the evolutionary history of African saturniids to lay down a new classification.

### Taxonomic Implications

Our phylogenetic hypothesis has important and unexpected implications for the classification of Saturniidae (Fig. 2). We propose several taxonomic changes that are detailed below and summarized as a complete genus-level checklist (SI Appendix S1; see also Table S1). The unexpected placement of *Hirpida + Hirpsinjaevia* as sister to Agliinae as well as the age of the lineage justify the erection of a new subfamily: Hirpidinae Rougerie **subfam. nov**. As all three previously recognized Afrotropical tribes of Saturniinae (Bunaeini, Urotini, Micragonini) form a monophyletic group sister to all other Saturniinae (Attacini, Saturniini and Solus), we adopt here the classification proposed by Nässig et al. (2015) that recognizes this clade as subfamily Bunaeinae Bouvier, **1927 stat. rev.**. Within this subfamily, we consider Bunaeini, Micragonini, and Urotini as monophyletic tribes but with new circumscriptions ensuring their monophyly. To achieve this, we describe a new tribe and resurrect three others involving the following seven genera: *Eochroa* (previously in Bunaeini), *Parusta* and *Usta* are assigned to Eochroini Cooper, 2002 **stat. rev.**; *Pseudaphelia* and *Pselaphelia* to Pseudapheliini Packard, 1914 **stat. rev.**; *Eudaemonia* to Eudaemoniini Packard, 1902 **stat. rev.**; and *Antistathmoptera* to Antistathmopterini Rougerie **trib. nov.**. The rare, enigmatic east African genus *Eosia* Le Cerf,1911 (Bouyer 2002), previously placed within Micragonini (Bunaeinae), is recovered as sister to *Cricula* Walker, 1855 of the Saturniini (Saturniinae) to which it is transferred, a position which agrees with evidence from the morphology of first instar larva (Rougerie 2005). The genus *Solus*, whose phylogenetic placement is clarified here for the first time, is sister to Attacini + Saturniini, a position that requires the erection of a new tribe: Solini Rougerie **trib. nov.**. We also propose to follow Nässig (1991) in considering *Graellsia* Grote, 1896 to be a junior synonym of *Actias* Leach, 1815. Since our results propose the first robust, comprehensive phylogeny for the two most diverse subfamilies of new-world Saturniidae, we propose a new tribal arrangement for their genera (Fig. 2; see Table S1 and Appendix S1 for details): (1) in Ceratocampinae, we resurrect the following three tribes: Bathyphlebiini Travassos & Noronha, 1967 **stat. rev.**, Ceratocampini Harris, 1841 **stat. rev.**, and Dryocampini Grote & Robinson, 1866 **stat. rev.**; and (2) in Hemileucinae, we expand the previously monotypic tribe Polythysanini Michener 1952 by adding the genera *Adetomeris*, *Callodirphia*, and *Cinommata*; resurrect tribe Lonomiini Bouvier, 1930 **stat. rev.** for genera *Lonomia* and *Periga*; and exclude *Catharisa cerina* Jordan 1911, considered until now as an hemileucine (Lemaire 2002, Smith et al. 2013), while we recovered it nested within the genus *Eacles* of the Ceratocampinae. This unexpected placement agrees with recent observations of the morphology of its first instar caterpillars (P. Smith, pers. comm.) and leads us to treat *Catharisa* Jordan, 1911 **syn. nov.** as a junior synonym of *Eacles* Hübner, [1819], with *Eacles cerina* **comb. nov.** transferred within Bathyphlebiini of the Ceratocampinae. Within the Hemileucinae, the monotypic genus *Eubergioides* Michener, 1949 **syn. nov.** appears nested within *Automeris* Hübner, [1819], an affinity already suggested by Michener (1952) based on the comparison of male genitalia; it is considered a junior synonym of the latter, with *Automeris bertha* (Schaus, 1896) **comb. nov.** transferred to genus *Automeris*. Among the African bunaeine saturniids, we resurrect the genus *Pinheyella* Cooper, 2002 **stat. rev.** for *P. anna* Maassen, 1985 **comb. nov.** whose current placement in genus *Gonimbrasia* Butler, 1878 makes that genus polyphyletic. Finally, we note that our results question the monophyly of six genera, but we refrain from proposing nomenclatural change as limited sampling within each genus impedes proper circumscription of natural groups: *Neodiphthera* Fletcher, 1982; *Ludia* Wallengren, 1865; *Molippa* Walker, 1855; *Pseudodirphia* Bouvier, 1928; *Kentroleuca* Draudt, 1929; and *Syssphinx* Hübner, [1819]. From here on, subfamily, tribe and genus names refer to the new classification proposed in this study (Fig. 2; Appendix S1).

### Historical Biogeography

This work represents a first investigation of the historical biogeography of saturniids (Fig. 3) based on a robust dated phylogeny that includes all known extant genera. It reveals that the family originated in the Neotropics shortly after the Cretaceous-Paleogene boundary. Its diversification was marked by a key split about 46 Ma between lineages AACHH (formed by subfamilies Agliinae, Arsenurinae, Ceratocampinae, Hemileucinae and Hirpidinae; >2,000 extant species in 96 genera) and BSS (Bunaeinae, Salassinae and Saturniinae; >1,350 species in 86 genera). The former remained confined to the New World except for the Agliinae (see below), while the later diversified in all biogeographical regions, before recolonizing the New World on several occasions. This split had been preceded by two earlier branching lineages (first Oxyteninae 52 Ma, then Cercophaninae 50 Ma) that have remained restricted to South America where they comprise 70 and 122 extant species respectively (Kitching et al. 2018, Brechlin 2020).

Dispersal out of the Neotropics and into the Old World of the ancestors of lineage BSS occurred during the middle Eocene (46-43 Ma). As dispersal across the oceanic barriers separating the New and Old Worlds seems unlikely, it is more likely that dispersal occurred via the Nearctic during the mid-Eocene Climatic Optimum, a transient period of global warming (Westerhold et al. 2020). This would have required over-water dispersal through an archipelago in the Panamanian gap (Pindell et al. 2005, Iturralde-Vinent 2006) or via the Caribbean Arc that subsided toward the end of the Eocene (Pindell et al. 2005, Philippon et al. 2020, Cornée et al. 2021, Garrocq et al. 2021) before their northward dispersal. At this time period, further dispersal of the BSS ancestors from the Nearctic into the Eastern Palearctic, and from there southwards into the Oriental region would have been possible through the Boreotropical forest belt (Morley 2003, 2011), which extended as far as the Arctic Circle, facilitating faunal exchanges (Jahren 2007). Our results also suggest that ancestors of the Agliinae, within the AACHH lineage, likely used the same route through Beringia about 43 Ma as an independent long-distance dispersal from the New to Old World. Interestingly, the food plants (Ballesteros-Mejia et al 2020) of both Agliinae and early lineages of the BSS lineage (e.g., beech, oak, maple, plum trees, sumac) were well represented in the forests of the northern hemisphere during middle Eocene (Jahren, 2007), thus lending support to this dispersal route.

During this same period (Fig. 3; 40-37 Ma), ancestors of Bunaeinae dispersed from the Oriental to African region. This dispersal was facilitated by the mixed deciduous and evergreen forests occurring in Central Asia and by the presence of archipelagoes and land connections. Between 41 and 35 Ma, Balkanotolia, an archipelago that spanned the Neotethyan margin (Licht et al., 2022) and the Alboran/Apulian platforms emerged (Vandenberghe et al. 2012), possibly facilitating the colonization of Africa. This dispersal may have been driven by the climatic deterioration (or “Grande Coupure”) (Hartenberger 1998, Licht et al. 2022) that occurred during the Eocene-Oligocene transition period, effectively forcing the Bunaeinae ancestor to seek warmer conditions southward. Interestingly, several groups of mammals (e.g., rodents, primates) with excellent fossil records also colonized Africa near the Bartonian-Priabonian boundary (*ca* 37.8 Ma) (Huchon et al. 2007, Chaimanee et al. 2012, Marivaux et al. 2014, Coster et al. 2015, Licht et al. 2022). After colonizing Africa, the ancestral Bunaeinae experienced a spectacular diversification as all extant tribes had appeared by the early Oligocene. Further investigation of the importance of different key innovations (ecological or morphological) that have triggered this diversification burst constitute an interesting perspective.

As cooling during the Eocene–Oligocene disrupted the belt of Boreotropical forests, the accretion of ice sheets broke the Beringian connection, leading to the divergence of ancestral Saturniinae in these forests. Our analyses (Figs. 3 and S5) show that the Saturniinae and most of its lineages originated in the Oriental + Eastern Palearctic region, but as the Boreotropical forests contracted drastically during the “Grande Coupure”, this likely forced southward displacements of their ranges into refuges in the Sino-Himalayan mountains of the Eastern Palearctic region (LePage et al. 2005, Xing and Ree 2017, Jia and Bartish 2018). The observation that the diversity of non-African members of the BSS clade peaks in the Sino-Himalayan mountains may reflect the role of this transition zone as a hub for further diversification of these moths. A number of Saturniinae lineages subsequently dispersed and diversified extensively in the Oriental and Palearctic regions, colonizing the Western Palearctic four times: *Neoris* (stem age 20 Ma), *Perisomena* (13 Ma), *Saturnia* (12Ma), and *Actias* (7 Ma). They also colonized Australasia (the *Opodiphthera* group (21 Ma) and *Coscinocera* (11 Ma) within the Attacini), the Afrotropics (*Epiphora* (14 Ma), *Eosia* (17 Ma) and *Argema* (14 Ma)), and the Nearctic and Neotropical regions (*Eupackardia* + *Rothschildia* (22 Ma), *Copaxa* (21 Ma), *Calosaturnia* (13 Ma), and *Agapema* (12 Ma)). Interestingly, a single dispersal event by a member of the genus *Sinobirma* (subfamily Bunaeinae) occurred from Africa to the Eastern Palearctic region, nearly synchronous (14 Ma) with the colonization of the Afrotropics by *Epiphora*, *Eosia,* and *Argema.* These range expansions may have been facilitated by the collision of the Arabian plate with Eurasia *ca.* 15 Ma (Rogl 1999, Harzhauser et al. 2007).

### Final Remarks

Our knowledge of insect evolution remains strongly impeded by both the scale of their diversity and the difficulty to generate accurate and robust spatial and evolutionary frameworks based on comprehensive coverage. Currently, only a few densely sampled and time-calibrated phylogenies have been generated in diverse groups of insects, bringing insights into the origins of observed diversity patterns such as the latitudinal diversity gradient (e.g., Condamine et al. 2012, Chazot et al. 2021 for butterflies; Economo et al. 2018 for ants), or the influence of life-history traits on diversification dynamics (Sota et al. 2022). We believe that the Saturniidae family, as one of the best-documented family of insects with global distribution and high species-richness, represents a new promising model to further investigate the mechanisms driving diversification in insects. Our results have revealed that the spatial dynamism of Saturniinae (33 genera, 741 species) is in striking contrast (Fig. 3) to the situation in other subfamilies that also diversified extensively (Bunaeinae: 50 genera, 580 species; Ceratocampinae: 30 genera, ca. 300 species; Hemileucinae: 55 genera, ca. 1600 species; see Kitching et al. (2018)), but remained restricted to the biogeographical region where they originated: Bunaeinae in the Afrotropical region (except for *Sinobirma*, and including two independent colonization of Madagascar); Ceratocampinae and Hemileucinae in the Neotropics with limited dispersal to the Nearctic region. The reasons for these contrasting dispersal patterns remain unclear, but differing life-history traits, such as larval diet and pupation modes (cocoons in Saturniinae are among the strongest within the family) may have facilitated long-distance dispersal and range expansion. In general, the new spatial and temporal framework produced here for the evolution of wild silkmoths opens new avenues of research to understand the intriguing success of these capital-breeding moths that managed to occupy all biogeographical regions today and to represent one of the most diverse and abundant group of large-sized insects.

## Acknowledgements

We are thankful to the Canadian Centre for DNA Barcoding and Centre for Biodiversity Genomics at University of Guelph (Ontario, Canada) as well as to all the participants to the Saturniidae DNA barcoding campaign for their contributions to the assembly of samples. We thank Corinne Poitout, Hélène Vignes and Audrey Weber (INRAE and CIRAD, AGAP, France) for their assistance to sequencing, and the Genotoul bioinformatics platform (Toulouse, Occitanie, France) for providing computing resources.

## Taxonomic Appendix – New Higher Taxa

Hirpidinae Rougerie, **subfamilia nova**

http://zoobank.org/LSID######

Type genus: *Hirpida* Draudt, 1930

This subfamily comprises two Andean genera, *Hirpida* and *Hirpsinjaevia*. A suggested combination of diagnostic features includes the following characters of the adults proposed by Michener (1952) as distinctive for genus *Hirpida*: head with the sides of the frons slightly convex; tarsomeres with spines on their ventral surface; well-developed arolia and pulvilli; absence or reduction of vein 3A in hindwings; tegulae long, reaching the anterior median angle of mesothoracic scutellum; anepisternal suture largely horizontal but directed slightly downward posteriorly; male genitalia with free juxta and articulated valvae. The status of these characters remains unknown in the recently described, rare genus *Hirpsinjaevia* as it could not be examined for this study. Further comparative morphology and a formal phylogenetic analysis of characters are needed to identify unique characters or more likely a unique combination of characters that can accurately diagnose members of the subfamily Hirpidinae.

Antistathmopterini Rougerie, **tribus nova**

http://zoobank.org/LSID######

Type genus: *Antistathmoptera* Tams, 1935

This tribe only includes species in the genus *Antistathmoptera*, all of which appear restricted to Eastern Africa. Because of their rarity in collections, little is known of their morphology, but they can be easily distinguished by a unique combination of adult characters: long tailed hindwings; pointed forewing apex; both pairs of wings with a discal spot formed of two contiguous hyaline areas.

Solini Rougerie, **tribus nova**

http://zoobank.org/LSID######

Type genus: *Solus* Watson, 1913

This tribe only includes the genus *Solus* which is restricted to the continental part of the Oriental region. The following combination of characters in adults are diagnostic: vein d3 of forewing discal cell missing, resulting in an open discal cell; lower branch of fork at the start of forewing vein 1A+2A much thinner than the upper branch; extremity of axillary sclerite 1 of forewings rounded; anapleural suture of mesothorax reduced; galeae present, short and broad.

## Author Contributions

Designed the study: AC, JYR, PA, RR

Obtained funding: AC, CLV, FLC, JM, JYR, ME, RR, SL, TD

Contributed samples: CLV, JYR, PDNH, RR, TD

Identified samples: IJK, JYR, RR, TD

Performed laboratory work: AC, DG, JYR, PA, RR, SN, YC

Analyzed data: AC, JYR, PA, RR

Discussed results: AC, CLV, FLC, IJK, JM, JYR, LBM, ME, PA, RR, SL, TD

Drafted the manuscript: AC, JYR, PA, RR

All authors commented on the manuscript

## Data Availability

Fastq paired reads for analyzed samples are available as a NCBI Sequence Read Archive (ID#PRJNA XXXXX). Custom script for random sampling of sites within data sets is available from https://github.com/acruaud/saturniidae_phylogenomics_2022. Data sets (concatenated UCEs) and trees have been uploaded on Dryad (http://dx.doi.org/10.5061/dryad.[NNNN]).

