## Supplementary material for "Phylogenomics Illuminates the Evolutionary History of Wild Silkmoths in Space and Time (Lepidoptera: Saturniidae)": SI Appendix S1

**Appendix S1 –** Revised classification of Saturniidae (in alphabetical order within each rank); a description and a diagnosis of newly described taxa are provided after the list.

### Family Saturniidae Boisduval, [1837]

#### Subfamily Agliinae Packard, 1893

*Aglia* Ochsenheimer, 1810

#### Subfamily Arsenurinae Jordan, 1922

Tribe Almeidaiini Lemaire, 1980

*Almeidaia* Travassos, 1937

Tribe Arsenurini Jordan, 1922

*Arsenura* Duncan [& Westwood], 1841

*Caio* Travassos & Noronha, 1968

*Copiopteryx* Duncan [& Westwood], 1841

*Dysdaemonia* Hübner, [1819]

*Grammopelta* Rothschild, 1907

*Loxolomia* Maassen, 1869

*Paradaemonia* Bouvier, 1925

*Rhescyntis* Hübner, [1819]

*Titaea* Hübner, [1823]

#### Subfamily Bunaeinae Packard, 1902

Tribe Antistathmopterini Rougerie, 2022 **trib. nov.**

*Antistathmoptera* Tams, 1935

Tribe Bunaeini Packard, 1902

- *Athletes* Karsch, 1896
- *Aurivillius* Packard, 1902
- *Bunaea* Hübner, [1819]
- *Bunaeopsis* Bouvier, 1927
- *Cinabra* Sonthonnax, 1901
- *Cirina* Walker, 1855
- *Gonimbrasia* Butler, 1878
    - *Gonimbrasia* Butler, 1878
    - *Nudaurelia* Rothschild, 1895
- *Gynanisa* Walker, 1855
- *Heniocha* Hübner, [1819]
- *Imbrasia* Hübner, [1819]
- *Leucopteryx* Packard, 1903
- *Lobobunaea* Packard, 1901
- *Melanocera* Sonthonnax, 1901
- *Pinheyella* Cooper, 2002
- *Protogynanisa* Rougeot, 1971
- *Pseudimbrasia* Rougeot, 1962
- *Pseudobunaea* Bouvier, 1927
- *Rohaniella* Bouvier, 1927
- *Ubaena* Karsch, 1900

Tribe Eochroini Cooper, 2002

- *Eochroa* Felder, C. & Felder, R., 1874
- *Parusta* Rothschild, 1907

*Usta* Wallengren, 1863

Tribe Eudaemoniiini Packard, 1902

*Eudaemonia* Hübner, [1819]

Tribe Micragonini Cockerell, 1914

*Adafroptilum* Darge, 2004

*Adrougeriana* Darge, 2015

*Basquiniana* Darge, 2015

*Campimoptilum* Karsch, 1896

*Carnegia* Holland, 1896

*Cyrtogone* Walker, 1855

*Decachorda* Aurivillius, 1898

*Dogoia* Bouyer, 2015

*Goodia* Holland, 1893

*Holocerina* Pinhey, 1956

*Holocerina* Pinhey, 1956

*Ludia* Wallengren, 1865

*Micragone* Walker, 1855

*Mucidomorpha* Darge, 2015

*Orthogonioptilum* Karsch, 1893

*Pseudoludia* Strand, 1911

*Vegetia* Jordan, 1922

Tribe Pseudapheliini Packard, 1914

*Pselaphelia Aurivillius, 1904*

*Pseudaphelia Kirby, 1892*

Tribe Urotini Packard, 1902

*Maltagorea* Bouyer, 1993

*Pseudantheraea* Weymer, 1892

*Sinobirma* Bryk, 1944

*Tagoropsiella* Darge, 2008

*Tagoropsis* Felder, C. & Felder, R., 1874

*Urota* Westwood, 1849

**Subfamily Ceratocampinae Harris, 1841**

Tribe Bathyphlebiini Travassos & Noronha, 1967

*Bathyphlebia* Felder, C. & Felder, R., 1874

*Eacles* Hübner, [1819]

*Catharisa* Jordan, 1911 **syn. nov.**

*Neorcarnegia* Draudt, 1930

*Schausiella* Bouvier, 1930

Tribe Citheroniini Harris, 1841

*Citheronia* Hübner, [1819]

*Citheronioides* Lemaire, 1988

*Citheronula* Michener, 1949

*Procitheronia* Michener, 1949

Tribe Dryocampini Grote & Robinson, 1866

*Adeloneivaia* Travassos, 1940

*Adelowalkeria* Travassos, 1941

*Almeidella* Oiticica Filho, 1946

*Anisota* Hübner, [1820]

*Ceratesa* Michener, 1949

*Ceropoda* Michener, 1949

*Cicia* Oiticica Filho, 1964

*Citioica* Travassos & Noronha, 1965

*Dacunju* Travassos & Noronha, 1965

*Dryocampa* Harris, 1933

*Giacomellia* Bouvier, 1930

*Jaiba* Lemaire, Tangerini & Mielke, 1999

*Megaceresa* Michener, 1949

*Mielkesia* Lemaire, 1988

*Oiticella* Travassos & Noronha, 1965

*Othorene* Boisduval, 1872

*Psilopygida* Michener, 1949

*Psigida* Oiticica Filho, 1959

*Psilopygida* Michener, 1949

*Psilopygoides* Michener, 1949

*Ptiloscola* Michener, 1949

*Rachesa* Michener, 1949

*Scolesa* Michener, 1949

*Syssphinx* Hübner, [1819]

**Subfamily Cercophaninae Jordan, 1924**

Tribe Cercophanini Jordan, 1924

*Cercophana* Felder, C., 1862

*Microdulia* Jordan, 1924

*Neocercophana* Izquierdo, 1895

Tribe Janiodini Jordan, 1924

*Janiodes* Jordan, 1924

**Subfamily Hemileucinae Grote & Robinson, 1866**

Tribe Hemileucini Grote & Robinson, 1866

Subtribe Automeriina Bouvier, 1928

*Ancistrota* Hübner, [1819]

*Auroraia* Brechlin & Meister, 2011

*Austrolippa* Brechlin & Meister, 2011

*Automerella* Felder, C. & Felder, R., 1874

*Automerina* Michener, 1949

*Automeris* Hübner, [1819]

*Eubergioides* Michener, 1949 **syn. nov.**

*Automeropsis* Lemaire, 1969

*Catacantha* Bouvier, 1930

*Erythromeris* Lemaire, 1969

*Eubergia* Bouvier, 1929

*Gamelia* Hübner, [1819]

*Gamelioides* Lemaire, 1988

*Hylesia* Hübner, [1820]

*Bertrandylesia* Brechlin & Meister, 2016

*Extremylesia* Brechlin & Meister, 2016

*Gamylesia* Brechlin & Meister, 2016

*Hylesia* Hübner, [1820]

*Linylesia* Brechlin & Meister, 2016

*Micrattacus* Walker, 1855

*Hylesiopsis* Bouvier, 1929

*Hyperchiria* Hübner, [1819]

*Hyperchirioides* Lemaire, 1981

*Hypermerina* Lemaire, 1969

*Leucanella* Lemaire, 1969

*Mexicantha* Naumann, Nässig & Nogueira G, 2012

*Molippa* Walker, 1855

*Prohylesia* Draudt, 1929

*Pseudautomeris* Lemaire, 1967

*Travassosula* Michener, 1949

Subtribe Hemileucina Grote & Robinson, 1866

*Arias* Lemaire, 1995

*Cerodirphia* Michener, 1949

*Coloradia* Blake, 1863

*Dirphia* Hübner, [1819]

*Dirphiella* Michener, 1949

*Dirphiopsis* Bouvier, 1928

*Eudyaria* Grote, 1896

*Heliconisa* Walker, 1855

*Hemileuca* Walker, 1855

*Hidripa* Draudt, 1929

*Hispaniodirphia* Lemaire, 1999

*Ithomisa* Oberthür, 1881

*Kentroleuca* Draudt, 1929

*Lemaireodirphia* Brechlin & Meister, 2012

*Manodirphia* Brechlin & Meister, 2012

*Meroleuca* Packard, 1904

*Dihirpa* Draudt, 1929

*Meroleuca* Packard, 1904

*Meroleucoides* Michener, 1949

*Ormiscodes* Blanchard, 1852

*Paradirphia* Michener, 1949

*Periphoba* Hübner, [1820]

*Pseudodirphia* Bouvier, 1928

*Rhodirphia* Michener, 1949

*Winbrechlinia* Brechlin, 2016

*Xanthodirphia* Michener, 1949

Tribe Lonomiini Bouvier, 1930

*Lonomia* Walker, 1855

*Periga* Walker, 1855

Tribe Polythysanini Michener, 1952

*Adetomeris* Michener, 1949

*Callodirphia* Michener, 1949

*Cinommata* Butler, 1882

*Polythysana* Walker, 1855

**Subfamily Hirpidinae Rougerie, 2022 subfam. nov.**

*Hirpida* Draudt, 1930

*Hirpsinjaevia* Brechlin, 2019

**Subfamily Oxyteninae Jordan, 1924**

*Homoeopteryx* Felder, C. & Felder, R., 1874

*Oxytenis* Hübner, [1819]

*Therinia* Hübner, [1823]

**Subfamily Salassinae Michener, 1949**

*Salassa* Moore, 1859

**Subfamily Saturniinae Boisduval, [1837]**

Tribe Attacini Blanchard, 1840

*Archaeoattacus* Watson, 1914

*Attacus* Linnaeus, 1767

*Callosamia* Packard, 1864

*Coscinocera* Butler, 1879

*Epiphora* Wallengren, 1860

*Eupackardia* Cockerell, 1912

*Hyalophora* Duncan [& Westwood], 1841

*Rhodinia* Staudinger, 1892

*Rothschildia* Grote, 1896

*Samia* Hübner, [1819]

Tribe Saturniini Boisduval, [1837]

*Actias* Leach, 1815

*Agapema* Neumoegen & Dyar, 1894

*Antheraea* Hübner, [1819]

*Antheraea* Hübner, [1819]

*Antheraeopsis* Wood-Mason, 1886

*Telea* Hübner, [1819]

*Antherina* Sonthonnax, 1901

*Argema* Wallengren, 1858

*Austrocaligula* Cockerell, 1914

*Cachosaturnia* Naumann, Löffler & Nässig, 2012

*Calosaturnia* Smith, 1886

*Ceranchia* Butler, 1878

*Copaxa* Walker, 1855

*Cricula* Walker, 1855

*Eosia* Le Cerf, 1911

*Lemaireia* Nässig & Holloway, 1988

*Loepa* Moore, [1860]

*Neodiphthera* Fletcher, 1982

*Neoris* Moore, 1862

*Opodiphthera* Wallengren, 1858

*Pararhodia* Cockerell, 1914

*Perisomena* Walker, 1855

*Rinaca* Walker, 1855

*Saturnia* Schrank, 1802

*Syntherata* Maassen, [1873]

Tribe Solini Rougerie, 2022 **trib. nov.**

*Solus* Watson, 1913

Hirpidinae Rougerie, **subfam. Nov**

Type genus: *Hirpida* Draudt, 1930

This description is based mainly on the detailed descriptions of *Hirpida* by Michener (1952) and Lemaire (2002) who only had knowledge of few (3 and 4, respectively) of the 42 currently known species within the genus. Furthermore, they had no knowledge of the recently described new genus and species *Hirpsinjaevia viksinjaevi* (Brechlin, 2019a). Their descriptions are here critically considered in the light of the information about adult habitus and male genitalia available from the description of genus *Hirpsinjaevia* and from multiple species descriptions of *Hirpida* species (e.g., Brechlin, 2019b).

**Description:** Small to medium-sized (forewing length: 19-36mm) Andean moths of brown or pink background coloration, with reduced discal markings; upperside patterns of forewing marked by distinct antemedian and postmedian bands and a narrow or reduced submarginal band; hindwing uniform in color with faint markings (see illustrations and species description and diagnoses for *Hirpsinjaevia* in Brechlin (2019a) and for *Hirpida* in Brechlin (2019b), Michener (1952) and Lemaire (2002)). Underside with faint markings. Forewing subtriangular, hindwing rounded; male and female individuals of similar size and color patterns. Venation of forewing and hindwing in *Hirpida* (undocumented in *Hirpsinjaevia*) with vein M2 arising from the middle of the apex of the discal cell, and M1 stalked with Rs or arising from the distal angle of the discal cell (vein d1 absent); R and Rs veins of forewing separated, the former arising from the anterior border of the discal cell. Hindwing with vein 3A absent. Head with frons slightly convex, but laterofrontal sutures visible and extremely close to the eye margin (Michener, 1952). Antennae quadripectinate to the apex in males, bidentate and filiform in females; labial palpi three-segmented. Forelegs of both sexes with a long epiphysis; tibial spines absent and tibial spurs number 0-2-2 (Lemaire, 2002); ventral surface of tarsal segments with spines. In males, eighth abdominal segment undifferentiated; genital apparatus with uncus apically simple, but showing much variation in size and shape, sometimes bilobed dorsally; gnathos absent or reduced to lateral arms in *Hirpida*, but forming two lobes in *Hirpsinjaevia* (Brechlin, 2019a); the juxta is free form the valvae and vinculum, and it extends posteriorly, sometimes forming a long hooked median process (see Brechlin (2019b)). Vinculum short and rounded; valvae broadly square-shaped with two distinct lobes, the lower one being either pointed and reduced, or as long as the upper one and bearing two distinctive lobes. The aedeagus is variable in size, as is the development of its caecum penis; cornuti either absent or forming two strongly sclerotized and dentated plates bordering the opening of the vesica. Female genitalia are only known in a limited number of species of *Hirpida* and were described only in Michener (1952) and Lemaire (2002); they are characterized by a very broad ventral plate resulting from the fusion of lamellae ante- and post-vaginalis; ductus bursae variable in length, opening in a broad ostium bursae. The corpus bursae is small and spherical, membranous, with the ductus seminalis arising mid-dorsally near the junction with the ductus. Nothing is known from the immature stages and food plant used by the species in this subfamily.

**Diagnosis:** Michener (1952) pointed at the slightly convex sides of the frons, the presence of spines on the ventral surface of tarsomeres, at the free juxta and at the articulated valvae as the main distinctive characters for the genus *Hirpida* within subfamily Hemileucinae. These characters are however known to be variable within Saturniidae and similar conditions are found in other subfamilies (e.g., convex sides of frons in members of the Attacini tribe (Arora & Gupta 1979, Rougerie 2005) or in Ceratocampinae (Balcazar-Lara & Wolfe, 1997); tarsomeres with spines in Arsenurinae, Agliinae, many Saturniinae, and with their presence considered to be plesiomorphic within Saturniidae (Michener 1952, Rougerie 2005)). Several other characters were mentioned by Michener (1952): the course of the mesothoracic anepisternal suture; the length of the tegulae; the visible frontal protuberance; well-developed arolia and pulvilli; the absence or reduction of vein 3A. They all are however insufficiently known throughout the family and further comparative morphology and a formal phylogenetic analysis of characters are needed to identify unique characters, if any, or more likely a unique combination of characters that would accurately define the subfamily Hirpidinae.

Antistathmopterini Rougerie, **tribus nova**

Type genus: *Antistathmoptera* Tams, 1935

**Description:** This tribe is defined for a single genus that includes only four described large-sized species, all restricted to Eastern Africa. The habitus of Antistathmopterini moths is unique, with extremely long hindwing tails, oval leaf-shaped forewings with pointed apex, wing patterns characterized by brown to reddish-brown ground color and a well-marked dark external line in forewings that contributes to the overall leaf-mimic aspect of the moths in resting posture; eyespots of both pairs of wings form a translucent spot composed of two adjacent hyaline areas, sometimes merged and always bigger in female specimens. Sexual dimorphism is low, but for shorter tails in female specimens and differences in antenna pectinations; male antennae are bipectinate with long and thin rami from base to tip of antennae, while females have much shorter rami. Labial palpi three-segmented. Tibial epiphysis of forelegs is present in both sexes. Forewing with three stalked Rs veins. Hindwing tail formed by veins M3, CuA1 and CuA2, with the former only reaching the tip of the tail, the other two reaching the basis or half of the twisted end of the tail. Anal vein (1A+2A) is short and stops at anal margin before the birth of the tail. The immature stages and biology of Antistathmopterini members remain unknown.

**Diagnosis:** Oberprieler (1997) emphasized that members of genus *Antistathmoptera* do not share any conspicuous character suggestive of phylogenetic affinities with any other saturniid moths. He considered that its placement within the tribe Urotini was mainly tentative since it only relied on the bipectinate condition of the antennae. Bouvier (1936) placed the genus along with *Eudaemonia* in a “macroure” section of his Pseudapheliini tribe, based on the sharing of tailed hindwings, but Oberprieler (1997) explicitly considered these affinities as unsupported by any other similarity and thus doubtful. Probably because of their rarity in natural history collections, little is known about the anatomical characters of these moths (see Testout (1941) for a superficial description of the genus *Antistathmoptera*) and further studies are needed to unveil additional diagnostic characters. General characters of the habitus however, in particular the tailed hindwings, pointed forewing and typical hyaline eyespots, make their identification unequivocal and Antistathmopterini moths cannot be confused with any other tailed saturniids in the Afrotropics (genera *Eudaemonia* and *Argema*).

Solini Rougerie, **tribus nova**

Type genus: *Solus* Watson, 1913

**Description:** This new tribe includes only the genus *Solus*, with seven described species (and three subspecies), all distributed in continental Asia. Small sized moths with triangular apically pointed forewings; pale brown in background color with well-marked ante-median, post-median and submarginal bands on both pairs of wings. On forewing, post-median band forming a straight line from the anal margin toward the apex; it is however curved toward the costal margin well before the apex, and the illusion of a continuous line to the wing-tip is caused by a strongly marked distal end of the submarginal line. This line contributes to the dead-leaf aspect of the moth that is also reinforced by several small patches of pale scales and small hyaline areas in and around the discal area of both forewings and hindwings. Sexual dimorphism is low, but for the more rounded shape of female wings and the slightly larger size of female individuals. Head with short antennae; galeae non-functional but present, short and broad. Forewing venation with vein d3 of discal cell missing, resulting in an open discal cell; R arising after the split between veins Rs4 and the merged veins Rs1+Rs2 and Rs3; lower branch of the fork at the start of forewing vein 1A+2A much thinner than the upper branch. The axillary sclerite 1 of forewing articulation with its tip rounded; anapleural suture of the mesothorax reduced; galeae present, short and broad. Tarsomeres of all legs lacking spines. Male genitalia short, stout with short valvae divided in a dorsal lobe and a ventral hook that are both curved toward the median axis. Uncus square-shaped ending distally in two small lobes oriented ventrally. Aedeagus with caecum penis undifferentiated; vesica with numerous cornuti forming a spiral of strongly sclerotized small teeth. Immature stages and biology unknown.

**Diagnosis:** Until now, the genus *Solus* has been treated as a member of tribe Saturniini, a position only questioned in Rougerie (2005) on the basis of a morphological phylogenetic analysis of the Saturniinae that placed *Solus* (along with genus *Cricula*) sister to Attacini + Saturniini. Our result here concurs with that finding for genus *Solus*, but suggests that its relationships with *Cricula* was superficial and based on morphological convergences (a result already pointed by Nässig (1989)) since the latter genus is nested within Saturniini in our analyses. We propose the following combination of characters in adults as diagnostic for this newly erected tribe: vein d3 of forewing discal cell missing, resulting in an open discal cell; lower branch of fork at the start of forewing vein 1A+2A much thinner than the upper branch; extremity of axillary sclerite 1 of forewings rounded; anapleural suture of mesothorax reduced; galeae present, short and broad.
