## Supplementary material for "Phylogenomics Illuminates the Evolutionary History of Wild Silkmoths in Space and Time (Lepidoptera: Saturniidae)": Figure S: FigureS2_trees_data_subsets_saturation.pdf

### Figure S2A. Data subset (20% least saturated loci) IQ-TREE unpartitioned

Best fit model = GTR+F+R10 chosen according to BIC
SH-aLRT /UFBoot

The new classification proposed in this manuscript is used to annotate tips.
Current classification is reported between square brackets

Nota : IQ-TREE analyses of data subsets were only performed without partitioning of the data to save computation time (and given that trees were similar when the initial data set was partitioned or not)

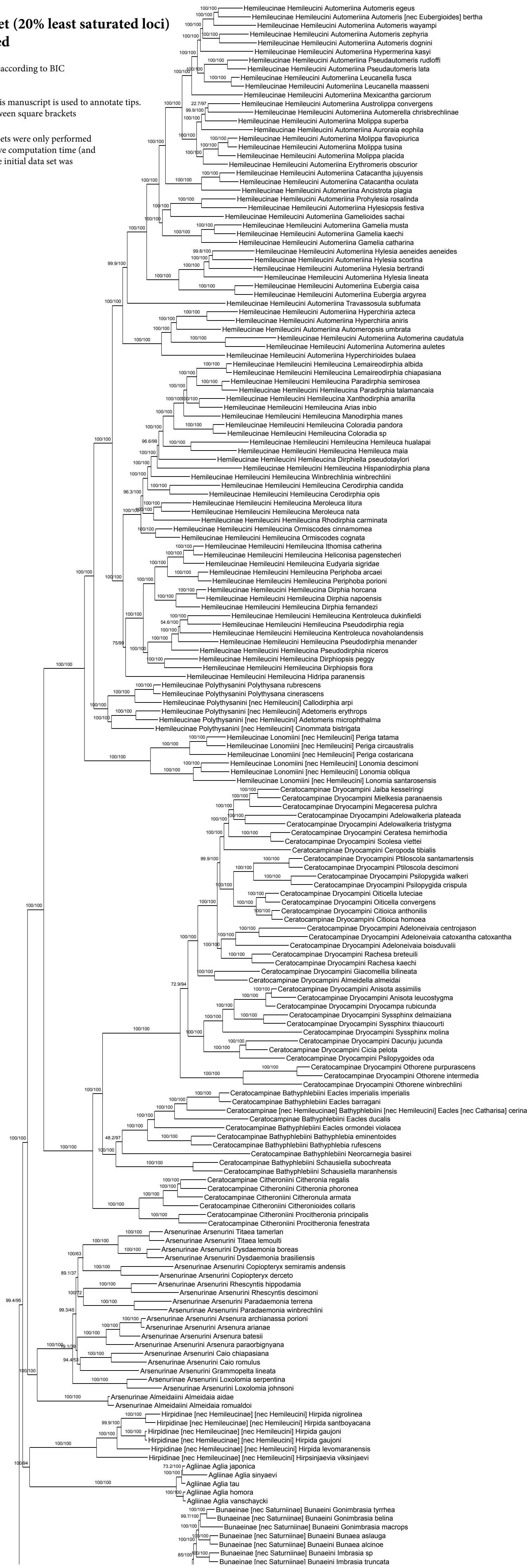

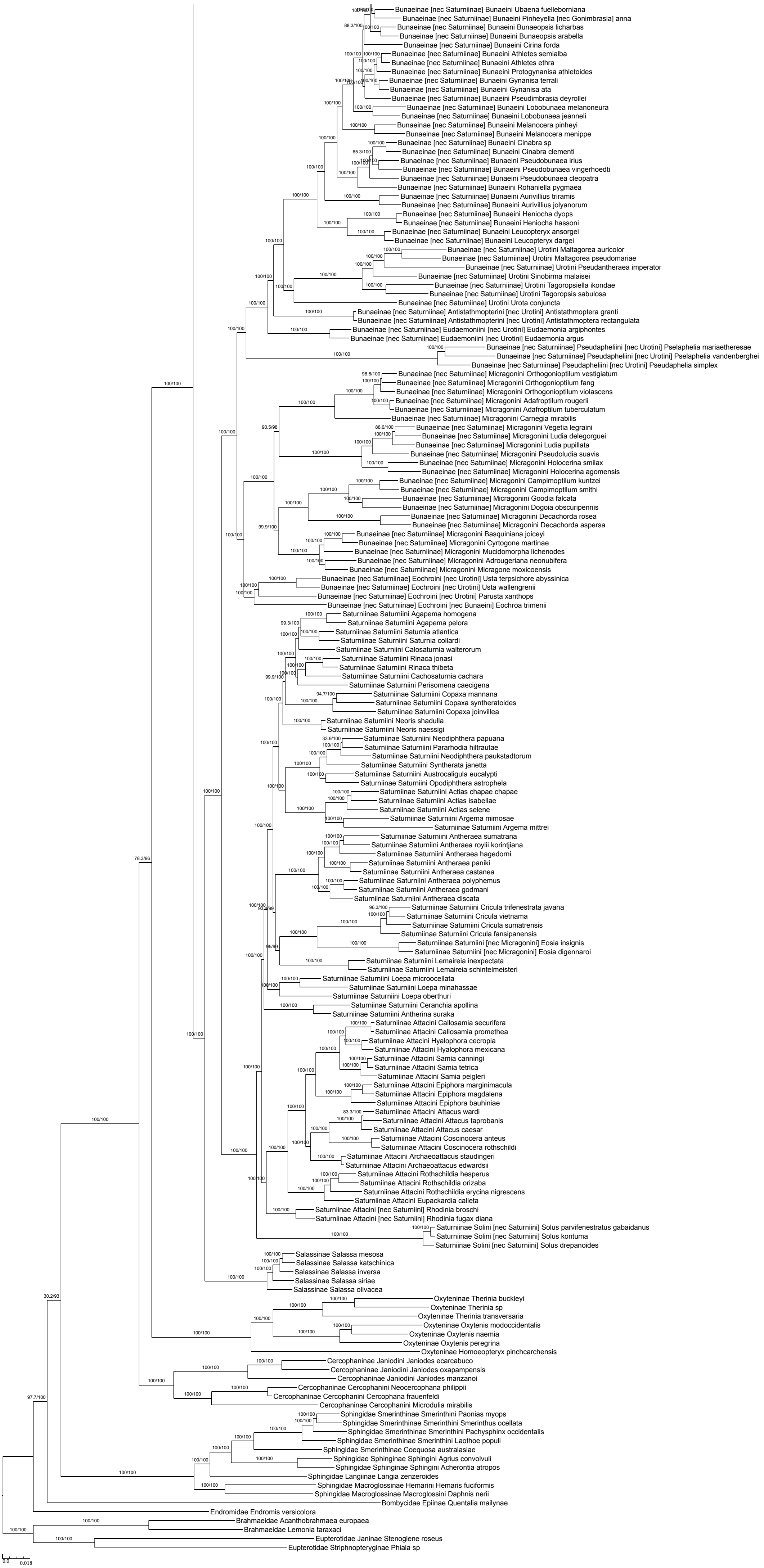

Bunaeinae [nec Saturniinae] Bunaeini Ubaena fuelleborniana
Bunaeinae [nec Saturniinae] Bunaeini Pinheyella [nec Gonimbrasia] anna
Bunaeinae [nec Saturniinae] Bunaeini Bunaeopsis licharbas
Bunaeinae [nec Saturniinae] Bunaeini Bunaeopsis arabella
Bunaeinae [nec Saturniinae] Bunaeini Cirina forda
Bunaeinae [nec Saturniinae] Bunaeini Athletes semialba
Bunaeinae [nec Saturniinae] Bunaeini Athletes ethra
Bunaeinae [nec Saturniinae] Bunaeini Protogynanisa athletoides
Bunaeinae [nec Saturniinae] Bunaeini Gynanisa terrali
Bunaeinae [nec Saturniinae] Bunaeini Gynanisa ata
Bunaeinae [nec Saturniinae] Bunaeini Pseudimbrasia deyrollei
Bunaeinae [nec Saturniinae] Bunaeini Lobobunaea melanoneura
Bunaeinae [nec Saturniinae] Bunaeini Lobobunaea jeanneli
Bunaeinae [nec Saturniinae] Bunaeini Melanocera pinheyi
Bunaeinae [nec Saturniinae] Bunaeini Melanocera menippe
Bunaeinae [nec Saturniinae] Bunaeini Cinabra sp
Bunaeinae [nec Saturniinae] Bunaeini Cinabra clementi
Bunaeinae [nec Saturniinae] Bunaeini Pseudobunaea irius
Bunaeinae [nec Saturniinae] Bunaeini Pseudobunaea vingerhoedti
Bunaeinae [nec Saturniinae] Bunaeini Pseudobunaea cleopatra
Bunaeinae [nec Saturniinae] Bunaeini Rohaniella pygmaea
Bunaeinae [nec Saturniinae] Bunaeini Aurivillius triramis
Bunaeinae [nec Saturniinae] Bunaeini Aurivillius jolyanorum
Bunaeinae [nec Saturniinae] Bunaeini Heniocha dyops
Bunaeinae [nec Saturniinae] Bunaeini Heniocha hassoni
Bunaeinae [nec Saturniinae] Bunaeini Leucopteryx ansorgei
Bunaeinae [nec Saturniinae] Bunaeini Leucopteryx dargei
Bunaeinae [nec Saturniinae] Urotini Maltagorea auricolor
Bunaeinae [nec Saturniinae] Urotini Maltagorea pseudomariae
Bunaeinae [nec Saturniinae] Urotini Pseudantheraea imperator
Bunaeinae [nec Saturniinae] Urotini Sinobirma malaisei
Bunaeinae [nec Saturniinae] Urotini Tagoropsiella ikondae
Bunaeinae [nec Saturniinae] Urotini Tagoropsis sabulosa
Bunaeinae [nec Saturniinae] Urotini Urota conjuncta
Bunaeinae [nec Saturniinae] Antistathmopterini [nec Urotini] Antistathmoptera granti
Bunaeinae [nec Saturniinae] Antistathmopterini [nec Urotini] Antistathmoptera rectangulata
Bunaeinae [nec Saturniinae] Eudaemoniini [nec Urotini] Eudaemonia argiphontes
Bunaeinae [nec Saturniinae] Eudaemoniini [nec Urotini] Eudaemonia argus
Bunaeinae [nec Saturniinae] Pseudapheliini [nec Urotini] Pselaphelia mariaetheresae
Bunaeinae [nec Saturniinae] Pseudapheliini [nec Urotini] Pselaphelia vandenberghei
Bunaeinae [nec Saturniinae] Pseudapheliini [nec Urotini] Pseudaphelia simplex
Bunaeinae [nec Saturniinae] Micragonini Orthogonioptilum vestigiatum
Bunaeinae [nec Saturniinae] Micragonini Orthogonioptilum fang
Bunaeinae [nec Saturniinae] Micragonini Orthogonioptilum violascens
Bunaeinae [nec Saturniinae] Micragonini Adafroptilum rougerii
Bunaeinae [nec Saturniinae] Micragonini Adafroptilum tuberculatum
Bunaeinae [nec Saturniinae] Micragonini Carnegia mirabilis
Bunaeinae [nec Saturniinae] Micragonini Vegetia legraini
Bunaeinae [nec Saturniinae] Micragonini Ludia delegorguei
Bunaeinae [nec Saturniinae] Micragonini Ludia pupillata
Bunaeinae [nec Saturniinae] Micragonini Pseudoludia suavis
Bunaeinae [nec Saturniinae] Micragonini Holocerina smilax
Bunaeinae [nec Saturniinae] Micragonini Holocerina agomensis
Bunaeinae [nec Saturniinae] Micragonini Campimoptilum kuntzei
Bunaeinae [nec Saturniinae] Micragonini Campimoptilum smithi
Bunaeinae [nec Saturniinae] Micragonini Goodia falcata
Bunaeinae [nec Saturniinae] Micragonini Dogoia obscuripennis
Bunaeinae [nec Saturniinae] Micragonini Decachorda rosea
Bunaeinae [nec Saturniinae] Micragonini Decachorda aspersa
Bunaeinae [nec Saturniinae] Micragonini Basquiniana joiceyi
Bunaeinae [nec Saturniinae] Micragonini Cyrtogone martinae
Bunaeinae [nec Saturniinae] Micragonini Mucidomorpha lichenodes
Bunaeinae [nec Saturniinae] Micragonini Adrougeriana neonubifera
Bunaeinae [nec Saturniinae] Micragonini Micragone moxicoensis
Bunaeinae [nec Saturniinae] Eochroini [nec Urotini] Usta terpsichore abyssinica
Bunaeinae [nec Saturniinae] Eochroini [nec Urotini] Usta wallengrenii
Bunaeinae [nec Saturniinae] Eochroini [nec Urotini] Parusta xanthops
Bunaeinae [nec Saturniinae] Eochroini [nec Bunaeini] Eochroa trimenii
Saturniinae Saturniini Agapema homogena
Saturniinae Saturniini Agapema pelora
Saturniinae Saturniini Saturnia atlantica
Saturniinae Saturniini Saturnia collardi
Saturniinae Saturniini Calosaturnia walterorum
Saturniinae Saturniini Rinaca jonasi
Saturniinae Saturniini Rinaca thibeta
Saturniinae Saturniini Cachosaturnia cachara
Saturniinae Saturniini Perisomena caecigena
Saturniinae Saturniini Copaxa mannana
Saturniinae Saturniini Copaxa syntheratoides
Saturniinae Saturniini Copaxa joinvillea
Saturniinae Saturniini Neoris shadulla
Saturniinae Saturniini Neoris naessigi
Saturniinae Saturniini Neodiphthera papuana
Saturniinae Saturniini Pararhodia hiltrautae
Saturniinae Saturniini Neodiphthera paukstadtorum
Saturniinae Saturniini Syntherata janetta
Saturniinae Saturniini Austrocaligula eucalypti
Saturniinae Saturniini Opodiphthera astrophela
Saturniinae Saturniini Actias chapae chapae
Saturniinae Saturniini Actias isabellae
Saturniinae Saturniini Actias selene
Saturniinae Saturniini Argema mimosae
Saturniinae Saturniini Argema mittrei
Saturniinae Saturniini Antheraea sumatrana
Saturniinae Saturniini Antheraea roylii korintjiana
Saturniinae Saturniini Antheraea hagedorni
Saturniinae Saturniini Antheraea paniki
Saturniinae Saturniini Antheraea castanea
Saturniinae Saturniini Antheraea polyphemus
Saturniinae Saturniini Antheraea godmani
Saturniinae Saturniini Antheraea discata
Saturniinae Saturniini Cricula trifenestrata javana
Saturniinae Saturniini Cricula vietnama
Saturniinae Saturniini Cricula sumatrensis
Saturniinae Saturniini Cricula fansipanensis
Saturniinae Saturniini [nec Micragonini] Eosia insignis
Saturniinae Saturniini [nec Micragonini] Eosia digennaroi
Saturniinae Saturniini Lemaireia inexpectata
Saturniinae Saturniini Lemaireia schintelmeisteri
Saturniinae Saturniini Loepa microocellata
Saturniinae Saturniini Loepa minahassae
Saturniinae Saturniini Loepa oberthuri
Saturniinae Saturniini Ceranchia apollina
Saturniinae Saturniini Antherina suraka
Saturniinae Attacini Callosamia securifera
Saturniinae Attacini Callosamia promethea
Saturniinae Attacini Hyalophora cecropia
Saturniinae Attacini Hyalophora mexicana
Saturniinae Attacini Samia canningi
Saturniinae Attacini Samia tetrica
Saturniinae Attacini Samia peigleri
Saturniinae Attacini Epiphora marginimacula
Saturniinae Attacini Epiphora magdalena
Saturniinae Attacini Epiphora bauhiniae
Saturniinae Attacini Attacus wardi
Saturniinae Attacini Attacus taprobanis
Saturniinae Attacini Attacus caesar
Saturniinae Attacini Coscinocera anteus
Saturniinae Attacini Coscinocera rothschildi
Saturniinae Attacini Archaeoattacus staudingeri
Saturniinae Attacini Archaeoattacus edwardsii
Saturniinae Attacini Rothschildia hesperus
Saturniinae Attacini Rothschildia orizaba
Saturniinae Attacini Rothschildia erycina nigrescens
Saturniinae Attacini Eupackardia calleta
Saturniinae Attacini [nec Saturniini] Rhodinia broschi
Saturniinae Attacini [nec Saturniini] Rhodinia fugax diana
Saturniinae Solini [nec Saturniini] Solus parvifenestratus gabaidanus
Saturniinae Solini [nec Saturniini] Solus kontuma
Saturniinae Solini [nec Saturniini] Solus drepanoides
Salassinae Salassa mesosa
Salassinae Salassa katschinica
Salassinae Salassa inversa
Salassinae Salassa siriae
Salassinae Salassa olivacea
Oxyteninae Therinia buckleyi
Oxyteninae Therinia sp
Oxyteninae Therinia transversaria
Oxyteninae Oxytenis modoccidentalis
Oxyteninae Oxytenis naemia
Oxyteninae Oxytenis peregrina
Oxyteninae Homoeopteryx pinchcarchensis
Cercophaninae Janiodini Janiodes ecarcabuco
Cercophaninae Janiodini Janiodes oxapampensis
Cercophaninae Janiodini Janiodes manzanoi
Cercophaninae Cercophanini Neocercophana philippii
Cercophaninae Cercophanini Cercophana frauenfeldi
Cercophaninae Cercophanini Microdulia mirabilis
Sphingidae Smerinthinae Smerinthini Paonias myops
Sphingidae Smerinthinae Smerinthini Smerinthus ocellata
Sphingidae Smerinthinae Smerinthini Pachysphinx occidentalis
Sphingidae Smerinthinae Smerinthini Laothoe populi
Sphingidae Smerinthinae Coequosa australasiae
Sphingidae Sphinginae Sphingini Agrius convolvuli
Sphingidae Sphinginae Sphingini Acherontia atropos
Sphingidae Langiinae Langia zenzeroides
Sphingidae Macroglossinae Hemarini Hemaris fuciformis
Sphingidae Macroglossinae Macroglossini Daphnis nerii
Bombycidae Epiinae Quentalia mailynae
Endromidae Endromis versicolora
Brahmaeidae Acanthobrahmaea europaea
Brahmaeidae Lemonia taraxaci
Eupterotidae Janinae Stenoglene roseus
Eupterotidae Striphnopteryginae Phiala sp
0.0
0.018

### Figure S2B. Data subset (20% least saturated loci) ASTRAL50

Species tree obtained when nodes with UFBoot<50 are collapsed in each gene tree

Final normalized quartet score : 0.95
Local posterior probabilities at nodes

The new classification proposed in this manuscript is used to annotate tips. Current classification is reported between square brackets

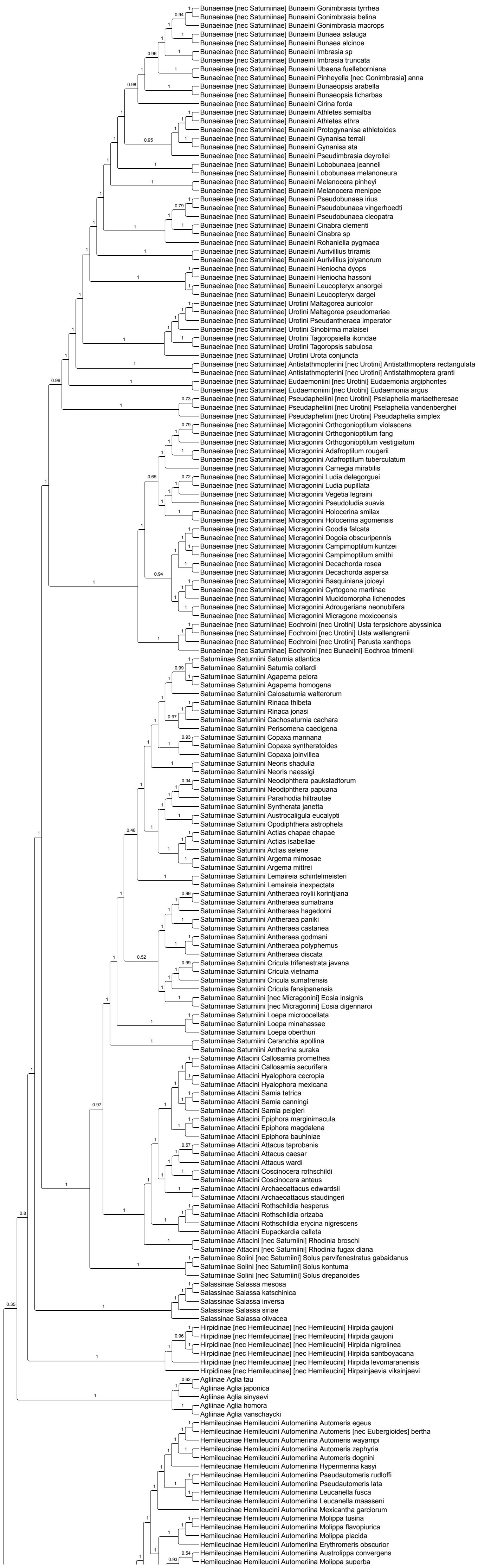

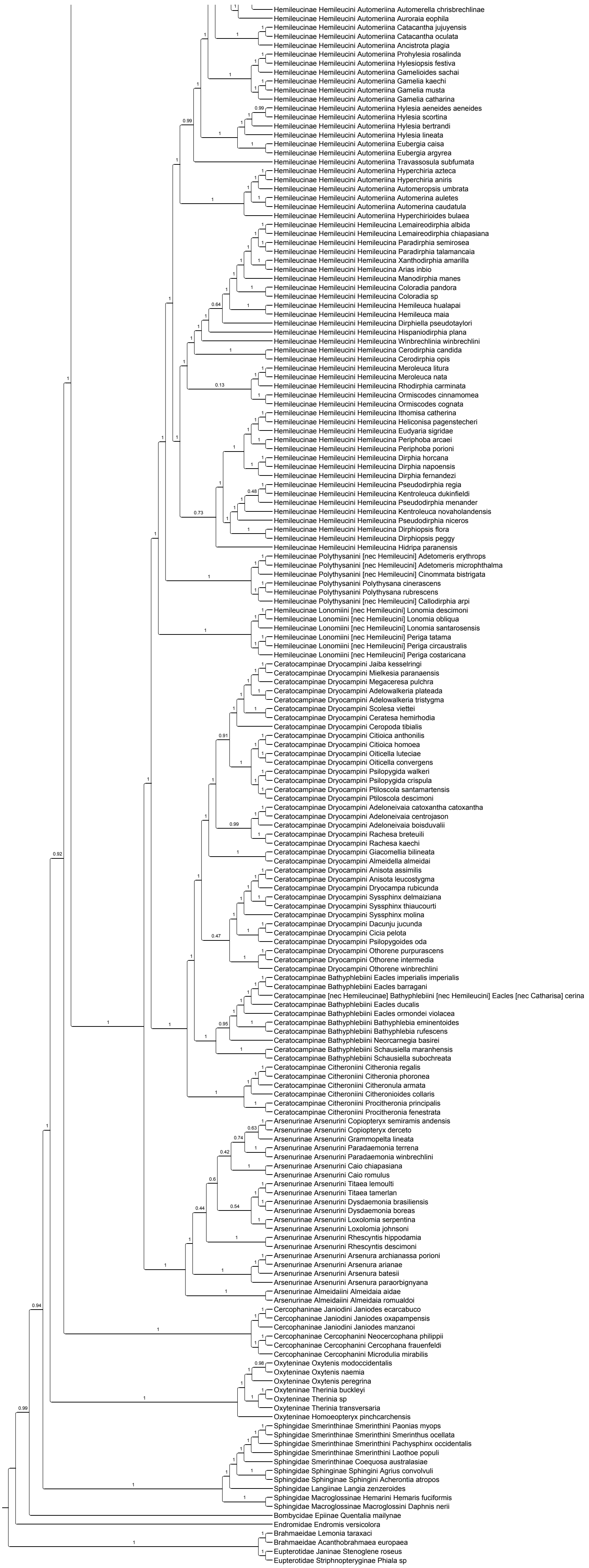

Hemileucinae Hemileucini Automeriina Automerella chrisbrechlinae
Hemileucinae Hemileucini Automeriina Auroraia eophila
Hemileucinae Hemileucini Automeriina Catacantha jujuyensis
Hemileucinae Hemileucini Automeriina Catacantha oculata
Hemileucinae Hemileucini Automeriina Ancistrota plagia
Hemileucinae Hemileucini Automeriina Prohylesia rosalinda
Hemileucinae Hemileucini Automeriina Hylesiopsis festiva
Hemileucinae Hemileucini Automeriina Gamelioides sachai
Hemileucinae Hemileucini Automeriina Gamelia kaechi
Hemileucinae Hemileucini Automeriina Gamelia musta
Hemileucinae Hemileucini Automeriina Gamelia catharina
Hemileucinae Hemileucini Automeriina Hylesia aeneides aeneides
Hemileucinae Hemileucini Automeriina Hylesia scortina
Hemileucinae Hemileucini Automeriina Hylesia bertrandi
Hemileucinae Hemileucini Automeriina Hylesia lineata
Hemileucinae Hemileucini Automeriina Eubergia caisa
Hemileucinae Hemileucini Automeriina Eubergia argyrea
Hemileucinae Hemileucini Automeriina Travassosula subfumata
Hemileucinae Hemileucini Automeriina Hyperchiria azteca
Hemileucinae Hemileucini Automeriina Hyperchiria aniris
Hemileucinae Hemileucini Automeriina Automeropsis umbrata
Hemileucinae Hemileucini Automeriina Automerina auletes
Hemileucinae Hemileucini Automeriina Automerina caudatula
Hemileucinae Hemileucini Automeriina Hyperchirioides bulaea
Hemileucinae Hemileucini Hemileucina Lemaireodirphia albida
Hemileucinae Hemileucini Hemileucina Lemaireodirphia chiapasiana
Hemileucinae Hemileucini Hemileucina Paradirphia semirosea
Hemileucinae Hemileucini Hemileucina Paradirphia talamancaia
Hemileucinae Hemileucini Hemileucina Xanthodirphia amarilla
Hemileucinae Hemileucini Hemileucina Arias inbio
Hemileucinae Hemileucini Hemileucina Manodirphia manes
Hemileucinae Hemileucini Hemileucina Coloradia pandora
Hemileucinae Hemileucini Hemileucina Coloradia sp
Hemileucinae Hemileucini Hemileucina Hemileuca hualapai
Hemileucinae Hemileucini Hemileucina Hemileuca maia
Hemileucinae Hemileucini Hemileucina Dirphiella pseudotaylori
Hemileucinae Hemileucini Hemileucina Hispaniodirphia plana
Hemileucinae Hemileucini Hemileucina Winbrechlinia winbrechlini
Hemileucinae Hemileucini Hemileucina Cerodirphia candida
Hemileucinae Hemileucini Hemileucina Cerodirphia opis
Hemileucinae Hemileucini Hemileucina Meroleuca litura
Hemileucinae Hemileucini Hemileucina Meroleuca nata
Hemileucinae Hemileucini Hemileucina Rhodirphia carminata
Hemileucinae Hemileucini Hemileucina Ormiscodes cinnamomea
Hemileucinae Hemileucini Hemileucina Ormiscodes cognata
Hemileucinae Hemileucini Hemileucina Ithomisa catherina
Hemileucinae Hemileucini Hemileucina Heliconisa pagenstecheri
Hemileucinae Hemileucini Hemileucina Eudyaria sigridae
Hemileucinae Hemileucini Hemileucina Periphoba arcaei
Hemileucinae Hemileucini Hemileucina Periphoba porioni
Hemileucinae Hemileucini Hemileucina Dirphia horcana
Hemileucinae Hemileucini Hemileucina Dirphia napoensis
Hemileucinae Hemileucini Hemileucina Dirphia fernandezi
Hemileucinae Hemileucini Hemileucina Pseudodirphia regia
Hemileucinae Hemileucini Hemileucina Kentroleuca dukinfieldi
Hemileucinae Hemileucini Hemileucina Pseudodirphia menander
Hemileucinae Hemileucini Hemileucina Kentroleuca novaholandensis
Hemileucinae Hemileucini Hemileucina Pseudodirphia niceros
Hemileucinae Hemileucini Hemileucina Dirphiopsis flora
Hemileucinae Hemileucini Hemileucina Dirphiopsis peggy
Hemileucinae Hemileucini Hemileucina Hidripa paranensis
Hemileucinae Polythysanini [nec Hemileucini] Adetomeris erythrops
Hemileucinae Polythysanini [nec Hemileucini] Adetomeris microphthalma
Hemileucinae Polythysanini [nec Hemileucini] Cinommata bistrigata
Hemileucinae Polythysanini Polythysana cinerascens
Hemileucinae Polythysanini Polythysana rubrescens
Hemileucinae Polythysanini [nec Hemileucini] Callodirphia arpi
Hemileucinae Lonomiini [nec Hemileucini] Lonomia descimoni
Hemileucinae Lonomiini [nec Hemileucini] Lonomia obliqua
Hemileucinae Lonomiini [nec Hemileucini] Lonomia santarosensis
Hemileucinae Lonomiini [nec Hemileucini] Periga tatama
Hemileucinae Lonomiini [nec Hemileucini] Periga circaustralis
Hemileucinae Lonomiini [nec Hemileucini] Periga costaricana
Ceratocampinae Dryocampini Jaiba kesselringi
Ceratocampinae Dryocampini Mielkesia paranaensis
Ceratocampinae Dryocampini Megaceresa pulchra
Ceratocampinae Dryocampini Adelowalkeria plateada
Ceratocampinae Dryocampini Adelowalkeria tristygma
Ceratocampinae Dryocampini Scolesa viettei
Ceratocampinae Dryocampini Ceratesa hemirhodia
Ceratocampinae Dryocampini Ceropoda tibialis
Ceratocampinae Dryocampini Citioica anthonilis
Ceratocampinae Dryocampini Citioica homoea
Ceratocampinae Dryocampini Oiticella luteciae
Ceratocampinae Dryocampini Oiticella convergens
Ceratocampinae Dryocampini Psilopygida walkeri
Ceratocampinae Dryocampini Psilopygida crispula
Ceratocampinae Dryocampini Ptiloscola santamartensis
Ceratocampinae Dryocampini Ptiloscola descimoni
Ceratocampinae Dryocampini Adeloneivaia catoxantha catoxantha
Ceratocampinae Dryocampini Adeloneivaia centrojason
Ceratocampinae Dryocampini Adeloneivaia boisduvalii
Ceratocampinae Dryocampini Rachesa breteuili
Ceratocampinae Dryocampini Rachesa kaechi
Ceratocampinae Dryocampini Giacomellia bilineata
Ceratocampinae Dryocampini Almeidella almeidai
Ceratocampinae Dryocampini Anisota assimilis
Ceratocampinae Dryocampini Anisota leucostygma
Ceratocampinae Dryocampini Dryocampa rubicunda
Ceratocampinae Dryocampini Syssphinx delmaiziana
Ceratocampinae Dryocampini Syssphinx thiaucourti
Ceratocampinae Dryocampini Syssphinx molina
Ceratocampinae Dryocampini Dacunju jucunda
Ceratocampinae Dryocampini Cicia pelota
Ceratocampinae Dryocampini Psilopygoides oda
Ceratocampinae Dryocampini Othorene purpurascens
Ceratocampinae Dryocampini Othorene intermedia
Ceratocampinae Dryocampini Othorene winbrechlini
Ceratocampinae Bathyphlebiini Eacles imperialis imperialis
Ceratocampinae Bathyphlebiini Eacles barragani
Ceratocampinae [nec Hemileucinae] Bathyphlebiini [nec Hemileucini] Eacles [nec Catharisa] cerina
Ceratocampinae Bathyphlebiini Eacles ducalis
Ceratocampinae Bathyphlebiini Eacles ormondei violacea
Ceratocampinae Bathyphlebiini Bathyphlebia eminentoides
Ceratocampinae Bathyphlebiini Bathyphlebia rufescens
Ceratocampinae Bathyphlebiini Neorcarnegia basirei
Ceratocampinae Bathyphlebiini Schausiella maranhensis
Ceratocampinae Bathyphlebiini Schausiella subochreata
Ceratocampinae Citheroniini Citheronia regalis
Ceratocampinae Citheroniini Citheronia phoronea
Ceratocampinae Citheroniini Citheronula armata
Ceratocampinae Citheroniini Citheronioides collaris
Ceratocampinae Citheroniini Procitheronia principalis
Ceratocampinae Citheroniini Procitheronia fenestrata
Arsenurinae Arsenurini Copiopteryx semiramis andensis
Arsenurinae Arsenurini Copiopteryx derceto
Arsenurinae Arsenurini Grammopelta lineata
Arsenurinae Arsenurini Paradaemonia terrena
Arsenurinae Arsenurini Paradaemonia winbrechlini
Arsenurinae Arsenurini Caio chiapasiana
Arsenurinae Arsenurini Caio romulus
Arsenurinae Arsenurini Titaea lemoulti
Arsenurinae Arsenurini Titaea tamerlan
Arsenurinae Arsenurini Dysdaemonia brasiliensis
Arsenurinae Arsenurini Dysdaemonia boreas
Arsenurinae Arsenurini Loxolomia serpentina
Arsenurinae Arsenurini Loxolomia johnsoni
Arsenurinae Arsenurini Rhescyntis hippodamia
Arsenurinae Arsenurini Rhescyntis descimoni
Arsenurinae Arsenurini Arsenura archianassa porioni
Arsenurinae Arsenurini Arsenura arianae
Arsenurinae Arsenurini Arsenura batesii
Arsenurinae Arsenurini Arsenura paraorbignyana
Arsenurinae Almeidaiini Almeidaia aidae
Arsenurinae Almeidaiini Almeidaia romualdoi
Cercophaninae Janiodini Janiodes ecarcabuco
Cercophaninae Janiodini Janiodes oxapampensis
Cercophaninae Janiodini Janiodes manzanoi
Cercophaninae Cercophanini Neocercophana philippii
Cercophaninae Cercophanini Cercophana frauenfeldi
Cercophaninae Cercophanini Microdulia mirabilis
Oxyteninae Oxytenis modoccidentalis
Oxyteninae Oxytenis naemia
Oxyteninae Oxytenis peregrina
Oxyteninae Therinia buckleyi
Oxyteninae Therinia sp
Oxyteninae Therinia transversaria
Oxyteninae Homoeopteryx pinchcarchensis
Sphingidae Smerinthinae Smerinthini Paonias myops
Sphingidae Smerinthinae Smerinthini Smerinthus ocellata
Sphingidae Smerinthinae Smerinthini Pachysphinx occidentalis
Sphingidae Smerinthinae Smerinthini Laothoe populi
Sphingidae Smerinthinae Coequosa australasiae
Sphingidae Sphinginae Sphingini Agrius convolvuli
Sphingidae Sphinginae Sphingini Acherontia atropos
Sphingidae Langiinae Langia zenzeroides
Sphingidae Macroglossinae Hemarini Hemaris fuciformis
Sphingidae Macroglossinae Macroglossini Daphnis nerii
Bombycidae Epiinae Quentalia mailynae
Endromidae Endromis versicolora
Brahmaeidae Lemonia taraxaci
Brahmaeidae Acanthobrahmaea europaea
Eupterotidae Janinae Stenoglene roseus
Eupterotidae Striphnopteryginae Phiala sp

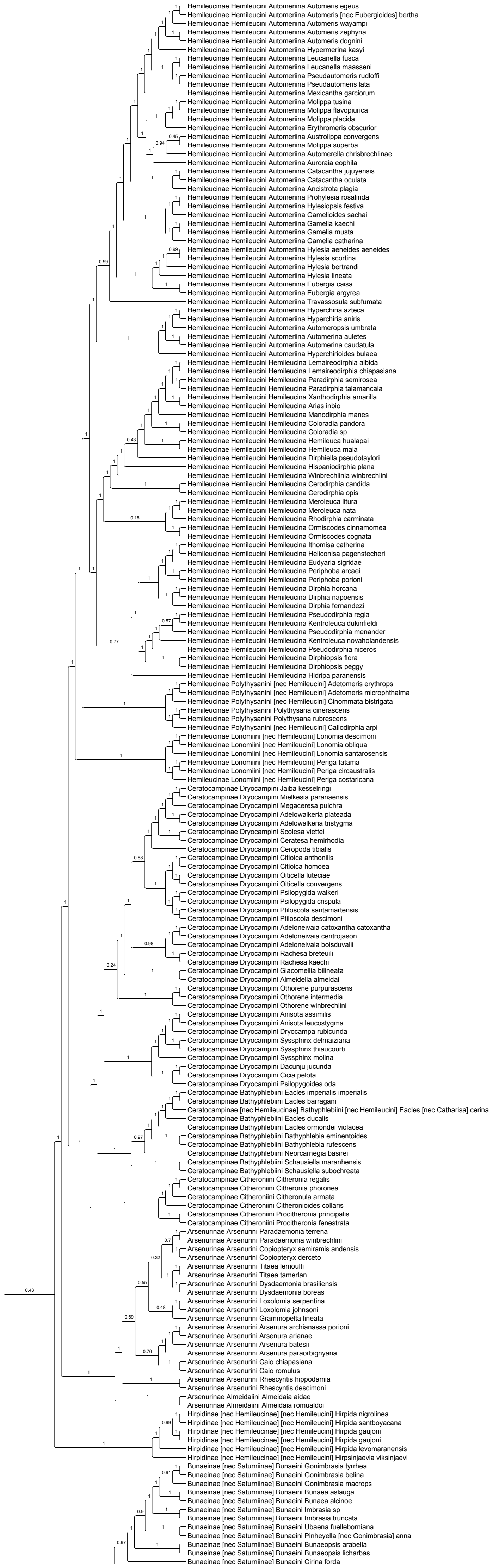


#### Figure S2C. Data subset (20% least saturated loci) ASTRAL70

Species tree obtained when nodes with UFBoot<70 are collapsed in each gene tree

Final normalized quartet score : 0.97
Local posterior probabilities at nodes

The new classification proposed in this manuscript is used to annotate tips. Current classification is reported between square brackets

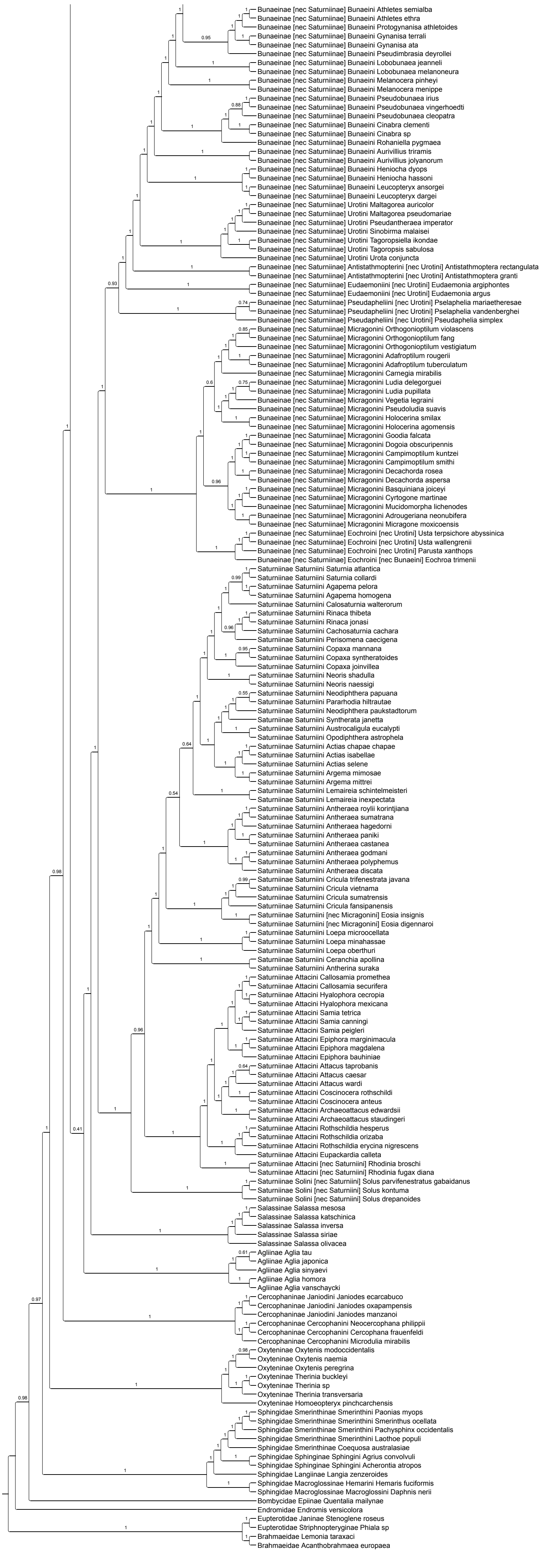
Bunaeinae [nec Saturniinae] Bunaeini Athletes semialba
Bunaeinae [nec Saturniinae] Bunaeini Athletes ethra
Bunaeinae [nec Saturniinae] Bunaeini Protogynanisa athletoides
Bunaeinae [nec Saturniinae] Bunaeini Gynanisa terrali
Bunaeinae [nec Saturniinae] Bunaeini Gynanisa ata
Bunaeinae [nec Saturniinae] Bunaeini Pseudimbrasia deyrollei
Bunaeinae [nec Saturniinae] Bunaeini Lobobunaea jeanneli
Bunaeinae [nec Saturniinae] Bunaeini Lobobunaea melanoneura
Bunaeinae [nec Saturniinae] Bunaeini Melanocera pinheyi
Bunaeinae [nec Saturniinae] Bunaeini Melanocera menippe
Bunaeinae [nec Saturniinae] Bunaeini Pseudobunaea irius
Bunaeinae [nec Saturniinae] Bunaeini Pseudobunaea vingerhoedti
Bunaeinae [nec Saturniinae] Bunaeini Pseudobunaea cleopatra
Bunaeinae [nec Saturniinae] Bunaeini Cinabra clementi
Bunaeinae [nec Saturniinae] Bunaeini Cinabra sp
Bunaeinae [nec Saturniinae] Bunaeini Rohaniella pygmaea
Bunaeinae [nec Saturniinae] Bunaeini Aurivillius triramis
Bunaeinae [nec Saturniinae] Bunaeini Aurivillius jolyanorum
Bunaeinae [nec Saturniinae] Bunaeini Heniocha dyops
Bunaeinae [nec Saturniinae] Bunaeini Heniocha hassoni
Bunaeinae [nec Saturniinae] Bunaeini Leucopteryx ansorgei
Bunaeinae [nec Saturniinae] Bunaeini Leucopteryx dargei
Bunaeinae [nec Saturniinae] Urotini Maltagorea auricolor
Bunaeinae [nec Saturniinae] Urotini Maltagorea pseudomariae
Bunaeinae [nec Saturniinae] Urotini Pseudantheraea imperator
Bunaeinae [nec Saturniinae] Urotini Sinobirma malaisei
Bunaeinae [nec Saturniinae] Urotini Tagoropsiella ikondae
Bunaeinae [nec Saturniinae] Urotini Tagoropsis sabulosa
Bunaeinae [nec Saturniinae] Urotini Urota conjuncta
Bunaeinae [nec Saturniinae] Antistathmopterini [nec Urotini] Antistathmoptera rectangulata
Bunaeinae [nec Saturniinae] Antistathmopterini [nec Urotini] Antistathmoptera granti
Bunaeinae [nec Saturniinae] Eudaemoniini [nec Urotini] Eudaemonia argiphontes
Bunaeinae [nec Saturniinae] Eudaemoniini [nec Urotini] Eudaemonia argus
Bunaeinae [nec Saturniinae] Pseudapheliini [nec Urotini] Pselaphelia mariaetheresae
Bunaeinae [nec Saturniinae] Pseudapheliini [nec Urotini] Pselaphelia vandenberghei
Bunaeinae [nec Saturniinae] Pseudapheliini [nec Urotini] Pseudaphelia simplex
Bunaeinae [nec Saturniinae] Micragonini Orthogonioptilum violascens
Bunaeinae [nec Saturniinae] Micragonini Orthogonioptilum fang
Bunaeinae [nec Saturniinae] Micragonini Orthogonioptilum vestigiatum
Bunaeinae [nec Saturniinae] Micragonini Adafroptilum rougerii
Bunaeinae [nec Saturniinae] Micragonini Adafroptilum tuberculatum
Bunaeinae [nec Saturniinae] Micragonini Carnegia mirabilis
Bunaeinae [nec Saturniinae] Micragonini Ludia delegorguei
Bunaeinae [nec Saturniinae] Micragonini Ludia pupillata
Bunaeinae [nec Saturniinae] Micragonini Vegetia legraini
Bunaeinae [nec Saturniinae] Micragonini Pseudoludia suavis
Bunaeinae [nec Saturniinae] Micragonini Holocerina smilax
Bunaeinae [nec Saturniinae] Micragonini Holocerina agomensis
Bunaeinae [nec Saturniinae] Micragonini Goodia falcata
Bunaeinae [nec Saturniinae] Micragonini Dogoia obscuripennis
Bunaeinae [nec Saturniinae] Micragonini Campimoptilum kuntzei
Bunaeinae [nec Saturniinae] Micragonini Campimoptilum smithi
Bunaeinae [nec Saturniinae] Micragonini Decachorda rosea
Bunaeinae [nec Saturniinae] Micragonini Decachorda aspersa
Bunaeinae [nec Saturniinae] Micragonini Basquiniana joiceyi
Bunaeinae [nec Saturniinae] Micragonini Cyrtogone martinae
Bunaeinae [nec Saturniinae] Micragonini Mucidomorpha lichenodes
Bunaeinae [nec Saturniinae] Micragonini Adrougeriana neonubifera
Bunaeinae [nec Saturniinae] Micragonini Micragone moxicoensis
Bunaeinae [nec Saturniinae] Eochroini [nec Urotini] Usta terpsichore abyssinica
Bunaeinae [nec Saturniinae] Eochroini [nec Urotini] Usta wallengrenii
Bunaeinae [nec Saturniinae] Eochroini [nec Urotini] Parusta xanthops
Bunaeinae [nec Saturniinae] Eochroini [nec Bunaeini] Eochroa trimenii
Saturniinae Saturniini Saturnia atlantica
Saturniinae Saturniini Saturnia collardi
Saturniinae Saturniini Agapema pelora
Saturniinae Saturniini Agapema homogena
Saturniinae Saturniini Calosaturnia walterorum
Saturniinae Saturniini Rinaca thibeta
Saturniinae Saturniini Rinaca jonasi
Saturniinae Saturniini Cachosaturnia cachara
Saturniinae Saturniini Perisomena caecigena
Saturniinae Saturniini Copaxa mannana
Saturniinae Saturniini Copaxa syntheratoides
Saturniinae Saturniini Copaxa joinvillea
Saturniinae Saturniini Neoris shadulla
Saturniinae Saturniini Neoris naessigi
Saturniinae Saturniini Neodiphthera papuana
Saturniinae Saturniini Pararhodia hiltrautae
Saturniinae Saturniini Neodiphthera paukstadtorum
Saturniinae Saturniini Syntherata janetta
Saturniinae Saturniini Austrocaligula eucalypti
Saturniinae Saturniini Opodiphthera astrophela
Saturniinae Saturniini Actias chapae chapae
Saturniinae Saturniini Actias isabellae
Saturniinae Saturniini Actias selene
Saturniinae Saturniini Argema mimosae
Saturniinae Saturniini Argema mittrei
Saturniinae Saturniini Lemaireia schintelmeisteri
Saturniinae Saturniini Lemaireia inexpectata
Saturniinae Saturniini Antheraea roylii korintjiana
Saturniinae Saturniini Antheraea sumatrana
Saturniinae Saturniini Antheraea hagedorni
Saturniinae Saturniini Antheraea paniki
Saturniinae Saturniini Antheraea castanea
Saturniinae Saturniini Antheraea godmani
Saturniinae Saturniini Antheraea polyphemus
Saturniinae Saturniini Antheraea discata
Saturniinae Saturniini Cricula trifenestrata javana
Saturniinae Saturniini Cricula vietnama
Saturniinae Saturniini Cricula sumatrensis
Saturniinae Saturniini Cricula fansipanensis
Saturniinae Saturniini [nec Micragonini] Eosia insignis
Saturniinae Saturniini [nec Micragonini] Eosia digennaroi
Saturniinae Saturniini Loepa microocellata
Saturniinae Saturniini Loepa minahassae
Saturniinae Saturniini Loepa oberthuri
Saturniinae Saturniini Ceranchia apollina
Saturniinae Saturniini Antherina suraka
Saturniinae Attacini Callosamia promethea
Saturniinae Attacini Callosamia securifera
Saturniinae Attacini Hyalophora cecropia
Saturniinae Attacini Hyalophora mexicana
Saturniinae Attacini Samia tetrica
Saturniinae Attacini Samia canningi
Saturniinae Attacini Samia peigleri
Saturniinae Attacini Epiphora marginimacula
Saturniinae Attacini Epiphora magdalena
Saturniinae Attacini Epiphora bauhiniae
Saturniinae Attacini Attacus taprobanis
Saturniinae Attacini Attacus caesar
Saturniinae Attacini Attacus wardi
Saturniinae Attacini Coscinocera rothschildi
Saturniinae Attacini Coscinocera anteus
Saturniinae Attacini Archaeoattacus edwardsii
Saturniinae Attacini Archaeoattacus staudingeri
Saturniinae Attacini Rothschildia hesperus
Saturniinae Attacini Rothschildia orizaba
Saturniinae Attacini Rothschildia erycina nigrescens
Saturniinae Attacini Eupackardia calleta
Saturniinae Attacini [nec Saturniini] Rhodinia broschi
Saturniinae Attacini [nec Saturniini] Rhodinia fugax diana
Saturniinae Solini [nec Saturniini] Solus parvifenestratus gabaidanus
Saturniinae Solini [nec Saturniini] Solus kontuma
Saturniinae Solini [nec Saturniini] Solus drepanoides
Salassinae Salassa mesosa
Salassinae Salassa katschinica
Salassinae Salassa inversa
Salassinae Salassa siriae
Salassinae Salassa olivacea
Agliinae Aglia tau
Agliinae Aglia japonica
Agliinae Aglia sinyaevi
Agliinae Aglia homora
Agliinae Aglia vanschaycki
Cercophaninae Janiodini Janiodes ecarcabuco
Cercophaninae Janiodini Janiodes oxapampensis
Cercophaninae Janiodini Janiodes manzanoi
Cercophaninae Cercophanini Neocercophana philippii
Cercophaninae Cercophanini Cercophana frauenfeldi
Cercophaninae Cercophanini Microdulia mirabilis
Oxyteninae Oxytenis modoccidentalis
Oxyteninae Oxytenis naemia
Oxyteninae Oxytenis peregrina
Oxyteninae Therinia buckleyi
Oxyteninae Therinia sp
Oxyteninae Therinia transversaria
Oxyteninae Homoeopteryx pinchcarchensis
Sphingidae Smerinthinae Smerinthini Paonias myops
Sphingidae Smerinthinae Smerinthini Smerinthus ocellata
Sphingidae Smerinthinae Smerinthini Pachysphinx occidentalis
Sphingidae Smerinthinae Smerinthini Laothoe populi
Sphingidae Smerinthinae Coequosa australasiae
Sphingidae Sphinginae Sphingini Agrius convolvuli
Sphingidae Sphinginae Sphingini Acherontia atropos
Sphingidae Langiinae Langia zenzeroides
Sphingidae Macroglossinae Hemarini Hemaris fuciformis
Sphingidae Macroglossinae Macroglossini Daphnis nerii
Bombycidae Epiinae Quentalia mailynae
Endromidae Endromis versicolora
Eupterotidae Janinae Stenoglene roseus
Eupterotidae Striphnopteryginae Phiala sp
Brahmaeidae Lemonia taraxaci
Brahmaeidae Acanthobrahmaea europaea

### Figure S2D. Data subset (20% least saturated loci) ASTRAL90

Species tree obtained when nodes with UFBoot<90 are collapsed in each gene tree

Final normalized quartet score : 0.99
Local posterior probabilities at nodes

The new classification proposed in this manuscript is used to annotate tips. Current classification is reported between square brackets

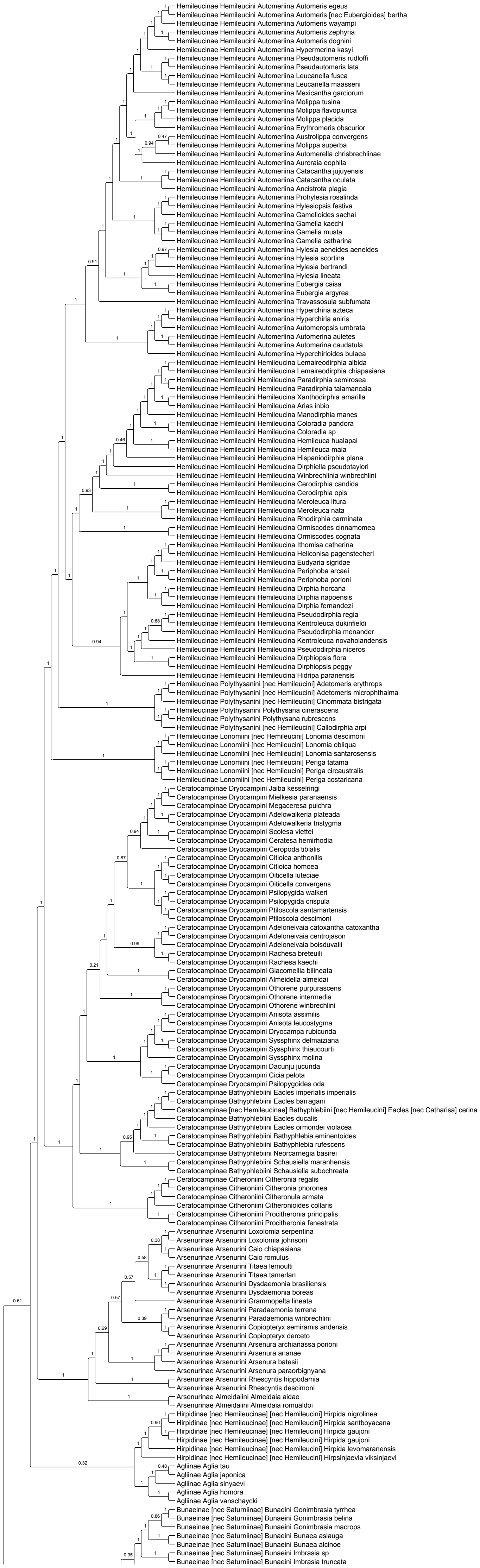

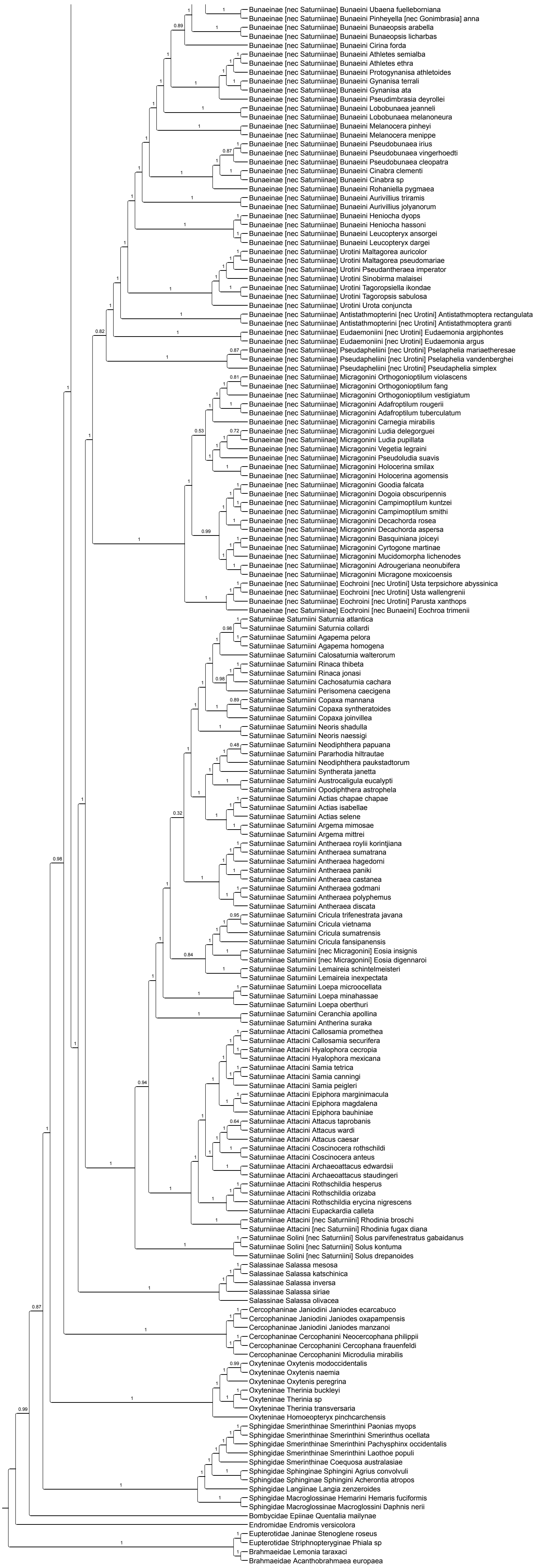

Bunaeinae [nec Saturniinae] Bunaeini Ubaena fuelleborniana
Bunaeinae [nec Saturniinae] Bunaeini Pinheyella [nec Gonimbrasia] anna
Bunaeinae [nec Saturniinae] Bunaeini Bunaeopsis arabella
Bunaeinae [nec Saturniinae] Bunaeini Bunaeopsis licharbas
Bunaeinae [nec Saturniinae] Bunaeini Cirina forda
Bunaeinae [nec Saturniinae] Bunaeini Athletes semialba
Bunaeinae [nec Saturniinae] Bunaeini Athletes ethra
Bunaeinae [nec Saturniinae] Bunaeini Protogynanisa athletoides
Bunaeinae [nec Saturniinae] Bunaeini Gynanisa terrali
Bunaeinae [nec Saturniinae] Bunaeini Gynanisa ata
Bunaeinae [nec Saturniinae] Bunaeini Pseudimbrasia deyrollei
Bunaeinae [nec Saturniinae] Bunaeini Lobobunaea jeanneli
Bunaeinae [nec Saturniinae] Bunaeini Lobobunaea melanoneura
Bunaeinae [nec Saturniinae] Bunaeini Melanocera pinheyi
Bunaeinae [nec Saturniinae] Bunaeini Melanocera menippe
Bunaeinae [nec Saturniinae] Bunaeini Pseudobunaea irius
Bunaeinae [nec Saturniinae] Bunaeini Pseudobunaea vingerhoedti
Bunaeinae [nec Saturniinae] Bunaeini Pseudobunaea cleopatra
Bunaeinae [nec Saturniinae] Bunaeini Cinabra clementi
Bunaeinae [nec Saturniinae] Bunaeini Cinabra sp
Bunaeinae [nec Saturniinae] Bunaeini Rohaniella pygmaea
Bunaeinae [nec Saturniinae] Bunaeini Aurivillius triramis
Bunaeinae [nec Saturniinae] Bunaeini Aurivillius jolyanorum
Bunaeinae [nec Saturniinae] Bunaeini Heniocha dyops
Bunaeinae [nec Saturniinae] Bunaeini Heniocha hassoni
Bunaeinae [nec Saturniinae] Bunaeini Leucopteryx ansorgei
Bunaeinae [nec Saturniinae] Bunaeini Leucopteryx dargei
Bunaeinae [nec Saturniinae] Urotini Maltagorea auricolor
Bunaeinae [nec Saturniinae] Urotini Maltagorea pseudomariae
Bunaeinae [nec Saturniinae] Urotini Pseudantheraea imperator
Bunaeinae [nec Saturniinae] Urotini Sinobirma malaisei
Bunaeinae [nec Saturniinae] Urotini Tagoropsiella ikondae
Bunaeinae [nec Saturniinae] Urotini Tagoropsis sabulosa
Bunaeinae [nec Saturniinae] Urotini Urota conjuncta
Bunaeinae [nec Saturniinae] Antistathmopterini [nec Urotini] Antistathmoptera rectangulata
Bunaeinae [nec Saturniinae] Antistathmopterini [nec Urotini] Antistathmoptera granti
Bunaeinae [nec Saturniinae] Eudaemoniini [nec Urotini] Eudaemonia argiphontes
Bunaeinae [nec Saturniinae] Eudaemoniini [nec Urotini] Eudaemonia argus
Bunaeinae [nec Saturniinae] Pseudapheliini [nec Urotini] Pselaphelia mariaetheresae
Bunaeinae [nec Saturniinae] Pseudapheliini [nec Urotini] Pselaphelia vandenberghei
Bunaeinae [nec Saturniinae] Pseudapheliini [nec Urotini] Pseudaphelia simplex
Bunaeinae [nec Saturniinae] Micragonini Orthogonioptilum violascens
Bunaeinae [nec Saturniinae] Micragonini Orthogonioptilum fang
Bunaeinae [nec Saturniinae] Micragonini Orthogonioptilum vestigiatum
Bunaeinae [nec Saturniinae] Micragonini Adafroptilum rougerii
Bunaeinae [nec Saturniinae] Micragonini Adafroptilum tuberculatum
Bunaeinae [nec Saturniinae] Micragonini Carnegia mirabilis
Bunaeinae [nec Saturniinae] Micragonini Ludia delegorguei
Bunaeinae [nec Saturniinae] Micragonini Ludia pupillata
Bunaeinae [nec Saturniinae] Micragonini Vegetia legraini
Bunaeinae [nec Saturniinae] Micragonini Pseudoludia suavis
Bunaeinae [nec Saturniinae] Micragonini Holocerina smilax
Bunaeinae [nec Saturniinae] Micragonini Holocerina agomensis
Bunaeinae [nec Saturniinae] Micragonini Goodia falcata
Bunaeinae [nec Saturniinae] Micragonini Dogoia obscuripennis
Bunaeinae [nec Saturniinae] Micragonini Campimoptilum kuntzei
Bunaeinae [nec Saturniinae] Micragonini Campimoptilum smithi
Bunaeinae [nec Saturniinae] Micragonini Decachorda rosea
Bunaeinae [nec Saturniinae] Micragonini Decachorda aspersa
Bunaeinae [nec Saturniinae] Micragonini Basquiniana joiceyi
Bunaeinae [nec Saturniinae] Micragonini Cyrtogone martinae
Bunaeinae [nec Saturniinae] Micragonini Mucidomorpha lichenodes
Bunaeinae [nec Saturniinae] Micragonini Adrougeriana neonubifera
Bunaeinae [nec Saturniinae] Micragonini Micragone moxicoensis
Bunaeinae [nec Saturniinae] Eochroini [nec Urotini] Usta terpsichore abyssinica
Bunaeinae [nec Saturniinae] Eochroini [nec Urotini] Usta wallengrenii
Bunaeinae [nec Saturniinae] Eochroini [nec Urotini] Parusta xanthops
Bunaeinae [nec Saturniinae] Eochroini [nec Bunaeini] Eochroa trimenii
Saturniinae Saturniini Saturnia atlantica
Saturniinae Saturniini Saturnia collardi
Saturniinae Saturniini Agapema pelora
Saturniinae Saturniini Agapema homogena
Saturniinae Saturniini Calosaturnia walterorum
Saturniinae Saturniini Rinaca thibeta
Saturniinae Saturniini Rinaca jonasi
Saturniinae Saturniini Cachosaturnia cachara
Saturniinae Saturniini Perisomena caecigena
Saturniinae Saturniini Copaxa mannana
Saturniinae Saturniini Copaxa syntheratoides
Saturniinae Saturniini Copaxa joinvillea
Saturniinae Saturniini Neoris shadulla
Saturniinae Saturniini Neoris naessigi
Saturniinae Saturniini Neodiphthera papuana
Saturniinae Saturniini Pararhodia hiltrautae
Saturniinae Saturniini Neodiphthera paukstadtorum
Saturniinae Saturniini Syntherata janetta
Saturniinae Saturniini Austrocaligula eucalypti
Saturniinae Saturniini Opodiphthera astrophela
Saturniinae Saturniini Actias chapae chapae
Saturniinae Saturniini Actias isabellae
Saturniinae Saturniini Actias selene
Saturniinae Saturniini Argema mimosae
Saturniinae Saturniini Argema mittrei
Saturniinae Saturniini Antheraea roylii korintjiana
Saturniinae Saturniini Antheraea sumatrana
Saturniinae Saturniini Antheraea hagedorni
Saturniinae Saturniini Antheraea paniki
Saturniinae Saturniini Antheraea castanea
Saturniinae Saturniini Antheraea godmani
Saturniinae Saturniini Antheraea polyphemus
Saturniinae Saturniini Antheraea discata
Saturniinae Saturniini Cricula trifenestrata javana
Saturniinae Saturniini Cricula vietnama
Saturniinae Saturniini Cricula sumatrensis
Saturniinae Saturniini Cricula fansipanensis
Saturniinae Saturniini [nec Micragonini] Eosia insignis
Saturniinae Saturniini [nec Micragonini] Eosia digennaroi
Saturniinae Saturniini Lemaireia schintelmeisteri
Saturniinae Saturniini Lemaireia inexpectata
Saturniinae Saturniini Loepa microocellata
Saturniinae Saturniini Loepa minahassae
Saturniinae Saturniini Loepa oberthuri
Saturniinae Saturniini Ceranchia apollina
Saturniinae Saturniini Antherina suraka
Saturniinae Attacini Callosamia promethea
Saturniinae Attacini Callosamia securifera
Saturniinae Attacini Hyalophora cecropia
Saturniinae Attacini Hyalophora mexicana
Saturniinae Attacini Samia tetrica
Saturniinae Attacini Samia canningi
Saturniinae Attacini Samia peigleri
Saturniinae Attacini Epiphora marginimacula
Saturniinae Attacini Epiphora magdalena
Saturniinae Attacini Epiphora bauhiniae
Saturniinae Attacini Attacus taprobanis
Saturniinae Attacini Attacus wardi
Saturniinae Attacini Attacus caesar
Saturniinae Attacini Coscinocera rothschildi
Saturniinae Attacini Coscinocera anteus
Saturniinae Attacini Archaeoattacus edwardsii
Saturniinae Attacini Archaeoattacus staudingeri
Saturniinae Attacini Rothschildia hesperus
Saturniinae Attacini Rothschildia orizaba
Saturniinae Attacini Rothschildia erycina nigrescens
Saturniinae Attacini Eupackardia calleta
Saturniinae Attacini [nec Saturniini] Rhodinia broschi
Saturniinae Attacini [nec Saturniini] Rhodinia fugax diana
Saturniinae Solini [nec Saturniini] Solus parvifenestratus gabaidanus
Saturniinae Solini [nec Saturniini] Solus kontuma
Saturniinae Solini [nec Saturniini] Solus drepanoides
Salassinae Salassa mesosa
Salassinae Salassa katschinica
Salassinae Salassa inversa
Salassinae Salassa siriae
Salassinae Salassa olivacea
Cercophaninae Janiodini Janiodes ecarcabuco
Cercophaninae Janiodini Janiodes oxapampensis
Cercophaninae Janiodini Janiodes manzanoi
Cercophaninae Cercophanini Neocercophana philippii
Cercophaninae Cercophanini Cercophana frauenfeldi
Cercophaninae Cercophanini Microdulia mirabilis
Oxyteninae Oxytenis modocccidentalis
Oxyteninae Oxytenis naemia
Oxyteninae Oxytenis peregrina
Oxyteninae Therinia buckleyi
Oxyteninae Therinia sp
Oxyteninae Therinia transversaria
Oxyteninae Homoeopteryx pinchcarchensis
Sphingidae Smerinthinae Smerinthini Paonias myops
Sphingidae Smerinthinae Smerinthini Smerinthus ocellata
Sphingidae Smerinthinae Smerinthini Pachysphinx occidentalis
Sphingidae Smerinthinae Smerinthini Laothoe populi
Sphingidae Smerinthinae Coequosa australasiae
Sphingidae Sphinginae Sphingini Agrius convolvuli
Sphingidae Sphinginae Sphingini Acherontia atropos
Sphingidae Langiinae Langia zenzeroides
Sphingidae Macroglossinae Hemarini Hemaris fuciformis
Sphingidae Macroglossinae Macroglossini Daphnis nerii
Bombycidae Epiinae Quentalia mailynae
Endromidae Endromis versicolora
Eupterotidae Janinae Stenoglene roseus
Eupterotidae Striphnopteryginae Phiala sp
Brahmaeidae Lemonia taraxaci
Brahmaeidae Acanthobrahmaea europaea

#### Figure S2E. Data subset (20% most saturated loci) IQ-TREE unpartitioned

Best fit model = GTR+F+R10 chosen according to BIC
SH-aLRT /UFBoot

The new classification proposed in this manuscript is used to annotate tips. Current classification is reported between square brackets

Nota : IQ-TREE analyses of data subsets were only performed without partitioning of the data to save computation time (and given that trees were similar when the initial data set was partitioned or not)

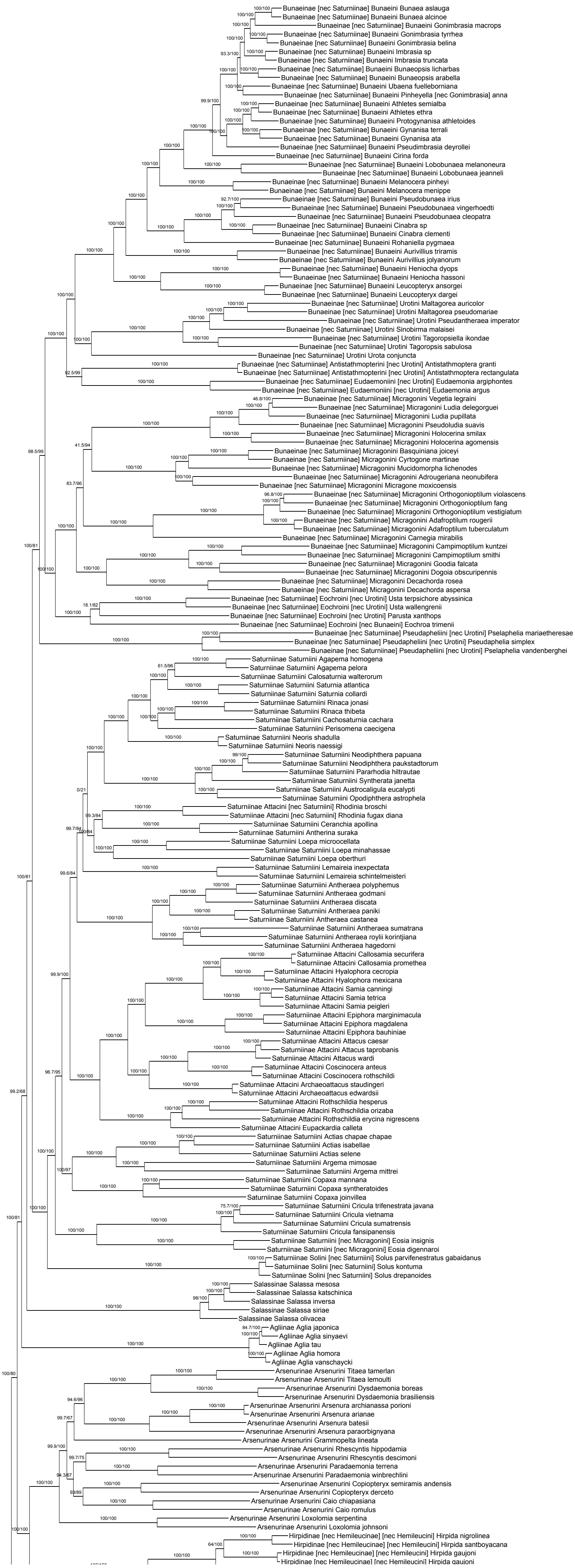

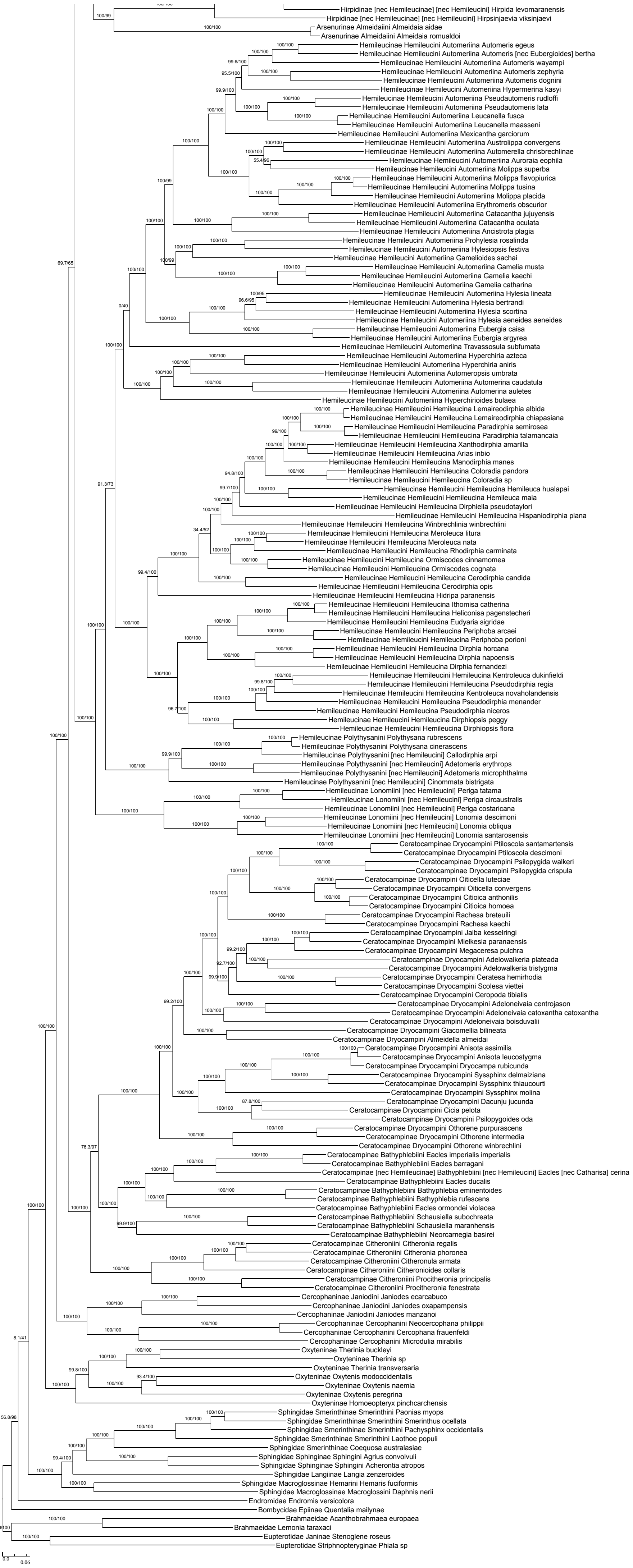
Hirpidinae [nec Hemileucinae] [nec Hemileucini] Hirpida levomaranensis
Hirpidinae [nec Hemileucinae] [nec Hemileucini] Hirpsinjaevia viksinjaevi
Arsenurinae Almeidaiini Almeidaia aidae
Arsenurinae Almeidaiini Almeidaia romualdoi
Hemileucinae Hemileucini Automeriina Automeris egeus
Hemileucinae Hemileucini Automeriina Automeris [nec Eubergioides] bertha
Hemileucinae Hemileucini Automeriina Automeris wayampi
Hemileucinae Hemileucini Automeriina Automeris zephyria
Hemileucinae Hemileucini Automeriina Automeris dognini
Hemileucinae Hemileucini Automeriina Hypermerina kasyi
Hemileucinae Hemileucini Automeriina Pseudautomeris rudloffi
Hemileucinae Hemileucini Automeriina Pseudautomeris lata
Hemileucinae Hemileucini Automeriina Leucanella fusca
Hemileucinae Hemileucini Automeriina Leucanella maasseni
Hemileucinae Hemileucini Automeriina Mexicantha garciorum
Hemileucinae Hemileucini Automeriina Austrolippa convergens
Hemileucinae Hemileucini Automeriina Automerella chrisbrechlinae
Hemileucinae Hemileucini Automeriina Auroraia eophila
Hemileucinae Hemileucini Automeriina Molippa superba
Hemileucinae Hemileucini Automeriina Molippa flavopiurica
Hemileucinae Hemileucini Automeriina Molippa tusina
Hemileucinae Hemileucini Automeriina Molippa placida
Hemileucinae Hemileucini Automeriina Erythromeris obscurior
Hemileucinae Hemileucini Automeriina Catacantha jujuyensis
Hemileucinae Hemileucini Automeriina Catacantha oculata
Hemileucinae Hemileucini Automeriina Ancistrota plagia
Hemileucinae Hemileucini Automeriina Prohylesia rosalinda
Hemileucinae Hemileucini Automeriina Hylesiopsis festiva
Hemileucinae Hemileucini Automeriina Gamelioides sachai
Hemileucinae Hemileucini Automeriina Gamelia musta
Hemileucinae Hemileucini Automeriina Gamelia kaechi
Hemileucinae Hemileucini Automeriina Gamelia catharina
Hemileucinae Hemileucini Automeriina Hylesia lineata
Hemileucinae Hemileucini Automeriina Hylesia bertrandi
Hemileucinae Hemileucini Automeriina Hylesia scortina
Hemileucinae Hemileucini Automeriina Hylesia aeneides aeneides
Hemileucinae Hemileucini Automeriina Eubergia caisa
Hemileucinae Hemileucini Automeriina Eubergia argyrea
Hemileucinae Hemileucini Automeriina Travassosula subfumata
Hemileucinae Hemileucini Automeriina Hyperchiria azteca
Hemileucinae Hemileucini Automeriina Hyperchiria aniris
Hemileucinae Hemileucini Automeriina Automeropsis umbrata
Hemileucinae Hemileucini Automeriina Automerina caudatula
Hemileucinae Hemileucini Automeriina Automerina auletes
Hemileucinae Hemileucini Automeriina Hyperchirioides bulaea
Hemileucinae Hemileucini Hemileucina Lemaireodirphia albida
Hemileucinae Hemileucini Hemileucina Lemaireodirphia chiapasiana
Hemileucinae Hemileucini Hemileucina Paradirphia semirosea
Hemileucinae Hemileucini Hemileucina Paradirphia talamancaia
Hemileucinae Hemileucini Hemileucina Xanthodirphia amarilla
Hemileucinae Hemileucini Hemileucina Arias inbio
Hemileucinae Hemileucini Hemileucina Manodirphia manes
Hemileucinae Hemileucini Hemileucina Coloradia pandora
Hemileucinae Hemileucini Hemileucina Coloradia sp
Hemileucinae Hemileucini Hemileucina Hemileuca hualapai
Hemileucinae Hemileucini Hemileucina Hemileuca maia
Hemileucinae Hemileucini Hemileucina Dirphiella pseudotaylori
Hemileucinae Hemileucini Hemileucina Hispaniodirphia plana
Hemileucinae Hemileucini Hemileucina Winbrechlinia winbrechlini
Hemileucinae Hemileucini Hemileucina Meroleuca litura
Hemileucinae Hemileucini Hemileucina Meroleuca nata
Hemileucinae Hemileucini Hemileucina Rhodirphia carminata
Hemileucinae Hemileucini Hemileucina Ormiscodes cinnamomea
Hemileucinae Hemileucini Hemileucina Ormiscodes cognata
Hemileucinae Hemileucini Hemileucina Cerodirphia candida
Hemileucinae Hemileucini Hemileucina Cerodirphia opis
Hemileucinae Hemileucini Hemileucina Hidripa paranensis
Hemileucinae Hemileucini Hemileucina Ithomisa catherina
Hemileucinae Hemileucini Hemileucina Heliconisa pagenstecheri
Hemileucinae Hemileucini Hemileucina Eudyaria sigridae
Hemileucinae Hemileucini Hemileucina Periphoba arcaei
Hemileucinae Hemileucini Hemileucina Periphoba porioni
Hemileucinae Hemileucini Hemileucina Dirphia horcana
Hemileucinae Hemileucini Hemileucina Dirphia napoensis
Hemileucinae Hemileucini Hemileucina Dirphia fernandezi
Hemileucinae Hemileucini Hemileucina Kentroleuca dukinfieldi
Hemileucinae Hemileucini Hemileucina Pseudodirphia regia
Hemileucinae Hemileucini Hemileucina Kentroleuca novaholandensis
Hemileucinae Hemileucini Hemileucina Pseudodirphia menander
Hemileucinae Hemileucini Hemileucina Pseudodirphia niceros
Hemileucinae Hemileucini Hemileucina Dirphiopsis peggy
Hemileucinae Hemileucini Hemileucina Dirphiopsis flora
Hemileucinae Polythysanini Polythysana rubrescens
Hemileucinae Polythysanini Polythysana cinerascens
Hemileucinae Polythysanini [nec Hemileucini] Callodirphia arpi
Hemileucinae Polythysanini [nec Hemileucini] Adetomeris erythrops
Hemileucinae Polythysanini [nec Hemileucini] Adetomeris microphthalma
Hemileucinae Polythysanini [nec Hemileucini] Cinommata bistrigata
Hemileucinae Lonomiini [nec Hemileucini] Periga tatama
Hemileucinae Lonomiini [nec Hemileucini] Periga circaustralis
Hemileucinae Lonomiini [nec Hemileucini] Periga costaricana
Hemileucinae Lonomiini [nec Hemileucini] Lonomia descimoni
Hemileucinae Lonomiini [nec Hemileucini] Lonomia obliqua
Hemileucinae Lonomiini [nec Hemileucini] Lonomia santarosensis
Ceratocampinae Dryocampini Ptiloscola santamartensis
Ceratocampinae Dryocampini Ptiloscola descimoni
Ceratocampinae Dryocampini Psilopygida walkeri
Ceratocampinae Dryocampini Psilopygida crispula
Ceratocampinae Dryocampini Oiticella luteciae
Ceratocampinae Dryocampini Oiticella convergens
Ceratocampinae Dryocampini Citioica anthonilis
Ceratocampinae Dryocampini Citioica homoea
Ceratocampinae Dryocampini Rachesa breteuili
Ceratocampinae Dryocampini Rachesa kaechi
Ceratocampinae Dryocampini Jaiba kesselringi
Ceratocampinae Dryocampini Mielkesia paranaensis
Ceratocampinae Dryocampini Megaceresa pulchra
Ceratocampinae Dryocampini Adelowalkeria plateada
Ceratocampinae Dryocampini Adelowalkeria tristygma
Ceratocampinae Dryocampini Ceratesa hemirhodia
Ceratocampinae Dryocampini Scolesa viettei
Ceratocampinae Dryocampini Ceropoda tibialis
Ceratocampinae Dryocampini Adeloneivaia centrojason
Ceratocampinae Dryocampini Adeloneivaia catoxantha catoxantha
Ceratocampinae Dryocampini Adeloneivaia boisduvalii
Ceratocampinae Dryocampini Giacomellia bilineata
Ceratocampinae Dryocampini Almeidella almeidai
Ceratocampinae Dryocampini Anisota assimilis
Ceratocampinae Dryocampini Anisota leucostygma
Ceratocampinae Dryocampini Dryocampa rubicunda
Ceratocampinae Dryocampini Syssphinx delmaiziana
Ceratocampinae Dryocampini Syssphinx thiaucourti
Ceratocampinae Dryocampini Syssphinx molina
Ceratocampinae Dryocampini Dacunju jucunda
Ceratocampinae Dryocampini Cicia pelota
Ceratocampinae Dryocampini Psilopygoides oda
Ceratocampinae Dryocampini Othorene purpurascens
Ceratocampinae Dryocampini Othorene intermedia
Ceratocampinae Dryocampini Othorene winbrechlini
Ceratocampinae Bathyphlebiini Eacles imperialis imperialis
Ceratocampinae Bathyphlebiini Eacles barragani
Ceratocampinae [nec Hemileucinae] Bathyphlebiini [nec Hemileucini] Eacles [nec Catharisa] cerina
Ceratocampinae Bathyphlebiini Eacles ducalis
Ceratocampinae Bathyphlebiini Bathyphlebia eminentoides
Ceratocampinae Bathyphlebiini Bathyphlebia rufescens
Ceratocampinae Bathyphlebiini Eacles ormondei violacea
Ceratocampinae Bathyphlebiini Schausiella subochreata
Ceratocampinae Bathyphlebiini Schausiella maranhensis
Ceratocampinae Bathyphlebiini Neorcarnegia basirei
Ceratocampinae Citheroniini Citheronia regalis
Ceratocampinae Citheroniini Citheronia phoronea
Ceratocampinae Citheroniini Citheronula armata
Ceratocampinae Citheroniini Citheronioides collaris
Ceratocampinae Citheroniini Procitheronia principalis
Ceratocampinae Citheroniini Procitheronia fenestrata
Cercophaninae Janiodini Janiodes ecarcabuco
Cercophaninae Janiodini Janiodes oxapampensis
Cercophaninae Janiodini Janiodes manzanoi
Cercophaninae Cercophanini Neocercophana philippii
Cercophaninae Cercophanini Cercophana frauenfeldi
Cercophaninae Cercophanini Microdulia mirabilis
Oxyteninae Therinia buckleyi
Oxyteninae Therinia sp
Oxyteninae Therinia transversaria
Oxyteninae Oxytenis modoccidentalis
Oxyteninae Oxytenis naemia
Oxyteninae Oxytenis peregrina
Oxyteninae Homoeopteryx pinchcarchensis
Sphingidae Smerinthinae Smerinthini Paonias myops
Sphingidae Smerinthinae Smerinthini Smerinthus ocellata
Sphingidae Smerinthinae Smerinthini Pachysphinx occidentalis
Sphingidae Smerinthinae Smerinthini Laothoe populi
Sphingidae Smerinthinae Coequosa australasiae
Sphingidae Sphinginae Sphingini Agrius convolvuli
Sphingidae Sphinginae Sphingini Acherontia atropos
Sphingidae Langiinae Langia zenzeroides
Sphingidae Macroglossinae Hemarini Hemaris fuciformis
Sphingidae Macroglossinae Macroglossini Daphnis nerii
Endromidae Endromis versicolora
Bombycidae Epiinae Quentalia mailynae
Brahmaeidae Acanthobrahmaea europaea
Brahmaeidae Lemonia taraxaci
Eupterotidae Janinae Stenoglene roseus
Eupterotidae Striphnopteryginae Phiala sp
0.0
0.06

#### Figure S2F. Data subset (20% most saturated loci) ASTRAL50

Species tree obtained when nodes with UFBoot<50 are collapsed in each gene tree

Final normalized quartet score : 0.79
Local posterior probabilities at nodes

The new classification proposed in this manuscript is used to annotate tips. Current classification is reported between square brackets

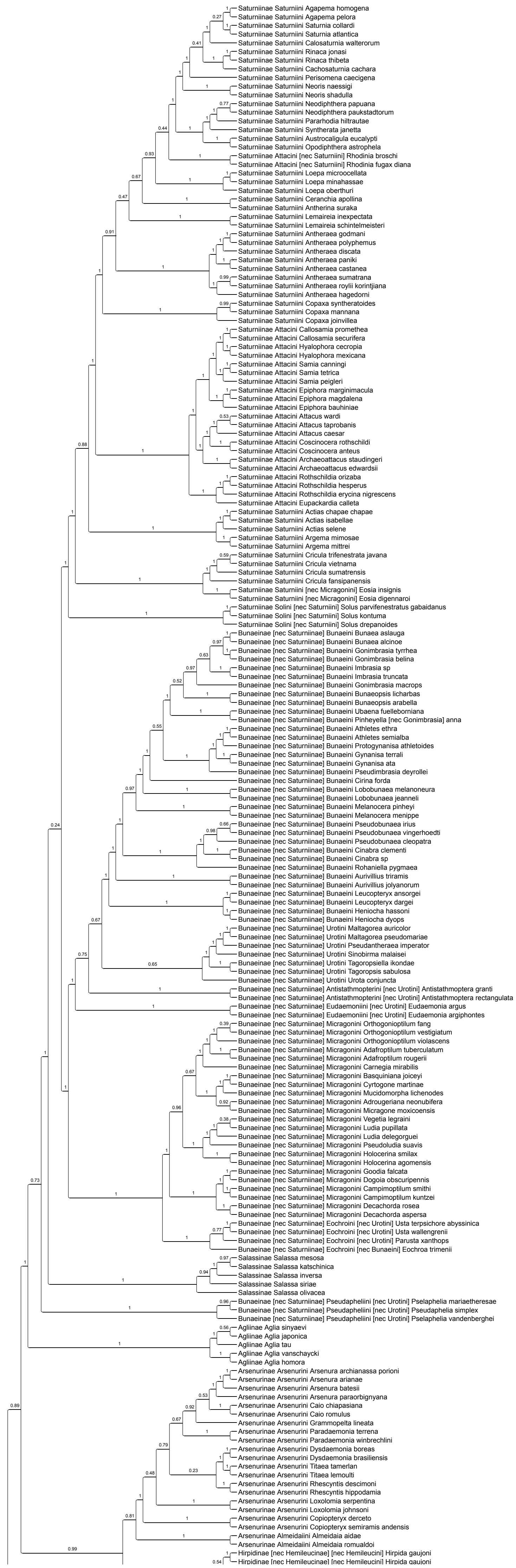

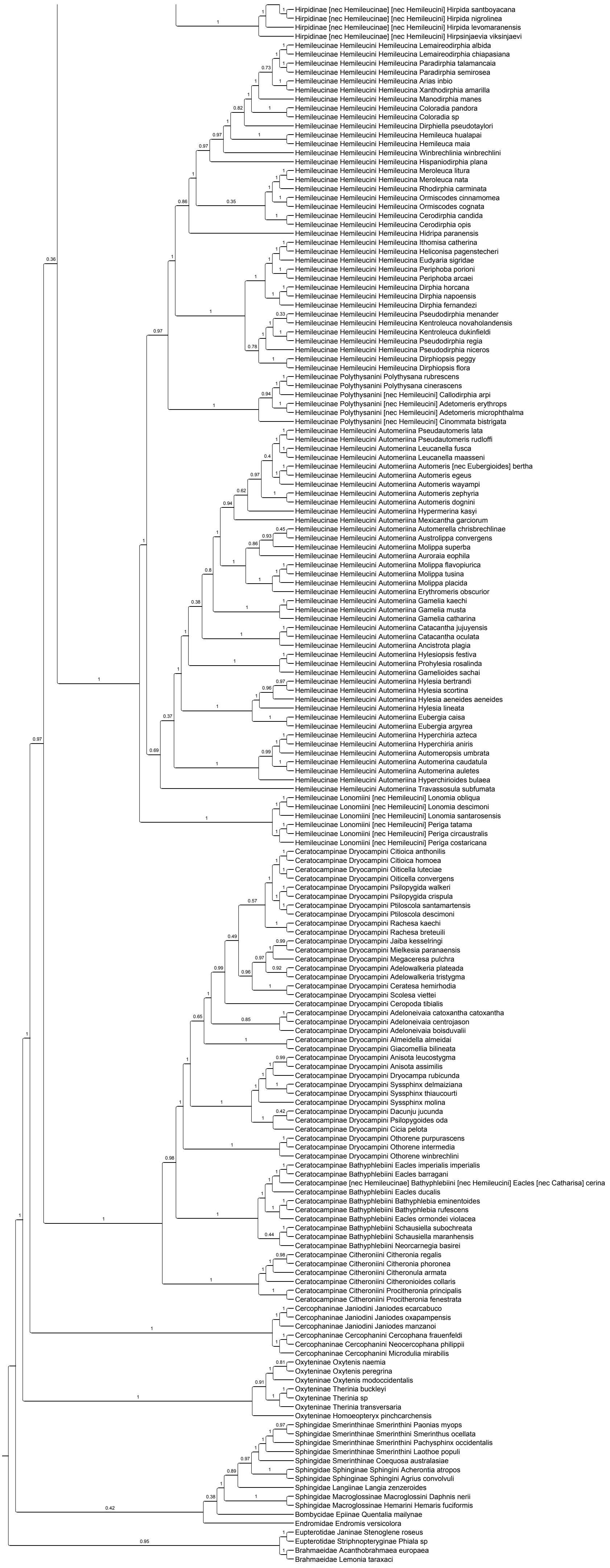
Hirpidinae [nec Hemileucinae] [nec Hemileucini] Hirpida santboyacana
Hirpidinae [nec Hemileucinae] [nec Hemileucini] Hirpida nigrolinea
Hirpidinae [nec Hemileucinae] [nec Hemileucini] Hirpida levomaranensis
Hirpidinae [nec Hemileucinae] [nec Hemileucini] Hirpsinjaevia viksinjaevi
Hemileucinae Hemileucini Hemileucina Lemaireodirphia albida
Hemileucinae Hemileucini Hemileucina Lemaireodirphia chiapasiana
Hemileucinae Hemileucini Hemileucina Paradirphia talamancaia
Hemileucinae Hemileucini Hemileucina Paradirphia semirosea
Hemileucinae Hemileucini Hemileucina Arias inbio
Hemileucinae Hemileucini Hemileucina Xanthodirphia amarilla
Hemileucinae Hemileucini Hemileucina Manodirphia manes
Hemileucinae Hemileucini Hemileucina Coloradia pandora
Hemileucinae Hemileucini Hemileucina Coloradia sp
Hemileucinae Hemileucini Hemileucina Dirphiella pseudotaylori
Hemileucinae Hemileucini Hemileucina Hemileuca hualapai
Hemileucinae Hemileucini Hemileucina Hemileuca maia
Hemileucinae Hemileucini Hemileucina Winbrechlinia winbrechlini
Hemileucinae Hemileucini Hemileucina Hispaniodirphia plana
Hemileucinae Hemileucini Hemileucina Meroleuca litura
Hemileucinae Hemileucini Hemileucina Meroleuca nata
Hemileucinae Hemileucini Hemileucina Rhodirphia carminata
Hemileucinae Hemileucini Hemileucina Ormiscodes cinnamomea
Hemileucinae Hemileucini Hemileucina Ormiscodes cognata
Hemileucinae Hemileucini Hemileucina Cerodirphia candida
Hemileucinae Hemileucini Hemileucina Cerodirphia opis
Hemileucinae Hemileucini Hemileucina Hidripa paranensis
Hemileucinae Hemileucini Hemileucina Ithomisa catherina
Hemileucinae Hemileucini Hemileucina Heliconisa pagenstecheri
Hemileucinae Hemileucini Hemileucina Eudyaria sigridae
Hemileucinae Hemileucini Hemileucina Periphoba porioni
Hemileucinae Hemileucini Hemileucina Periphoba arcaei
Hemileucinae Hemileucini Hemileucina Dirphia horcana
Hemileucinae Hemileucini Hemileucina Dirphia napoensis
Hemileucinae Hemileucini Hemileucina Dirphia fernandezi
Hemileucinae Hemileucini Hemileucina Pseudodirphia menander
Hemileucinae Hemileucini Hemileucina Kentroleuca novaholandensis
Hemileucinae Hemileucini Hemileucina Kentroleuca dukinfieldi
Hemileucinae Hemileucini Hemileucina Pseudodirphia regia
Hemileucinae Hemileucini Hemileucina Pseudodirphia niceros
Hemileucinae Hemileucini Hemileucina Dirphiopsis peggy
Hemileucinae Hemileucini Hemileucina Dirphiopsis flora
Hemileucinae Polythysanini Polythysana rubrescens
Hemileucinae Polythysanini Polythysana cinerascens
Hemileucinae Polythysanini [nec Hemileucini] Callodirphia arpi
Hemileucinae Polythysanini [nec Hemileucini] Adetomeris erythrops
Hemileucinae Polythysanini [nec Hemileucini] Adetomeris microphthalma
Hemileucinae Polythysanini [nec Hemileucini] Cinommata bistrigata
Hemileucinae Hemileucini Automeriina Pseudautomeris lata
Hemileucinae Hemileucini Automeriina Pseudautomeris rudloffi
Hemileucinae Hemileucini Automeriina Leucanella fusca
Hemileucinae Hemileucini Automeriina Leucanella maasseni
Hemileucinae Hemileucini Automeriina Automeris [nec Eubergioides] bertha
Hemileucinae Hemileucini Automeriina Automeris egeus
Hemileucinae Hemileucini Automeriina Automeris wayampi
Hemileucinae Hemileucini Automeriina Automeris zephyria
Hemileucinae Hemileucini Automeriina Automeris dognini
Hemileucinae Hemileucini Automeriina Hypermerina kasyi
Hemileucinae Hemileucini Automeriina Mexicantha garciorum
Hemileucinae Hemileucini Automeriina Automerella chrisbrechlinae
Hemileucinae Hemileucini Automeriina Austrolippa convergens
Hemileucinae Hemileucini Automeriina Molippa superba
Hemileucinae Hemileucini Automeriina Auroraia eophila
Hemileucinae Hemileucini Automeriina Molippa flavopiurica
Hemileucinae Hemileucini Automeriina Molippa tusina
Hemileucinae Hemileucini Automeriina Molippa placida
Hemileucinae Hemileucini Automeriina Erythromeris obscurior
Hemileucinae Hemileucini Automeriina Gamelia kaechi
Hemileucinae Hemileucini Automeriina Gamelia musta
Hemileucinae Hemileucini Automeriina Gamelia catharina
Hemileucinae Hemileucini Automeriina Catacantha jujuyensis
Hemileucinae Hemileucini Automeriina Catacantha oculata
Hemileucinae Hemileucini Automeriina Ancistrota plagia
Hemileucinae Hemileucini Automeriina Hylesiopsis festiva
Hemileucinae Hemileucini Automeriina Prohylesia rosalinda
Hemileucinae Hemileucini Automeriina Gamelioides sachai
Hemileucinae Hemileucini Automeriina Hylesia bertrandi
Hemileucinae Hemileucini Automeriina Hylesia scortina
Hemileucinae Hemileucini Automeriina Hylesia aeneides aeneides
Hemileucinae Hemileucini Automeriina Hylesia lineata
Hemileucinae Hemileucini Automeriina Eubergia caisa
Hemileucinae Hemileucini Automeriina Eubergia argyrea
Hemileucinae Hemileucini Automeriina Hyperchiria azteca
Hemileucinae Hemileucini Automeriina Hyperchiria aniris
Hemileucinae Hemileucini Automeriina Automeropsis umbrata
Hemileucinae Hemileucini Automeriina Automerina caudatula
Hemileucinae Hemileucini Automeriina Automerina auletes
Hemileucinae Hemileucini Automeriina Hyperchirioides bulaea
Hemileucinae Hemileucini Automeriina Travassosula subfumata
Hemileucinae Lonomiini [nec Hemileucini] Lonomia obliqua
Hemileucinae Lonomiini [nec Hemileucini] Lonomia descimoni
Hemileucinae Lonomiini [nec Hemileucini] Lonomia santarosensis
Hemileucinae Lonomiini [nec Hemileucini] Periga tatama
Hemileucinae Lonomiini [nec Hemileucini] Periga circaustralis
Hemileucinae Lonomiini [nec Hemileucini] Periga costaricana
Ceratocampinae Dryocampini Citioica anthonilis
Ceratocampinae Dryocampini Citioica homoea
Ceratocampinae Dryocampini Oiticella luteciae
Ceratocampinae Dryocampini Oiticella convergens
Ceratocampinae Dryocampini Psilopygida walkeri
Ceratocampinae Dryocampini Psilopygida crispula
Ceratocampinae Dryocampini Ptiloscola santamartensis
Ceratocampinae Dryocampini Ptiloscola descimoni
Ceratocampinae Dryocampini Rachesa kaechi
Ceratocampinae Dryocampini Rachesa breteuili
Ceratocampinae Dryocampini Jaiba kesselringi
Ceratocampinae Dryocampini Mielkesia paranaensis
Ceratocampinae Dryocampini Megaceresa pulchra
Ceratocampinae Dryocampini Adelowalkeria plateada
Ceratocampinae Dryocampini Adelowalkeria tristygma
Ceratocampinae Dryocampini Ceratesa hemirhodia
Ceratocampinae Dryocampini Scolesa viettei
Ceratocampinae Dryocampini Ceropoda tibialis
Ceratocampinae Dryocampini Adeloneivaia catoxantha catoxantha
Ceratocampinae Dryocampini Adeloneivaia centrojason
Ceratocampinae Dryocampini Adeloneivaia boisduvalii
Ceratocampinae Dryocampini Almeidella almeidai
Ceratocampinae Dryocampini Giacomellia bilineata
Ceratocampinae Dryocampini Anisota leucostygma
Ceratocampinae Dryocampini Anisota assimilis
Ceratocampinae Dryocampini Dryocampa rubicunda
Ceratocampinae Dryocampini Syssphinx delmaiziana
Ceratocampinae Dryocampini Syssphinx thiaucourti
Ceratocampinae Dryocampini Syssphinx molina
Ceratocampinae Dryocampini Dacunju jucunda
Ceratocampinae Dryocampini Psilopygoides oda
Ceratocampinae Dryocampini Cicia pelota
Ceratocampinae Dryocampini Othorene purpurascens
Ceratocampinae Dryocampini Othorene intermedia
Ceratocampinae Dryocampini Othorene winbrechlini
Ceratocampinae Bathyphlebiini Eacles imperialis imperialis
Ceratocampinae Bathyphlebiini Eacles barragani
Ceratocampinae [nec Hemileucinae] Bathyphlebiini [nec Hemileucini] Eacles [nec Catharisa] cerina
Ceratocampinae Bathyphlebiini Eacles ducalis
Ceratocampinae Bathyphlebiini Bathyphlebia eminentoides
Ceratocampinae Bathyphlebiini Bathyphlebia rufescens
Ceratocampinae Bathyphlebiini Eacles ormondei violacea
Ceratocampinae Bathyphlebiini Schausiella subochreata
Ceratocampinae Bathyphlebiini Schausiella maranhensis
Ceratocampinae Bathyphlebiini Neorcarnegia basirei
Ceratocampinae Citheroniini Citheronia regalis
Ceratocampinae Citheroniini Citheronia phoronea
Ceratocampinae Citheroniini Citheronula armata
Ceratocampinae Citheroniini Citheronioides collaris
Ceratocampinae Citheroniini Procitheronia principalis
Ceratocampinae Citheroniini Procitheronia fenestrata
Cercophaninae Janiodini Janiodes ecarcabuco
Cercophaninae Janiodini Janiodes oxapampensis
Cercophaninae Janiodini Janiodes manzanoi
Cercophaninae Cercophanini Cercophana frauenfeldi
Cercophaninae Cercophanini Neocercophana philippii
Cercophaninae Cercophanini Microdulia mirabilis
Oxyteninae Oxytenis naemia
Oxyteninae Oxytenis peregrina
Oxyteninae Oxytenis modoccidentalis
Oxyteninae Therinia buckleyi
Oxyteninae Therinia sp
Oxyteninae Therinia transversaria
Oxyteninae Homoeopteryx pinchcarchensis
Sphingidae Smerinthinae Smerinthini Paonias myops
Sphingidae Smerinthinae Smerinthini Smerinthus ocellata
Sphingidae Smerinthinae Smerinthini Pachysphinx occidentalis
Sphingidae Smerinthinae Smerinthini Laothoe populi
Sphingidae Smerinthinae Coequosa australasiae
Sphingidae Sphinginae Sphingini Acherontia atropos
Sphingidae Sphinginae Sphingini Agrius convolvuli
Sphingidae Langiinae Langia zenzeroides
Sphingidae Macroglossinae Macroglossini Daphnis nerii
Sphingidae Macroglossinae Hemarini Hemaris fuciformis
Bombycidae Epiinae Quentalia mailynae
Endromidae Endromis versicolora
Eupterotidae Janinae Stenoglene roseus
Eupterotidae Striphnopteryginae Phiala sp
Brahmaeidae Acanthobrahmaea europaea
Brahmaeidae Lemonia taraxaci

### Figure S2G. Data subset (20% most saturated loci) ASTRAL70

Species tree obtained when nodes with UFBoot<70 are collapsed in each gene tree

Final normalized quartet score : 0.83
Local posterior probabilities at nodes

The new classification proposed in this manuscript is used to annotate tips. Current classification is reported between square brackets

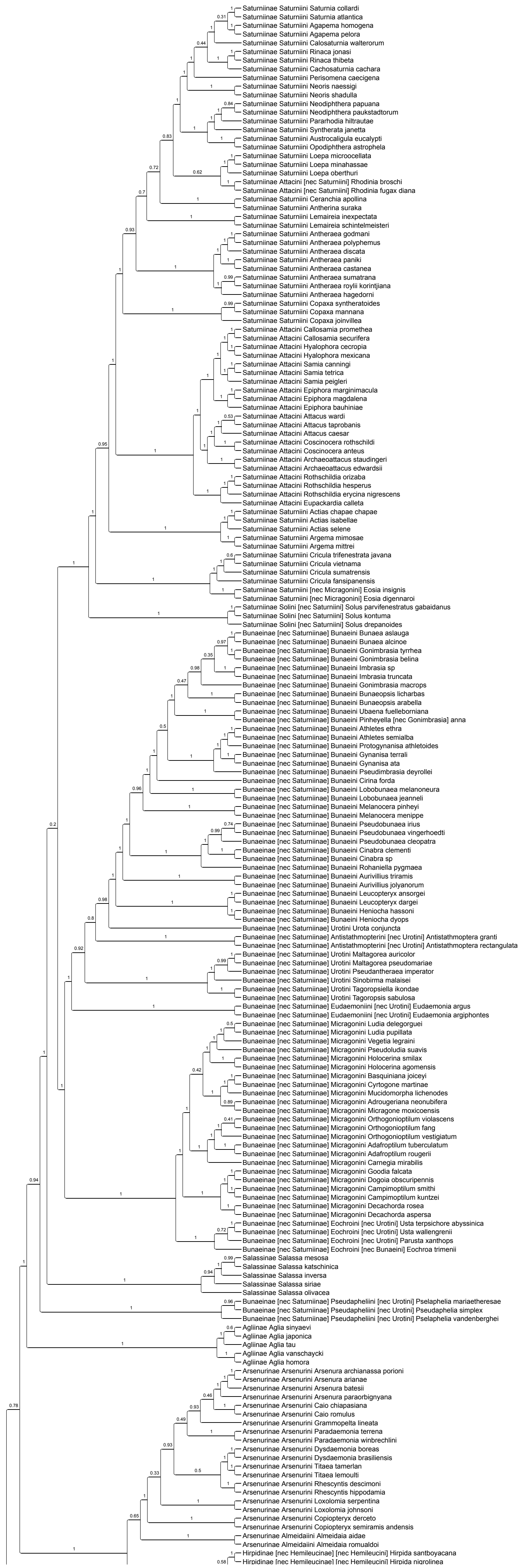

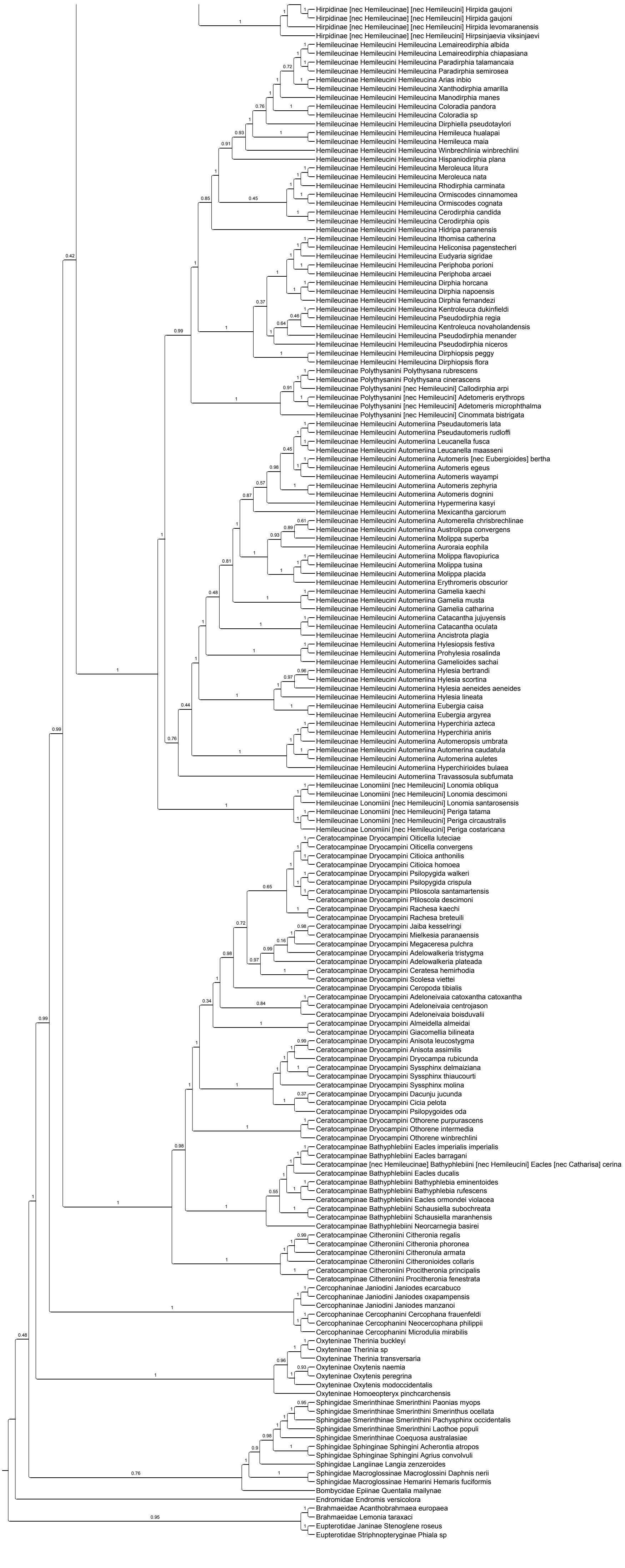
Hirpidinae [nec Hemileucinae] [nec Hemileucini] Hirpida gaujoni
Hirpidinae [nec Hemileucinae] [nec Hemileucini] Hirpida gaujoni
Hirpidinae [nec Hemileucinae] [nec Hemileucini] Hirpida levomaranensis
Hirpidinae [nec Hemileucinae] [nec Hemileucini] Hirpsinjaevia viksinjaevi
Hemileucinae Hemileucini Hemileucina Lemaireodirphia albida
Hemileucinae Hemileucini Hemileucina Lemaireodirphia chiapasiana
Hemileucinae Hemileucini Hemileucina Paradirphia talamancaia
Hemileucinae Hemileucini Hemileucina Paradirphia semirosea
Hemileucinae Hemileucini Hemileucina Arias inbio
Hemileucinae Hemileucini Hemileucina Xanthodirphia amarilla
Hemileucinae Hemileucini Hemileucina Manodirphia manes
Hemileucinae Hemileucini Hemileucina Coloradia pandora
Hemileucinae Hemileucini Hemileucina Coloradia sp
Hemileucinae Hemileucini Hemileucina Dirphiella pseudotaylori
Hemileucinae Hemileucini Hemileucina Hemileuca hualapai
Hemileucinae Hemileucini Hemileucina Hemileuca maia
Hemileucinae Hemileucini Hemileucina Winbrechlinia winbrechlini
Hemileucinae Hemileucini Hemileucina Hispaniodirphia plana
Hemileucinae Hemileucini Hemileucina Meroleuca litura
Hemileucinae Hemileucini Hemileucina Meroleuca nata
Hemileucinae Hemileucini Hemileucina Rhodirphia carminata
Hemileucinae Hemileucini Hemileucina Ormiscodes cinnamomea
Hemileucinae Hemileucini Hemileucina Ormiscodes cognata
Hemileucinae Hemileucini Hemileucina Cerodirphia candida
Hemileucinae Hemileucini Hemileucina Cerodirphia opis
Hemileucinae Hemileucini Hemileucina Hidripa paranensis
Hemileucinae Hemileucini Hemileucina Ithomisa catherina
Hemileucinae Hemileucini Hemileucina Heliconisa pagenstecheri
Hemileucinae Hemileucini Hemileucina Eudyaria sigridae
Hemileucinae Hemileucini Hemileucina Periphoba porioni
Hemileucinae Hemileucini Hemileucina Periphoba arcaei
Hemileucinae Hemileucini Hemileucina Dirphia horcana
Hemileucinae Hemileucini Hemileucina Dirphia napoensis
Hemileucinae Hemileucini Hemileucina Dirphia fernandezi
Hemileucinae Hemileucini Hemileucina Kentroleuca dukinfieldi
Hemileucinae Hemileucini Hemileucina Pseudodirphia regia
Hemileucinae Hemileucini Hemileucina Kentroleuca novaholandensis
Hemileucinae Hemileucini Hemileucina Pseudodirphia menander
Hemileucinae Hemileucini Hemileucina Pseudodirphia niceros
Hemileucinae Hemileucini Hemileucina Dirphiopsis peggy
Hemileucinae Hemileucini Hemileucina Dirphiopsis flora
Hemileucinae Polythysanini Polythysana rubrescens
Hemileucinae Polythysanini Polythysana cinerascens
Hemileucinae Polythysanini [nec Hemileucini] Callodirphia arpi
Hemileucinae Polythysanini [nec Hemileucini] Adetomeris erythrops
Hemileucinae Polythysanini [nec Hemileucini] Adetomeris microphthalma
Hemileucinae Polythysanini [nec Hemileucini] Cinommata bistrigata
Hemileucinae Hemileucini Automeriina Pseudautomeris lata
Hemileucinae Hemileucini Automeriina Pseudautomeris rudloffi
Hemileucinae Hemileucini Automeriina Leucanella fusca
Hemileucinae Hemileucini Automeriina Leucanella maasseni
Hemileucinae Hemileucini Automeriina Automeris [nec Eubergioides] bertha
Hemileucinae Hemileucini Automeriina Automeris egeus
Hemileucinae Hemileucini Automeriina Automeris wayampi
Hemileucinae Hemileucini Automeriina Automeris zephyria
Hemileucinae Hemileucini Automeriina Automeris dognini
Hemileucinae Hemileucini Automeriina Hypermerina kasyi
Hemileucinae Hemileucini Automeriina Mexicantha garciorum
Hemileucinae Hemileucini Automeriina Automerella chrisbrechlinae
Hemileucinae Hemileucini Automeriina Austrolippa convergens
Hemileucinae Hemileucini Automeriina Molippa superba
Hemileucinae Hemileucini Automeriina Auroraia eophila
Hemileucinae Hemileucini Automeriina Molippa flavopiurica
Hemileucinae Hemileucini Automeriina Molippa tusina
Hemileucinae Hemileucini Automeriina Molippa placida
Hemileucinae Hemileucini Automeriina Erythromeris obscurior
Hemileucinae Hemileucini Automeriina Gamelia kaechi
Hemileucinae Hemileucini Automeriina Gamelia musta
Hemileucinae Hemileucini Automeriina Gamelia catharina
Hemileucinae Hemileucini Automeriina Catacantha jujuyensis
Hemileucinae Hemileucini Automeriina Catacantha oculata
Hemileucinae Hemileucini Automeriina Ancistrota plagia
Hemileucinae Hemileucini Automeriina Hylesiopsis festiva
Hemileucinae Hemileucini Automeriina Prohylesia rosalinda
Hemileucinae Hemileucini Automeriina Gamelioides sachai
Hemileucinae Hemileucini Automeriina Hylesia bertrandi
Hemileucinae Hemileucini Automeriina Hylesia scortina
Hemileucinae Hemileucini Automeriina Hylesia aeneides aeneides
Hemileucinae Hemileucini Automeriina Hylesia lineata
Hemileucinae Hemileucini Automeriina Eubergia caisa
Hemileucinae Hemileucini Automeriina Eubergia argyrea
Hemileucinae Hemileucini Automeriina Hyperchiria azteca
Hemileucinae Hemileucini Automeriina Hyperchiria aniris
Hemileucinae Hemileucini Automeriina Automeropsis umbrata
Hemileucinae Hemileucini Automeriina Automerina caudatula
Hemileucinae Hemileucini Automeriina Automerina auletes
Hemileucinae Hemileucini Automeriina Hyperchirioides bulaea
Hemileucinae Hemileucini Automeriina Travassosula subfumata
Hemileucinae Lonomiini [nec Hemileucini] Lonomia obliqua
Hemileucinae Lonomiini [nec Hemileucini] Lonomia descimoni
Hemileucinae Lonomiini [nec Hemileucini] Lonomia santarosensis
Hemileucinae Lonomiini [nec Hemileucini] Periga tatama
Hemileucinae Lonomiini [nec Hemileucini] Periga circaustralis
Hemileucinae Lonomiini [nec Hemileucini] Periga costaricana
Ceratocampinae Dryocampini Oiticella luteciae
Ceratocampinae Dryocampini Oiticella convergens
Ceratocampinae Dryocampini Citioica anthonilis
Ceratocampinae Dryocampini Citioica homoea
Ceratocampinae Dryocampini Psilopygida walkeri
Ceratocampinae Dryocampini Psilopygida crispula
Ceratocampinae Dryocampini Ptiloscola santamartensis
Ceratocampinae Dryocampini Ptiloscola descimoni
Ceratocampinae Dryocampini Rachesa kaechi
Ceratocampinae Dryocampini Rachesa breteuili
Ceratocampinae Dryocampini Jaiba kesselringi
Ceratocampinae Dryocampini Mielkesia paranaensis
Ceratocampinae Dryocampini Megaceresa pulchra
Ceratocampinae Dryocampini Adelowalkeria tristygma
Ceratocampinae Dryocampini Adelowalkeria plateada
Ceratocampinae Dryocampini Ceratesa hemirhodia
Ceratocampinae Dryocampini Scolesa viettei
Ceratocampinae Dryocampini Ceropoda tibialis
Ceratocampinae Dryocampini Adeloneivaia catoxantha catoxantha
Ceratocampinae Dryocampini Adeloneivaia centrojason
Ceratocampinae Dryocampini Adeloneivaia boisduvalii
Ceratocampinae Dryocampini Almeidella almeidai
Ceratocampinae Dryocampini Giacomellia bilineata
Ceratocampinae Dryocampini Anisota leucostygma
Ceratocampinae Dryocampini Anisota assimilis
Ceratocampinae Dryocampini Dryocampa rubicunda
Ceratocampinae Dryocampini Syssphinx delmaiziana
Ceratocampinae Dryocampini Syssphinx thiaucourti
Ceratocampinae Dryocampini Syssphinx molina
Ceratocampinae Dryocampini Dacunju jucunda
Ceratocampinae Dryocampini Cicia pelota
Ceratocampinae Dryocampini Psilopygoides oda
Ceratocampinae Dryocampini Othorene purpurascens
Ceratocampinae Dryocampini Othorene intermedia
Ceratocampinae Dryocampini Othorene winbrechlini
Ceratocampinae Bathyphlebiini Eacles imperialis imperialis
Ceratocampinae Bathyphlebiini Eacles barragani
Ceratocampinae [nec Hemileucinae] Bathyphlebiini [nec Hemileucini] Eacles [nec Catharisa] cerina
Ceratocampinae Bathyphlebiini Eacles ducalis
Ceratocampinae Bathyphlebiini Bathyphlebia eminentoides
Ceratocampinae Bathyphlebiini Bathyphlebia rufescens
Ceratocampinae Bathyphlebiini Eacles ormondei violacea
Ceratocampinae Bathyphlebiini Schausiella subochreata
Ceratocampinae Bathyphlebiini Schausiella maranhensis
Ceratocampinae Bathyphlebiini Neorcarnegia basirei
Ceratocampinae Citheroniini Citheronia regalis
Ceratocampinae Citheroniini Citheronia phoronea
Ceratocampinae Citheroniini Citheronula armata
Ceratocampinae Citheroniini Citheronioides collaris
Ceratocampinae Citheroniini Procitheronia principalis
Ceratocampinae Citheroniini Procitheronia fenestrata
Cercophaninae Janiodini Janiodes ecarcabuco
Cercophaninae Janiodini Janiodes oxapampensis
Cercophaninae Janiodini Janiodes manzanoi
Cercophaninae Cercophanini Cercophana frauenfeldi
Cercophaninae Cercophanini Neocercophana philippii
Cercophaninae Cercophanini Microdulia mirabilis
Oxyteninae Therinia buckleyi
Oxyteninae Therinia sp
Oxyteninae Therinia transversaria
Oxyteninae Oxytenis naemia
Oxyteninae Oxytenis peregrina
Oxyteninae Oxytenis modoccidentalis
Oxyteninae Homoeopteryx pinchcarchensis
Sphingidae Smerinthinae Smerinthini Paonias myops
Sphingidae Smerinthinae Smerinthini Smerinthus ocellata
Sphingidae Smerinthinae Smerinthini Pachysphinx occidentalis
Sphingidae Smerinthinae Smerinthini Laothoe populi
Sphingidae Smerinthinae Coequosa australasiae
Sphingidae Sphinginae Sphingini Acherontia atropos
Sphingidae Sphinginae Sphingini Agrius convolvuli
Sphingidae Langiinae Langia zenzeroides
Sphingidae Macroglossinae Macroglossini Daphnis nerii
Sphingidae Macroglossinae Hemarini Hemaris fuciformis
Bombycidae Epiinae Quentalia mailynae
Endromidae Endromis versicolora
Brahmaeidae Acanthobrahmaea europaea
Brahmaeidae Lemonia taraxaci
Eupterotidae Janinae Stenoglene roseus
Eupterotidae Striphnopteryginae Phiala sp

### Figure S2H. Data subset (20% most saturated loci) ASTRAL90

Species tree obtained when nodes with UFBoot<90 are collapsed in each gene tree

Final normalized quartet score : 0.92
Local posterior probabilities at nodes

The new classification proposed in this manuscript is used to annotate tips. Current classification is reported between square brackets

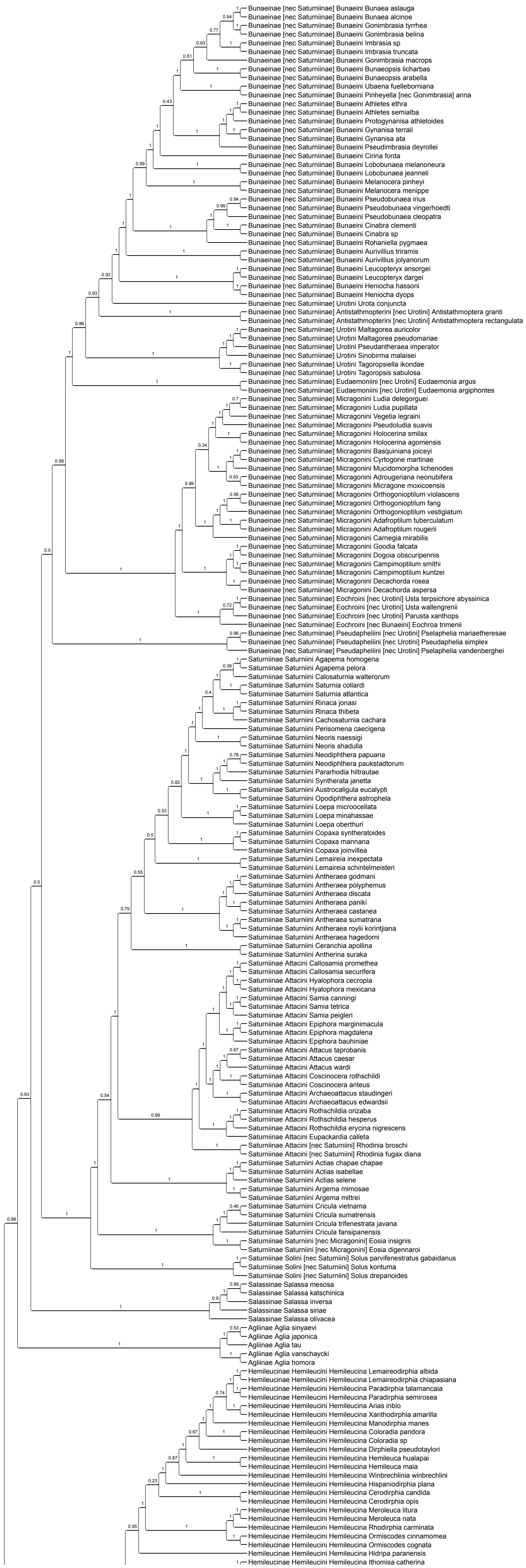

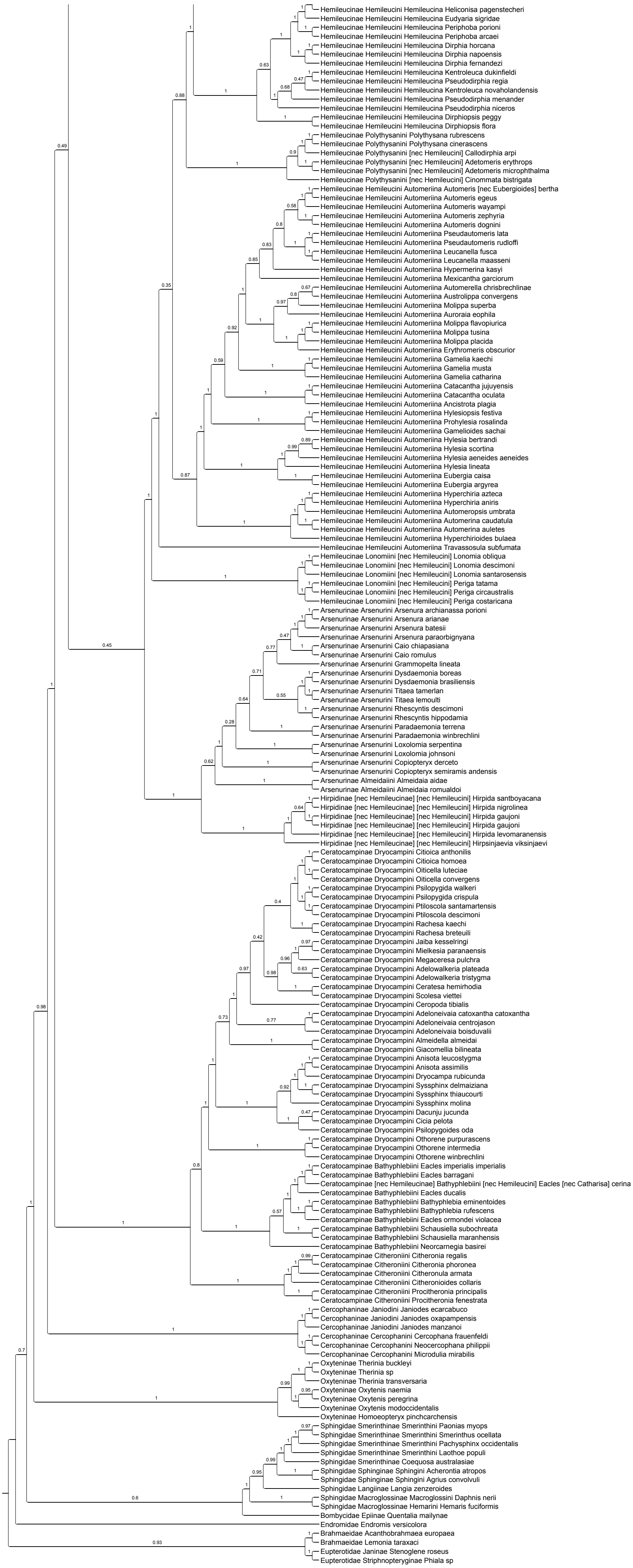
Hemileucinae Hemileucini Hemileucina Heliconisa pagenstecheri
Hemileucinae Hemileucini Hemileucina Eudyaria sigridae
Hemileucinae Hemileucini Hemileucina Periphoba porioni
Hemileucinae Hemileucini Hemileucina Periphoba arcaei
Hemileucinae Hemileucini Hemileucina Dirphia horcana
Hemileucinae Hemileucini Hemileucina Dirphia napoensis
Hemileucinae Hemileucini Hemileucina Dirphia fernandezi
Hemileucinae Hemileucini Hemileucina Kentroleuca dukinfieldi
Hemileucinae Hemileucini Hemileucina Pseudodirphia regia
Hemileucinae Hemileucini Hemileucina Kentroleuca novaholandensis
Hemileucinae Hemileucini Hemileucina Pseudodirphia menander
Hemileucinae Hemileucini Hemileucina Pseudodirphia niceros
Hemileucinae Hemileucini Hemileucina Dirphiopsis peggy
Hemileucinae Hemileucini Hemileucina Dirphiopsis flora
Hemileucinae Polythysanini Polythysana rubrescens
Hemileucinae Polythysanini Polythysana cinerascens
Hemileucinae Polythysanini [nec Hemileucini] Callodirphia arpi
Hemileucinae Polythysanini [nec Hemileucini] Adetomeris erythrops
Hemileucinae Polythysanini [nec Hemileucini] Adetomeris microphthalma
Hemileucinae Polythysanini [nec Hemileucini] Cinommata bistrigata
Hemileucinae Hemileucini Automeriina Automeris [nec Eubergioides] bertha
Hemileucinae Hemileucini Automeriina Automeris egeus
Hemileucinae Hemileucini Automeriina Automeris wayampi
Hemileucinae Hemileucini Automeriina Automeris zephyria
Hemileucinae Hemileucini Automeriina Automeris dognini
Hemileucinae Hemileucini Automeriina Pseudautomeris lata
Hemileucinae Hemileucini Automeriina Pseudautomeris rudloffi
Hemileucinae Hemileucini Automeriina Leucanella fusca
Hemileucinae Hemileucini Automeriina Leucanella maasseni
Hemileucinae Hemileucini Automeriina Hypermerina kasyi
Hemileucinae Hemileucini Automeriina Mexicantha garciorum
Hemileucinae Hemileucini Automeriina Automerella chrisbrechlinae
Hemileucinae Hemileucini Automeriina Austrolippa convergens
Hemileucinae Hemileucini Automeriina Molippa superba
Hemileucinae Hemileucini Automeriina Auroraia eophila
Hemileucinae Hemileucini Automeriina Molippa flavopiurica
Hemileucinae Hemileucini Automeriina Molippa tusina
Hemileucinae Hemileucini Automeriina Molippa placida
Hemileucinae Hemileucini Automeriina Erythromeris obscurior
Hemileucinae Hemileucini Automeriina Gamelia kaechi
Hemileucinae Hemileucini Automeriina Gamelia musta
Hemileucinae Hemileucini Automeriina Gamelia catharina
Hemileucinae Hemileucini Automeriina Catacantha jujuyensis
Hemileucinae Hemileucini Automeriina Catacantha oculata
Hemileucinae Hemileucini Automeriina Ancistrota plagia
Hemileucinae Hemileucini Automeriina Hylesiopsis festiva
Hemileucinae Hemileucini Automeriina Prohylesia rosalinda
Hemileucinae Hemileucini Automeriina Gamelioides sachai
Hemileucinae Hemileucini Automeriina Hylesia bertrandi
Hemileucinae Hemileucini Automeriina Hylesia scortina
Hemileucinae Hemileucini Automeriina Hylesia aeneides aeneides
Hemileucinae Hemileucini Automeriina Hylesia lineata
Hemileucinae Hemileucini Automeriina Eubergia caisa
Hemileucinae Hemileucini Automeriina Eubergia argyrea
Hemileucinae Hemileucini Automeriina Hyperchiria azteca
Hemileucinae Hemileucini Automeriina Hyperchiria aniris
Hemileucinae Hemileucini Automeriina Automeropsis umbrata
Hemileucinae Hemileucini Automeriina Automerina caudatula
Hemileucinae Hemileucini Automeriina Automerina auletes
Hemileucinae Hemileucini Automeriina Hyperchirioides bulaea
Hemileucinae Hemileucini Automeriina Travassosula subfumata
Hemileucinae Lonomiini [nec Hemileucini] Lonomia obliqua
Hemileucinae Lonomiini [nec Hemileucini] Lonomia descimoni
Hemileucinae Lonomiini [nec Hemileucini] Lonomia santarosensis
Hemileucinae Lonomiini [nec Hemileucini] Periga tatama
Hemileucinae Lonomiini [nec Hemileucini] Periga circaustralis
Hemileucinae Lonomiini [nec Hemileucini] Periga costaricana
Arsenurinae Arsenurini Arsenura archianassa porioni
Arsenurinae Arsenurini Arsenura arianae
Arsenurinae Arsenurini Arsenura batesii
Arsenurinae Arsenurini Arsenura paraorbignyana
Arsenurinae Arsenurini Caio chiapasiana
Arsenurinae Arsenurini Caio romulus
Arsenurinae Arsenurini Grammopelta lineata
Arsenurinae Arsenurini Dysdaemonia boreas
Arsenurinae Arsenurini Dysdaemonia brasiliensis
Arsenurinae Arsenurini Titaea tamerlan
Arsenurinae Arsenurini Titaea lemoulti
Arsenurinae Arsenurini Rhescyntis descimoni
Arsenurinae Arsenurini Rhescyntis hippodamia
Arsenurinae Arsenurini Paradaemonia terrena
Arsenurinae Arsenurini Paradaemonia winbrechlini
Arsenurinae Arsenurini Loxolomia serpentina
Arsenurinae Arsenurini Loxolomia johnsoni
Arsenurinae Arsenurini Copiopteryx derceto
Arsenurinae Arsenurini Copiopteryx semiramis andensis
Arsenurinae Almeidaiini Almeidaia aidae
Arsenurinae Almeidaiini Almeidaia romualdoi
Hirpidinae [nec Hemileucinae] [nec Hemileucini] Hirpida santboyacana
Hirpidinae [nec Hemileucinae] [nec Hemileucini] Hirpida nigrolinea
Hirpidinae [nec Hemileucinae] [nec Hemileucini] Hirpida gaujoni
Hirpidinae [nec Hemileucinae] [nec Hemileucini] Hirpida gaujoni
Hirpidinae [nec Hemileucinae] [nec Hemileucini] Hirpida levomaranensis
Hirpidinae [nec Hemileucinae] [nec Hemileucini] Hirpsinjaevia viksinjaevi
Ceratocampinae Dryocampini Citioica anthonilis
Ceratocampinae Dryocampini Citioica homoea
Ceratocampinae Dryocampini Oiticella luteciae
Ceratocampinae Dryocampini Oiticella convergens
Ceratocampinae Dryocampini Psilopygida walkeri
Ceratocampinae Dryocampini Psilopygida crispula
Ceratocampinae Dryocampini Ptiloscola santamartensis
Ceratocampinae Dryocampini Ptiloscola descimoni
Ceratocampinae Dryocampini Rachesa kaechi
Ceratocampinae Dryocampini Rachesa breteuili
Ceratocampinae Dryocampini Jaiba kesselringi
Ceratocampinae Dryocampini Mielkesia paranaensis
Ceratocampinae Dryocampini Megaceresa pulchra
Ceratocampinae Dryocampini Adelowalkeria plateada
Ceratocampinae Dryocampini Adelowalkeria tristygma
Ceratocampinae Dryocampini Ceratesa hemirhodia
Ceratocampinae Dryocampini Scolesa viettei
Ceratocampinae Dryocampini Ceropoda tibialis
Ceratocampinae Dryocampini Adeloneivaia catoxantha catoxantha
Ceratocampinae Dryocampini Adeloneivaia centrojason
Ceratocampinae Dryocampini Adeloneivaia boisduvalii
Ceratocampinae Dryocampini Almeidella almeidai
Ceratocampinae Dryocampini Giacomellia bilineata
Ceratocampinae Dryocampini Anisota leucostygma
Ceratocampinae Dryocampini Anisota assimilis
Ceratocampinae Dryocampini Dryocampa rubicunda
Ceratocampinae Dryocampini Syssphinx delmaiziana
Ceratocampinae Dryocampini Syssphinx thiaucourti
Ceratocampinae Dryocampini Syssphinx molina
Ceratocampinae Dryocampini Dacunju jucunda
Ceratocampinae Dryocampini Cicia pelota
Ceratocampinae Dryocampini Psilopygoides oda
Ceratocampinae Dryocampini Othorene purpurascens
Ceratocampinae Dryocampini Othorene intermedia
Ceratocampinae Dryocampini Othorene winbrechlini
Ceratocampinae Bathyphlebiini Eacles imperialis imperialis
Ceratocampinae Bathyphlebiini Eacles barragani
Ceratocampinae [nec Hemileucinae] Bathyphlebiini [nec Hemileucini] Eacles [nec Catharisa] cerina
Ceratocampinae Bathyphlebiini Eacles ducalis
Ceratocampinae Bathyphlebiini Bathyphlebia eminentoides
Ceratocampinae Bathyphlebiini Bathyphlebia rufescens
Ceratocampinae Bathyphlebiini Eacles ormondei violacea
Ceratocampinae Bathyphlebiini Schausiella subochreata
Ceratocampinae Bathyphlebiini Schausiella maranhensis
Ceratocampinae Bathyphlebiini Neorcarnegia basirei
Ceratocampinae Citheroniini Citheronia regalis
Ceratocampinae Citheroniini Citheronia phoronea
Ceratocampinae Citheroniini Citheronula armata
Ceratocampinae Citheroniini Citheronioides collaris
Ceratocampinae Citheroniini Procitheronia principalis
Ceratocampinae Citheroniini Procitheronia fenestrata
Cercophaninae Janiodini Janiodes ecarcabuco
Cercophaninae Janiodini Janiodes oxapampensis
Cercophaninae Janiodini Janiodes manzanoi
Cercophaninae Cercophanini Cercophana frauenfeldi
Cercophaninae Cercophanini Neocercophana philippii
Cercophaninae Cercophanini Microdulia mirabilis
Oxyteninae Therinia buckleyi
Oxyteninae Therinia sp
Oxyteninae Therinia transversaria
Oxyteninae Oxytenis naemia
Oxyteninae Oxytenis peregrina
Oxyteninae Oxytenis modoccidentalis
Oxyteninae Homoeopteryx pinchcarchensis
Sphingidae Smerinthinae Smerinthini Paonias myops
Sphingidae Smerinthinae Smerinthini Smerinthus ocellata
Sphingidae Smerinthinae Smerinthini Pachysphinx occidentalis
Sphingidae Smerinthinae Smerinthini Laothoe populi
Sphingidae Smerinthinae Coequosa australasiae
Sphingidae Sphinginae Sphingini Acherontia atropos
Sphingidae Sphinginae Sphingini Agrius convolvuli
Sphingidae Langiinae Langia zenzeroides
Sphingidae Macroglossinae Macroglossini Daphnis nerii
Sphingidae Macroglossinae Hemarini Hemaris fuciformis
Bombycidae Epiinae Quentalia mailynae
Endromidae Endromis versicolora
Brahmaeidae Acanthobrahmaea europaea
Brahmaeidae Lemonia taraxaci
Eupterotidae Janinae Stenoglene roseus
Eupterotidae Striphnopteryginae Phiala sp

**Figure S2I. Data subset (50% least saturated loci)**
**IQ-TREE unpartitioned**

Best fit model = GTR+F+R10 chosen according to BIC
SH-aLRT /UFBoot

The new classification proposed in this manuscript is used to annotate tips. Current classification is reported between square brackets

Nota : IQ-TREE analyses of data subsets were only performed without partitioning of the data to save computation time (and given that trees were similar when the initial data set was partitioned or not)

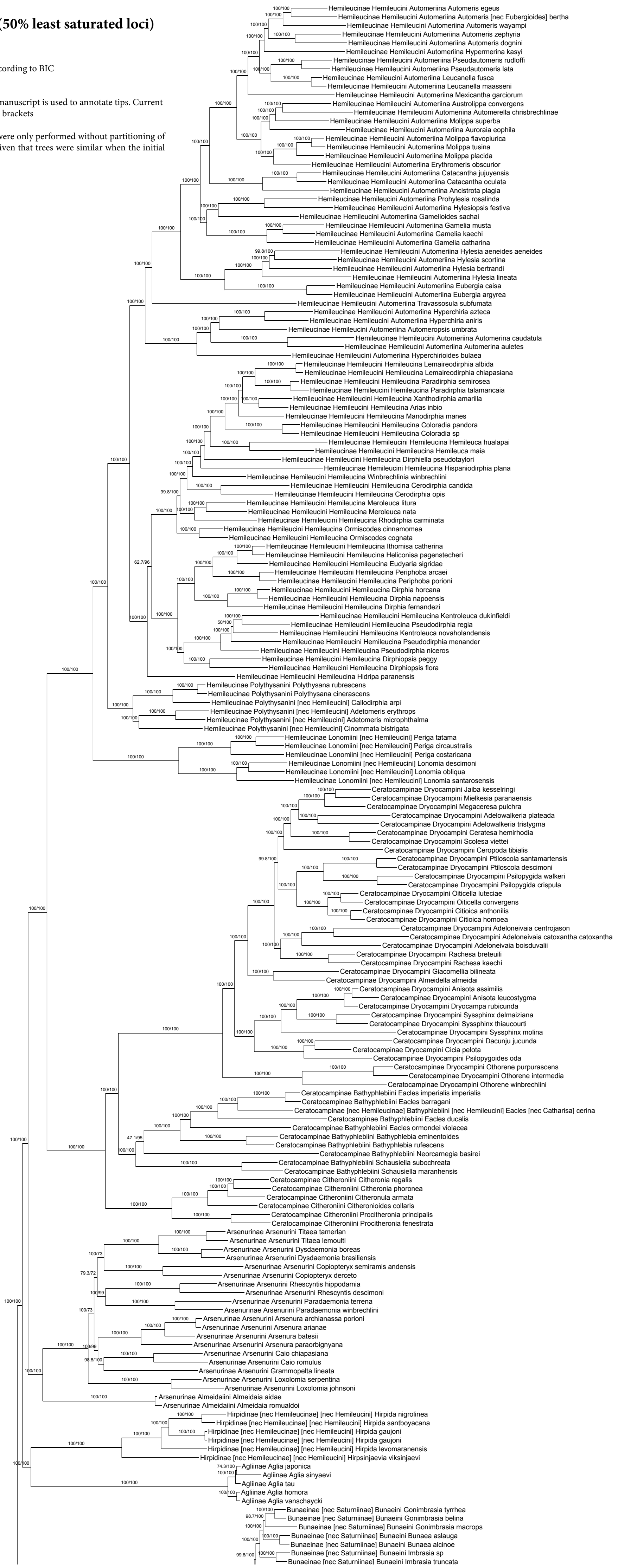

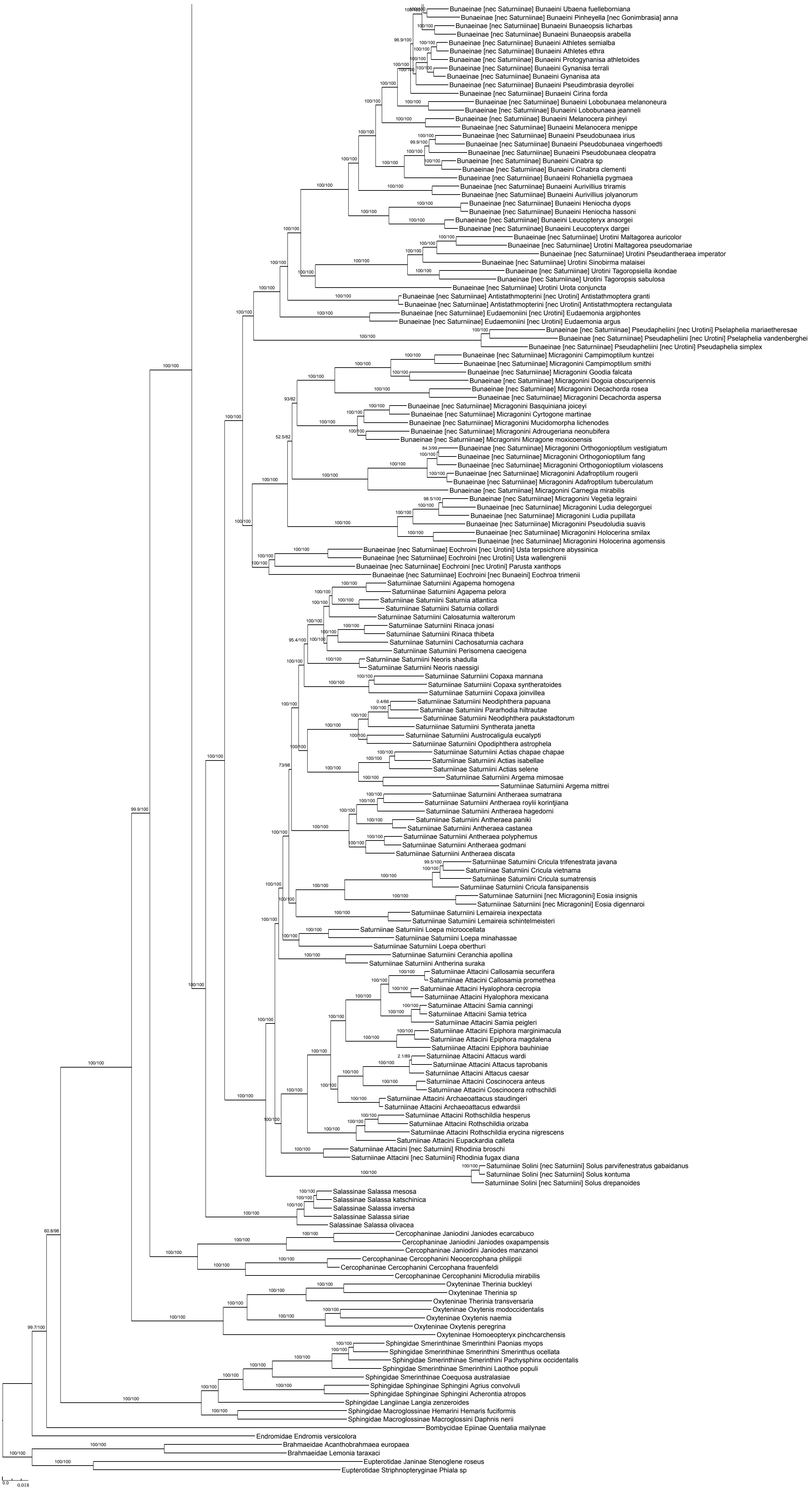
Bunaeinae [nec Saturniinae] Bunaeini Ubaena fuelleborniana
Bunaeinae [nec Saturniinae] Bunaeini Pinheyella [nec Gonimbrasia] anna
Bunaeinae [nec Saturniinae] Bunaeini Bunaeopsis licharbas
Bunaeinae [nec Saturniinae] Bunaeini Bunaeopsis arabella
Bunaeinae [nec Saturniinae] Bunaeini Athletes semialba
Bunaeinae [nec Saturniinae] Bunaeini Athletes ethra
Bunaeinae [nec Saturniinae] Bunaeini Protogynanisa athletoides
Bunaeinae [nec Saturniinae] Bunaeini Gynanisa terrali
Bunaeinae [nec Saturniinae] Bunaeini Gynanisa ata
Bunaeinae [nec Saturniinae] Bunaeini Pseudimbrasia deyrollei
Bunaeinae [nec Saturniinae] Bunaeini Cirina forda
Bunaeinae [nec Saturniinae] Bunaeini Lobobunaea melanoneura
Bunaeinae [nec Saturniinae] Bunaeini Lobobunaea jeanneli
Bunaeinae [nec Saturniinae] Bunaeini Melanocera pinheyi
Bunaeinae [nec Saturniinae] Bunaeini Melanocera menippe
Bunaeinae [nec Saturniinae] Bunaeini Pseudobunaea irius
Bunaeinae [nec Saturniinae] Bunaeini Pseudobunaea vingerhoedti
Bunaeinae [nec Saturniinae] Bunaeini Pseudobunaea cleopatra
Bunaeinae [nec Saturniinae] Bunaeini Cinabra sp
Bunaeinae [nec Saturniinae] Bunaeini Cinabra clementi
Bunaeinae [nec Saturniinae] Bunaeini Rohaniella pygmaea
Bunaeinae [nec Saturniinae] Bunaeini Aurivillius triramis
Bunaeinae [nec Saturniinae] Bunaeini Aurivillius jolyanorum
Bunaeinae [nec Saturniinae] Bunaeini Heniocha dyops
Bunaeinae [nec Saturniinae] Bunaeini Heniocha hassoni
Bunaeinae [nec Saturniinae] Bunaeini Leucopteryx ansorgei
Bunaeinae [nec Saturniinae] Bunaeini Leucopteryx dargei
Bunaeinae [nec Saturniinae] Urotini Maltagorea auricolor
Bunaeinae [nec Saturniinae] Urotini Maltagorea pseudomariae
Bunaeinae [nec Saturniinae] Urotini Pseudantheraea imperator
Bunaeinae [nec Saturniinae] Urotini Sinobirma malaisei
Bunaeinae [nec Saturniinae] Urotini Tagoropsiella ikondae
Bunaeinae [nec Saturniinae] Urotini Tagoropsis sabulosa
Bunaeinae [nec Saturniinae] Urotini Urota conjuncta
Bunaeinae [nec Saturniinae] Antistathmopterini [nec Urotini] Antistathmoptera granti
Bunaeinae [nec Saturniinae] Antistathmopterini [nec Urotini] Antistathmoptera rectangulata
Bunaeinae [nec Saturniinae] Eudaemoniini [nec Urotini] Eudaemonia argiphontes
Bunaeinae [nec Saturniinae] Eudaemoniini [nec Urotini] Eudaemonia argus
Bunaeinae [nec Saturniinae] Pseudapheliini [nec Urotini] Pselaphelia mariaetheresae
Bunaeinae [nec Saturniinae] Pseudapheliini [nec Urotini] Pselaphelia vandenberghei
Bunaeinae [nec Saturniinae] Pseudapheliini [nec Urotini] Pseudaphelia simplex
Bunaeinae [nec Saturniinae] Micragonini Campimoptilum kuntzei
Bunaeinae [nec Saturniinae] Micragonini Campimoptilum smithi
Bunaeinae [nec Saturniinae] Micragonini Goodia falcata
Bunaeinae [nec Saturniinae] Micragonini Dogoia obscuripennis
Bunaeinae [nec Saturniinae] Micragonini Decachorda rosea
Bunaeinae [nec Saturniinae] Micragonini Decachorda aspersa
Bunaeinae [nec Saturniinae] Micragonini Basquiniana joiceyi
Bunaeinae [nec Saturniinae] Micragonini Cyrtogone martinae
Bunaeinae [nec Saturniinae] Micragonini Mucidomorpha lichenodes
Bunaeinae [nec Saturniinae] Micragonini Adrougeriana neonubifera
Bunaeinae [nec Saturniinae] Micragonini Micragone moxicoensis
Bunaeinae [nec Saturniinae] Micragonini Orthogonioptilum vestigiatum
Bunaeinae [nec Saturniinae] Micragonini Orthogonioptilum fang
Bunaeinae [nec Saturniinae] Micragonini Orthogonioptilum violascens
Bunaeinae [nec Saturniinae] Micragonini Adafroptilum rougerii
Bunaeinae [nec Saturniinae] Micragonini Adafroptilum tuberculatum
Bunaeinae [nec Saturniinae] Micragonini Carnegia mirabilis
Bunaeinae [nec Saturniinae] Micragonini Vegetia legraini
Bunaeinae [nec Saturniinae] Micragonini Ludia delegorguei
Bunaeinae [nec Saturniinae] Micragonini Ludia pupillata
Bunaeinae [nec Saturniinae] Micragonini Pseudoludia suavis
Bunaeinae [nec Saturniinae] Micragonini Holocerina smilax
Bunaeinae [nec Saturniinae] Micragonini Holocerina agomensis
Bunaeinae [nec Saturniinae] Eochroini [nec Urotini] Usta terpsichore abyssinica
Bunaeinae [nec Saturniinae] Eochroini [nec Urotini] Usta wallengrenii
Bunaeinae [nec Saturniinae] Eochroini [nec Urotini] Parusta xanthops
Bunaeinae [nec Saturniinae] Eochroini [nec Bunaeini] Eochroa trimenii
Saturniinae Saturniini Agapema homogena
Saturniinae Saturniini Agapema pelora
Saturniinae Saturniini Saturnia atlantica
Saturniinae Saturniini Saturnia collardi
Saturniinae Saturniini Calosaturnia walterorum
Saturniinae Saturniini Rinaca jonasi
Saturniinae Saturniini Rinaca thibeta
Saturniinae Saturniini Cachosaturnia cachara
Saturniinae Saturniini Perisomena caecigena
Saturniinae Saturniini Neoris shadulla
Saturniinae Saturniini Neoris naessigi
Saturniinae Saturniini Copaxa mannana
Saturniinae Saturniini Copaxa syntheratoides
Saturniinae Saturniini Copaxa joinvillea
Saturniinae Saturniini Neodiphthera papuana
Saturniinae Saturniini Pararhodia hiltrautae
Saturniinae Saturniini Neodiphthera paukstadtorum
Saturniinae Saturniini Syntherata janetta
Saturniinae Saturniini Austrocaligula eucalypti
Saturniinae Saturniini Opodiphthera astrophela
Saturniinae Saturniini Actias chapae chapae
Saturniinae Saturniini Actias isabellae
Saturniinae Saturniini Actias selene
Saturniinae Saturniini Argema mimosae
Saturniinae Saturniini Argema mittrei
Saturniinae Saturniini Antheraea sumatrana
Saturniinae Saturniini Antheraea roylii korintjiana
Saturniinae Saturniini Antheraea hagedorni
Saturniinae Saturniini Antheraea paniki
Saturniinae Saturniini Antheraea castanea
Saturniinae Saturniini Antheraea polyphemus
Saturniinae Saturniini Antheraea godmani
Saturniinae Saturniini Antheraea discata
Saturniinae Saturniini Cricula trifenestrata javana
Saturniinae Saturniini Cricula vietnama
Saturniinae Saturniini Cricula sumatrensis
Saturniinae Saturniini Cricula fansipanensis
Saturniinae Saturniini [nec Micragonini] Eosia insignis
Saturniinae Saturniini [nec Micragonini] Eosia digennaroi
Saturniinae Saturniini Lemaireia inexpectata
Saturniinae Saturniini Lemaireia schintelmeisteri
Saturniinae Saturniini Loepa microocellata
Saturniinae Saturniini Loepa minahassae
Saturniinae Saturniini Loepa oberthuri
Saturniinae Saturniini Ceranchia apollina
Saturniinae Saturniini Antherina suraka
Saturniinae Attacini Callosamia securifera
Saturniinae Attacini Callosamia promethea
Saturniinae Attacini Hyalophora cecropia
Saturniinae Attacini Hyalophora mexicana
Saturniinae Attacini Samia canningi
Saturniinae Attacini Samia tetrica
Saturniinae Attacini Samia peigleri
Saturniinae Attacini Epiphora marginimacula
Saturniinae Attacini Epiphora magdalena
Saturniinae Attacini Epiphora bauhiniae
Saturniinae Attacini Attacus wardi
Saturniinae Attacini Attacus taprobanis
Saturniinae Attacini Attacus caesar
Saturniinae Attacini Coscinocera anteus
Saturniinae Attacini Coscinocera rothschildi
Saturniinae Attacini Archaeoattacus staudingeri
Saturniinae Attacini Archaeoattacus edwardsii
Saturniinae Attacini Rothschildia hesperus
Saturniinae Attacini Rothschildia orizaba
Saturniinae Attacini Rothschildia erycina nigrescens
Saturniinae Attacini Eupackardia calleta
Saturniinae Attacini [nec Saturniini] Rhodinia broschi
Saturniinae Attacini [nec Saturniini] Rhodinia fugax diana
Saturniinae Solini [nec Saturniini] Solus parvifenestratus gabaidanus
Saturniinae Solini [nec Saturniini] Solus kontuma
Saturniinae Solini [nec Saturniini] Solus drepanoides
Salassinae Salassa mesosa
Salassinae Salassa katschinica
Salassinae Salassa inversa
Salassinae Salassa siriae
Salassinae Salassa olivacea
Cercophaninae Janiodini Janiodes ecarcabuco
Cercophaninae Janiodini Janiodes oxapampensis
Cercophaninae Janiodini Janiodes manzanoi
Cercophaninae Cercophanini Neocercophana philippii
Cercophaninae Cercophanini Cercophana frauenfeldi
Cercophaninae Cercophanini Microdulia mirabilis
Oxyteninae Therinia buckleyi
Oxyteninae Therinia sp
Oxyteninae Therinia transversaria
Oxyteninae Oxytenis modocidentalis
Oxyteninae Oxytenis naemia
Oxyteninae Oxytenis peregrina
Oxyteninae Homoeopteryx pinchcarchensis
Sphingidae Smerinthinae Smerinthini Paonias myops
Sphingidae Smerinthinae Smerinthini Smerinthus ocellata
Sphingidae Smerinthinae Smerinthini Pachysphinx occidentalis
Sphingidae Smerinthinae Smerinthini Laothoe populi
Sphingidae Smerinthinae Coequosa australasiae
Sphingidae Sphinginae Sphingini Agrius convolvuli
Sphingidae Sphinginae Sphingini Acherontia atropos
Sphingidae Langiinae Langia zenzeroides
Sphingidae Macroglossinae Hemarini Hemaris fuciformis
Sphingidae Macroglossinae Macroglossini Daphnis nerii
Bombycidae Epiinae Quentalia mailynae
Endromidae Endromis versicolora
Brahmaeidae Acanthobrahmaea europaea
Brahmaeidae Lemonia taraxaci
Eupterotidae Janinae Stenoglene roseus
Eupterotidae Striphnopteryginae Phiala sp
0.0 0.018

#### Figure S2J. Data subset (50% least saturated loci) ASTRAL50

Species tree obtained when nodes with UFBoot<50 are collapsed in each gene tree

Final normalized quartet score : 0.93
Local posterior probabilities at nodes

The new classification proposed in this manuscript is used to annotate tips. Current classification is reported between square brackets

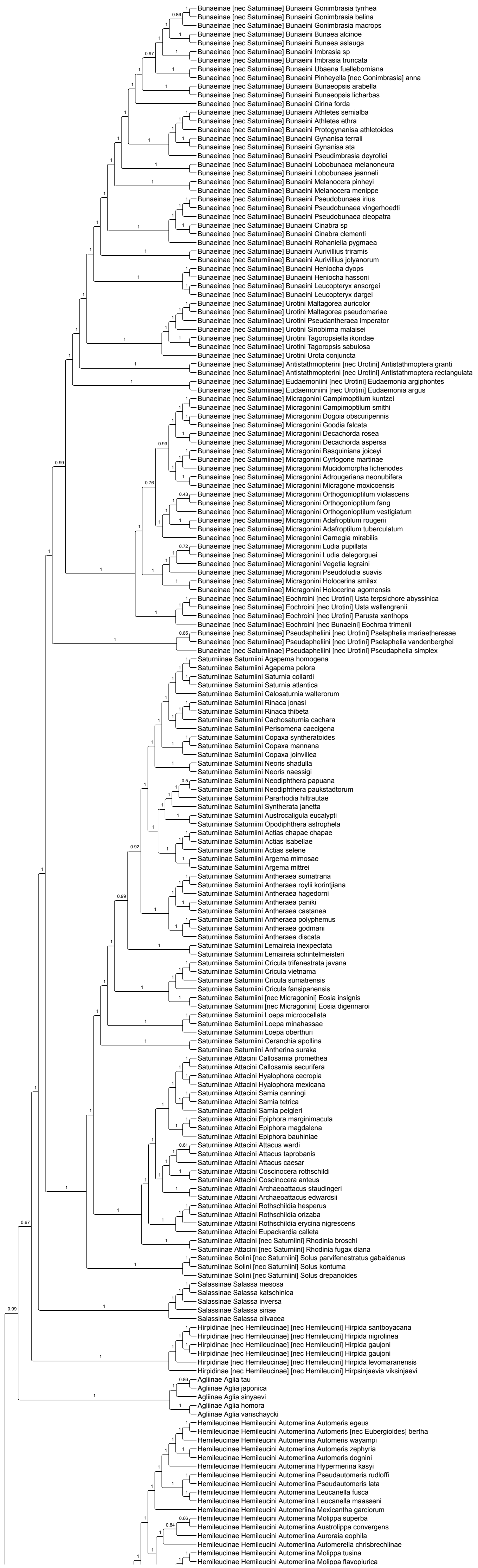

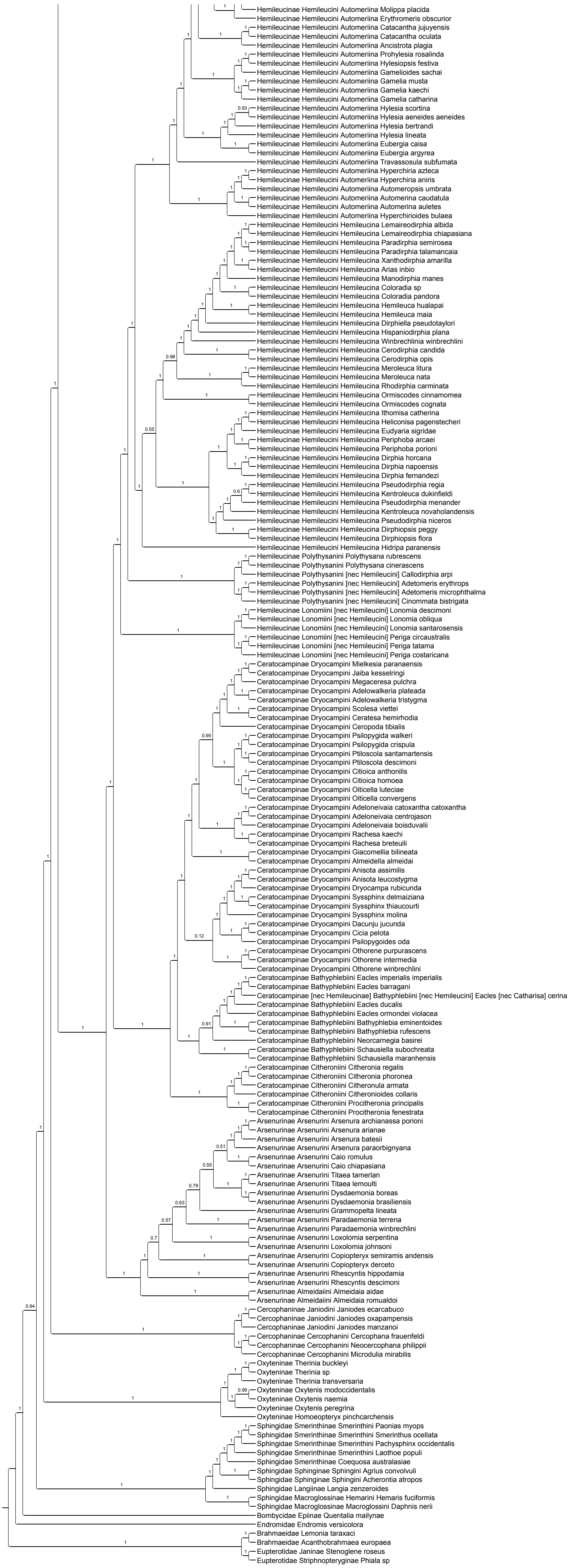
Hemileucinae Hemileucini Automeriina Molippa placida
Hemileucinae Hemileucini Automeriina Erythromeris obscurior
Hemileucinae Hemileucini Automeriina Catacantha jujuyensis
Hemileucinae Hemileucini Automeriina Catacantha oculata
Hemileucinae Hemileucini Automeriina Ancistrota plagia
Hemileucinae Hemileucini Automeriina Prohylesia rosalinda
Hemileucinae Hemileucini Automeriina Hylesiopsis festiva
Hemileucinae Hemileucini Automeriina Gamelioides sachai
Hemileucinae Hemileucini Automeriina Gamelia musta
Hemileucinae Hemileucini Automeriina Gamelia kaechi
Hemileucinae Hemileucini Automeriina Gamelia catharina
Hemileucinae Hemileucini Automeriina Hylesia scortina
Hemileucinae Hemileucini Automeriina Hylesia aeneides aeneides
Hemileucinae Hemileucini Automeriina Hylesia bertrandi
Hemileucinae Hemileucini Automeriina Hylesia lineata
Hemileucinae Hemileucini Automeriina Eubergia caisa
Hemileucinae Hemileucini Automeriina Eubergia argyrea
Hemileucinae Hemileucini Automeriina Travassosula subfumata
Hemileucinae Hemileucini Automeriina Hyperchiria azteca
Hemileucinae Hemileucini Automeriina Hyperchiria aniris
Hemileucinae Hemileucini Automeriina Automeropsis umbrata
Hemileucinae Hemileucini Automeriina Automerina caudatula
Hemileucinae Hemileucini Automeriina Automerina auletes
Hemileucinae Hemileucini Automeriina Hyperchirioides bulaea
Hemileucinae Hemileucini Hemileucina Lemaireodirphia albida
Hemileucinae Hemileucini Hemileucina Lemaireodirphia chiapasiana
Hemileucinae Hemileucini Hemileucina Paradirphia semirosea
Hemileucinae Hemileucini Hemileucina Paradirphia talamancaia
Hemileucinae Hemileucini Hemileucina Xanthodirphia amarilla
Hemileucinae Hemileucini Hemileucina Arias inbio
Hemileucinae Hemileucini Hemileucina Manodirphia manes
Hemileucinae Hemileucini Hemileucina Coloradia sp
Hemileucinae Hemileucini Hemileucina Coloradia pandora
Hemileucinae Hemileucini Hemileucina Hemileuca hualapai
Hemileucinae Hemileucini Hemileucina Hemileuca maia
Hemileucinae Hemileucini Hemileucina Dirphiella pseudotaylori
Hemileucinae Hemileucini Hemileucina Hispaniodirphia plana
Hemileucinae Hemileucini Hemileucina Winbrechlinia winbrechlini
Hemileucinae Hemileucini Hemileucina Cerodirphia candida
Hemileucinae Hemileucini Hemileucina Cerodirphia opis
Hemileucinae Hemileucini Hemileucina Meroleuca litura
Hemileucinae Hemileucini Hemileucina Meroleuca nata
Hemileucinae Hemileucini Hemileucina Rhodirphia carminata
Hemileucinae Hemileucini Hemileucina Ormiscodes cinnamomea
Hemileucinae Hemileucini Hemileucina Ormiscodes cognata
Hemileucinae Hemileucini Hemileucina Ithomisa catherina
Hemileucinae Hemileucini Hemileucina Heliconisa pagenstecheri
Hemileucinae Hemileucini Hemileucina Eudyaria sigridae
Hemileucinae Hemileucini Hemileucina Periphoba arcaei
Hemileucinae Hemileucini Hemileucina Periphoba porioni
Hemileucinae Hemileucini Hemileucina Dirphia horcana
Hemileucinae Hemileucini Hemileucina Dirphia napoensis
Hemileucinae Hemileucini Hemileucina Dirphia fernandezi
Hemileucinae Hemileucini Hemileucina Pseudodirphia regia
Hemileucinae Hemileucini Hemileucina Kentroleuca dukinfieldi
Hemileucinae Hemileucini Hemileucina Pseudodirphia menander
Hemileucinae Hemileucini Hemileucina Kentroleuca novaholandensis
Hemileucinae Hemileucini Hemileucina Pseudodirphia niceros
Hemileucinae Hemileucini Hemileucina Dirphiopsis peggy
Hemileucinae Hemileucini Hemileucina Dirphiopsis flora
Hemileucinae Hemileucini Hemileucina Hidripa paranensis
Hemileucinae Polythysanini Polythysana rubrescens
Hemileucinae Polythysanini Polythysana cinerascens
Hemileucinae Polythysanini [nec Hemileucini] Callodirphia arpi
Hemileucinae Polythysanini [nec Hemileucini] Adetomeris erythrops
Hemileucinae Polythysanini [nec Hemileucini] Adetomeris microphthalma
Hemileucinae Polythysanini [nec Hemileucini] Cinommata bistrigata
Hemileucinae Lonomiini [nec Hemileucini] Lonomia descimoni
Hemileucinae Lonomiini [nec Hemileucini] Lonomia obliqua
Hemileucinae Lonomiini [nec Hemileucini] Lonomia santarosensis
Hemileucinae Lonomiini [nec Hemileucini] Periga circaustralis
Hemileucinae Lonomiini [nec Hemileucini] Periga tatama
Hemileucinae Lonomiini [nec Hemileucini] Periga costaricana
Ceratocampinae Dryocampini Mielkesia paranaensis
Ceratocampinae Dryocampini Jaiba kesselringi
Ceratocampinae Dryocampini Megaceresa pulchra
Ceratocampinae Dryocampini Adelowalkeria plateada
Ceratocampinae Dryocampini Adelowalkeria tristygma
Ceratocampinae Dryocampini Scolesa viettei
Ceratocampinae Dryocampini Ceratesa hemirhodia
Ceratocampinae Dryocampini Ceropoda tibialis
Ceratocampinae Dryocampini Psilopygida walkeri
Ceratocampinae Dryocampini Psilopygida crispula
Ceratocampinae Dryocampini Ptiloscola santamartensis
Ceratocampinae Dryocampini Ptiloscola descimoni
Ceratocampinae Dryocampini Citioica anthonilis
Ceratocampinae Dryocampini Citioica homoea
Ceratocampinae Dryocampini Oiticella luteciae
Ceratocampinae Dryocampini Oiticella convergens
Ceratocampinae Dryocampini Adeloneivaia catoxantha catoxantha
Ceratocampinae Dryocampini Adeloneivaia centrojason
Ceratocampinae Dryocampini Adeloneivaia boisduvalii
Ceratocampinae Dryocampini Rachesa kaechi
Ceratocampinae Dryocampini Rachesa breteuili
Ceratocampinae Dryocampini Giacomellia bilineata
Ceratocampinae Dryocampini Almeidella almeidai
Ceratocampinae Dryocampini Anisota assimilis
Ceratocampinae Dryocampini Anisota leucostygma
Ceratocampinae Dryocampini Dryocampa rubicunda
Ceratocampinae Dryocampini Syssphinx delmaiziana
Ceratocampinae Dryocampini Syssphinx thiaucourti
Ceratocampinae Dryocampini Syssphinx molina
Ceratocampinae Dryocampini Dacunju jucunda
Ceratocampinae Dryocampini Cicia pelota
Ceratocampinae Dryocampini Psilopygoides oda
Ceratocampinae Dryocampini Othorene purpurascens
Ceratocampinae Dryocampini Othorene intermedia
Ceratocampinae Dryocampini Othorene winbrechlini
Ceratocampinae Bathyphlebiini Eacles imperialis imperialis
Ceratocampinae Bathyphlebiini Eacles barragani
Ceratocampinae [nec Hemileucinae] Bathyphlebiini [nec Hemileucini] Eacles [nec Catharisa] cerina
Ceratocampinae Bathyphlebiini Eacles ducalis
Ceratocampinae Bathyphlebiini Eacles ormondei violacea
Ceratocampinae Bathyphlebiini Bathyphlebia eminentoides
Ceratocampinae Bathyphlebiini Bathyphlebia rufescens
Ceratocampinae Bathyphlebiini Neorcarnegia basirei
Ceratocampinae Bathyphlebiini Schausiella subochreata
Ceratocampinae Bathyphlebiini Schausiella maranhensis
Ceratocampinae Citheroniini Citheronia regalis
Ceratocampinae Citheroniini Citheronia phoronea
Ceratocampinae Citheroniini Citheronula armata
Ceratocampinae Citheroniini Citheronioides collaris
Ceratocampinae Citheroniini Procitheronia principalis
Ceratocampinae Citheroniini Procitheronia fenestrata
Arsenurinae Arsenurini Arsenura archianassa porioni
Arsenurinae Arsenurini Arsenura arianae
Arsenurinae Arsenurini Arsenura batesii
Arsenurinae Arsenurini Arsenura paraorbignyana
Arsenurinae Arsenurini Caio romulus
Arsenurinae Arsenurini Caio chiapasiana
Arsenurinae Arsenurini Titaea tamerlan
Arsenurinae Arsenurini Titaea lemoulti
Arsenurinae Arsenurini Dysdaemonia boreas
Arsenurinae Arsenurini Dysdaemonia brasiliensis
Arsenurinae Arsenurini Grammopelta lineata
Arsenurinae Arsenurini Paradaemonia terrena
Arsenurinae Arsenurini Paradaemonia winbrechlini
Arsenurinae Arsenurini Loxolomia serpentina
Arsenurinae Arsenurini Loxolomia johnsoni
Arsenurinae Arsenurini Copiopteryx semiramis andensis
Arsenurinae Arsenurini Copiopteryx derceto
Arsenurinae Arsenurini Rhescyntis hippodamia
Arsenurinae Arsenurini Rhescyntis descimoni
Arsenurinae Almeidaiini Almeidaia aidae
Arsenurinae Almeidaiini Almeidaia romualdoi
Cercophaninae Janiodini Janiodes ecarcabuco
Cercophaninae Janiodini Janiodes oxapampensis
Cercophaninae Janiodini Janiodes manzanoi
Cercophaninae Cercophanini Cercophana frauenfeldi
Cercophaninae Cercophanini Neocercophana philippii
Cercophaninae Cercophanini Microdulia mirabilis
Oxyteninae Therinia buckleyi
Oxyteninae Therinia sp
Oxyteninae Therinia transversaria
Oxyteninae Oxytenis modocidentalis
Oxyteninae Oxytenis naemia
Oxyteninae Oxytenis peregrina
Oxyteninae Homoeopteryx pinchcarchensis
Sphingidae Smerinthinae Smerinthini Paonias myops
Sphingidae Smerinthinae Smerinthini Smerinthus ocellata
Sphingidae Smerinthinae Smerinthini Pachysphinx occidentalis
Sphingidae Smerinthinae Smerinthini Laothoe populi
Sphingidae Smerinthinae Coequosa australasiae
Sphingidae Sphinginae Sphingini Agrius convolvuli
Sphingidae Sphinginae Sphingini Acherontia atropos
Sphingidae Langiinae Langia zenzeroides
Sphingidae Macroglossinae Hemarini Hemaris fuciformis
Sphingidae Macroglossinae Macroglossini Daphnis nerii
Bombycidae Epiinae Quentalia mailynae
Endromidae Endromis versicolora
Brahmaeidae Lemonia taraxaci
Brahmaeidae Acanthobrahmaea europaea
Eupterotidae Janinae Stenoglene roseus
Eupterotidae Striphnopteryginae Phiala sp

### Figure S2K. Data subset (50% least saturated loci) ASTRAL70

Species tree obtained when nodes with UFBoot<70 are collapsed in each gene tree

Final normalized quartet score : 0.95
Local posterior probabilities at nodes

The new classification proposed in this manuscript is used to annotate tips. Current classification is reported between square brackets

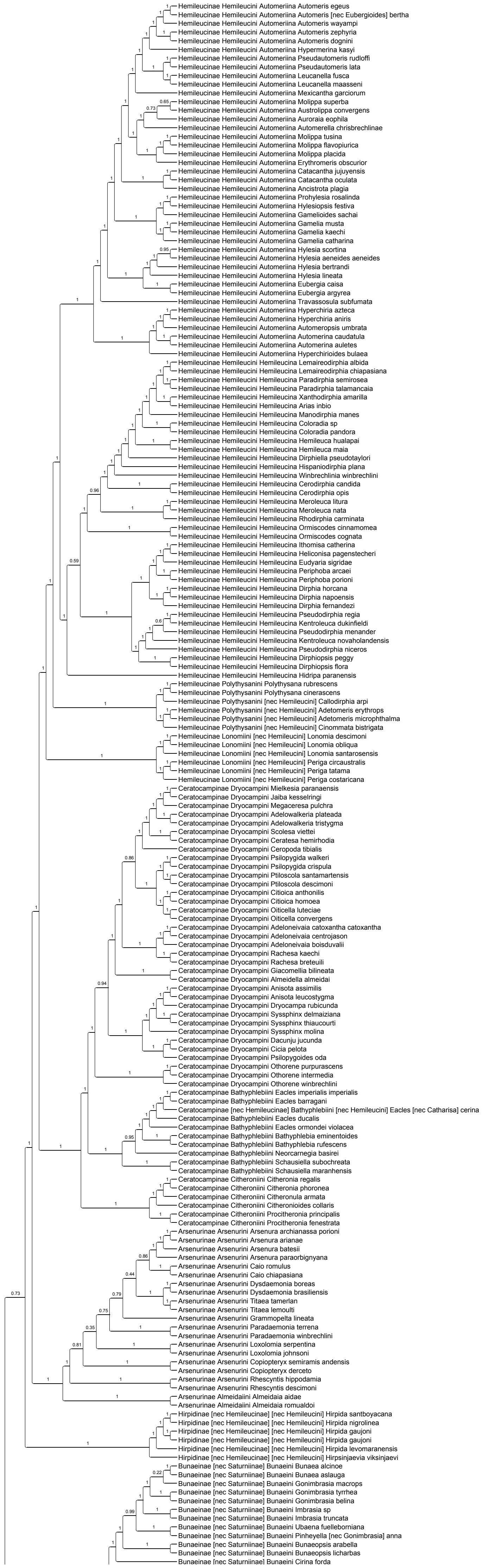

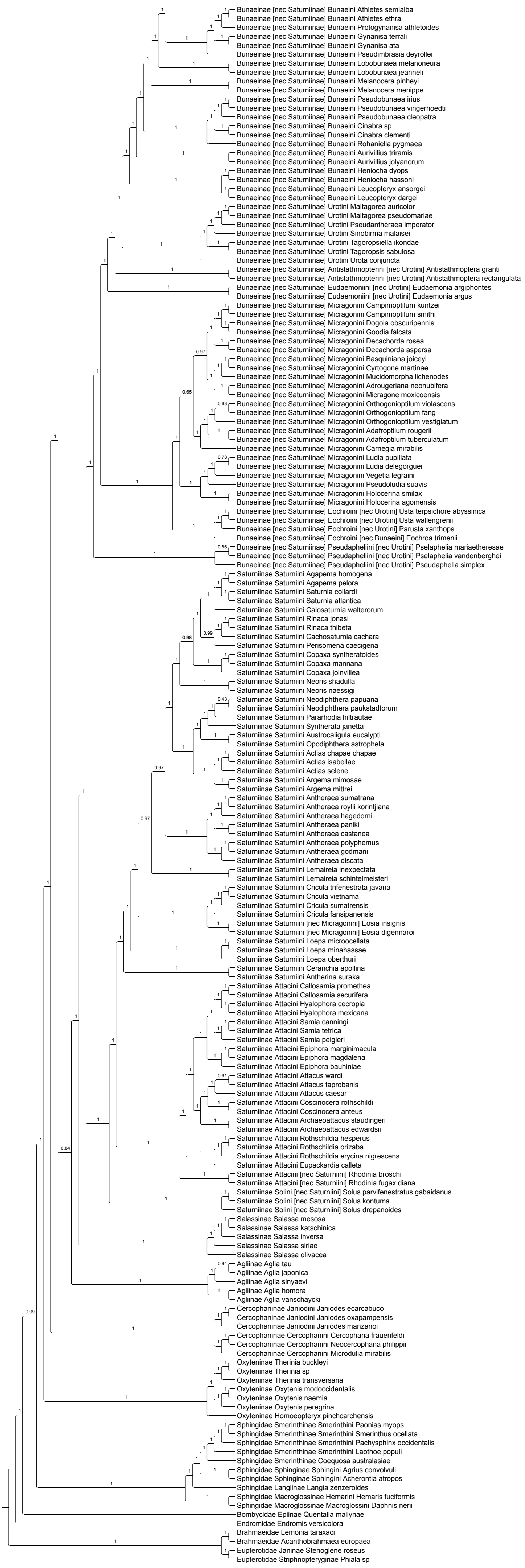
Bunaeinae [nec Saturniinae] Bunaeini Athletes semialba
Bunaeinae [nec Saturniinae] Bunaeini Athletes ethra
Bunaeinae [nec Saturniinae] Bunaeini Protogynanisa athletoides
Bunaeinae [nec Saturniinae] Bunaeini Gynanisa terrali
Bunaeinae [nec Saturniinae] Bunaeini Gynanisa ata
Bunaeinae [nec Saturniinae] Bunaeini Pseudimbrasia deyrollei
Bunaeinae [nec Saturniinae] Bunaeini Lobobunaea melanoneura
Bunaeinae [nec Saturniinae] Bunaeini Lobobunaea jeanneli
Bunaeinae [nec Saturniinae] Bunaeini Melanocera pinheyi
Bunaeinae [nec Saturniinae] Bunaeini Melanocera menippe
Bunaeinae [nec Saturniinae] Bunaeini Pseudobunaea irius
Bunaeinae [nec Saturniinae] Bunaeini Pseudobunaea vingerhoedti
Bunaeinae [nec Saturniinae] Bunaeini Pseudobunaea cleopatra
Bunaeinae [nec Saturniinae] Bunaeini Cinabra sp
Bunaeinae [nec Saturniinae] Bunaeini Cinabra clementi
Bunaeinae [nec Saturniinae] Bunaeini Rohaniella pygmaea
Bunaeinae [nec Saturniinae] Bunaeini Aurivillius triramis
Bunaeinae [nec Saturniinae] Bunaeini Aurivillius jolyanorum
Bunaeinae [nec Saturniinae] Bunaeini Heniocha dyops
Bunaeinae [nec Saturniinae] Bunaeini Heniocha hassoni
Bunaeinae [nec Saturniinae] Bunaeini Leucopteryx ansorgei
Bunaeinae [nec Saturniinae] Bunaeini Leucopteryx dargei
Bunaeinae [nec Saturniinae] Urotini Maltagorea auricolor
Bunaeinae [nec Saturniinae] Urotini Maltagorea pseudomariae
Bunaeinae [nec Saturniinae] Urotini Pseudantheraea imperator
Bunaeinae [nec Saturniinae] Urotini Sinobirma malaisei
Bunaeinae [nec Saturniinae] Urotini Tagoropsiella ikondae
Bunaeinae [nec Saturniinae] Urotini Tagoropsis sabulosa
Bunaeinae [nec Saturniinae] Urotini Urota conjuncta
Bunaeinae [nec Saturniinae] Antistathmopterini [nec Urotini] Antistathmoptera granti
Bunaeinae [nec Saturniinae] Antistathmopterini [nec Urotini] Antistathmoptera rectangulata
Bunaeinae [nec Saturniinae] Eudaemoniini [nec Urotini] Eudaemonia argiphontes
Bunaeinae [nec Saturniinae] Eudaemoniini [nec Urotini] Eudaemonia argus
Bunaeinae [nec Saturniinae] Micragonini Campimoptilum kuntzei
Bunaeinae [nec Saturniinae] Micragonini Campimoptilum smithi
Bunaeinae [nec Saturniinae] Micragonini Dogoia obscuripennis
Bunaeinae [nec Saturniinae] Micragonini Goodia falcata
Bunaeinae [nec Saturniinae] Micragonini Decachorda rosea
Bunaeinae [nec Saturniinae] Micragonini Decachorda aspersa
Bunaeinae [nec Saturniinae] Micragonini Basquiniana joiceyi
Bunaeinae [nec Saturniinae] Micragonini Cyrtogone martinae
Bunaeinae [nec Saturniinae] Micragonini Mucidomorpha lichenodes
Bunaeinae [nec Saturniinae] Micragonini Adrougeriana neonubifera
Bunaeinae [nec Saturniinae] Micragonini Micragone moxicoensis
Bunaeinae [nec Saturniinae] Micragonini Orthogonioptilum violascens
Bunaeinae [nec Saturniinae] Micragonini Orthogonioptilum fang
Bunaeinae [nec Saturniinae] Micragonini Orthogonioptilum vestigiatum
Bunaeinae [nec Saturniinae] Micragonini Adafroptilum rougerii
Bunaeinae [nec Saturniinae] Micragonini Adafroptilum tuberculatum
Bunaeinae [nec Saturniinae] Micragonini Carnegia mirabilis
Bunaeinae [nec Saturniinae] Micragonini Ludia pupillata
Bunaeinae [nec Saturniinae] Micragonini Ludia delegorguei
Bunaeinae [nec Saturniinae] Micragonini Vegetia legraini
Bunaeinae [nec Saturniinae] Micragonini Pseudoludia suavis
Bunaeinae [nec Saturniinae] Micragonini Holocerina smilax
Bunaeinae [nec Saturniinae] Micragonini Holocerina agomensis
Bunaeinae [nec Saturniinae] Eochroini [nec Urotini] Usta terpsichore abyssinica
Bunaeinae [nec Saturniinae] Eochroini [nec Urotini] Usta wallengrenii
Bunaeinae [nec Saturniinae] Eochroini [nec Urotini] Parusta xanthops
Bunaeinae [nec Saturniinae] Eochroini [nec Bunaeini] Eochroa trimenii
Bunaeinae [nec Saturniinae] Pseudapheliini [nec Urotini] Pselaphelia mariaetheresae
Bunaeinae [nec Saturniinae] Pseudapheliini [nec Urotini] Pselaphelia vandenberghei
Bunaeinae [nec Saturniinae] Pseudapheliini [nec Urotini] Pseudaphelia simplex
Saturniinae Saturniini Agapema homogena
Saturniinae Saturniini Agapema pelora
Saturniinae Saturniini Saturnia collardi
Saturniinae Saturniini Saturnia atlantica
Saturniinae Saturniini Calosaturnia walterorum
Saturniinae Saturniini Rinaca jonasi
Saturniinae Saturniini Rinaca thibeta
Saturniinae Saturniini Cachosaturnia cachara
Saturniinae Saturniini Perisomena caecigena
Saturniinae Saturniini Copaxa syntheratoides
Saturniinae Saturniini Copaxa mannana
Saturniinae Saturniini Copaxa joinvillea
Saturniinae Saturniini Neoris shadulla
Saturniinae Saturniini Neoris naessigi
Saturniinae Saturniini Neodiphthera papuana
Saturniinae Saturniini Neodiphthera paukstadtorum
Saturniinae Saturniini Pararhodia hiltrautae
Saturniinae Saturniini Syntherata janetta
Saturniinae Saturniini Austrocaligula eucalypti
Saturniinae Saturniini Opodiphthera astrophela
Saturniinae Saturniini Actias chapae chapae
Saturniinae Saturniini Actias isabellae
Saturniinae Saturniini Actias selene
Saturniinae Saturniini Argema mimosae
Saturniinae Saturniini Argema mittrei
Saturniinae Saturniini Antheraea sumatrana
Saturniinae Saturniini Antheraea roylii korintjiana
Saturniinae Saturniini Antheraea hagedorni
Saturniinae Saturniini Antheraea paniki
Saturniinae Saturniini Antheraea castanea
Saturniinae Saturniini Antheraea polyphemus
Saturniinae Saturniini Antheraea godmani
Saturniinae Saturniini Antheraea discata
Saturniinae Saturniini Lemaireia inexpectata
Saturniinae Saturniini Lemaireia schintelmeisteri
Saturniinae Saturniini Cricula trifenestrata javana
Saturniinae Saturniini Cricula vietnama
Saturniinae Saturniini Cricula sumatrensis
Saturniinae Saturniini Cricula fansipanensis
Saturniinae Saturniini [nec Micragonini] Eosia insignis
Saturniinae Saturniini [nec Micragonini] Eosia digennaroi
Saturniinae Saturniini Loepa microocellata
Saturniinae Saturniini Loepa minahassae
Saturniinae Saturniini Loepa oberthuri
Saturniinae Saturniini Ceranchia apollina
Saturniinae Saturniini Antherina suraka
Saturniinae Attacini Callosamia promethea
Saturniinae Attacini Callosamia securifera
Saturniinae Attacini Hyalophora cecropia
Saturniinae Attacini Hyalophora mexicana
Saturniinae Attacini Samia canningi
Saturniinae Attacini Samia tetrica
Saturniinae Attacini Samia peigleri
Saturniinae Attacini Epiphora marginimacula
Saturniinae Attacini Epiphora magdalena
Saturniinae Attacini Epiphora bauhiniae
Saturniinae Attacini Attacus wardi
Saturniinae Attacini Attacus taprobanis
Saturniinae Attacini Attacus caesar
Saturniinae Attacini Coscinocera rothschildi
Saturniinae Attacini Coscinocera anteus
Saturniinae Attacini Archaeoattacus staudingeri
Saturniinae Attacini Archaeoattacus edwardsii
Saturniinae Attacini Rothschildia hesperus
Saturniinae Attacini Rothschildia orizaba
Saturniinae Attacini Rothschildia erycina nigrescens
Saturniinae Attacini Eupackardia calleta
Saturniinae Attacini [nec Saturniini] Rhodinia broschi
Saturniinae Attacini [nec Saturniini] Rhodinia fugax diana
Saturniinae Solini [nec Saturniini] Solus parvifenestratus gabaidanus
Saturniinae Solini [nec Saturniini] Solus kontuma
Saturniinae Solini [nec Saturniini] Solus drepanoides
Salassinae Salassa mesosa
Salassinae Salassa katschinica
Salassinae Salassa inversa
Salassinae Salassa siriae
Salassinae Salassa olivacea
Agliinae Aglia tau
Agliinae Aglia japonica
Agliinae Aglia sinyaevi
Agliinae Aglia homora
Agliinae Aglia vanschaycki
Cercophaninae Janiodini Janiodes ecarcabuco
Cercophaninae Janiodini Janiodes oxapampensis
Cercophaninae Janiodini Janiodes manzanoi
Cercophaninae Cercophanini Cercophana frauenfeldi
Cercophaninae Cercophanini Neocercophana philippii
Cercophaninae Cercophanini Microdulia mirabilis
Oxyteninae Therinia buckleyi
Oxyteninae Therinia sp
Oxyteninae Therinia transversaria
Oxyteninae Oxytenis modocccidentalis
Oxyteninae Oxytenis naemia
Oxyteninae Oxytenis peregrina
Oxyteninae Homoeopteryx pinchcarchensis
Sphingidae Smerinthinae Smerinthini Paonias myops
Sphingidae Smerinthinae Smerinthini Smerinthus ocellata
Sphingidae Smerinthinae Smerinthini Pachysphinx occidentalis
Sphingidae Smerinthinae Smerinthini Laothoe populi
Sphingidae Smerinthinae Coequosa australasiae
Sphingidae Sphinginae Sphingini Agrius convolvuli
Sphingidae Sphinginae Sphingini Acherontia atropos
Sphingidae Langiinae Langia zenzeroides
Sphingidae Macroglossinae Hemarini Hemaris fuciformis
Sphingidae Macroglossinae Macroglossini Daphnis nerii
Bombycidae Epiinae Quentalia mailynae
Endromidae Endromis versicolora
Brahmaeidae Lemonia taraxaci
Brahmaeidae Acanthobrahmaea europaea
Eupterotidae Janinae Stenoglene roseus
Eupterotidae Striphnopteryginae Phiala sp

### Figure S2L. Data subset (50% least saturated loci) ASTRAL90

Species tree obtained when nodes with UFBoot<90 are collapsed in each gene tree

Final normalized quartet score : 0.98
Local posterior probabilities at nodes

The new classification proposed in this manuscript is used to annotate tips. Current classification is reported between square brackets

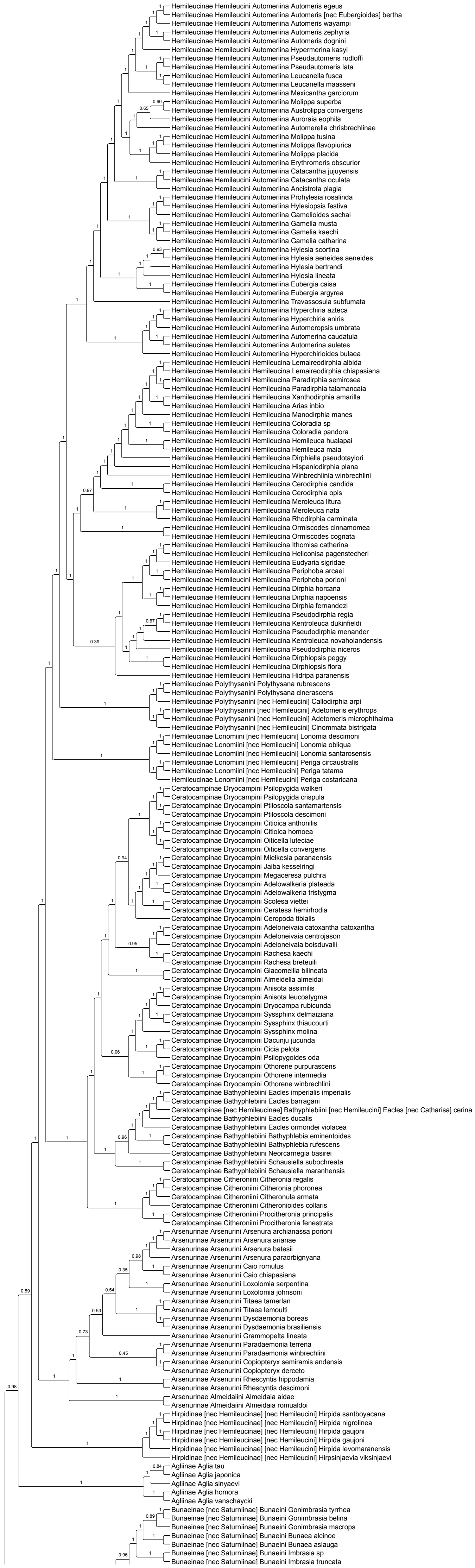

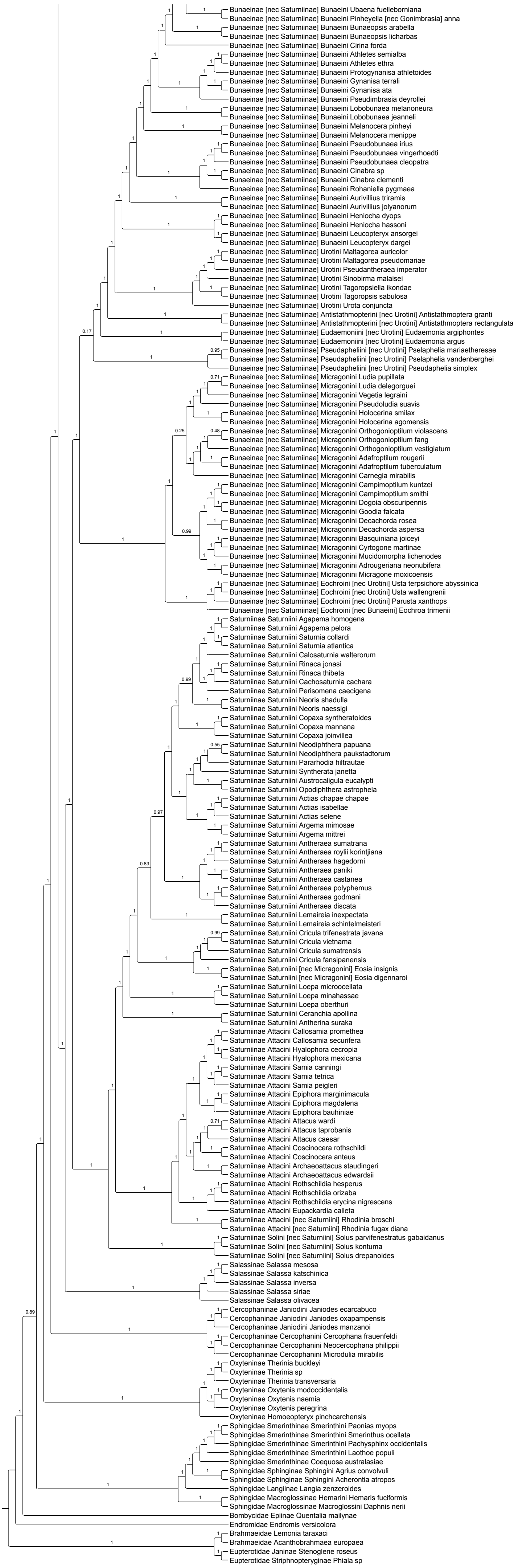

Bunaeinae [nec Saturniinae] Bunaeini Ubaena fuelleborniana
Bunaeinae [nec Saturniinae] Bunaeini Pinheyella [nec Gonimbrasia] anna
Bunaeinae [nec Saturniinae] Bunaeini Bunaeopsis arabella
Bunaeinae [nec Saturniinae] Bunaeini Bunaeopsis licharbas
Bunaeinae [nec Saturniinae] Bunaeini Cirina forda
Bunaeinae [nec Saturniinae] Bunaeini Athletes semialba
Bunaeinae [nec Saturniinae] Bunaeini Athletes ethra
Bunaeinae [nec Saturniinae] Bunaeini Protogynanisa athletoides
Bunaeinae [nec Saturniinae] Bunaeini Gynanisa terrali
Bunaeinae [nec Saturniinae] Bunaeini Gynanisa ata
Bunaeinae [nec Saturniinae] Bunaeini Pseudimbrasia deyrollei
Bunaeinae [nec Saturniinae] Bunaeini Lobobunaea melanoneura
Bunaeinae [nec Saturniinae] Bunaeini Lobobunaea jeanneli
Bunaeinae [nec Saturniinae] Bunaeini Melanocera pinheyi
Bunaeinae [nec Saturniinae] Bunaeini Melanocera menippe
Bunaeinae [nec Saturniinae] Bunaeini Pseudobunaea irius
Bunaeinae [nec Saturniinae] Bunaeini Pseudobunaea vingerhoedti
Bunaeinae [nec Saturniinae] Bunaeini Pseudobunaea cleopatra
Bunaeinae [nec Saturniinae] Bunaeini Cinabra sp
Bunaeinae [nec Saturniinae] Bunaeini Cinabra clementi
Bunaeinae [nec Saturniinae] Bunaeini Rohaniella pygmaea
Bunaeinae [nec Saturniinae] Bunaeini Aurivillius triramis
Bunaeinae [nec Saturniinae] Bunaeini Aurivillius jolyanorum
Bunaeinae [nec Saturniinae] Bunaeini Heniocha dyops
Bunaeinae [nec Saturniinae] Bunaeini Heniocha hassoni
Bunaeinae [nec Saturniinae] Bunaeini Leucopteryx ansorgei
Bunaeinae [nec Saturniinae] Bunaeini Leucopteryx dargei
Bunaeinae [nec Saturniinae] Urotini Maltagorea auricolor
Bunaeinae [nec Saturniinae] Urotini Maltagorea pseudomariae
Bunaeinae [nec Saturniinae] Urotini Pseudantheraea imperator
Bunaeinae [nec Saturniinae] Urotini Sinobirma malaisei
Bunaeinae [nec Saturniinae] Urotini Tagoropsiella ikondae
Bunaeinae [nec Saturniinae] Urotini Tagoropsis sabulosa
Bunaeinae [nec Saturniinae] Urotini Urota conjuncta
Bunaeinae [nec Saturniinae] Antistathmopterini [nec Urotini] Antistathmoptera granti
Bunaeinae [nec Saturniinae] Antistathmopterini [nec Urotini] Antistathmoptera rectangulata
Bunaeinae [nec Saturniinae] Eudaemoniini [nec Urotini] Eudaemonia argiphontes
Bunaeinae [nec Saturniinae] Eudaemoniini [nec Urotini] Eudaemonia argus
Bunaeinae [nec Saturniinae] Pseudapheliini [nec Urotini] Pselaphelia mariaetheresae
Bunaeinae [nec Saturniinae] Pseudapheliini [nec Urotini] Pselaphelia vandenberghei
Bunaeinae [nec Saturniinae] Pseudapheliini [nec Urotini] Pseudaphelia simplex
Bunaeinae [nec Saturniinae] Micragonini Ludia pupillata
Bunaeinae [nec Saturniinae] Micragonini Ludia delegorguei
Bunaeinae [nec Saturniinae] Micragonini Vegetia legraini
Bunaeinae [nec Saturniinae] Micragonini Pseudoludia suavis
Bunaeinae [nec Saturniinae] Micragonini Holocerina smilax
Bunaeinae [nec Saturniinae] Micragonini Holocerina agomensis
Bunaeinae [nec Saturniinae] Micragonini Orthogonioptilum violascens
Bunaeinae [nec Saturniinae] Micragonini Orthogonioptilum fang
Bunaeinae [nec Saturniinae] Micragonini Orthogonioptilum vestigiatum
Bunaeinae [nec Saturniinae] Micragonini Adafroptilum rougerii
Bunaeinae [nec Saturniinae] Micragonini Adafroptilum tuberculatum
Bunaeinae [nec Saturniinae] Micragonini Carnegia mirabilis
Bunaeinae [nec Saturniinae] Micragonini Campimoptilum kuntzei
Bunaeinae [nec Saturniinae] Micragonini Campimoptilum smithi
Bunaeinae [nec Saturniinae] Micragonini Dogoia obscuripennis
Bunaeinae [nec Saturniinae] Micragonini Goodia falcata
Bunaeinae [nec Saturniinae] Micragonini Decachorda rosea
Bunaeinae [nec Saturniinae] Micragonini Decachorda aspersa
Bunaeinae [nec Saturniinae] Micragonini Basquiniana joiceyi
Bunaeinae [nec Saturniinae] Micragonini Cyrtogone martinae
Bunaeinae [nec Saturniinae] Micragonini Mucidomorpha lichenodes
Bunaeinae [nec Saturniinae] Micragonini Adrougeriana neonubifera
Bunaeinae [nec Saturniinae] Micragonini Micragone moxicoensis
Bunaeinae [nec Saturniinae] Eochroini [nec Urotini] Usta terpsichore abyssinica
Bunaeinae [nec Saturniinae] Eochroini [nec Urotini] Usta wallengrenii
Bunaeinae [nec Saturniinae] Eochroini [nec Urotini] Parusta xanthops
Bunaeinae [nec Saturniinae] Eochroini [nec Bunaeini] Eochroa trimenii
Saturniinae Saturniini Agapema homogena
Saturniinae Saturniini Agapema pelora
Saturniinae Saturniini Saturnia collardi
Saturniinae Saturniini Saturnia atlantica
Saturniinae Saturniini Calosaturnia walterorum
Saturniinae Saturniini Rinaca jonasi
Saturniinae Saturniini Rinaca thibeta
Saturniinae Saturniini Cachosaturnia cachara
Saturniinae Saturniini Perisomena caecigena
Saturniinae Saturniini Neoris shadulla
Saturniinae Saturniini Neoris naessigi
Saturniinae Saturniini Copaxa syntheratoides
Saturniinae Saturniini Copaxa mannana
Saturniinae Saturniini Copaxa joinvillea
Saturniinae Saturniini Neodiphthera papuana
Saturniinae Saturniini Neodiphthera paukstadtorum
Saturniinae Saturniini Pararhodia hiltrautae
Saturniinae Saturniini Syntherata janetta
Saturniinae Saturniini Austrocaligula eucalypti
Saturniinae Saturniini Opodiphthera astrophela
Saturniinae Saturniini Actias chapae chapae
Saturniinae Saturniini Actias isabellae
Saturniinae Saturniini Actias selene
Saturniinae Saturniini Argema mimosae
Saturniinae Saturniini Argema mittrei
Saturniinae Saturniini Antheraea sumatrana
Saturniinae Saturniini Antheraea roylii korintjiana
Saturniinae Saturniini Antheraea hagedorni
Saturniinae Saturniini Antheraea paniki
Saturniinae Saturniini Antheraea castanea
Saturniinae Saturniini Antheraea polyphemus
Saturniinae Saturniini Antheraea godmani
Saturniinae Saturniini Antheraea discata
Saturniinae Saturniini Lemaireia inexpectata
Saturniinae Saturniini Lemaireia schintelmeisteri
Saturniinae Saturniini Cricula trifenestrata javana
Saturniinae Saturniini Cricula vietnama
Saturniinae Saturniini Cricula sumatrensis
Saturniinae Saturniini Cricula fansipanensis
Saturniinae Saturniini [nec Micragonini] Eosia insignis
Saturniinae Saturniini [nec Micragonini] Eosia digennaroi
Saturniinae Saturniini Loepa microocellata
Saturniinae Saturniini Loepa minahassae
Saturniinae Saturniini Loepa oberthuri
Saturniinae Saturniini Ceranchia apollina
Saturniinae Saturniini Antherina suraka
Saturniinae Attacini Callosamia promethea
Saturniinae Attacini Callosamia securifera
Saturniinae Attacini Hyalophora cecropia
Saturniinae Attacini Hyalophora mexicana
Saturniinae Attacini Samia canningi
Saturniinae Attacini Samia tetrica
Saturniinae Attacini Samia peigleri
Saturniinae Attacini Epiphora marginimacula
Saturniinae Attacini Epiphora magdalena
Saturniinae Attacini Epiphora bauhiniae
Saturniinae Attacini Attacus wardi
Saturniinae Attacini Attacus taprobanis
Saturniinae Attacini Attacus caesar
Saturniinae Attacini Coscinocera rothschildi
Saturniinae Attacini Coscinocera anteus
Saturniinae Attacini Archaeoattacus staudingeri
Saturniinae Attacini Archaeoattacus edwardsii
Saturniinae Attacini Rothschildia hesperus
Saturniinae Attacini Rothschildia orizaba
Saturniinae Attacini Rothschildia erycina nigrescens
Saturniinae Attacini Eupackardia calleta
Saturniinae Attacini [nec Saturniini] Rhodinia broschi
Saturniinae Attacini [nec Saturniini] Rhodinia fugax diana
Saturniinae Solini [nec Saturniini] Solus parvifenestratus gabaidanus
Saturniinae Solini [nec Saturniini] Solus kontuma
Saturniinae Solini [nec Saturniini] Solus drepanoides
Salassinae Salassa mesosa
Salassinae Salassa katschinica
Salassinae Salassa inversa
Salassinae Salassa siriae
Salassinae Salassa olivacea
Cercophaninae Janiodini Janiodes ecarcabuco
Cercophaninae Janiodini Janiodes oxapampensis
Cercophaninae Janiodini Janiodes manzanoi
Cercophaninae Cercophanini Cercophana frauenfeldi
Cercophaninae Cercophanini Neocercophana philippii
Cercophaninae Cercophanini Microdulia mirabilis
Oxyteninae Therinia buckleyi
Oxyteninae Therinia sp
Oxyteninae Therinia transversaria
Oxyteninae Oxytenis modoccidentalis
Oxyteninae Oxytenis naemia
Oxyteninae Oxytenis peregrina
Oxyteninae Homoeopteryx pinchcarchensis
Sphingidae Smerinthinae Smerinthini Paonias myops
Sphingidae Smerinthinae Smerinthini Smerinthus ocellata
Sphingidae Smerinthinae Smerinthini Pachysphinx occidentalis
Sphingidae Smerinthinae Smerinthini Laothoe populi
Sphingidae Smerinthinae Coequosa australasiae
Sphingidae Sphinginae Sphingini Agrius convolvuli
Sphingidae Sphinginae Sphingini Acherontia atropos
Sphingidae Langiinae Langia zenzeroides
Sphingidae Macroglossinae Hemarini Hemaris fuciformis
Sphingidae Macroglossinae Macroglossini Daphnis nerii
Bombycidae Epiinae Quentalia mailynae
Endromidae Endromis versicolora
Brahmaeidae Lemonia taraxaci
Brahmaeidae Acanthobrahmaea europaea
Eupterotidae Janinae Stenoglene roseus
Eupterotidae Striphnopteryginae Phiala sp

### Figure S2M. Data subset (50% most saturated loci) IQ-TREE unpartitioned

Best fit model = GTR+F+R10 chosen according to BIC
SH-aLRT /UFBoot

The new classification proposed in this manuscript is used to annotate tips. Current classification is reported between square brackets

Nota : IQ-TREE analyses of data subsets were only performed without partitioning of the data to save computation time (and given that trees were similar when the initial data set was partitioned or not)

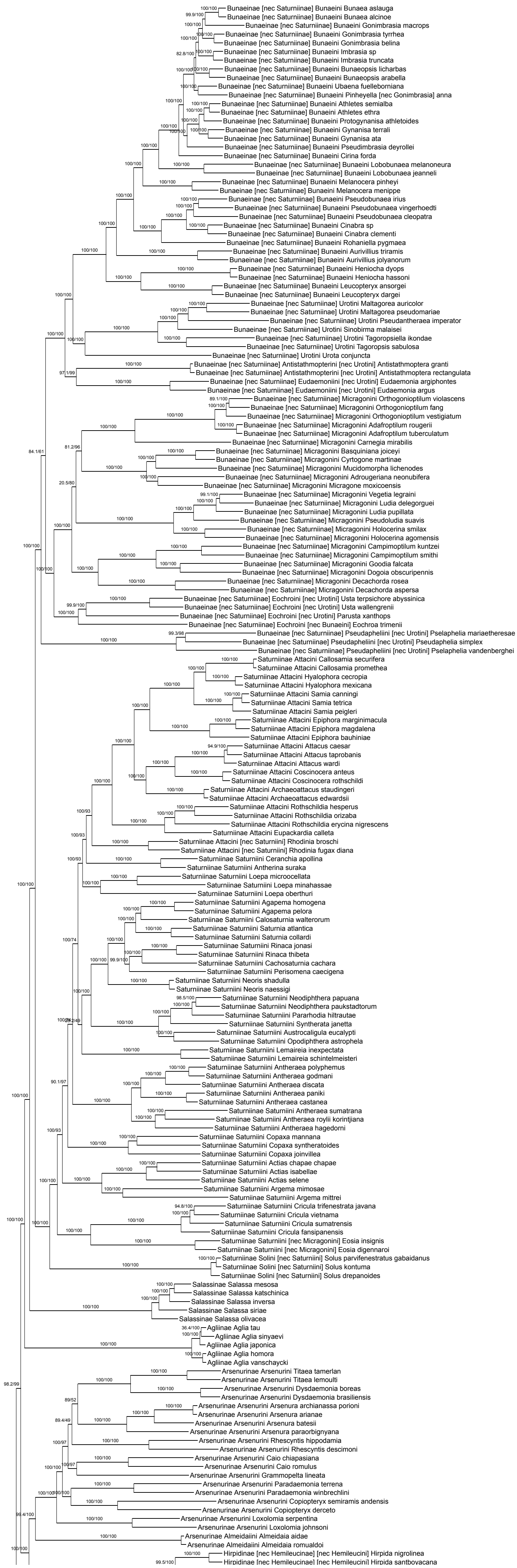

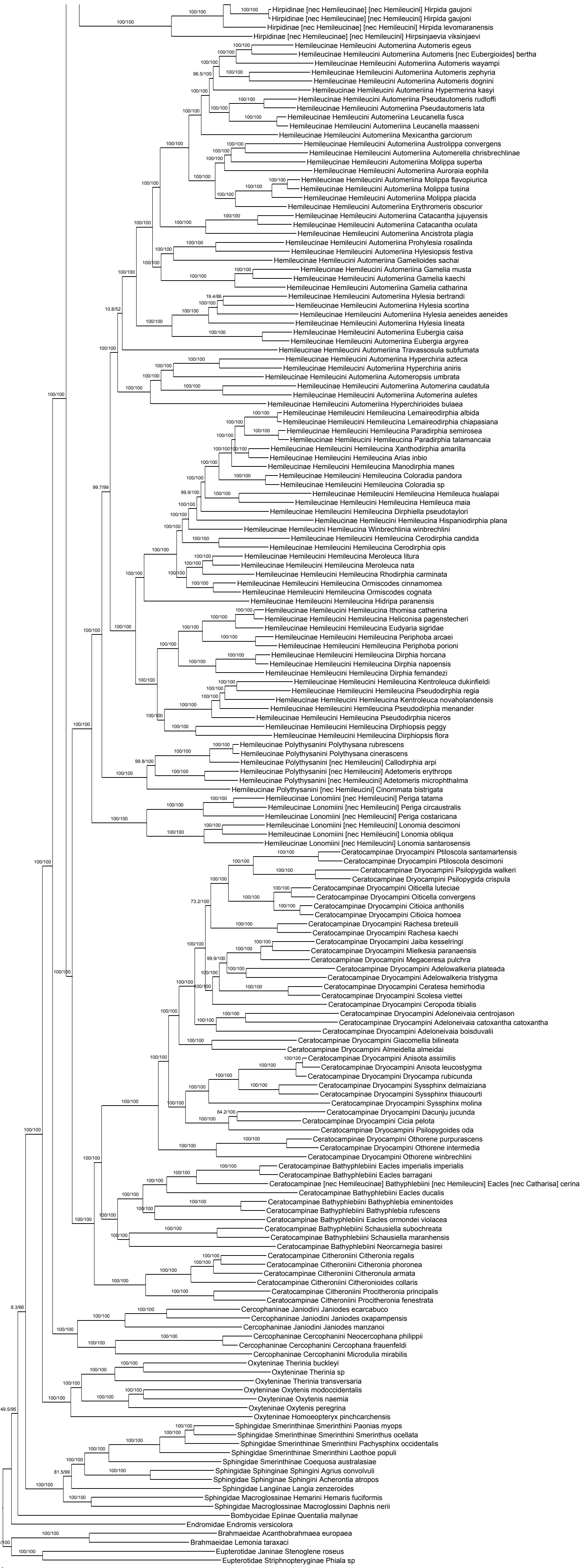

Hirpidinae [nec Hemileucinae] [nec Hemileucini] Hirpida gaujoni
Hirpidinae [nec Hemileucinae] [nec Hemileucini] Hirpida gaujoni
Hirpidinae [nec Hemileucinae] [nec Hemileucini] Hirpida levomaranensis
Hirpidinae [nec Hemileucinae] [nec Hemileucini] Hirpsinjaevia viksinjaevi
Hemileucinae Hemileucini Automeriina Automeris egeus
Hemileucinae Hemileucini Automeriina Automeris [nec Eubergioides] bertha
Hemileucinae Hemileucini Automeriina Automeris wayampi
Hemileucinae Hemileucini Automeriina Automeris zephyria
Hemileucinae Hemileucini Automeriina Automeris dognini
Hemileucinae Hemileucini Automeriina Hypermerina kasyi
Hemileucinae Hemileucini Automeriina Pseudautomeris rudloffi
Hemileucinae Hemileucini Automeriina Pseudautomeris lata
Hemileucinae Hemileucini Automeriina Leucanella fusca
Hemileucinae Hemileucini Automeriina Leucanella maasseni
Hemileucinae Hemileucini Automeriina Mexicantha garciorum
Hemileucinae Hemileucini Automeriina Austrolippa convergens
Hemileucinae Hemileucini Automeriina Automerella chrisbrechlinae
Hemileucinae Hemileucini Automeriina Molippa superba
Hemileucinae Hemileucini Automeriina Auroraia eophila
Hemileucinae Hemileucini Automeriina Molippa flavopiurica
Hemileucinae Hemileucini Automeriina Molippa tusina
Hemileucinae Hemileucini Automeriina Molippa placida
Hemileucinae Hemileucini Automeriina Erythromeris obscurior
Hemileucinae Hemileucini Automeriina Catacantha jujuyensis
Hemileucinae Hemileucini Automeriina Catacantha oculata
Hemileucinae Hemileucini Automeriina Ancistrota plagia
Hemileucinae Hemileucini Automeriina Prohylesia rosalinda
Hemileucinae Hemileucini Automeriina Hylesiopsis festiva
Hemileucinae Hemileucini Automeriina Gamelioides sachai
Hemileucinae Hemileucini Automeriina Gamelia musta
Hemileucinae Hemileucini Automeriina Gamelia kaechi
Hemileucinae Hemileucini Automeriina Gamelia catharina
Hemileucinae Hemileucini Automeriina Hylesia bertrandi
Hemileucinae Hemileucini Automeriina Hylesia scortina
Hemileucinae Hemileucini Automeriina Hylesia aeneides aeneides
Hemileucinae Hemileucini Automeriina Hylesia lineata
Hemileucinae Hemileucini Automeriina Eubergia caisa
Hemileucinae Hemileucini Automeriina Eubergia argyrea
Hemileucinae Hemileucini Automeriina Travassosula subfumata
Hemileucinae Hemileucini Automeriina Hyperchiria azteca
Hemileucinae Hemileucini Automeriina Hyperchiria aniris
Hemileucinae Hemileucini Automeriina Automeropsis umbrata
Hemileucinae Hemileucini Automeriina Automerina caudatula
Hemileucinae Hemileucini Automeriina Automerina auletes
Hemileucinae Hemileucini Automeriina Hyperchirioides bulaea
Hemileucinae Hemileucini Hemileucina Lemaireodirphia albida
Hemileucinae Hemileucini Hemileucina Lemaireodirphia chiapasiana
Hemileucinae Hemileucini Hemileucina Paradirphia semirosea
Hemileucinae Hemileucini Hemileucina Paradirphia talamancaia
Hemileucinae Hemileucini Hemileucina Xanthodirphia amarilla
Hemileucinae Hemileucini Hemileucina Arias inbio
Hemileucinae Hemileucini Hemileucina Manodirphia manes
Hemileucinae Hemileucini Hemileucina Coloradia pandora
Hemileucinae Hemileucini Hemileucina Coloradia sp
Hemileucinae Hemileucini Hemileucina Hemileuca hualapai
Hemileucinae Hemileucini Hemileucina Hemileuca maia
Hemileucinae Hemileucini Hemileucina Dirphiella pseudotaylori
Hemileucinae Hemileucini Hemileucina Hispaniodirphia plana
Hemileucinae Hemileucini Hemileucina Winbrechlinia winbrechlini
Hemileucinae Hemileucini Hemileucina Cerodirphia candida
Hemileucinae Hemileucini Hemileucina Cerodirphia opis
Hemileucinae Hemileucini Hemileucina Meroleuca litura
Hemileucinae Hemileucini Hemileucina Meroleuca nata
Hemileucinae Hemileucini Hemileucina Rhodirphia carminata
Hemileucinae Hemileucini Hemileucina Ormiscodes cinnamomea
Hemileucinae Hemileucini Hemileucina Ormiscodes cognata
Hemileucinae Hemileucini Hemileucina Hidripa paranensis
Hemileucinae Hemileucini Hemileucina Ithomisa catherina
Hemileucinae Hemileucini Hemileucina Heliconisa pagenstecheri
Hemileucinae Hemileucini Hemileucina Eudyaria sigridae
Hemileucinae Hemileucini Hemileucina Periphoba arcaei
Hemileucinae Hemileucini Hemileucina Periphoba porioni
Hemileucinae Hemileucini Hemileucina Dirphia horcana
Hemileucinae Hemileucini Hemileucina Dirphia napoensis
Hemileucinae Hemileucini Hemileucina Dirphia fernandezi
Hemileucinae Hemileucini Hemileucina Kentroleuca dukinfieldi
Hemileucinae Hemileucini Hemileucina Pseudodirphia regia
Hemileucinae Hemileucini Hemileucina Kentroleuca novaholandensis
Hemileucinae Hemileucini Hemileucina Pseudodirphia menander
Hemileucinae Hemileucini Hemileucina Pseudodirphia niceros
Hemileucinae Hemileucini Hemileucina Dirphiopsis peggy
Hemileucinae Hemileucini Hemileucina Dirphiopsis flora
Hemileucinae Polythysanini Polythysana rubrescens
Hemileucinae Polythysanini Polythysana cinerascens
Hemileucinae Polythysanini [nec Hemileucini] Callodirphia arpi
Hemileucinae Polythysanini [nec Hemileucini] Adetomeris erythrops
Hemileucinae Polythysanini [nec Hemileucini] Adetomeris microphthalma
Hemileucinae Polythysanini [nec Hemileucini] Cinommata bistrigata
Hemileucinae Lonomiini [nec Hemileucini] Periga tatama
Hemileucinae Lonomiini [nec Hemileucini] Periga circaustralis
Hemileucinae Lonomiini [nec Hemileucini] Periga costaricana
Hemileucinae Lonomiini [nec Hemileucini] Lonomia descimoni
Hemileucinae Lonomiini [nec Hemileucini] Lonomia obliqua
Hemileucinae Lonomiini [nec Hemileucini] Lonomia santarosensis
Ceratocampinae Dryocampini Ptiloscola santamartensis
Ceratocampinae Dryocampini Ptiloscola descimoni
Ceratocampinae Dryocampini Psilopygida walkeri
Ceratocampinae Dryocampini Psilopygida crispula
Ceratocampinae Dryocampini Oiticella luteciae
Ceratocampinae Dryocampini Oiticella convergens
Ceratocampinae Dryocampini Citioica anthonilis
Ceratocampinae Dryocampini Citioica homoea
Ceratocampinae Dryocampini Rachesa breteuili
Ceratocampinae Dryocampini Rachesa kaechi
Ceratocampinae Dryocampini Jaiba kesselringi
Ceratocampinae Dryocampini Mielkesia paranaensis
Ceratocampinae Dryocampini Megaceresa pulchra
Ceratocampinae Dryocampini Adelowalkeria plateada
Ceratocampinae Dryocampini Adelowalkeria tristygma
Ceratocampinae Dryocampini Ceratesa hemirhodia
Ceratocampinae Dryocampini Scolesa viettei
Ceratocampinae Dryocampini Ceropoda tibialis
Ceratocampinae Dryocampini Adeloneivaia centrojason
Ceratocampinae Dryocampini Adeloneivaia catoxantha catoxantha
Ceratocampinae Dryocampini Adeloneivaia boisduvalii
Ceratocampinae Dryocampini Giacomellia bilineata
Ceratocampinae Dryocampini Almeidella almeidai
Ceratocampinae Dryocampini Anisota assimilis
Ceratocampinae Dryocampini Anisota leucostygma
Ceratocampinae Dryocampini Dryocampa rubicunda
Ceratocampinae Dryocampini Syssphinx delmaiziana
Ceratocampinae Dryocampini Syssphinx thiaucourti
Ceratocampinae Dryocampini Syssphinx molina
Ceratocampinae Dryocampini Dacunju jucunda
Ceratocampinae Dryocampini Cicia pelota
Ceratocampinae Dryocampini Psilopygoides oda
Ceratocampinae Dryocampini Othorene purpurascens
Ceratocampinae Dryocampini Othorene intermedia
Ceratocampinae Dryocampini Othorene winbrechlini
Ceratocampinae Bathyphlebiini Eacles imperialis imperialis
Ceratocampinae Bathyphlebiini Eacles barragani
Ceratocampinae [nec Hemileucinae] Bathyphlebiini [nec Hemileucini] Eacles [nec Catharisa] cerina
Ceratocampinae Bathyphlebiini Eacles ducalis
Ceratocampinae Bathyphlebiini Bathyphlebia eminentoides
Ceratocampinae Bathyphlebiini Bathyphlebia rufescens
Ceratocampinae Bathyphlebiini Eacles ormondei violacea
Ceratocampinae Bathyphlebiini Schausiella subochreata
Ceratocampinae Bathyphlebiini Schausiella maranhensis
Ceratocampinae Bathyphlebiini Neorcarnegia basirei
Ceratocampinae Citheroniini Citheronia regalis
Ceratocampinae Citheroniini Citheronia phoronea
Ceratocampinae Citheroniini Citheronula armata
Ceratocampinae Citheroniini Citheronioides collaris
Ceratocampinae Citheroniini Procitheronia principalis
Ceratocampinae Citheroniini Procitheronia fenestrata
Cercophaninae Janiodini Janiodes ecarcabuco
Cercophaninae Janiodini Janiodes oxapampensis
Cercophaninae Janiodini Janiodes manzanoi
Cercophaninae Cercophanini Neocercophana philippii
Cercophaninae Cercophanini Cercophana frauenfeldi
Cercophaninae Cercophanini Microdulia mirabilis
Oxyteninae Therinia buckleyi
Oxyteninae Therinia sp
Oxyteninae Therinia transversaria
Oxyteninae Oxytenis modoccidentalis
Oxyteninae Oxytenis naemia
Oxyteninae Oxytenis peregrina
Oxyteninae Homoeopteryx pinchcarchensis
Sphingidae Smerinthinae Smerinthini Paonias myops
Sphingidae Smerinthinae Smerinthini Smerinthus ocellata
Sphingidae Smerinthinae Smerinthini Pachysphinx occidentalis
Sphingidae Smerinthinae Smerinthini Laothoe populi
Sphingidae Smerinthinae Coequosa australasiae
Sphingidae Sphinginae Sphingini Agrius convolvuli
Sphingidae Sphinginae Sphingini Acherontia atropos
Sphingidae Langiinae Langia zenzeroides
Sphingidae Macroglossinae Hemarini Hemaris fuciformis
Sphingidae Macroglossinae Macroglossini Daphnis nerii
Bombycidae Epiinae Quentalia mailynae
Endromidae Endromis versicolora
Brahmaeidae Acanthobrahmaea europaea
Brahmaeidae Lemonia taraxaci
Eupterotidae Janinae Stenoglene roseus
Eupterotidae Striphnopteryginae Phiala sp
0.0
0.06

### Figure S2N. Data subset (50% most saturated loci) ASTRAL50

Species tree obtained when nodes with UFBoot<50 are collapsed in each gene tree

Final normalized quartet score : 0.82
Local posterior probabilities at nodes

The new classification proposed in this manuscript is used to annotate tips. Current classification is reported between square brackets

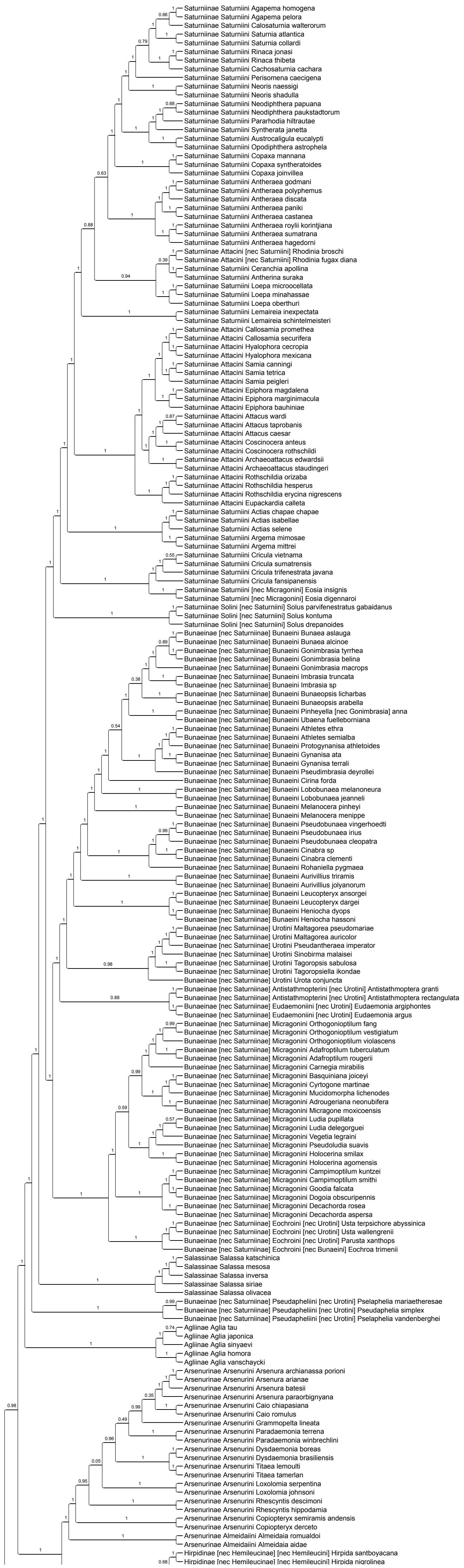

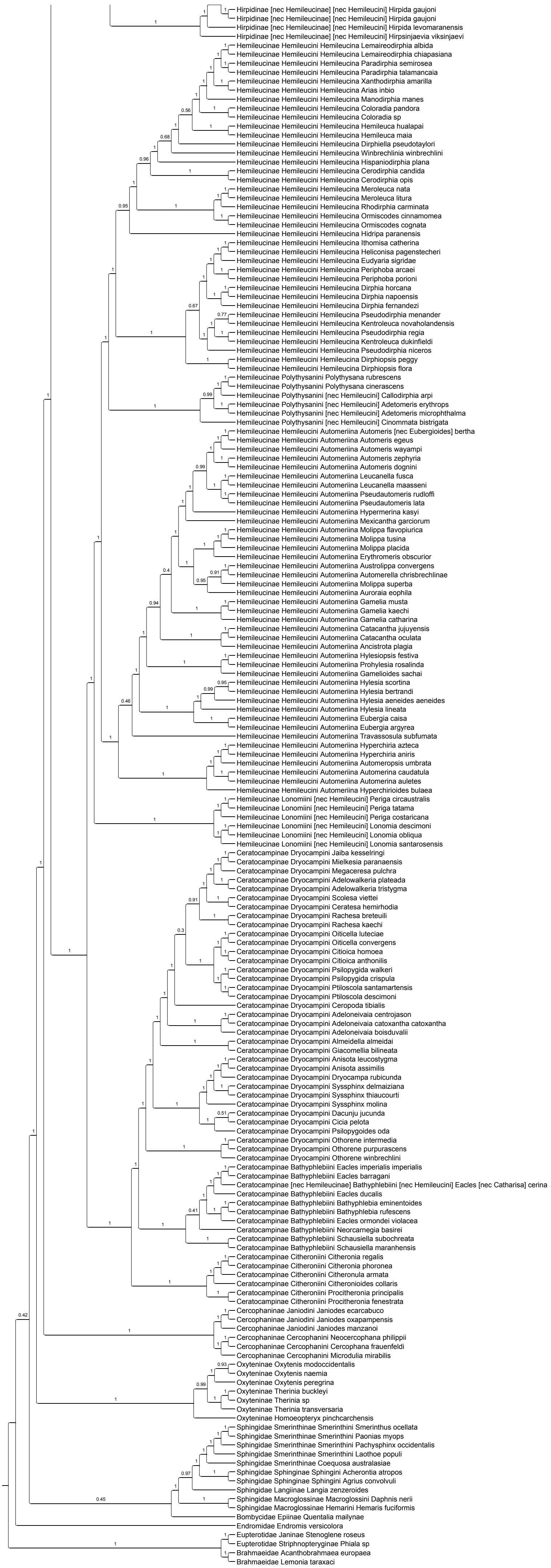
Hirpidinae [nec Hemileucinae] [nec Hemileucini] Hirpida gaujoni
Hirpidinae [nec Hemileucinae] [nec Hemileucini] Hirpida gaujoni
Hirpidinae [nec Hemileucinae] [nec Hemileucini] Hirpida levomaranensis
Hirpidinae [nec Hemileucinae] [nec Hemileucini] Hirpsinjaevia viksinjaevi
Hemileucinae Hemileucini Hemileucina Lemaireodirphia albida
Hemileucinae Hemileucini Hemileucina Lemaireodirphia chiapasiana
Hemileucinae Hemileucini Hemileucina Paradirphia semirosea
Hemileucinae Hemileucini Hemileucina Paradirphia talamancaia
Hemileucinae Hemileucini Hemileucina Xanthodirphia amarilla
Hemileucinae Hemileucini Hemileucina Arias inbio
Hemileucinae Hemileucini Hemileucina Manodirphia manes
Hemileucinae Hemileucini Hemileucina Coloradia pandora
Hemileucinae Hemileucini Hemileucina Coloradia sp
Hemileucinae Hemileucini Hemileucina Hemileuca hualapai
Hemileucinae Hemileucini Hemileucina Hemileuca maia
Hemileucinae Hemileucini Hemileucina Dirphiella pseudotaylori
Hemileucinae Hemileucini Hemileucina Winbrechlinia winbrechlini
Hemileucinae Hemileucini Hemileucina Hispaniodirphia plana
Hemileucinae Hemileucini Hemileucina Cerodirphia candida
Hemileucinae Hemileucini Hemileucina Cerodirphia opis
Hemileucinae Hemileucini Hemileucina Meroleuca nata
Hemileucinae Hemileucini Hemileucina Meroleuca litura
Hemileucinae Hemileucini Hemileucina Rhodirphia carminata
Hemileucinae Hemileucini Hemileucina Ormiscodes cinnamomea
Hemileucinae Hemileucini Hemileucina Ormiscodes cognata
Hemileucinae Hemileucini Hemileucina Hidripa paranensis
Hemileucinae Hemileucini Hemileucina Ithomisa catherina
Hemileucinae Hemileucini Hemileucina Heliconisa pagenstecheri
Hemileucinae Hemileucini Hemileucina Eudyaria sigridae
Hemileucinae Hemileucini Hemileucina Periphoba arcaei
Hemileucinae Hemileucini Hemileucina Periphoba porioni
Hemileucinae Hemileucini Hemileucina Dirphia horcana
Hemileucinae Hemileucini Hemileucina Dirphia napoensis
Hemileucinae Hemileucini Hemileucina Dirphia fernandezi
Hemileucinae Hemileucini Hemileucina Pseudodirphia menander
Hemileucinae Hemileucini Hemileucina Kentroleuca novaholandensis
Hemileucinae Hemileucini Hemileucina Pseudodirphia regia
Hemileucinae Hemileucini Hemileucina Kentroleuca dukinfieldi
Hemileucinae Hemileucini Hemileucina Pseudodirphia niceros
Hemileucinae Hemileucini Hemileucina Dirphiopsis peggy
Hemileucinae Hemileucini Hemileucina Dirphiopsis flora
Hemileucinae Polythysanini Polythysana rubrescens
Hemileucinae Polythysanini Polythysana cinerascens
Hemileucinae Polythysanini [nec Hemileucini] Callodirphia arpi
Hemileucinae Polythysanini [nec Hemileucini] Adetomeris erythrops
Hemileucinae Polythysanini [nec Hemileucini] Adetomeris microphthalma
Hemileucinae Polythysanini [nec Hemileucini] Cinommata bistrigata
Hemileucinae Hemileucini Automeriina Automeris [nec Eubergioides] bertha
Hemileucinae Hemileucini Automeriina Automeris egeus
Hemileucinae Hemileucini Automeriina Automeris wayampi
Hemileucinae Hemileucini Automeriina Automeris zephyria
Hemileucinae Hemileucini Automeriina Automeris dognini
Hemileucinae Hemileucini Automeriina Leucanella fusca
Hemileucinae Hemileucini Automeriina Leucanella maasseni
Hemileucinae Hemileucini Automeriina Pseudautomeris rudloffi
Hemileucinae Hemileucini Automeriina Pseudautomeris lata
Hemileucinae Hemileucini Automeriina Hypermerina kasyi
Hemileucinae Hemileucini Automeriina Mexicantha garciorum
Hemileucinae Hemileucini Automeriina Molippa flavopiurica
Hemileucinae Hemileucini Automeriina Molippa tusina
Hemileucinae Hemileucini Automeriina Molippa placida
Hemileucinae Hemileucini Automeriina Erythromeris obscurior
Hemileucinae Hemileucini Automeriina Austrolippa convergens
Hemileucinae Hemileucini Automeriina Automerella chrisbrechlinae
Hemileucinae Hemileucini Automeriina Molippa superba
Hemileucinae Hemileucini Automeriina Auroraia eophila
Hemileucinae Hemileucini Automeriina Gamelia musta
Hemileucinae Hemileucini Automeriina Gamelia kaechi
Hemileucinae Hemileucini Automeriina Gamelia catharina
Hemileucinae Hemileucini Automeriina Catacantha jujuyensis
Hemileucinae Hemileucini Automeriina Catacantha oculata
Hemileucinae Hemileucini Automeriina Ancistrota plagia
Hemileucinae Hemileucini Automeriina Hylesiopsis festiva
Hemileucinae Hemileucini Automeriina Prohylesia rosalinda
Hemileucinae Hemileucini Automeriina Gamelioides sachai
Hemileucinae Hemileucini Automeriina Hylesia scortina
Hemileucinae Hemileucini Automeriina Hylesia bertrandi
Hemileucinae Hemileucini Automeriina Hylesia aeneides aeneides
Hemileucinae Hemileucini Automeriina Hylesia lineata
Hemileucinae Hemileucini Automeriina Eubergia caisa
Hemileucinae Hemileucini Automeriina Eubergia argyrea
Hemileucinae Hemileucini Automeriina Travassosula subfumata
Hemileucinae Hemileucini Automeriina Hyperchiria azteca
Hemileucinae Hemileucini Automeriina Hyperchiria aniris
Hemileucinae Hemileucini Automeriina Automeropsis umbrata
Hemileucinae Hemileucini Automeriina Automerina caudatula
Hemileucinae Hemileucini Automeriina Automerina auletes
Hemileucinae Hemileucini Automeriina Hyperchirioides bulaea
Hemileucinae Lonomiini [nec Hemileucini] Periga circaustralis
Hemileucinae Lonomiini [nec Hemileucini] Periga tatama
Hemileucinae Lonomiini [nec Hemileucini] Periga costaricana
Hemileucinae Lonomiini [nec Hemileucini] Lonomia descimoni
Hemileucinae Lonomiini [nec Hemileucini] Lonomia obliqua
Hemileucinae Lonomiini [nec Hemileucini] Lonomia santarosensis
Ceratocampinae Dryocampini Jaiba kesselringi
Ceratocampinae Dryocampini Mielkesia paranaensis
Ceratocampinae Dryocampini Megaceresa pulchra
Ceratocampinae Dryocampini Adelowalkeria plateada
Ceratocampinae Dryocampini Adelowalkeria tristygma
Ceratocampinae Dryocampini Scolesa viettei
Ceratocampinae Dryocampini Ceratesa hemirhodia
Ceratocampinae Dryocampini Rachesa breteuili
Ceratocampinae Dryocampini Rachesa kaechi
Ceratocampinae Dryocampini Oiticella luteciae
Ceratocampinae Dryocampini Oiticella convergens
Ceratocampinae Dryocampini Citioica homoea
Ceratocampinae Dryocampini Citioica anthonilis
Ceratocampinae Dryocampini Psilopygida walkeri
Ceratocampinae Dryocampini Psilopygida crispula
Ceratocampinae Dryocampini Ptiloscola santamartensis
Ceratocampinae Dryocampini Ptiloscola descimoni
Ceratocampinae Dryocampini Ceropoda tibialis
Ceratocampinae Dryocampini Adeloneivaia centrojason
Ceratocampinae Dryocampini Adeloneivaia catoxantha catoxantha
Ceratocampinae Dryocampini Adeloneivaia boisduvalii
Ceratocampinae Dryocampini Almeidella almeidai
Ceratocampinae Dryocampini Giacomellia bilineata
Ceratocampinae Dryocampini Anisota leucostygma
Ceratocampinae Dryocampini Anisota assimilis
Ceratocampinae Dryocampini Dryocampa rubicunda
Ceratocampinae Dryocampini Syssphinx delmaiziana
Ceratocampinae Dryocampini Syssphinx thiaucourti
Ceratocampinae Dryocampini Syssphinx molina
Ceratocampinae Dryocampini Dacunju jucunda
Ceratocampinae Dryocampini Cicia pelota
Ceratocampinae Dryocampini Psilopygoides oda
Ceratocampinae Dryocampini Othorene intermedia
Ceratocampinae Dryocampini Othorene purpurascens
Ceratocampinae Dryocampini Othorene winbrechlini
Ceratocampinae Bathyphlebiini Eacles imperialis imperialis
Ceratocampinae Bathyphlebiini Eacles barragani
Ceratocampinae [nec Hemileucinae] Bathyphlebiini [nec Hemileucini] Eacles [nec Catharisa] cerina
Ceratocampinae Bathyphlebiini Eacles ducalis
Ceratocampinae Bathyphlebiini Bathyphlebia eminentoides
Ceratocampinae Bathyphlebiini Bathyphlebia rufescens
Ceratocampinae Bathyphlebiini Eacles ormondei violacea
Ceratocampinae Bathyphlebiini Neorcarnegia basirei
Ceratocampinae Bathyphlebiini Schausiella subochreata
Ceratocampinae Bathyphlebiini Schausiella maranhensis
Ceratocampinae Citheroniini Citheronia regalis
Ceratocampinae Citheroniini Citheronia phoronea
Ceratocampinae Citheroniini Citheronula armata
Ceratocampinae Citheroniini Citheronioides collaris
Ceratocampinae Citheroniini Procitheronia principalis
Ceratocampinae Citheroniini Procitheronia fenestrata
Cercophaninae Janiodini Janiodes ecarcabuco
Cercophaninae Janiodini Janiodes oxapampensis
Cercophaninae Janiodini Janiodes manzanoi
Cercophaninae Cercophanini Neocercophana philippii
Cercophaninae Cercophanini Cercophana frauenfeldi
Cercophaninae Cercophanini Microdulia mirabilis
Oxyteninae Oxytenis modoccidentalis
Oxyteninae Oxytenis naemia
Oxyteninae Oxytenis peregrina
Oxyteninae Therinia buckleyi
Oxyteninae Therinia sp
Oxyteninae Therinia transversaria
Oxyteninae Homoeopteryx pinchcarchensis
Sphingidae Smerinthinae Smerinthini Smerinthus ocellata
Sphingidae Smerinthinae Smerinthini Paonias myops
Sphingidae Smerinthinae Smerinthini Pachysphinx occidentalis
Sphingidae Smerinthinae Smerinthini Laothoe populi
Sphingidae Smerinthinae Coequosa australasiae
Sphingidae Sphinginae Sphingini Acherontia atropos
Sphingidae Sphinginae Sphingini Agrius convolvuli
Sphingidae Langiinae Langia zenzeroides
Sphingidae Macroglossinae Macroglossini Daphnis nerii
Sphingidae Macroglossinae Hemarini Hemaris fuciformis
Bombycidae Epiinae Quentalia mailynae
Endromidae Endromis versicolora
Eupterotidae Janinae Stenoglene roseus
Eupterotidae Striphnopteryginae Phiala sp
Brahmaeidae Acanthobrahmaea europaea
Brahmaeidae Lemonia taraxaci

#### Figure S2O. Data subset (50% most saturated loci) ASTRAL70

Species tree obtained when nodes with UFBoot<70 are collapsed in each gene tree

Final normalized quartet score : 0.86
Local posterior probabilities at nodes

The new classification proposed in this manuscript is used to annotate tips. Current classification is reported between square brackets

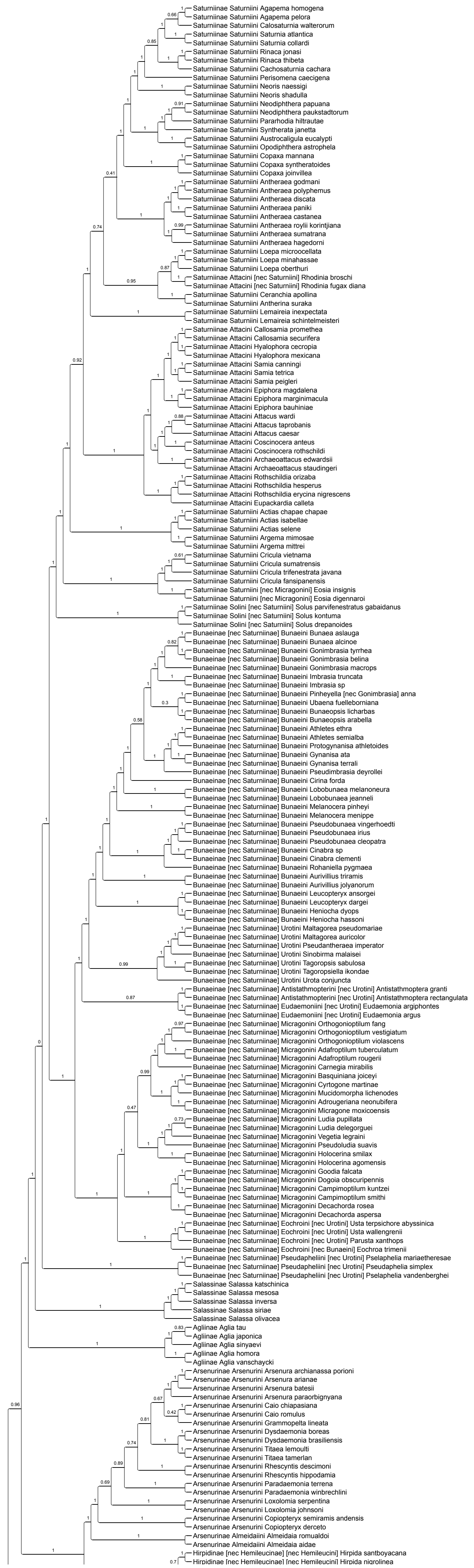

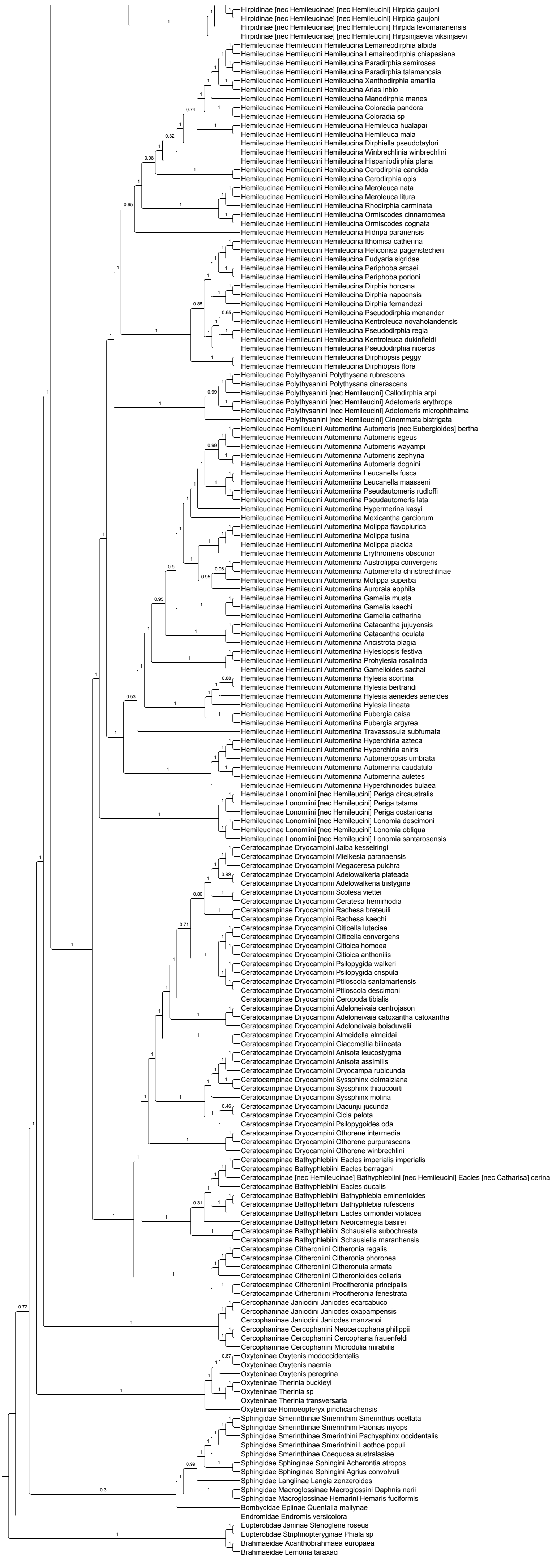
Hirpidinae [nec Hemileucinae] [nec Hemileucini] Hirpida gaujoni
Hirpidinae [nec Hemileucinae] [nec Hemileucini] Hirpida gaujoni
Hirpidinae [nec Hemileucinae] [nec Hemileucini] Hirpida levomaranensis
Hirpidinae [nec Hemileucinae] [nec Hemileucini] Hirpsinjaevia viksinjaevi
Hemileucinae Hemileucini Hemileucina Lemaireodirphia albida
Hemileucinae Hemileucini Hemileucina Lemaireodirphia chiapasiana
Hemileucinae Hemileucini Hemileucina Paradirphia semirosea
Hemileucinae Hemileucini Hemileucina Paradirphia talamancaia
Hemileucinae Hemileucini Hemileucina Xanthodirphia amarilla
Hemileucinae Hemileucini Hemileucina Arias inbio
Hemileucinae Hemileucini Hemileucina Manodirphia manes
Hemileucinae Hemileucini Hemileucina Coloradia pandora
Hemileucinae Hemileucini Hemileucina Coloradia sp
Hemileucinae Hemileucini Hemileucina Hemileuca hualapai
Hemileucinae Hemileucini Hemileucina Hemileuca maia
Hemileucinae Hemileucini Hemileucina Dirphiella pseudotaylori
Hemileucinae Hemileucini Hemileucina Winbrechlinia winbrechlini
Hemileucinae Hemileucini Hemileucina Hispaniodirphia plana
Hemileucinae Hemileucini Hemileucina Cerodirphia candida
Hemileucinae Hemileucini Hemileucina Cerodirphia opis
Hemileucinae Hemileucini Hemileucina Meroleuca nata
Hemileucinae Hemileucini Hemileucina Meroleuca litura
Hemileucinae Hemileucini Hemileucina Rhodirphia carminata
Hemileucinae Hemileucini Hemileucina Ormiscodes cinnamomea
Hemileucinae Hemileucini Hemileucina Ormiscodes cognata
Hemileucinae Hemileucini Hemileucina Hidripa paranensis
Hemileucinae Hemileucini Hemileucina Ithomisa catherina
Hemileucinae Hemileucini Hemileucina Heliconisa pagenstecheri
Hemileucinae Hemileucini Hemileucina Eudyaria sigridae
Hemileucinae Hemileucini Hemileucina Periphoba arcaei
Hemileucinae Hemileucini Hemileucina Periphoba porioni
Hemileucinae Hemileucini Hemileucina Dirphia horcana
Hemileucinae Hemileucini Hemileucina Dirphia napoensis
Hemileucinae Hemileucini Hemileucina Dirphia fernandezi
Hemileucinae Hemileucini Hemileucina Pseudodirphia menander
Hemileucinae Hemileucini Hemileucina Kentroleuca novaholandensis
Hemileucinae Hemileucini Hemileucina Pseudodirphia regia
Hemileucinae Hemileucini Hemileucina Kentroleuca dukinfieldi
Hemileucinae Hemileucini Hemileucina Pseudodirphia niceros
Hemileucinae Hemileucini Hemileucina Dirphiopsis peggy
Hemileucinae Hemileucini Hemileucina Dirphiopsis flora
Hemileucinae Polythysanini Polythysana rubrescens
Hemileucinae Polythysanini Polythysana cinerascens
Hemileucinae Polythysanini [nec Hemileucini] Callodirphia arpi
Hemileucinae Polythysanini [nec Hemileucini] Adetomeris erythrops
Hemileucinae Polythysanini [nec Hemileucini] Adetomeris microphthalma
Hemileucinae Polythysanini [nec Hemileucini] Cinommata bistrigata
Hemileucinae Hemileucini Automeriina Automeris [nec Eubergioides] bertha
Hemileucinae Hemileucini Automeriina Automeris egeus
Hemileucinae Hemileucini Automeriina Automeris wayampi
Hemileucinae Hemileucini Automeriina Automeris zephyria
Hemileucinae Hemileucini Automeriina Automeris dognini
Hemileucinae Hemileucini Automeriina Leucanella fusca
Hemileucinae Hemileucini Automeriina Leucanella maasseni
Hemileucinae Hemileucini Automeriina Pseudautomeris rudloffi
Hemileucinae Hemileucini Automeriina Pseudautomeris lata
Hemileucinae Hemileucini Automeriina Hypermerina kasyi
Hemileucinae Hemileucini Automeriina Mexicantha garciorum
Hemileucinae Hemileucini Automeriina Molippa flavopiurica
Hemileucinae Hemileucini Automeriina Molippa tusina
Hemileucinae Hemileucini Automeriina Molippa placida
Hemileucinae Hemileucini Automeriina Erythromeris obscurior
Hemileucinae Hemileucini Automeriina Austrolippa convergens
Hemileucinae Hemileucini Automeriina Automerella chrisbrechlinae
Hemileucinae Hemileucini Automeriina Molippa superba
Hemileucinae Hemileucini Automeriina Auroraia eophila
Hemileucinae Hemileucini Automeriina Gamelia musta
Hemileucinae Hemileucini Automeriina Gamelia kaechi
Hemileucinae Hemileucini Automeriina Gamelia catharina
Hemileucinae Hemileucini Automeriina Catacantha jujuyensis
Hemileucinae Hemileucini Automeriina Catacantha oculata
Hemileucinae Hemileucini Automeriina Ancistrota plagia
Hemileucinae Hemileucini Automeriina Hylesiopsis festiva
Hemileucinae Hemileucini Automeriina Prohylesia rosalinda
Hemileucinae Hemileucini Automeriina Gamelioides sachai
Hemileucinae Hemileucini Automeriina Hylesia scortina
Hemileucinae Hemileucini Automeriina Hylesia bertrandi
Hemileucinae Hemileucini Automeriina Hylesia aeneides aeneides
Hemileucinae Hemileucini Automeriina Hylesia lineata
Hemileucinae Hemileucini Automeriina Eubergia caisa
Hemileucinae Hemileucini Automeriina Eubergia argyrea
Hemileucinae Hemileucini Automeriina Travassosula subfumata
Hemileucinae Hemileucini Automeriina Hyperchiria azteca
Hemileucinae Hemileucini Automeriina Hyperchiria aniris
Hemileucinae Hemileucini Automeriina Automeropsis umbrata
Hemileucinae Hemileucini Automeriina Automerina caudatula
Hemileucinae Hemileucini Automeriina Automerina auletes
Hemileucinae Hemileucini Automeriina Hyperchirioides bulaea
Hemileucinae Lonomiini [nec Hemileucini] Periga circaustralis
Hemileucinae Lonomiini [nec Hemileucini] Periga tatama
Hemileucinae Lonomiini [nec Hemileucini] Periga costaricana
Hemileucinae Lonomiini [nec Hemileucini] Lonomia descimoni
Hemileucinae Lonomiini [nec Hemileucini] Lonomia obliqua
Hemileucinae Lonomiini [nec Hemileucini] Lonomia santarosensis
Ceratocampinae Dryocampini Jaiba kesselringi
Ceratocampinae Dryocampini Mielkesia paranaensis
Ceratocampinae Dryocampini Megaceresa pulchra
Ceratocampinae Dryocampini Adelowalkeria plateada
Ceratocampinae Dryocampini Adelowalkeria tristygma
Ceratocampinae Dryocampini Scolesa viettei
Ceratocampinae Dryocampini Ceratesa hemirhodia
Ceratocampinae Dryocampini Rachesa breteuili
Ceratocampinae Dryocampini Rachesa kaechi
Ceratocampinae Dryocampini Oiticella luteciae
Ceratocampinae Dryocampini Oiticella convergens
Ceratocampinae Dryocampini Citioica homoea
Ceratocampinae Dryocampini Citioica anthonilis
Ceratocampinae Dryocampini Psilopygida walkeri
Ceratocampinae Dryocampini Psilopygida crispula
Ceratocampinae Dryocampini Ptiloscola santamartensis
Ceratocampinae Dryocampini Ptiloscola descimoni
Ceratocampinae Dryocampini Ceropoda tibialis
Ceratocampinae Dryocampini Adeloneivaia centrojason
Ceratocampinae Dryocampini Adeloneivaia catoxantha catoxantha
Ceratocampinae Dryocampini Adeloneivaia boisduvalii
Ceratocampinae Dryocampini Almeidella almeidai
Ceratocampinae Dryocampini Giacomellia bilineata
Ceratocampinae Dryocampini Anisota leucostygma
Ceratocampinae Dryocampini Anisota assimilis
Ceratocampinae Dryocampini Dryocampa rubicunda
Ceratocampinae Dryocampini Syssphinx delmaiziana
Ceratocampinae Dryocampini Syssphinx thiaucourti
Ceratocampinae Dryocampini Syssphinx molina
Ceratocampinae Dryocampini Dacunju jucunda
Ceratocampinae Dryocampini Cicia pelota
Ceratocampinae Dryocampini Psilopygoides oda
Ceratocampinae Dryocampini Othorene intermedia
Ceratocampinae Dryocampini Othorene purpurascens
Ceratocampinae Dryocampini Othorene winbrechlini
Ceratocampinae Bathyphlebiini Eacles imperialis imperialis
Ceratocampinae Bathyphlebiini Eacles barragani
Ceratocampinae [nec Hemileucinae] Bathyphlebiini [nec Hemileucini] Eacles [nec Catharisa] cerina
Ceratocampinae Bathyphlebiini Eacles ducalis
Ceratocampinae Bathyphlebiini Bathyphlebia eminentoides
Ceratocampinae Bathyphlebiini Bathyphlebia rufescens
Ceratocampinae Bathyphlebiini Eacles ormondei violacea
Ceratocampinae Bathyphlebiini Neorcarnegia basirei
Ceratocampinae Bathyphlebiini Schausiella subochreata
Ceratocampinae Bathyphlebiini Schausiella maranhensis
Ceratocampinae Citheroniini Citheronia regalis
Ceratocampinae Citheroniini Citheronia phoronea
Ceratocampinae Citheroniini Citheronula armata
Ceratocampinae Citheroniini Citheronioides collaris
Ceratocampinae Citheroniini Procitheronia principalis
Ceratocampinae Citheroniini Procitheronia fenestrata
Cercophaninae Janiodini Janiodes ecarcabuco
Cercophaninae Janiodini Janiodes oxapampensis
Cercophaninae Janiodini Janiodes manzanoi
Cercophaninae Cercophanini Neocercophana philippii
Cercophaninae Cercophanini Cercophana frauenfeldi
Cercophaninae Cercophanini Microdulia mirabilis
Oxyteninae Oxytenis modocidentalis
Oxyteninae Oxytenis naemia
Oxyteninae Oxytenis peregrina
Oxyteninae Therinia buckleyi
Oxyteninae Therinia sp
Oxyteninae Therinia transversaria
Oxyteninae Homoeopteryx pinchcarchensis
Sphingidae Smerinthinae Smerinthini Smerinthus ocellata
Sphingidae Smerinthinae Smerinthini Paonias myops
Sphingidae Smerinthinae Smerinthini Pachysphinx occidentalis
Sphingidae Smerinthinae Smerinthini Laothoe populi
Sphingidae Smerinthinae Coequosa australasiae
Sphingidae Sphinginae Sphingini Acherontia atropos
Sphingidae Sphinginae Sphingini Agrius convolvuli
Sphingidae Langiinae Langia zenzeroides
Sphingidae Macroglossinae Macroglossini Daphnis nerii
Sphingidae Macroglossinae Hemarini Hemaris fuciformis
Bombycidae Epiinae Quentalia mailynae
Endromidae Endromis versicolora
Eupterotidae Janinae Stenoglene roseus
Eupterotidae Striphnopteryginae Phiala sp
Brahmaeidae Acanthobrahmaea europaea
Brahmaeidae Lemonia taraxaci

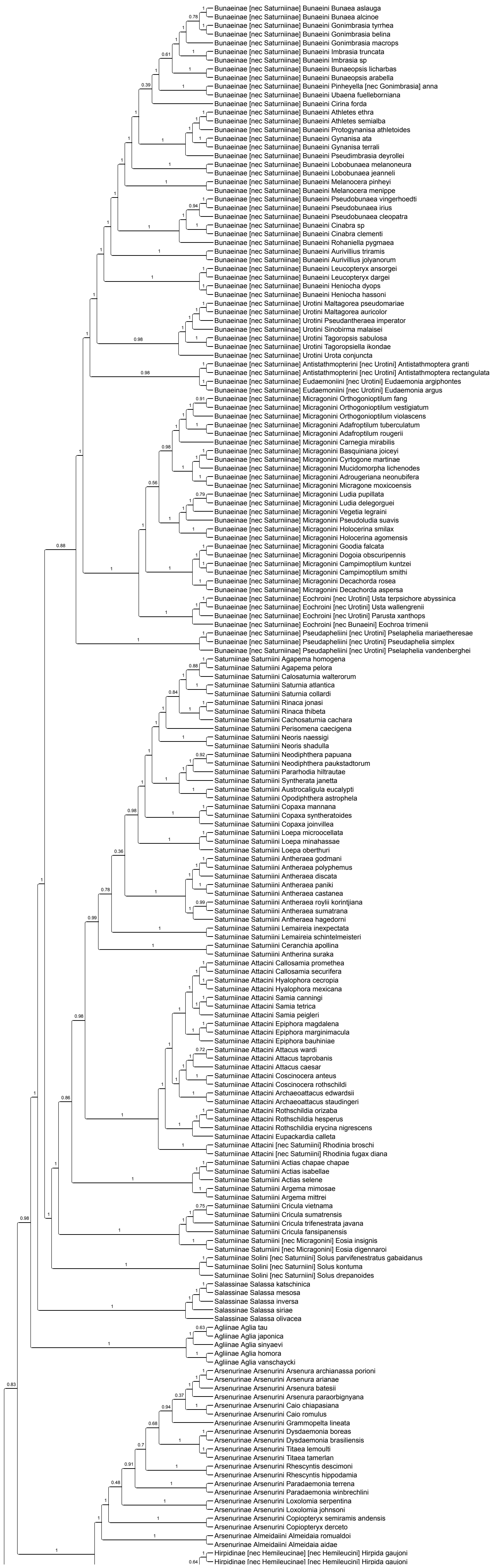


### Figure S2P. Data subset (50% most saturated loci) ASTRAL90

Species tree obtained when nodes with UFBoot<90 are collapsed in each gene tree

Final normalized quartet score : 0.94
Local posterior probabilities at nodes

The new classification proposed in this manuscript is used to annotate tips. Current classification is reported between square brackets

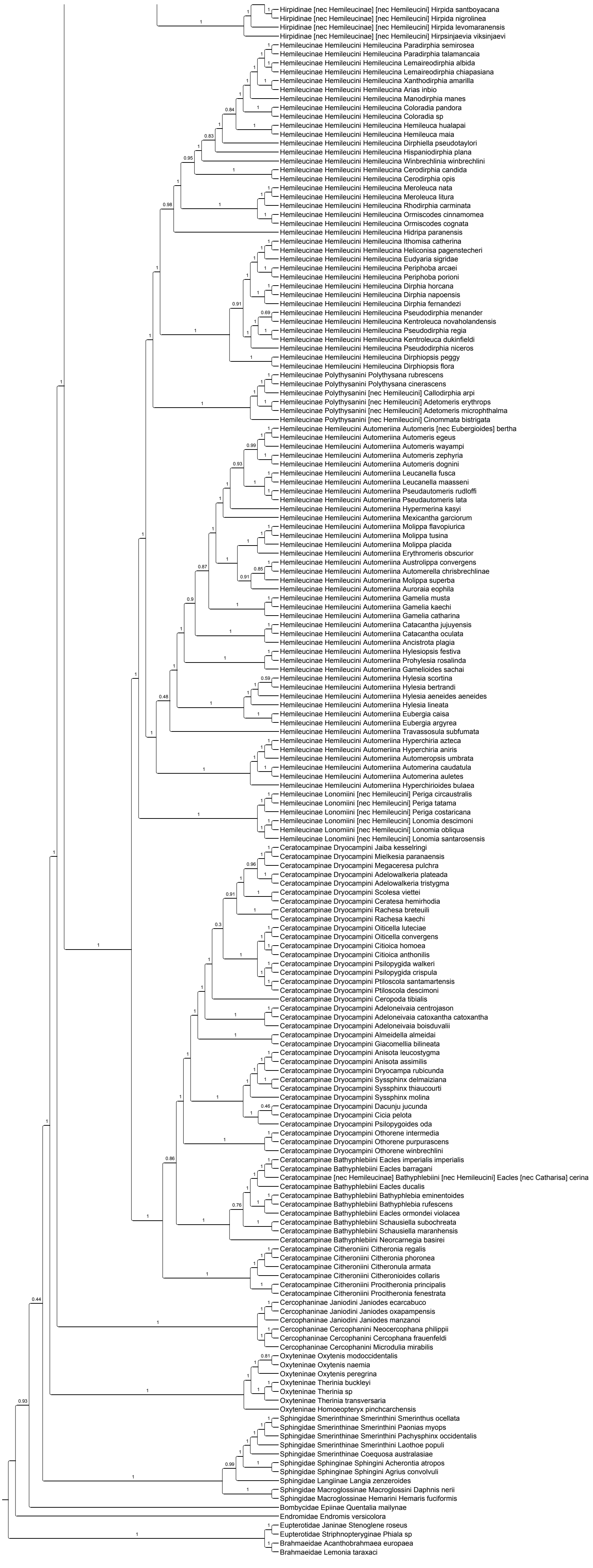
Hirpidinae [nec Hemileucinae] [nec Hemileucini] Hirpida santboyacana
Hirpidinae [nec Hemileucinae] [nec Hemileucini] Hirpida nigrolinea
Hirpidinae [nec Hemileucinae] [nec Hemileucini] Hirpida levomaranensis
Hirpidinae [nec Hemileucinae] [nec Hemileucini] Hirpsinjaevia viksinjaevi
Hemileucinae Hemileucini Hemileucina Paradirphia semirosea
Hemileucinae Hemileucini Hemileucina Paradirphia talamancaia
Hemileucinae Hemileucini Hemileucina Lemaireodirphia albida
Hemileucinae Hemileucini Hemileucina Lemaireodirphia chiapasiana
Hemileucinae Hemileucini Hemileucina Xanthodirphia amarilla
Hemileucinae Hemileucini Hemileucina Arias inbio
Hemileucinae Hemileucini Hemileucina Manodirphia manes
Hemileucinae Hemileucini Hemileucina Coloradia pandora
Hemileucinae Hemileucini Hemileucina Coloradia sp
Hemileucinae Hemileucini Hemileucina Hemileuca hualapai
Hemileucinae Hemileucini Hemileucina Hemileuca maia
Hemileucinae Hemileucini Hemileucina Dirphiella pseudotaylori
Hemileucinae Hemileucini Hemileucina Hispaniodirphia plana
Hemileucinae Hemileucini Hemileucina Winbrechlinia winbrechlini
Hemileucinae Hemileucini Hemileucina Cerodirphia candida
Hemileucinae Hemileucini Hemileucina Cerodirphia opis
Hemileucinae Hemileucini Hemileucina Meroleuca nata
Hemileucinae Hemileucini Hemileucina Meroleuca litura
Hemileucinae Hemileucini Hemileucina Rhodirphia carminata
Hemileucinae Hemileucini Hemileucina Ormiscodes cinnamomea
Hemileucinae Hemileucini Hemileucina Ormiscodes cognata
Hemileucinae Hemileucini Hemileucina Hidripa paranensis
Hemileucinae Hemileucini Hemileucina Ithomisa catherina
Hemileucinae Hemileucini Hemileucina Heliconisa pagenstecheri
Hemileucinae Hemileucini Hemileucina Eudyaria sigridae
Hemileucinae Hemileucini Hemileucina Periphoba arcaei
Hemileucinae Hemileucini Hemileucina Periphoba porioni
Hemileucinae Hemileucini Hemileucina Dirphia horcana
Hemileucinae Hemileucini Hemileucina Dirphia napoensis
Hemileucinae Hemileucini Hemileucina Dirphia fernandezi
Hemileucinae Hemileucini Hemileucina Pseudodirphia menander
Hemileucinae Hemileucini Hemileucina Kentroleuca novaholandensis
Hemileucinae Hemileucini Hemileucina Pseudodirphia regia
Hemileucinae Hemileucini Hemileucina Kentroleuca dukinfieldi
Hemileucinae Hemileucini Hemileucina Pseudodirphia niceros
Hemileucinae Hemileucini Hemileucina Dirphiopsis peggy
Hemileucinae Hemileucini Hemileucina Dirphiopsis flora
Hemileucinae Polythysanini Polythysana rubrescens
Hemileucinae Polythysanini Polythysana cinerascens
Hemileucinae Polythysanini [nec Hemileucini] Callodirphia arpi
Hemileucinae Polythysanini [nec Hemileucini] Adetomeris erythrops
Hemileucinae Polythysanini [nec Hemileucini] Adetomeris microphthalma
Hemileucinae Polythysanini [nec Hemileucini] Cinommata bistrigata
Hemileucinae Hemileucini Automeriina Automeris [nec Eubergioides] bertha
Hemileucinae Hemileucini Automeriina Automeris egeus
Hemileucinae Hemileucini Automeriina Automeris wayampi
Hemileucinae Hemileucini Automeriina Automeris zephyria
Hemileucinae Hemileucini Automeriina Automeris dognini
Hemileucinae Hemileucini Automeriina Leucanella fusca
Hemileucinae Hemileucini Automeriina Leucanella maasseni
Hemileucinae Hemileucini Automeriina Pseudautomeris rudloffi
Hemileucinae Hemileucini Automeriina Pseudautomeris lata
Hemileucinae Hemileucini Automeriina Hypermerina kasyi
Hemileucinae Hemileucini Automeriina Mexicantha garciorum
Hemileucinae Hemileucini Automeriina Molippa flavopiurica
Hemileucinae Hemileucini Automeriina Molippa tusina
Hemileucinae Hemileucini Automeriina Molippa placida
Hemileucinae Hemileucini Automeriina Erythromeris obscurior
Hemileucinae Hemileucini Automeriina Austrolippa convergens
Hemileucinae Hemileucini Automeriina Automerella chrisbrechlinae
Hemileucinae Hemileucini Automeriina Molippa superba
Hemileucinae Hemileucini Automeriina Auroraia eophila
Hemileucinae Hemileucini Automeriina Gamelia musta
Hemileucinae Hemileucini Automeriina Gamelia kaechi
Hemileucinae Hemileucini Automeriina Gamelia catharina
Hemileucinae Hemileucini Automeriina Catacantha jujuyensis
Hemileucinae Hemileucini Automeriina Catacantha oculata
Hemileucinae Hemileucini Automeriina Ancistrota plagia
Hemileucinae Hemileucini Automeriina Hylesiopsis festiva
Hemileucinae Hemileucini Automeriina Prohylesia rosalinda
Hemileucinae Hemileucini Automeriina Gamelioides sachai
Hemileucinae Hemileucini Automeriina Hylesia scortina
Hemileucinae Hemileucini Automeriina Hylesia bertrandi
Hemileucinae Hemileucini Automeriina Hylesia aeneides aeneides
Hemileucinae Hemileucini Automeriina Hylesia lineata
Hemileucinae Hemileucini Automeriina Eubergia caisa
Hemileucinae Hemileucini Automeriina Eubergia argyrea
Hemileucinae Hemileucini Automeriina Travassosula subfumata
Hemileucinae Hemileucini Automeriina Hyperchiria azteca
Hemileucinae Hemileucini Automeriina Hyperchiria aniris
Hemileucinae Hemileucini Automeriina Automeropsis umbrata
Hemileucinae Hemileucini Automeriina Automerina caudatula
Hemileucinae Hemileucini Automeriina Automerina auletes
Hemileucinae Hemileucini Automeriina Hyperchirioides bulaea
Hemileucinae Lonomiini [nec Hemileucini] Periga circaustralis
Hemileucinae Lonomiini [nec Hemileucini] Periga tatama
Hemileucinae Lonomiini [nec Hemileucini] Periga costaricana
Hemileucinae Lonomiini [nec Hemileucini] Lonomia descimoni
Hemileucinae Lonomiini [nec Hemileucini] Lonomia obliqua
Hemileucinae Lonomiini [nec Hemileucini] Lonomia santarosensis
Ceratocampinae Dryocampini Jaiba kesselringi
Ceratocampinae Dryocampini Mielkesia paranaensis
Ceratocampinae Dryocampini Megaceresa pulchra
Ceratocampinae Dryocampini Adelowalkeria plateada
Ceratocampinae Dryocampini Adelowalkeria tristygma
Ceratocampinae Dryocampini Scolesa viettei
Ceratocampinae Dryocampini Ceratesa hemirhodia
Ceratocampinae Dryocampini Rachesa breteuili
Ceratocampinae Dryocampini Rachesa kaechi
Ceratocampinae Dryocampini Oiticella luteciae
Ceratocampinae Dryocampini Oiticella convergens
Ceratocampinae Dryocampini Citioica homoea
Ceratocampinae Dryocampini Citioica anthonilis
Ceratocampinae Dryocampini Psilopygida walkeri
Ceratocampinae Dryocampini Psilopygida crispula
Ceratocampinae Dryocampini Ptiloscola santamartensis
Ceratocampinae Dryocampini Ptiloscola descimoni
Ceratocampinae Dryocampini Ceropoda tibialis
Ceratocampinae Dryocampini Adeloneivaia centrojason
Ceratocampinae Dryocampini Adeloneivaia catoxantha catoxantha
Ceratocampinae Dryocampini Adeloneivaia boisduvalii
Ceratocampinae Dryocampini Almeidella almeidai
Ceratocampinae Dryocampini Giacomellia bilineata
Ceratocampinae Dryocampini Anisota leucostygma
Ceratocampinae Dryocampini Anisota assimilis
Ceratocampinae Dryocampini Dryocampa rubicunda
Ceratocampinae Dryocampini Syssphinx delmaiziana
Ceratocampinae Dryocampini Syssphinx thiaucourti
Ceratocampinae Dryocampini Syssphinx molina
Ceratocampinae Dryocampini Dacunju jucunda
Ceratocampinae Dryocampini Cicia pelota
Ceratocampinae Dryocampini Psilopygoides oda
Ceratocampinae Dryocampini Othorene intermedia
Ceratocampinae Dryocampini Othorene purpurascens
Ceratocampinae Dryocampini Othorene winbrechlini
Ceratocampinae Bathyphlebiini Eacles imperialis imperialis
Ceratocampinae Bathyphlebiini Eacles barragani
Ceratocampinae [nec Hemileucinae] Bathyphlebiini [nec Hemileucini] Eacles [nec Catharisa] cerina
Ceratocampinae Bathyphlebiini Eacles ducalis
Ceratocampinae Bathyphlebiini Bathyphlebia eminentoides
Ceratocampinae Bathyphlebiini Bathyphlebia rufescens
Ceratocampinae Bathyphlebiini Eacles ormondei violacea
Ceratocampinae Bathyphlebiini Schausiella subochreata
Ceratocampinae Bathyphlebiini Schausiella maranhensis
Ceratocampinae Bathyphlebiini Neorcarnegia basirei
Ceratocampinae Citheroniini Citheronia regalis
Ceratocampinae Citheroniini Citheronia phoronea
Ceratocampinae Citheroniini Citheronula armata
Ceratocampinae Citheroniini Citheronioides collaris
Ceratocampinae Citheroniini Procitheronia principalis
Ceratocampinae Citheroniini Procitheronia fenestrata
Cercophaninae Janiodini Janiodes ecarcabuco
Cercophaninae Janiodini Janiodes oxapampensis
Cercophaninae Janiodini Janiodes manzanoi
Cercophaninae Cercophanini Neocercophana philippii
Cercophaninae Cercophanini Cercophana frauenfeldi
Cercophaninae Cercophanini Microdulia mirabilis
Oxyteninae Oxytenis modoccidentalis
Oxyteninae Oxytenis naemia
Oxyteninae Oxytenis peregrina
Oxyteninae Therinia buckleyi
Oxyteninae Therinia sp
Oxyteninae Therinia transversaria
Oxyteninae Homoeopteryx pinchcarchensis
Sphingidae Smerinthinae Smerinthini Smerinthus ocellata
Sphingidae Smerinthinae Smerinthini Paonias myops
Sphingidae Smerinthinae Smerinthini Pachysphinx occidentalis
Sphingidae Smerinthinae Smerinthini Laothoe populi
Sphingidae Smerinthinae Coequosa australasiae
Sphingidae Sphinginae Sphingini Acherontia atropos
Sphingidae Sphinginae Sphingini Agrius convolvuli
Sphingidae Langiinae Langia zenzeroides
Sphingidae Macroglossinae Macroglossini Daphnis nerii
Sphingidae Macroglossinae Hemarini Hemaris fuciformis
Bombycidae Epiinae Quentalia mailynae
Endromidae Endromis versicolora
Eupterotidae Janinae Stenoglene roseus
Eupterotidae Striphnopteryginae Phiala sp
Brahmaeidae Acanthobrahmaea europaea
Brahmaeidae Lemonia taraxaci
