## Supplementary material for "Phylogenomics Illuminates the Evolutionary History of Wild Silkmoths in Space and Time (Lepidoptera: Saturniidae)": Figure S: FigureS3_taxa_properties.pdf

Figure S3A. boxplot of GC content and LB score for the different groups (as calculated from the concatenation of the 1024 UCEs; cf Table S3)

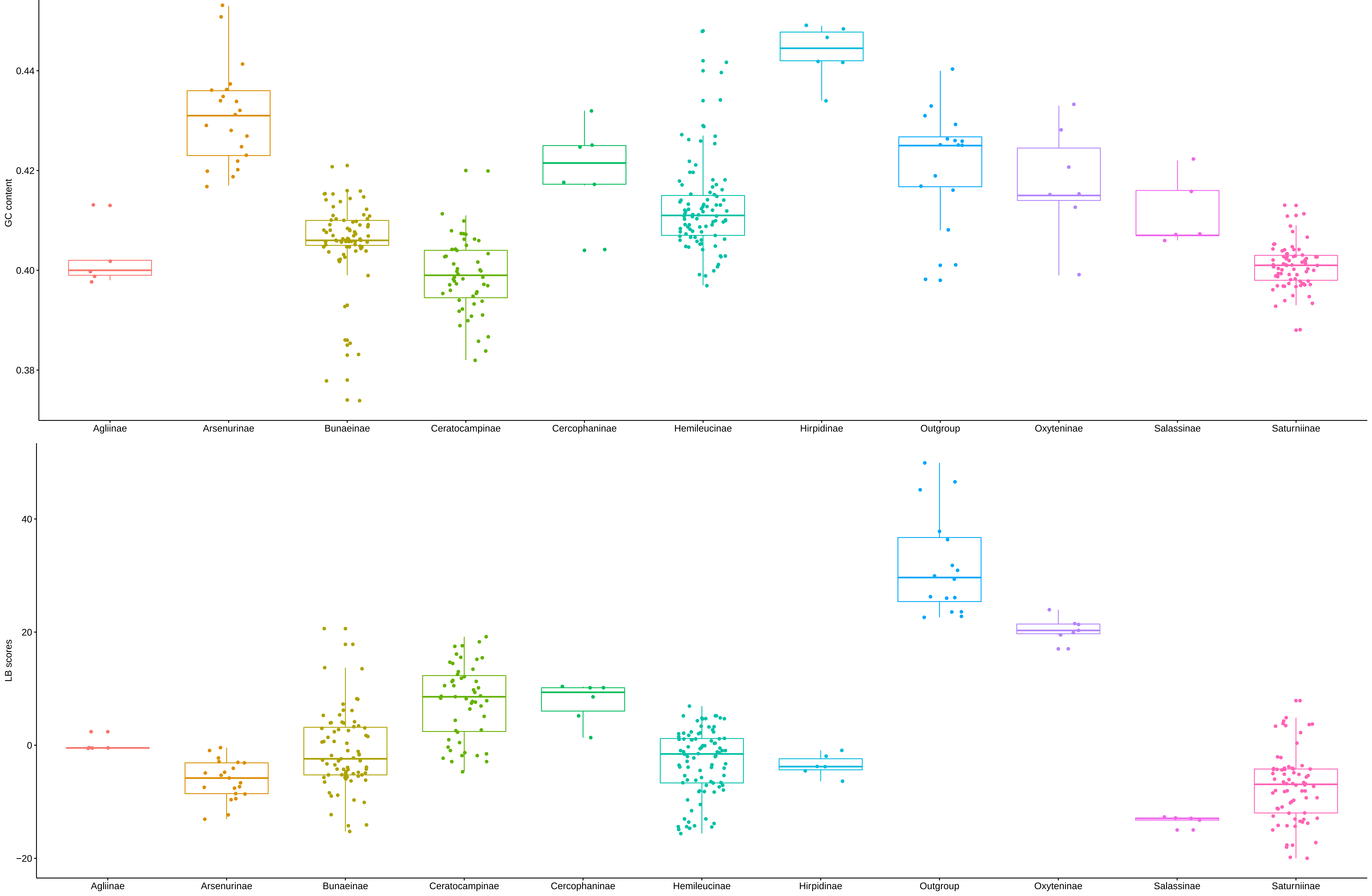

**Figure S3B. Dendrogram of samples based on their GC content in each of the 1024 UCEs**

(distance matrix: Gower; clustering method: Ward.D2)

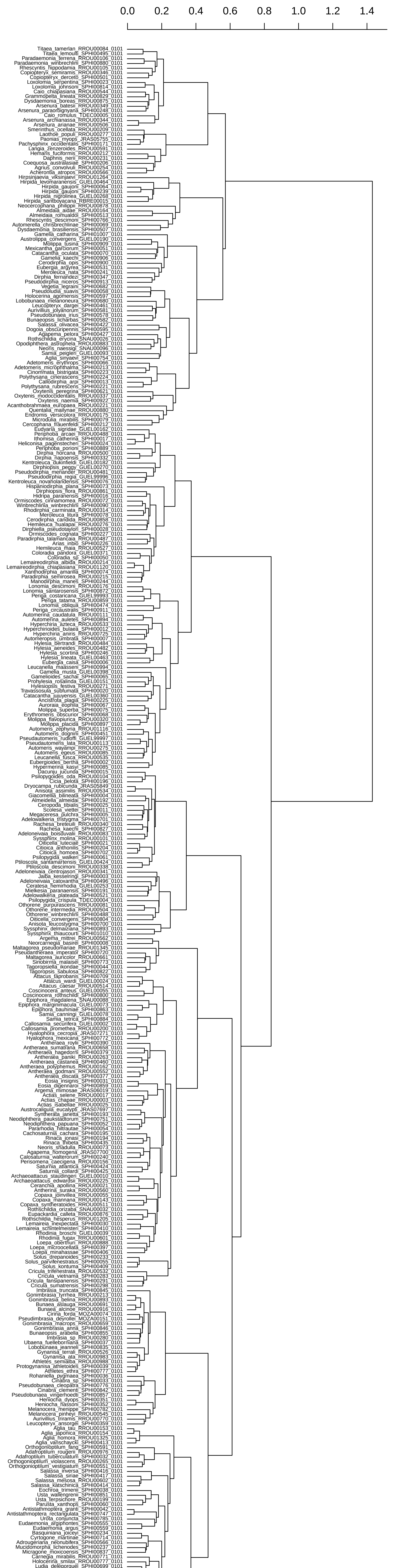

**Figure S3C. Dendrogram of samples based on their LB heterogeneity scores in each of the 1024 UCEs**

(distance matrix: Gower; clustering method: Ward.D2)

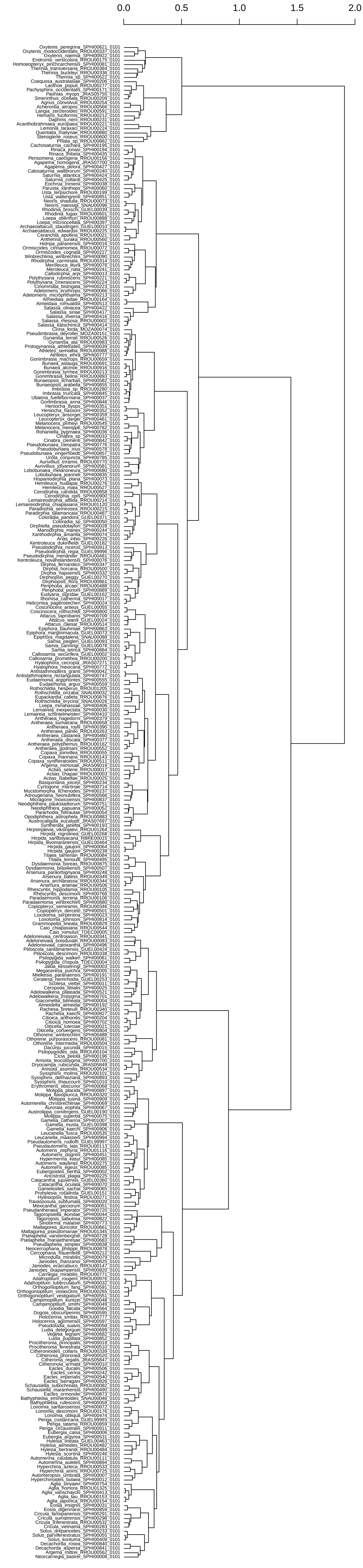
