## Supplementary material for "Phylogenomics Illuminates the Evolutionary History of Wild Silkmoths in Space and Time (Lepidoptera: Saturniidae)": Figure S: FigureS4_chronograms.pdf

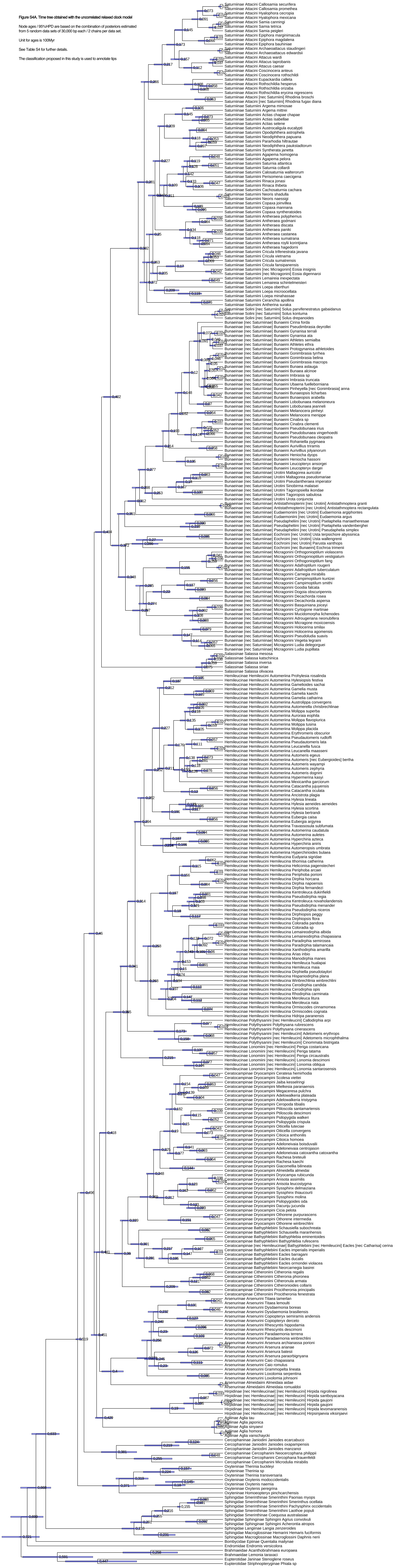

Figure S4A. Time tree obtained with the uncorrelated relaxed clock model

Node ages / 95%HPD are based on the combination of posteriors estimated from 5 random data sets of 30,000 bp each / 2 chains per data set.

Unit for ages is 100Myr

See Table S4 for further details.

The classification proposed in this study is used to annotate tips

Figure S4B. Time tree obtained with the correlated relaxed clock model

Node ages / 95%HPD are based on the combination of posteriors estimated from 5 random data sets of 30,000 bp each / 2 chains per data set.

Unit for ages is 100Myr

See Table S4 for further details.

The classification proposed in this study is used to annotate tips

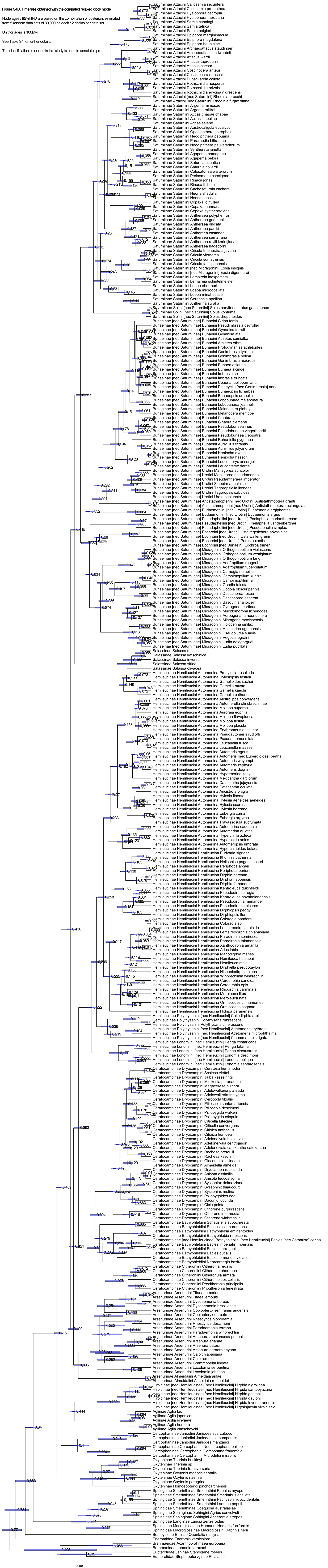
