## Supplementary material for "Phylogenomics Illuminates the Evolutionary History of Wild Silkmoths in Space and Time (Lepidoptera: Saturniidae)": Figure S: FigureS5_biogeography.pdf

**Figure S5A. BAYAREALIKE**

ancstates: global optim, 5 areas max. d=0.0068; e=0.0141; j=0; LnL=-385.06

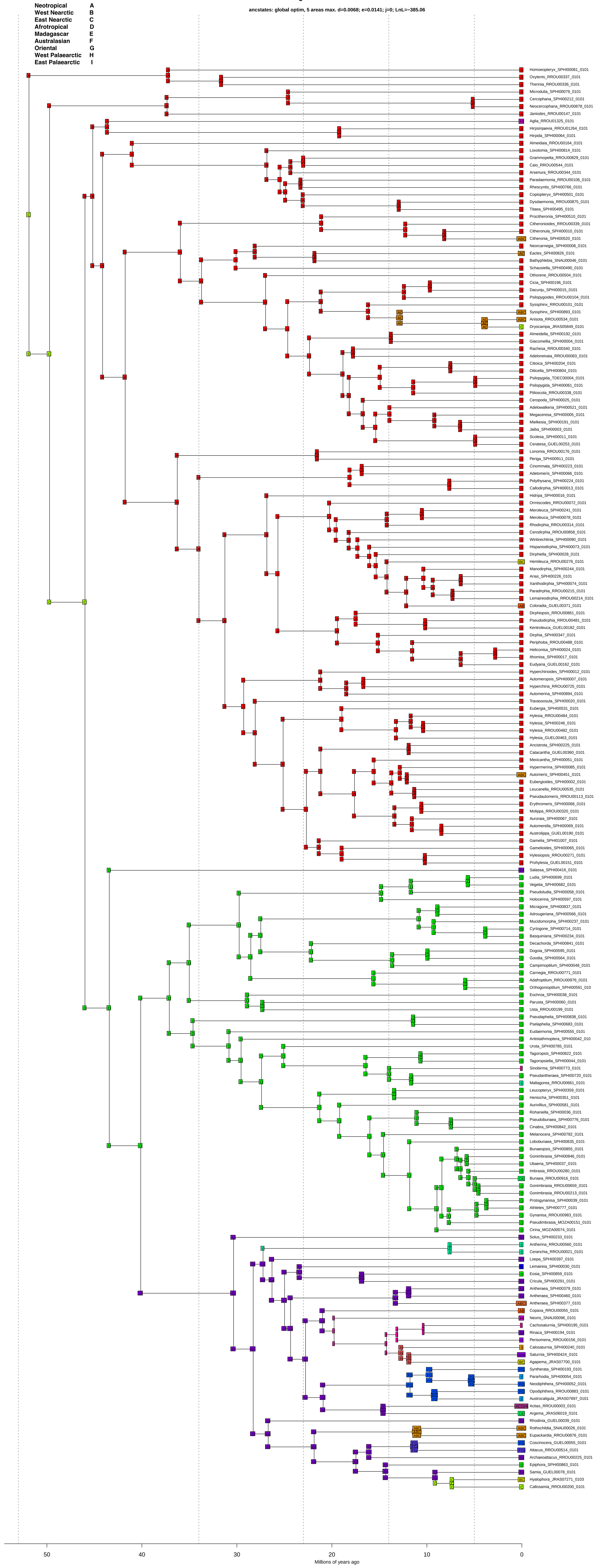

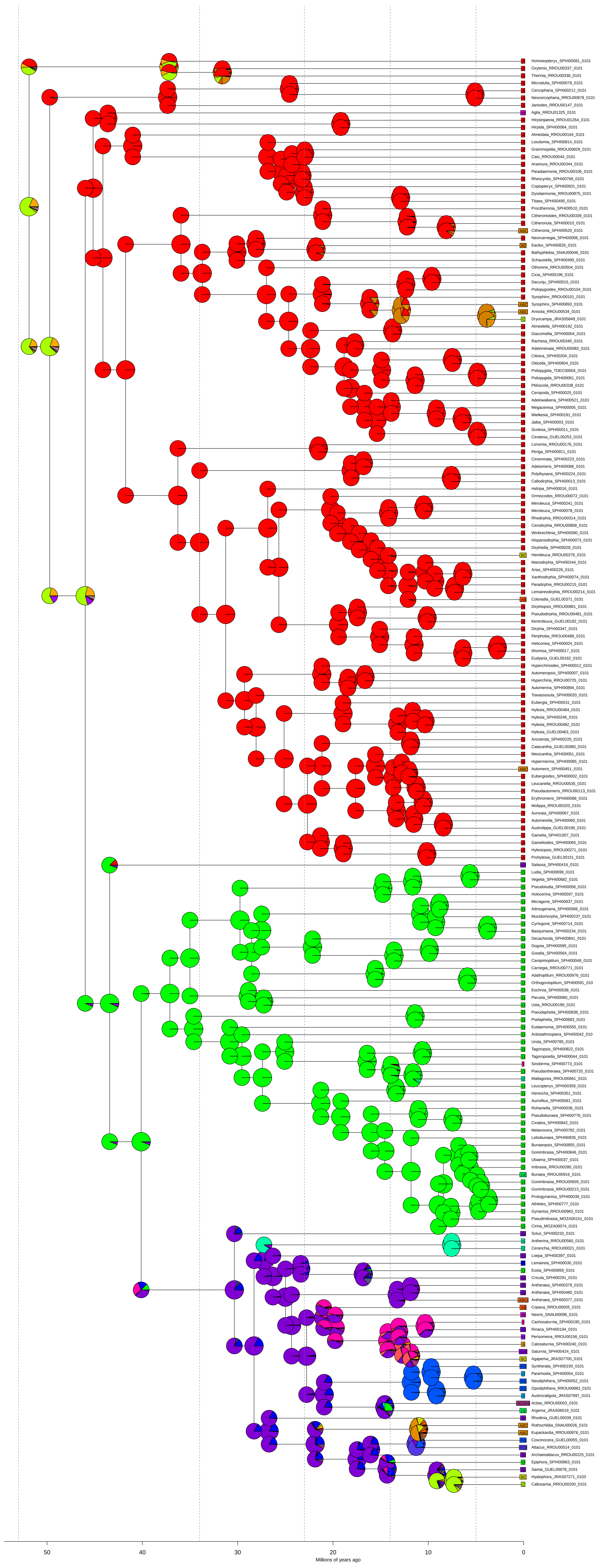
A Homoeopteryx_SPHI00081_0101
A Oxytenis_RROU00337_0101
A Therinia_RROU00336_0101
A Microdulia_SPHI00079_0101
A Cercophana_SPHI00212_0101
A Neocercophana_RROU00878_0101
A Janiodes_RROU00147_0101
HI Aglia_RROU01325_0101
A Hirpsinjaevia_RROU01264_0101
A Hirpida_SPHI00064_0101
A Almeidaia_RROU00164_0101
A Loxolomia_SPHI00814_0101
A Grammopelta_RROU00829_0101
A Caio_RROU00544_0101
A Arsenura_RROU00344_0101
A Paradaemonia_RROU00106_0101
A Rhescyntis_SPHI00766_0101
A Copiopteryx_SPHI00501_0101
A Dysdaemonia_RROU00875_0101
A Titaea_SPHI00495_0101
A Procitheronia_SPHI00510_0101
A Citheronioides_RROU00339_0101
A Citheronula_SPHI00010_0101
ABC Citheronia_SPHI00520_0101
A Neorcarnegia_SPHI00008_0101
AC Eacles_SPHI00826_0101
A Bathyphlebia_SNAU00046_0101
A Schausiella_SPHI00490_0101
A Othorene_RROU00504_0101
A Cicia_SPHI00196_0101
A Dacunju_SPHI00015_0101
A Psilopygoides_RROU00104_0101
A Syssphinx_RROU00101_0101
ABC Syssphinx_SPHI00893_0101
ABC Anisota_RROU00534_0101
C Dryocampa_JRAS05849_0101
A Almeidella_SPHI00192_0101
A Giacomellia_SPHI00004_0101
A Rachesa_RROU00340_0101
A Adeloneivaia_RROU00083_0101
A Citioica_SPHI00204_0101
A Oiticella_SPHI00804_0101
A Psilopygida_TDEC00004_0101
A Psilopygida_SPHI00061_0101
A Ptiloscola_RROU00338_0101
A Ceropoda_SPHI00025_0101
A Adelowalkeria_SPHI00521_0101
A Megaceresa_SPHI00005_0101
A Mielkesia_SPHI00191_0101
A Jaiba_SPHI00003_0101
A Scolesa_SPHI00011_0101
A Ceratesa_GUEL00253_0101
A Lonomia_RROU00176_0101
A Periga_SPHI00911_0101
A Cinommata_SPHI00223_0101
A Adetomeris_SPHI00066_0101
A Polythysana_SPHI00224_0101
A Callodirphia_SPHI00013_0101
A Hidripa_SPHI00016_0101
A Ormiscodes_RROU00072_0101
A Meroleuca_SPHI00241_0101
A Meroleuca_SPHI00078_0101
A Rhodirphia_RROU00314_0101
A Cerodirphia_RROU00858_0101
A Winbrechlinia_SPHI00090_0101
A Hispaniodirphia_SPHI00073_0101
A Dirphiella_SPHI00028_0101
BC Hemileuca_RROU00276_0101
A Manodirphia_SPHI00244_0101
A Arias_SPHI00226_0101
A Xanthodirphia_SPHI00074_0101
A Paradirphia_RROU00215_0101
A Lemaireodirphia_RROU00214_0101
AB Coloradia_GUEL00371_0101
A Dirphiopsis_RROU00861_0101
A Pseudodirphia_RROU00481_0101
A Kentroleuca_GUEL00182_0101
A Dirphia_SPHI00347_0101
A Periphoba_RROU00488_0101
A Heliconisa_SPHI00024_0101
A Ithomisa_SPHI00017_0101
A Eudyaria_GUEL00162_0101
A Hyperchirioides_SPHI00012_0101
A Automeropsis_SPHI00007_0101
A Hyperchiria_RROU00725_0101
A Automerina_SPHI00894_0101
A Travassosula_SPHI00020_0101
A Eubergia_SPHI00531_0101
A Hylesia_RROU00484_0101
A Hylesia_SPHI00246_0101
A Hylesia_RROU00482_0101
A Hylesia_GUEL00463_0101
A Ancistrota_SPHI00225_0101
A Catacantha_GUEL00360_0101
A Mexicantha_SPHI00051_0101
A Hypermerina_SPHI00085_0101
ABC Automeris_SPHI00451_0101
A Eubergioides_SPHI00002_0101
A Leucanella_RROU00535_0101
A Pseudautomeris_RROU00113_0101
A Erythromeris_SPHI00068_0101
A Molippa_RROU00320_0101
A Auroraia_SPHI00067_0101
A Automerella_SPHI00069_0101
A Austrolippa_GUEL00190_0101
A Gamelia_SPHI01007_0101
A Gamelioides_SPHI00065_0101
A Hylesiopsis_RROU00271_0101
A Prohylesia_GUEL00151_0101
GI Salassa_SPHI00416_0101
D Ludia_SPHI00699_0101
D Vegetia_SPHI00682_0101
D Pseudoludia_SPHI00058_0101
D Holocerina_SPHI00597_0101
D Micragone_SPHI00837_0101
D Adrougeriana_SPHI00566_0101
D Mucidomorpha_SPHI00237_0101
D Cyrtogone_SPHI00714_0101
D Basquiniana_SPHI00234_0101
D Decachorda_SPHI00841_0101
D Dogoia_SPHI00595_0101
D Goodia_SPHI00564_0101
D Campimoptilum_SPHI00048_0101
D Carnegia_RROU00771_0101
D Adafroptilum_RROU00976_0101
D Orthogonioptilum_SPHI00591_010
D Eochroa_SPHI00038_0101
D Parusta_SPHI00060_0101
D Usta_RROU00199_0101
D Pseudaphelia_SPHI00838_0101
D Pselaphelia_SPHI00683_0101
D Eudaemonia_SPHI00555_0101
D Antistathmoptera_SPHI00042_010
D Urota_SPHI00785_0101
D Tagoropsis_SPHI00822_0101
D Tagoropsiella_SPHI00044_0101
I Sinobirma_SPHI00773_0101
D Pseudantheraea_SPHI00720_0101
E Maltagorea_RROU00661_0101
D Leucopteryx_SPHI00359_0101
D Heniocha_SPHI00351_0101
D Aurivillius_SPHI00581_0101
D Rohaniella_SPHI00036_0101
D Pseudobunaea_SPHI00776_0101
D Cinabra_SPHI00842_0101
D Melanocera_SPHI00782_0101
D Lobobunaea_SPHI00835_0101
D Bunaeopsis_SPHI00855_0101
D Gonimbrasia_SPHI00846_0101
D Ubaena_SPHI00037_0101
D Imbrasia_RROU00280_0101
DE Bunaea_RROU00916_0101
D Gonimbrasia_RROU00659_0101
D Gonimbrasia_RROU00213_0101
D Protogynanisa_SPHI00039_0101
D Athletes_SPHI00777_0101
D Gynanisa_RROU00983_0101
D Pseudimbrasia_MOZA00151_0101
D Cirina_MOZA00074_0101
GI Solus_SPHI00233_0101
E Antherina_RROU00560_0101
E Ceranchia_RROU00021_0101
GI Loepa_SPHI00397_0101
G Lemaireia_SPHI00030_0101
D Eosia_SPHI00859_0101
GI Cricula_SPHI00291_0101
GI Antheraea_SPHI00379_0101
GI Antheraea_SPHI00460_0101
ABCI Antheraea_SPHI00377_0101
AB Copaxa_RROU00055_0101
HI Neoris_SNAU00096_0101
I Cachosaturnia_SPHI00195_0101
GI Rinaca_SPHI00194_0101
H Perisomena_RROU00156_0101
B Calosaturnia_SPHI00240_0101
GHI Saturnia_SPHI00424_0101
BC Agapema_JRAS07700_0101
FG Syntherata_SPHI00193_0101
F Pararhodia_SPHI00054_0101
FG Neodiphthera_SPHI00052_0101
FG Opodiphthera_RROU00883_0101
F Austrocaligula_JRAS07697_0101
ACGHI Actias_RROU00003_0101
DE Argema_JRAS06019_0101
GI Rhodinia_GUEL00039_0101
ABC Rothschildia_SNAU00026_0101
ABC Eupackardia_RROU00876_0101
FG Coscinocera_GUEL00055_0101
FGI Attacus_RROU00514_0101
GI Archaeoattacus_RROU00225_0101
D Epiphora_SPHI00863_0101
GI Samia_GUEL00078_0101
BC Hyalophora_JRAS07271_0103
C Callosamia_RROU00200_0101
50
40
30
20
10
0
Millions of years ago

**Figure S5B. BAYAREALIKE+J**

ancstates: global optim, 5 areas max. d=0.0033; e=0.0013; j=0.0194; LnL=-256.45

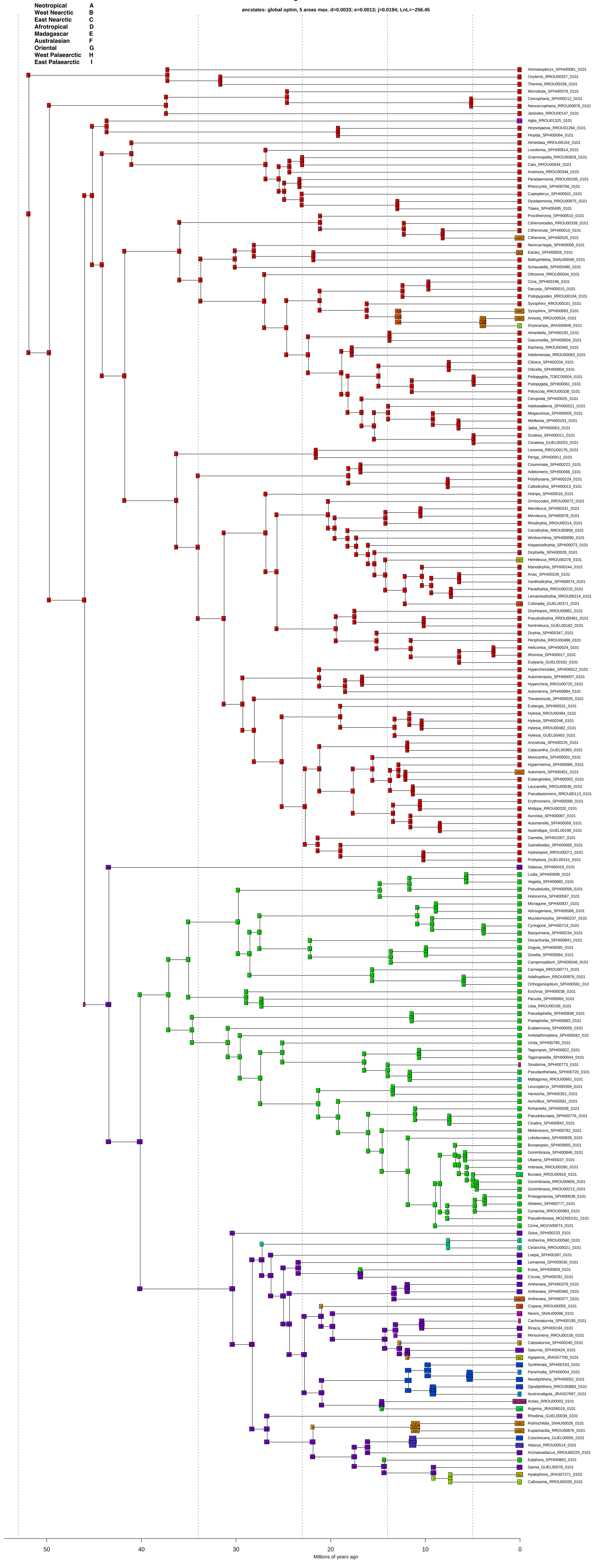

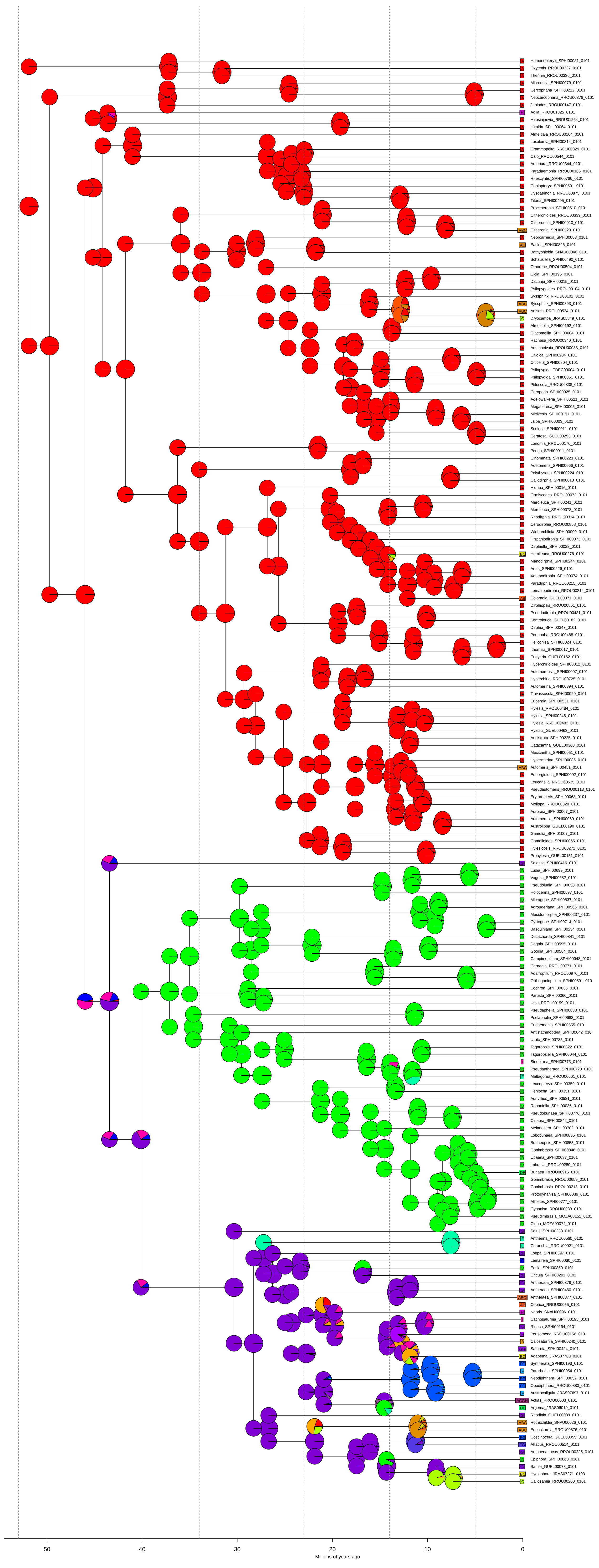

A Homoeopteryx_SPHI00081_0101
A Oxytenis_RROU00337_0101
A Therinia_RROU00336_0101
A Microdulia_SPHI00079_0101
A Cercophana_SPHI00212_0101
A Neocercophana_RROU00878_0101
A Janiodes_RROU00147_0101
HI Aglia_RROU01325_0101
A Hirpsinjaevia_RROU01264_0101
A Hirpida_SPHI00064_0101
A Almeidaia_RROU00164_0101
A Loxolomia_SPHI00814_0101
A Grammopelta_RROU00829_0101
A Caio_RROU00544_0101
A Arsenura_RROU00344_0101
A Paradaemonia_RROU00106_0101
A Rhescyntis_SPHI00766_0101
A Copiopteryx_SPHI00501_0101
A Dysdaemonia_RROU00875_0101
A Titaea_SPHI00495_0101
A Procitheronia_SPHI00510_0101
A Citheronioides_RROU00339_0101
A Citheronula_SPHI00010_0101
ABC Citheronia_SPHI00520_0101
A Neorcarnegia_SPHI00008_0101
AC Eacles_SPHI00826_0101
A Bathyphlebia_SNAU00046_0101
A Schausiella_SPHI00490_0101
A Othorene_RROU00504_0101
A Cicia_SPHI00196_0101
A Dacunju_SPHI00015_0101
A Psilopygoides_RROU00104_0101
A Syssphinx_RROU00101_0101
ABC Syssphinx_SPHI00893_0101
ABC Anisota_RROU00534_0101
C Dryocampa_JRAS05849_0101
A Almeidella_SPHI00192_0101
A Giacomellia_SPHI00004_0101
A Rachesa_RROU00340_0101
A Adeloneivaia_RROU00083_0101
A Citioica_SPHI00204_0101
A Oiticella_SPHI00804_0101
A Psilopygida_TDEC00004_0101
A Psilopygida_SPHI00061_0101
A Ptiloscola_RROU00338_0101
A Ceropoda_SPHI00025_0101
A Adelowalkeria_SPHI00521_0101
A Megaceresa_SPHI00005_0101
A Mielkesia_SPHI00191_0101
A Jaiba_SPHI00003_0101
A Scolesa_SPHI00011_0101
A Ceratesa_GUEL00253_0101
A Lonomia_RROU00176_0101
A Periga_SPHI00911_0101
A Cinommata_SPHI00223_0101
A Adetomeris_SPHI00066_0101
A Polythysana_SPHI00224_0101
A Callodirphia_SPHI00013_0101
A Hidripa_SPHI00016_0101
A Ormiscodes_RROU00072_0101
A Meroleuca_SPHI00241_0101
A Meroleuca_SPHI00078_0101
A Rhodirphia_RROU00314_0101
A Cerodirphia_RROU00858_0101
A Winbrechlinia_SPHI00090_0101
A Hispaniodirphia_SPHI00073_0101
A Dirphiella_SPHI00028_0101
BC Hemileuca_RROU00276_0101
A Manodirphia_SPHI00244_0101
A Arias_SPHI00226_0101
A Xanthodirphia_SPHI00074_0101
A Paradirphia_RROU00215_0101
A Lemaireodirphia_RROU00214_0101
AB Coloradia_GUEL00371_0101
A Dirphiopsis_RROU00861_0101
A Pseudodirphia_RROU00481_0101
A Kentroleuca_GUEL00182_0101
A Dirphia_SPHI00347_0101
A Periphoba_RROU00488_0101
A Heliconisa_SPHI00024_0101
A Ithomisa_SPHI00017_0101
A Eudyaria_GUEL00162_0101
A Hyperchirioides_SPHI00012_0101
A Automeropsis_SPHI00007_0101
A Hyperchiria_RROU00725_0101
A Automerina_SPHI00894_0101
A Travassosula_SPHI00020_0101
A Eubergia_SPHI00531_0101
A Hylesia_RROU00484_0101
A Hylesia_SPHI00246_0101
A Hylesia_RROU00482_0101
A Hylesia_GUEL00463_0101
A Ancistrota_SPHI00225_0101
A Catacantha_GUEL00360_0101
A Mexicantha_SPHI00051_0101
A Hypermerina_SPHI00085_0101
ABC Automeris_SPHI00451_0101
A Eubergioides_SPHI00002_0101
A Leucanella_RROU00535_0101
A Pseudautomeris_RROU00113_0101
A Erythromeris_SPHI00068_0101
A Molippa_RROU00320_0101
A Auroraia_SPHI00067_0101
A Automerella_SPHI00069_0101
A Austrolippa_GUEL00190_0101
A Gamelia_SPHI01007_0101
A Gamelioides_SPHI00065_0101
A Hylesiopsis_RROU00271_0101
A Prohylesia_GUEL00151_0101
GI Salassa_SPHI00416_0101
D Ludia_SPHI00699_0101
D Vegetia_SPHI00682_0101
D Pseudoludia_SPHI00058_0101
D Holocerina_SPHI00597_0101
D Micragone_SPHI00837_0101
D Adrougeriana_SPHI00566_0101
D Mucidomorpha_SPHI00237_0101
D Cyrtogone_SPHI00714_0101
D Basquiniana_SPHI00234_0101
D Decachorda_SPHI00841_0101
D Dogoia_SPHI00595_0101
D Goodia_SPHI00564_0101
D Campimoptilum_SPHI00048_0101
D Carnegia_RROU00771_0101
D Adafroptilum_RROU00976_0101
D Orthogonioptilum_SPHI00591_010
D Eochroa_SPHI00038_0101
D Parusta_SPHI00060_0101
D Usta_RROU00199_0101
D Pseudaphelia_SPHI00838_0101
D Pselaphelia_SPHI00683_0101
D Eudaemonia_SPHI00555_0101
D Antistathmoptera_SPHI00042_010
D Urota_SPHI00785_0101
D Tagoropsis_SPHI00822_0101
D Tagoropsiella_SPHI00044_0101
I Sinobirma_SPHI00773_0101
D Pseudantheraea_SPHI00720_0101
E Maltagorea_RROU00661_0101
D Leucopteryx_SPHI00359_0101
D Heniocha_SPHI00351_0101
D Aurivillius_SPHI00581_0101
D Rohaniella_SPHI00036_0101
D Pseudobunaea_SPHI00776_0101
D Cinabra_SPHI00842_0101
D Melanocera_SPHI00782_0101
D Lobobunaea_SPHI00835_0101
D Bunaeopsis_SPHI00855_0101
D Gonimbrasia_SPHI00846_0101
D Ubaena_SPHI00037_0101
D Imbrasia_RROU00280_0101
DE Bunaea_RROU00916_0101
D Gonimbrasia_RROU00659_0101
D Gonimbrasia_RROU00213_0101
D Protogynanisa_SPHI00039_0101
D Athletes_SPHI00777_0101
D Gynanisa_RROU00983_0101
D Pseudimbrasia_MOZA00151_0101
D Cirina_MOZA00074_0101
GI Solus_SPHI00233_0101
E Antherina_RROU00560_0101
E Ceranchia_RROU00021_0101
GI Loepa_SPHI00397_0101
G Lemaireia_SPHI00030_0101
D Eosia_SPHI00859_0101
GI Cricula_SPHI00291_0101
GI Antheraea_SPHI00379_0101
GI Antheraea_SPHI00460_0101
ABCI Antheraea_SPHI00377_0101
AB Copaxa_RROU00055_0101
HI Neoris_SNAU00096_0101
I Cachosaturnia_SPHI00195_0101
GI Rinaca_SPHI00194_0101
H Perisomena_RROU00156_0101
B Calosaturnia_SPHI00240_0101
GHI Saturnia_SPHI00424_0101
BC Agapema_JRAS07700_0101
FG Syntherata_SPHI00193_0101
F Pararhodia_SPHI00054_0101
FG Neodiphthera_SPHI00052_0101
FG Opodiphthera_RROU00883_0101
F Austrocaligula_JRAS07697_0101
ACGHI Actias_RROU00003_0101
DE Argema_JRAS06019_0101
GI Rhodinia_GUEL00039_0101
ABC Rothschildia_SNAU00026_0101
ABC Eupackardia_RROU00876_0101
FG Coscinocera_GUEL00055_0101
FGI Attacus_RROU00514_0101
GI Archaeoattacus_RROU00225_0101
D Epiphora_SPHI00863_0101
GI Samia_GUEL00078_0101
BC Hyalophora_JRAS07271_0103
C Callosamia_RROU00200_0101
50
40
30
20
10
0
Millions of years ago

### Figure S5C. DEC

ancstates: global optim, 5 areas max. d=0.0078; e=0.0053; j=0; LnL=-360.96

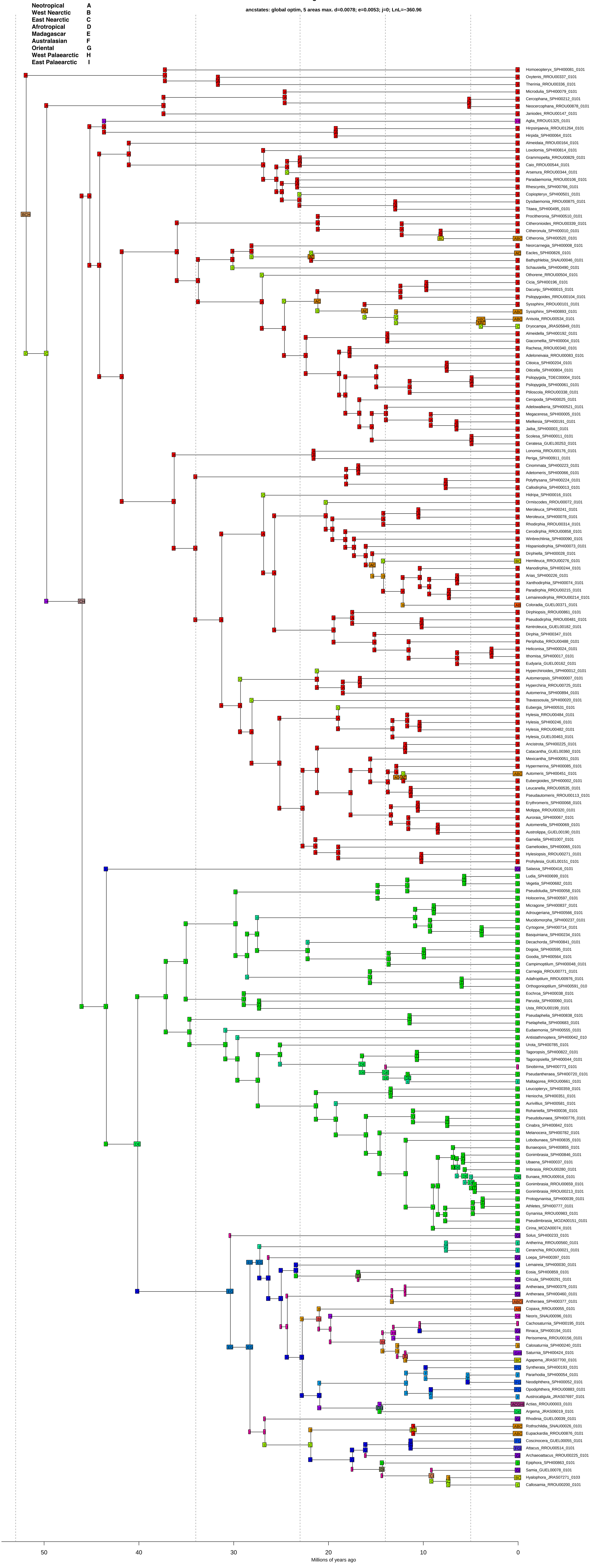

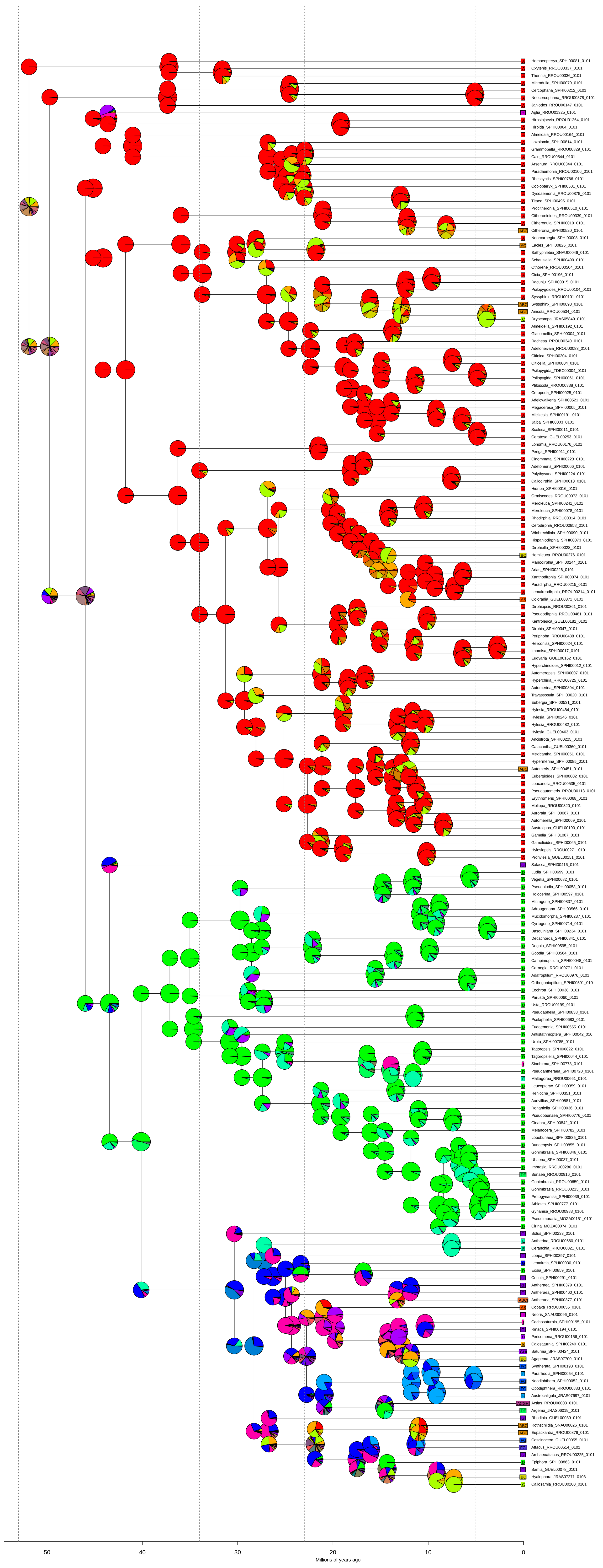
Homoeopteryx_SPHI00081_0101
Oxytenis_RROU00337_0101
Therinia_RROU00336_0101
Microdulia_SPHI00079_0101
Cercophana_SPHI00212_0101
Neocercophana_RROU00878_0101
Janiodes_RROU00147_0101
Aglia_RROU01325_0101
Hirpsinjaevia_RROU01264_0101
Hirpida_SPHI00064_0101
Almeidaia_RROU00164_0101
Loxolomia_SPHI00814_0101
Grammopelta_RROU00829_0101
Caio_RROU00544_0101
Arsenura_RROU00344_0101
Paradaemonia_RROU00106_0101
Rhescyntis_SPHI00766_0101
Copiopteryx_SPHI00501_0101
Dysdaemonia_RROU00875_0101
Titaea_SPHI00495_0101
Procitheronia_SPHI00510_0101
Citheronioides_RROU00339_0101
Citheronula_SPHI00010_0101
Citheronia_SPHI00520_0101
Neorcarnegia_SPHI00008_0101
Eacles_SPHI00826_0101
Bathyphlebia_SNAU00046_0101
Schausiella_SPHI00490_0101
Othorene_RROU00504_0101
Cicia_SPHI00196_0101
Dacunju_SPHI00015_0101
Psilopygoides_RROU00104_0101
Syssphinx_RROU00101_0101
Syssphinx_SPHI00893_0101
Anisota_RROU00534_0101
Dryocampa_JRAS05849_0101
Almeidella_SPHI00192_0101
Giacomellia_SPHI00004_0101
Rachesa_RROU00340_0101
Adeloneivaia_RROU00083_0101
Citioica_SPHI00204_0101
Oiticella_SPHI00804_0101
Psilopygida_TDEC00004_0101
Psilopygida_SPHI00061_0101
Ptiloscola_RROU00338_0101
Ceropoda_SPHI00025_0101
Adelowalkeria_SPHI00521_0101
Megaceresa_SPHI00005_0101
Mielkesia_SPHI00191_0101
Jaiba_SPHI00003_0101
Scolesa_SPHI00011_0101
Ceratesa_GUEL00253_0101
Lonomia_RROU00176_0101
Periga_SPHI00911_0101
Cinommata_SPHI00223_0101
Adetomeris_SPHI00066_0101
Polythysana_SPHI00224_0101
Callodirphia_SPHI00013_0101
Hidripa_SPHI00016_0101
Ormiscodes_RROU00072_0101
Meroleuca_SPHI00241_0101
Meroleuca_SPHI00078_0101
Rhodirphia_RROU00314_0101
Cerodirphia_RROU00858_0101
Winbrechlinia_SPHI00090_0101
Hispaniodirphia_SPHI00073_0101
Dirphiella_SPHI00028_0101
Hemileuca_RROU00276_0101
Manodirphia_SPHI00244_0101
Arias_SPHI00226_0101
Xanthodirphia_SPHI00074_0101
Paradirphia_RROU00215_0101
Lemaireodirphia_RROU00214_0101
Coloradia_GUEL00371_0101
Dirphiopsis_RROU00861_0101
Pseudodirphia_RROU00481_0101
Kentroleuca_GUEL00182_0101
Dirphia_SPHI00347_0101
Periphoba_RROU00488_0101
Heliconisa_SPHI00024_0101
Ithomisa_SPHI00017_0101
Eudyaria_GUEL00162_0101
Hyperchirioides_SPHI00012_0101
Automeropsis_SPHI00007_0101
Hyperchiria_RROU00725_0101
Automerina_SPHI00894_0101
Travassosula_SPHI00020_0101
Eubergia_SPHI00531_0101
Hylesia_RROU00484_0101
Hylesia_SPHI00246_0101
Hylesia_RROU00482_0101
Hylesia_GUEL00463_0101
Ancistrota_SPHI00225_0101
Catacantha_GUEL00360_0101
Mexicantha_SPHI00051_0101
Hypermerina_SPHI00085_0101
Automeris_SPHI00451_0101
Eubergioides_SPHI00002_0101
Leucanella_RROU00535_0101
Pseudautomeris_RROU00113_0101
Erythromeris_SPHI00068_0101
Molippa_RROU00320_0101
Auroraia_SPHI00067_0101
Automerella_SPHI00069_0101
Austrolippa_GUEL00190_0101
Gamelia_SPHI01007_0101
Gamelioides_SPHI00065_0101
Hylesiopsis_RROU00271_0101
Prohylesia_GUEL00151_0101
Salassa_SPHI00416_0101
Ludia_SPHI00699_0101
Vegetia_SPHI00682_0101
Pseudoludia_SPHI00058_0101
Holocerina_SPHI00597_0101
Micragone_SPHI00837_0101
Adrougeriana_SPHI00566_0101
Mucidomorpha_SPHI00237_0101
Cyrtogone_SPHI00714_0101
Basquiniana_SPHI00234_0101
Decachorda_SPHI00841_0101
Dogoia_SPHI00595_0101
Goodia_SPHI00564_0101
Campimoptilum_SPHI00048_0101
Carnegia_RROU00771_0101
Adafroptilum_RROU00976_0101
Orthogonioptilum_SPHI00591_010
Eochroa_SPHI00038_0101
Parusta_SPHI00060_0101
Usta_RROU00199_0101
Pseudaphelia_SPHI00838_0101
Pselaphelia_SPHI00683_0101
Eudaemonia_SPHI00555_0101
Antistathmoptera_SPHI00042_010
Urota_SPHI00785_0101
Tagoropsis_SPHI00822_0101
Tagoropsiella_SPHI00044_0101
Sinobirma_SPHI00773_0101
Pseudantheraea_SPHI00720_0101
Maltagorea_RROU00661_0101
Leucopteryx_SPHI00359_0101
Heniocha_SPHI00351_0101
Aurivillius_SPHI00581_0101
Rohaniella_SPHI00036_0101
Pseudobunaea_SPHI00776_0101
Cinabra_SPHI00842_0101
Melanocera_SPHI00782_0101
Lobobunaea_SPHI00835_0101
Bunaeopsis_SPHI00855_0101
Gonimbrasia_SPHI00846_0101
Ubaena_SPHI00037_0101
Imbrasia_RROU00280_0101
Bunaea_RROU00916_0101
Gonimbrasia_RROU00659_0101
Gonimbrasia_RROU00213_0101
Protogynanisa_SPHI00039_0101
Athletes_SPHI00777_0101
Gynanisa_RROU00983_0101
Pseudimbrasia_MOZA00151_0101
Cirina_MOZA00074_0101
Solus_SPHI00233_0101
Antherina_RROU00560_0101
Ceranchia_RROU00021_0101
Loepa_SPHI00397_0101
Lemaireia_SPHI00030_0101
Eosia_SPHI00859_0101
Cricula_SPHI00291_0101
Antheraea_SPHI00379_0101
Antheraea_SPHI00460_0101
Antheraea_SPHI00377_0101
Copaxa_RROU00055_0101
Neoris_SNAU00096_0101
Cachosaturnia_SPHI00195_0101
Rinaca_SPHI00194_0101
Perisomena_RROU00156_0101
Calosaturnia_SPHI00240_0101
Saturnia_SPHI00424_0101
Agapema_JRAS07700_0101
Syntherata_SPHI00193_0101
Pararhodia_SPHI00054_0101
Neodiphthera_SPHI00052_0101
Opodiphthera_RROU00883_0101
Austrocaligula_JRAS07697_0101
Actias_RROU00003_0101
Argema_JRAS06019_0101
Rhodinia_GUEL00039_0101
Rothschildia_SNAU00026_0101
Eupackardia_RROU00876_0101
Coscinocera_GUEL00055_0101
Attacus_RROU00514_0101
Archaeoattacus_RROU00225_0101
Epiphora_SPHI00863_0101
Samia_GUEL00078_0101
Hyalophora_JRAS07271_0103
Callosamia_RROU00200_0101
50
40
30
20
10
0
Millions of years ago

### Figure S5D. DEC+J

ancstates: global optim, 5 areas max. d=0.0053; e=0; j=0.0142; LnL=-267.23

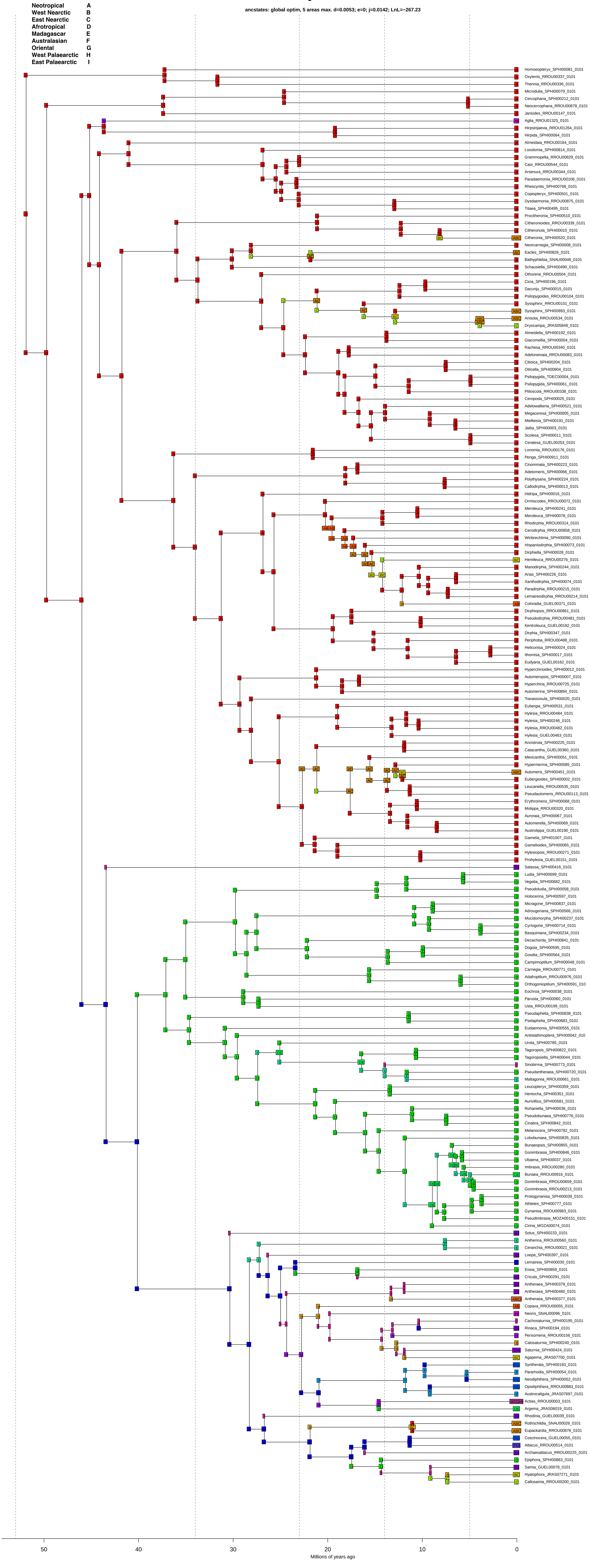

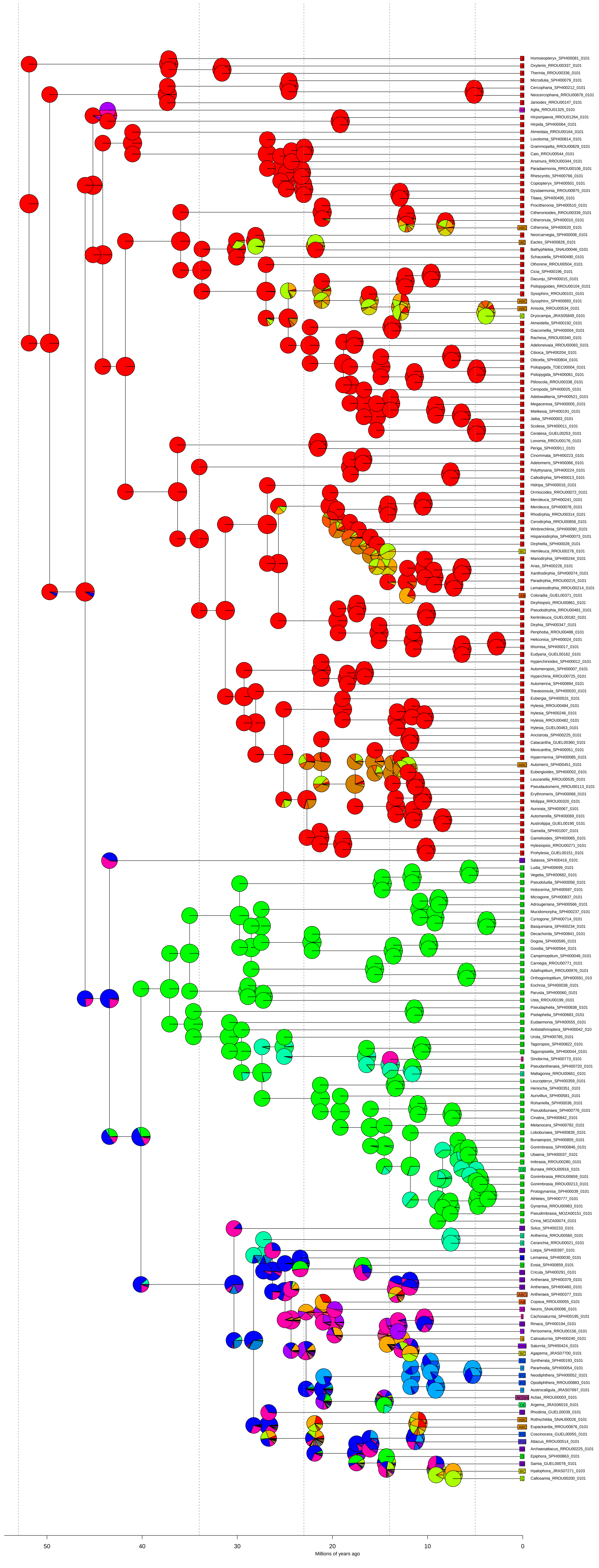
Homoeopteryx_SPHI00081_0101
Oxytenis_RROU00337_0101
Therinia_RROU00336_0101
Microdulia_SPHI00079_0101
Cercophana_SPHI00212_0101
Neocercophana_RROU00878_0101
Janiodes_RROU00147_0101
Aglia_RROU01325_0101
Hirpsinjaevia_RROU01264_0101
Hirpida_SPHI00064_0101
Almeidaia_RROU00164_0101
Loxolomia_SPHI00814_0101
Grammopelta_RROU00829_0101
Caio_RROU00544_0101
Arsenura_RROU00344_0101
Paradaemonia_RROU00106_0101
Rhescyntis_SPHI00766_0101
Copiopteryx_SPHI00501_0101
Dysdaemonia_RROU00875_0101
Titaea_SPHI00495_0101
Procitheronia_SPHI00510_0101
Citheronioides_RROU00339_0101
Citheronula_SPHI00010_0101
Citheronia_SPHI00520_0101
Neorcarnegia_SPHI00008_0101
Eacles_SPHI00826_0101
Bathyphlebia_SNAU00046_0101
Schausiella_SPHI00490_0101
Othorene_RROU00504_0101
Cicia_SPHI00196_0101
Dacunju_SPHI00015_0101
Psilopygoides_RROU00104_0101
Syssphinx_RROU00101_0101
Syssphinx_SPHI00893_0101
Anisota_RROU00534_0101
Dryocampa_JRAS05849_0101
Almeidella_SPHI00192_0101
Giacomellia_SPHI00004_0101
Rachesa_RROU00340_0101
Adeloneivaia_RROU00083_0101
Citioica_SPHI00204_0101
Oiticella_SPHI00804_0101
Psilopygida_TDEC00004_0101
Psilopygida_SPHI00061_0101
Ptiloscola_RROU00338_0101
Ceropoda_SPHI00025_0101
Adelowalkeria_SPHI00521_0101
Megaceresa_SPHI00005_0101
Mielkesia_SPHI00191_0101
Jaiba_SPHI00003_0101
Scolesa_SPHI00011_0101
Ceratesa_GUEL00253_0101
Lonomia_RROU00176_0101
Periga_SPHI00911_0101
Cinommata_SPHI00223_0101
Adetomeris_SPHI00066_0101
Polythysana_SPHI00224_0101
Callodirphia_SPHI00013_0101
Hidripa_SPHI00016_0101
Ormiscodes_RROU00072_0101
Meroleuca_SPHI00241_0101
Meroleuca_SPHI00078_0101
Rhodirphia_RROU00314_0101
Cerodirphia_RROU00858_0101
Winbrechlinia_SPHI00090_0101
Hispaniodirphia_SPHI00073_0101
Dirphiella_SPHI00028_0101
Hemileuca_RROU00276_0101
Manodirphia_SPHI00244_0101
Arias_SPHI00226_0101
Xanthodirphia_SPHI00074_0101
Paradirphia_RROU00215_0101
Lemaireodirphia_RROU00214_0101
Coloradia_GUEL00371_0101
Dirphiopsis_RROU00861_0101
Pseudodirphia_RROU00481_0101
Kentroleuca_GUEL00182_0101
Dirphia_SPHI00347_0101
Periphoba_RROU00488_0101
Heliconisa_SPHI00024_0101
Ithomisa_SPHI00017_0101
Eudyaria_GUEL00162_0101
Hyperchirioides_SPHI00012_0101
Automeropsis_SPHI00007_0101
Hyperchiria_RROU00725_0101
Automerina_SPHI00894_0101
Travassosula_SPHI00020_0101
Eubergia_SPHI00531_0101
Hylesia_RROU00484_0101
Hylesia_SPHI00246_0101
Hylesia_RROU00482_0101
Hylesia_GUEL00463_0101
Ancistrota_SPHI00225_0101
Catacantha_GUEL00360_0101
Mexicantha_SPHI00051_0101
Hypermerina_SPHI00085_0101
Automeris_SPHI00451_0101
Eubergioides_SPHI00002_0101
Leucanella_RROU00535_0101
Pseudautomeris_RROU00113_0101
Erythromeris_SPHI00068_0101
Molippa_RROU00320_0101
Auroraia_SPHI00067_0101
Automerella_SPHI00069_0101
Austrolippa_GUEL00190_0101
Gamelia_SPHI01007_0101
Gamelioides_SPHI00065_0101
Hylesiopsis_RROU00271_0101
Prohylesia_GUEL00151_0101
Salassa_SPHI00416_0101
Ludia_SPHI00699_0101
Vegetia_SPHI00682_0101
Pseudoludia_SPHI00058_0101
Holocerina_SPHI00597_0101
Micragone_SPHI00837_0101
Adrougeriana_SPHI00566_0101
Mucidomorpha_SPHI00237_0101
Cyrtogone_SPHI00714_0101
Basquiniana_SPHI00234_0101
Decachorda_SPHI00841_0101
Dogoia_SPHI00595_0101
Goodia_SPHI00564_0101
Campimoptilum_SPHI00048_0101
Carnegia_RROU00771_0101
Adafroptilum_RROU00976_0101
Orthogonioptilum_SPHI00591_010
Eochroa_SPHI00038_0101
Parusta_SPHI00060_0101
Usta_RROU00199_0101
Pseudaphelia_SPHI00838_0101
Pselaphelia_SPHI00683_0101
Eudaemonia_SPHI00555_0101
Antistathmoptera_SPHI00042_010
Urota_SPHI00785_0101
Tagoropsis_SPHI00822_0101
Tagoropsiella_SPHI00044_0101
Sinobirma_SPHI00773_0101
Pseudantheraea_SPHI00720_0101
Maltagorea_RROU00661_0101
Leucopteryx_SPHI00359_0101
Heniocha_SPHI00351_0101
Aurivillius_SPHI00581_0101
Rohaniella_SPHI00036_0101
Pseudobunaea_SPHI00776_0101
Cinabra_SPHI00842_0101
Melanocera_SPHI00782_0101
Lobobunaea_SPHI00835_0101
Bunaeopsis_SPHI00855_0101
Gonimbrasia_SPHI00846_0101
Ubaena_SPHI00037_0101
Imbrasia_RROU00280_0101
Bunaea_RROU00916_0101
Gonimbrasia_RROU00659_0101
Gonimbrasia_RROU00213_0101
Protogynanisa_SPHI00039_0101
Athletes_SPHI00777_0101
Gynanisa_RROU00983_0101
Pseudimbrasia_MOZA00151_0101
Cirina_MOZA00074_0101
Solus_SPHI00233_0101
Antherina_RROU00560_0101
Ceranchia_RROU00021_0101
Loepa_SPHI00397_0101
Lemaireia_SPHI00030_0101
Eosia_SPHI00859_0101
Cricula_SPHI00291_0101
Antheraea_SPHI00379_0101
Antheraea_SPHI00460_0101
Antheraea_SPHI00377_0101
Copaxa_RROU00055_0101
Neoris_SNAU00096_0101
Cachosaturnia_SPHI00195_0101
Rinaca_SPHI00194_0101
Perisomena_RROU00156_0101
Calosaturnia_SPHI00240_0101
Saturnia_SPHI00424_0101
Agapema_JRAS07700_0101
Syntherata_SPHI00193_0101
Pararhodia_SPHI00054_0101
Neodiphthera_SPHI00052_0101
Opodiphthera_RROU00883_0101
Austrocaligula_JRAS07697_0101
Actias_RROU00003_0101
Argema_JRAS06019_0101
Rhodinia_GUEL00039_0101
Rothschildia_SNAU00026_0101
Eupackardia_RROU00876_0101
Coscinocera_GUEL00055_0101
Attacus_RROU00514_0101
Archaeoattacus_RROU00225_0101
Epiphora_SPHI00863_0101
Samia_GUEL00078_0101
Hyalophora_JRAS07271_0103
Callosamia_RROU00200_0101
50
40
30
20
10
0
Millions of years ago

**Figure S5E. DIVALIKE**

ancstates: global optim, 5 areas max. d=0.0088; e=0.0055; j=0; LnL=-373.23

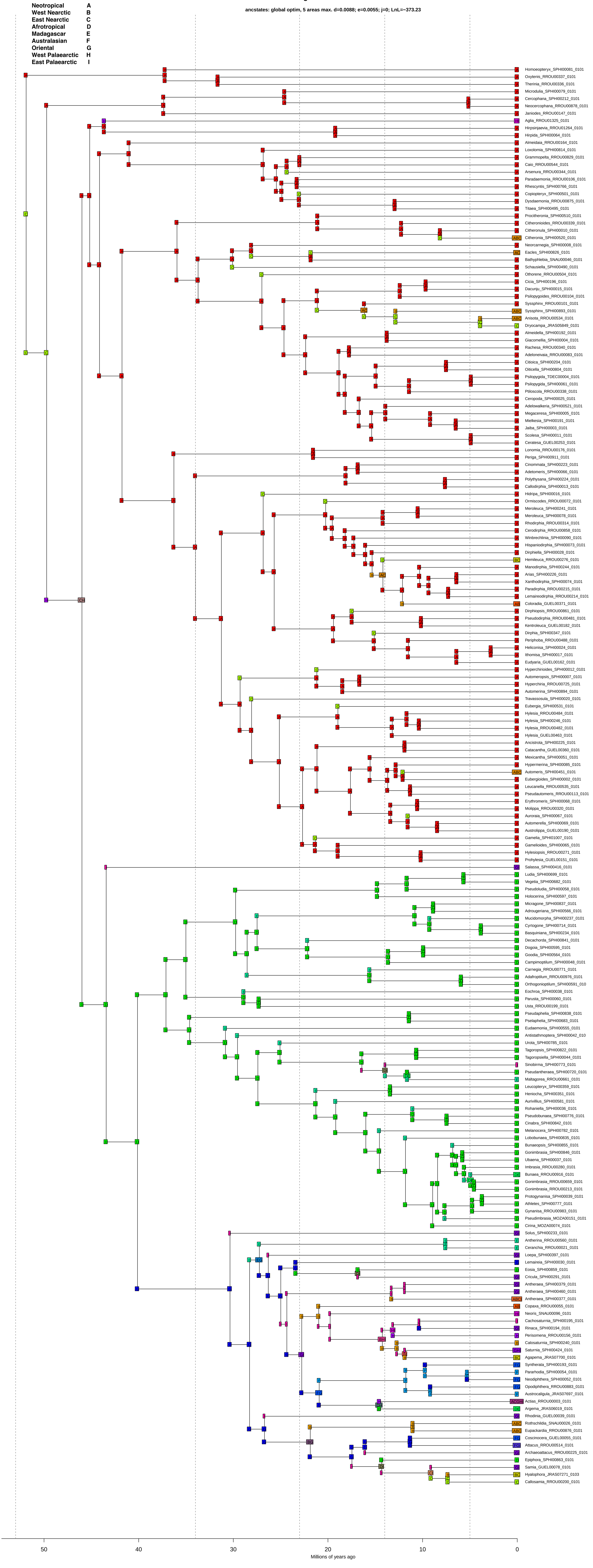

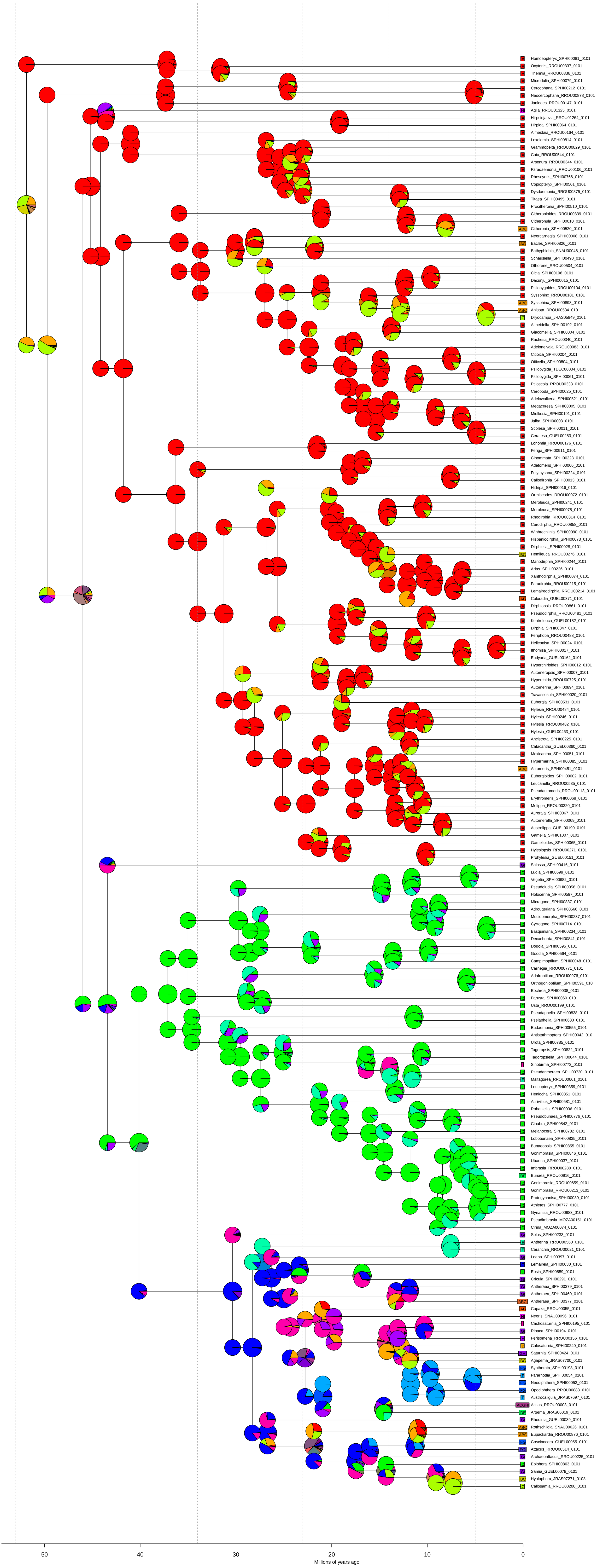
A Homoeopteryx_SPHI00081_0101
A Oxytenis_RROU00337_0101
A Therinia_RROU00336_0101
A Microdulia_SPHI00079_0101
A Cercophana_SPHI00212_0101
A Neocercophana_RROU00878_0101
A Janiodes_RROU00147_0101
HI Aglia_RROU01325_0101
A Hirpsinjaevia_RROU01264_0101
A Hirpida_SPHI00064_0101
A Almeidaia_RROU00164_0101
A Loxolomia_SPHI00814_0101
A Grammopelta_RROU00829_0101
A Caio_RROU00544_0101
A Arsenura_RROU00344_0101
A Paradaemonia_RROU00106_0101
A Rhescyntis_SPHI00766_0101
A Copiopteryx_SPHI00501_0101
A Dysdaemonia_RROU00875_0101
A Titaea_SPHI00495_0101
A Procitheronia_SPHI00510_0101
A Citheronioides_RROU00339_0101
A Citheronula_SPHI00010_0101
ABC Citheronia_SPHI00520_0101
A Neorcarnegia_SPHI00008_0101
AC Eacles_SPHI00826_0101
A Bathyphlebia_SNAU00046_0101
A Schausiella_SPHI00490_0101
A Othorene_RROU00504_0101
A Cicia_SPHI00196_0101
A Dacunju_SPHI00015_0101
A Psilopygoides_RROU00104_0101
A Syssphinx_RROU00101_0101
ABC Syssphinx_SPHI00893_0101
ABC Anisota_RROU00534_0101
C Dryocampa_JRAS05849_0101
A Almeidella_SPHI00192_0101
A Giacomellia_SPHI00004_0101
A Rachesa_RROU00340_0101
A Adeloneivaia_RROU00083_0101
A Citioica_SPHI00204_0101
A Oiticella_SPHI00804_0101
A Psilopygida_TDEC00004_0101
A Psilopygida_SPHI00061_0101
A Ptiloscola_RROU00338_0101
A Ceropoda_SPHI00025_0101
A Adelowalkeria_SPHI00521_0101
A Megaceresa_SPHI00005_0101
A Mielkesia_SPHI00191_0101
A Jaiba_SPHI00003_0101
A Scolesa_SPHI00011_0101
A Ceratesa_GUEL00253_0101
A Lonomia_RROU00176_0101
A Periga_SPHI00911_0101
A Cinommata_SPHI00223_0101
A Adetomeris_SPHI00066_0101
A Polythysana_SPHI00224_0101
A Callodirphia_SPHI00013_0101
A Hidripa_SPHI00016_0101
A Ormiscodes_RROU00072_0101
A Meroleuca_SPHI00241_0101
A Meroleuca_SPHI00078_0101
A Rhodirphia_RROU00314_0101
A Cerodirphia_RROU00858_0101
A Winbrechlinia_SPHI00090_0101
A Hispaniodirphia_SPHI00073_0101
A Dirphiella_SPHI00028_0101
BC Hemileuca_RROU00276_0101
A Manodirphia_SPHI00244_0101
A Arias_SPHI00226_0101
A Xanthodirphia_SPHI00074_0101
A Paradirphia_RROU00215_0101
A Lemaireodirphia_RROU00214_0101
AB Coloradia_GUEL00371_0101
A Dirphiopsis_RROU00861_0101
A Pseudodirphia_RROU00481_0101
A Kentroleuca_GUEL00182_0101
A Dirphia_SPHI00347_0101
A Periphoba_RROU00488_0101
A Heliconisa_SPHI00024_0101
A Ithomisa_SPHI00017_0101
A Eudyaria_GUEL00162_0101
A Hyperchirioides_SPHI00012_0101
A Automeropsis_SPHI00007_0101
A Hyperchiria_RROU00725_0101
A Automerina_SPHI00894_0101
A Travassosula_SPHI00020_0101
A Eubergia_SPHI00531_0101
A Hylesia_RROU00484_0101
A Hylesia_SPHI00246_0101
A Hylesia_RROU00482_0101
A Hylesia_GUEL00463_0101
A Ancistrota_SPHI00225_0101
A Catacantha_GUEL00360_0101
A Mexicantha_SPHI00051_0101
A Hypermerina_SPHI00085_0101
ABC Automeris_SPHI00451_0101
A Eubergioides_SPHI00002_0101
A Leucanella_RROU00535_0101
A Pseudautomeris_RROU00113_0101
A Erythromeris_SPHI00068_0101
A Molippa_RROU00320_0101
A Auroraia_SPHI00067_0101
A Automerella_SPHI00069_0101
A Austrolippa_GUEL00190_0101
A Gamelia_SPHI01007_0101
A Gamelioides_SPHI00065_0101
A Hylesiopsis_RROU00271_0101
A Prohylesia_GUEL00151_0101
GI Salassa_SPHI00416_0101
D Ludia_SPHI00699_0101
D Vegetia_SPHI00682_0101
D Pseudoludia_SPHI00058_0101
D Holocerina_SPHI00597_0101
D Micragone_SPHI00837_0101
D Adrougeriana_SPHI00566_0101
D Mucidomorpha_SPHI00237_0101
D Cyrtogone_SPHI00714_0101
D Basquiniana_SPHI00234_0101
D Decachorda_SPHI00841_0101
D Dogoia_SPHI00595_0101
D Goodia_SPHI00564_0101
D Campimoptilum_SPHI00048_0101
D Carnegia_RROU00771_0101
D Adafroptilum_RROU00976_0101
D Orthogonioptilum_SPHI00591_010
D Eochroa_SPHI00038_0101
D Parusta_SPHI00060_0101
D Usta_RROU00199_0101
D Pseudaphelia_SPHI00838_0101
D Pselaphelia_SPHI00683_0101
D Eudaemonia_SPHI00555_0101
D Antistathmoptera_SPHI00042_010
D Urota_SPHI00785_0101
D Tagoropsis_SPHI00822_0101
D Tagoropsiella_SPHI00044_0101
I Sinobirma_SPHI00773_0101
D Pseudantheraea_SPHI00720_0101
E Maltagorea_RROU00661_0101
D Leucopteryx_SPHI00359_0101
D Heniocha_SPHI00351_0101
D Aurivillius_SPHI00581_0101
D Rohaniella_SPHI00036_0101
D Pseudobunaea_SPHI00776_0101
D Cinabra_SPHI00842_0101
D Melanocera_SPHI00782_0101
D Lobobunaea_SPHI00835_0101
D Bunaeopsis_SPHI00855_0101
D Gonimbrasia_SPHI00846_0101
D Ubaena_SPHI00037_0101
D Imbrasia_RROU00280_0101
DE Bunaea_RROU00916_0101
D Gonimbrasia_RROU00659_0101
D Gonimbrasia_RROU00213_0101
D Protogynanisa_SPHI00039_0101
D Athletes_SPHI00777_0101
D Gynanisa_RROU00983_0101
D Pseudimbrasia_MOZA00151_0101
D Cirina_MOZA00074_0101
GI Solus_SPHI00233_0101
E Antherina_RROU00560_0101
E Ceranchia_RROU00021_0101
GI Loepa_SPHI00397_0101
G Lemaireia_SPHI00030_0101
D Eosia_SPHI00859_0101
GI Cricula_SPHI00291_0101
GI Antheraea_SPHI00379_0101
GI Antheraea_SPHI00460_0101
ABCI Antheraea_SPHI00377_0101
AB Copaxa_RROU00055_0101
HI Neoris_SNAU00096_0101
I Cachosaturnia_SPHI00195_0101
GI Rinaca_SPHI00194_0101
H Perisomena_RROU00156_0101
B Calosaturnia_SPHI00240_0101
GHI Saturnia_SPHI00424_0101
BC Agapema_JRAS07700_0101
FG Syntherata_SPHI00193_0101
F Pararhodia_SPHI00054_0101
FG Neodiphthera_SPHI00052_0101
FG Opodiphthera_RROU00883_0101
F Austrocaligula_JRAS07697_0101
ACGHI Actias_RROU00003_0101
DE Argema_JRAS06019_0101
GI Rhodinia_GUEL00039_0101
ABC Rothschildia_SNAU00026_0101
ABC Eupackardia_RROU00876_0101
FG Coscinocera_GUEL00055_0101
FGI Attacus_RROU00514_0101
GI Archaeoattacus_RROU00225_0101
D Epiphora_SPHI00863_0101
GI Samia_GUEL00078_0101
BC Hyalophora_JRAS07271_0103
C Callosamia_RROU00200_0101
50
40
30
20
10
0
Millions of years ago

**Figure S5F. DIVALIKE+J**

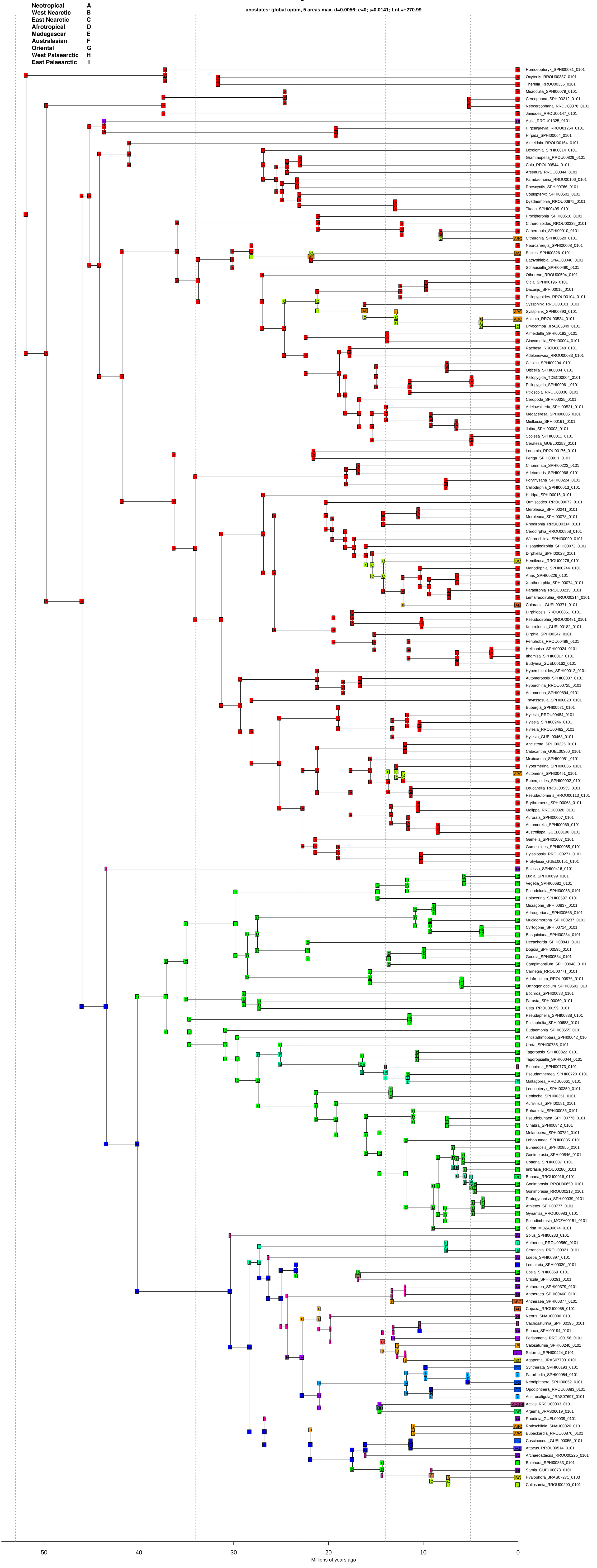

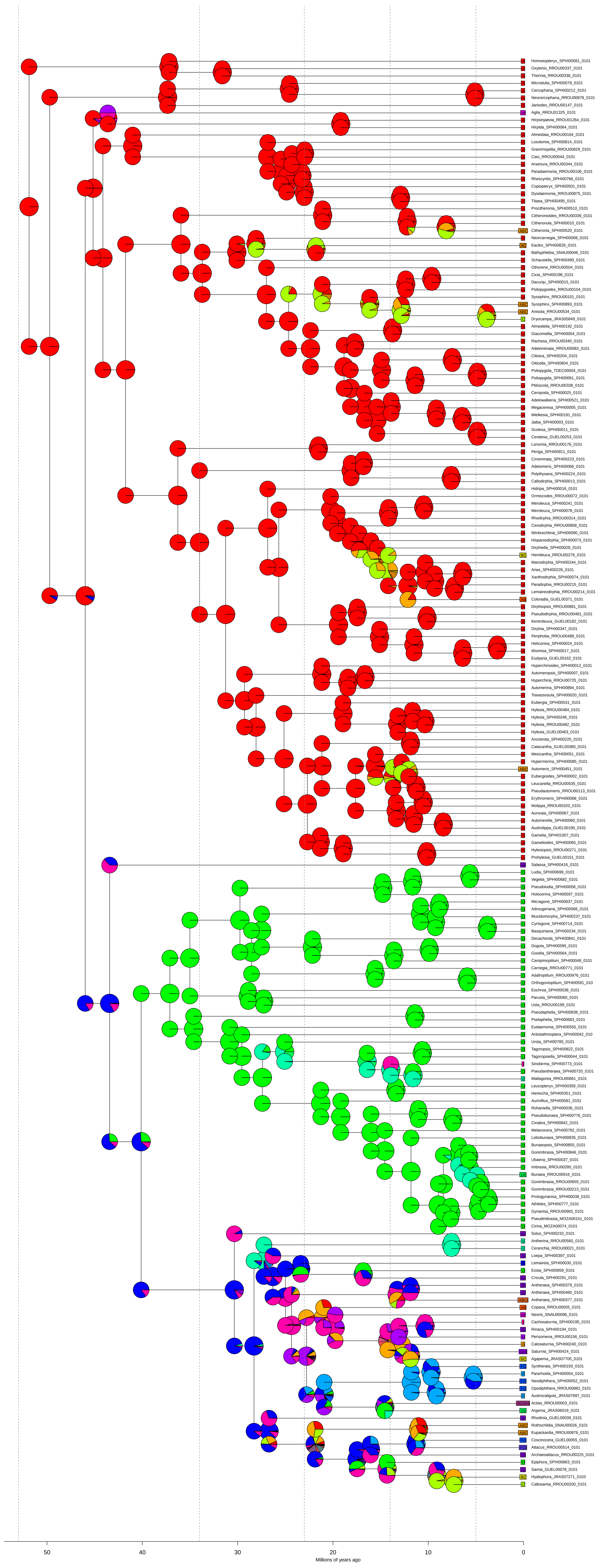
Homoeopteryx_SPHI00081_0101
Oxytenis_RROU00337_0101
Therinia_RROU00336_0101
Microdulia_SPHI00079_0101
Cercophana_SPHI00212_0101
Neocercophana_RROU00878_0101
Janiodes_RROU00147_0101
Aglia_RROU01325_0101
Hirpsinjaevia_RROU01264_0101
Hirpida_SPHI00064_0101
Almeidaia_RROU00164_0101
Loxolomia_SPHI00814_0101
Grammopelta_RROU00829_0101
Caio_RROU00544_0101
Arsenura_RROU00344_0101
Paradaemonia_RROU00106_0101
Rhescyntis_SPHI00766_0101
Copiopteryx_SPHI00501_0101
Dysdaemonia_RROU00875_0101
Titaea_SPHI00495_0101
Procitheronia_SPHI00510_0101
Citheronioides_RROU00339_0101
Citheronula_SPHI00010_0101
Citheronia_SPHI00520_0101
Neorcarnegia_SPHI00008_0101
Eacles_SPHI00826_0101
Bathyphlebia_SNAU00046_0101
Schausiella_SPHI00490_0101
Othorene_RROU00504_0101
Cicia_SPHI00196_0101
Dacunju_SPHI00015_0101
Psilopygoides_RROU00104_0101
Syssphinx_RROU00101_0101
Syssphinx_SPHI00893_0101
Anisota_RROU00534_0101
Dryocampa_JRAS05849_0101
Almeidella_SPHI00192_0101
Giacomellia_SPHI00004_0101
Rachesa_RROU00340_0101
Adeloneivaia_RROU00083_0101
Citioica_SPHI00204_0101
Oiticella_SPHI00804_0101
Psilopygida_TDEC00004_0101
Psilopygida_SPHI00061_0101
Ptiloscola_RROU00338_0101
Ceropoda_SPHI00025_0101
Adelowalkeria_SPHI00521_0101
Megaceresa_SPHI00005_0101
Mielkesia_SPHI00191_0101
Jaiba_SPHI00003_0101
Scolesa_SPHI00011_0101
Ceratesa_GUEL00253_0101
Lonomia_RROU00176_0101
Periga_SPHI00911_0101
Cinommata_SPHI00223_0101
Adetomeris_SPHI00066_0101
Polythysana_SPHI00224_0101
Callodirphia_SPHI00013_0101
Hidripa_SPHI00016_0101
Ormiscodes_RROU00072_0101
Meroleuca_SPHI00241_0101
Meroleuca_SPHI00078_0101
Rhodirphia_RROU00314_0101
Cerodirphia_RROU00858_0101
Winbrechlinia_SPHI00090_0101
Hispaniodirphia_SPHI00073_0101
Dirphiella_SPHI00028_0101
Hemileuca_RROU00276_0101
Manodirphia_SPHI00244_0101
Arias_SPHI00226_0101
Xanthodirphia_SPHI00074_0101
Paradirphia_RROU00215_0101
Lemaireodirphia_RROU00214_0101
Coloradia_GUEL00371_0101
Dirphiopsis_RROU00861_0101
Pseudodirphia_RROU00481_0101
Kentroleuca_GUEL00182_0101
Dirphia_SPHI00347_0101
Periphoba_RROU00488_0101
Heliconisa_SPHI00024_0101
Ithomisa_SPHI00017_0101
Eudyaria_GUEL00162_0101
Hyperchirioides_SPHI00012_0101
Automeropsis_SPHI00007_0101
Hyperchiria_RROU00725_0101
Automerina_SPHI00894_0101
Travassosula_SPHI00020_0101
Eubergia_SPHI00531_0101
Hylesia_RROU00484_0101
Hylesia_SPHI00246_0101
Hylesia_RROU00482_0101
Hylesia_GUEL00463_0101
Ancistrota_SPHI00225_0101
Catacantha_GUEL00360_0101
Mexicantha_SPHI00051_0101
Hypermerina_SPHI00085_0101
Automeris_SPHI00451_0101
Eubergioides_SPHI00002_0101
Leucanella_RROU00535_0101
Pseudautomeris_RROU00113_0101
Erythromeris_SPHI00068_0101
Molippa_RROU00320_0101
Auroraia_SPHI00067_0101
Automerella_SPHI00069_0101
Austrolippa_GUEL00190_0101
Gamelia_SPHI01007_0101
Gamelioides_SPHI00065_0101
Hylesiopsis_RROU00271_0101
Prohylesia_GUEL00151_0101
Salassa_SPHI00416_0101
Ludia_SPHI00699_0101
Vegetia_SPHI00682_0101
Pseudoludia_SPHI00058_0101
Holocerina_SPHI00597_0101
Micragone_SPHI00837_0101
Adrougeriana_SPHI00566_0101
Mucidomorpha_SPHI00237_0101
Cyrtogone_SPHI00714_0101
Basquiniana_SPHI00234_0101
Decachorda_SPHI00841_0101
Dogoia_SPHI00595_0101
Goodia_SPHI00564_0101
Campimoptilum_SPHI00048_0101
Carnegia_RROU00771_0101
Adafroptilum_RROU00976_0101
Orthogonioptilum_SPHI00591_010
Eochroa_SPHI00038_0101
Parusta_SPHI00060_0101
Usta_RROU00199_0101
Pseudaphelia_SPHI00838_0101
Pselaphelia_SPHI00683_0101
Eudaemonia_SPHI00555_0101
Antistathmoptera_SPHI00042_010
Urota_SPHI00785_0101
Tagoropsis_SPHI00822_0101
Tagoropsiella_SPHI00044_0101
Sinobirma_SPHI00773_0101
Pseudantheraea_SPHI00720_0101
Maltagorea_RROU00661_0101
Leucopteryx_SPHI00359_0101
Heniocha_SPHI00351_0101
Aurivillius_SPHI00581_0101
Rohaniella_SPHI00036_0101
Pseudobunaea_SPHI00776_0101
Cinabra_SPHI00842_0101
Melanocera_SPHI00782_0101
Lobobunaea_SPHI00835_0101
Bunaeopsis_SPHI00855_0101
Gonimbrasia_SPHI00846_0101
Ubaena_SPHI00037_0101
Imbrasia_RROU00280_0101
Bunaea_RROU00916_0101
Gonimbrasia_RROU00659_0101
Gonimbrasia_RROU00213_0101
Protogynanisa_SPHI00039_0101
Athletes_SPHI00777_0101
Gynanisa_RROU00983_0101
Pseudimbrasia_MOZA00151_0101
Cirina_MOZA00074_0101
Solus_SPHI00233_0101
Antherina_RROU00560_0101
Ceranchia_RROU00021_0101
Loepa_SPHI00397_0101
Lemaireia_SPHI00030_0101
Eosia_SPHI00859_0101
Cricula_SPHI00291_0101
Antheraea_SPHI00379_0101
Antheraea_SPHI00460_0101
Antheraea_SPHI00377_0101
Copaxa_RROU00055_0101
Neoris_SNAU00096_0101
Cachosaturnia_SPHI00195_0101
Rinaca_SPHI00194_0101
Perisomena_RROU00156_0101
Calosaturnia_SPHI00240_0101
Saturnia_SPHI00424_0101
Agapema_JRAS07700_0101
Syntherata_SPHI00193_0101
Pararhodia_SPHI00054_0101
Neodiphthera_SPHI00052_0101
Opodiphthera_RROU00883_0101
Austrocaligula_JRAS07697_0101
Actias_RROU00003_0101
Argema_JRAS06019_0101
Rhodinia_GUEL00039_0101
Rothschildia_SNAU00026_0101
Eupackardia_RROU00876_0101
Coscinocera_GUEL00055_0101
Attacus_RROU00514_0101
Archaeoattacus_RROU00225_0101
Epiphora_SPHI00863_0101
Samia_GUEL00078_0101
Hyalophora_JRAS07271_0103
Callosamia_RROU00200_0101
50
40
30
20
10
0
Millions of years ago
